# Cerebellar granular neuron progenitors exit their germinative niche via Barhl1 mediated silencing of T-Cell Factor transcriptional activity

**DOI:** 10.1101/2023.05.25.542248

**Authors:** Johnny Bou-Rouphael, Mohammed Doulazmi, Alexis Eschstruth, Asna Abdou, Béatrice C. Durand

## Abstract

T-Cell Factors (TCFs) are the main transcriptional effectors of Wnt/β-catenin signaling. TCF responsiveness is a hallmark of self-renewal in mouse embryonic, and adult, neural stem cells (NSC). However, *in vivo* contribution(s) of TCF activities in long-lived NSCs are poorly understood. Granule neuron progenitors (GNP) in the upper rhombic lip (URL) are long-lived NSCs which express *Atoh1* and generate cerebellar granule neurons. Using functional and transcriptomic approaches in amphibian, we demonstrate that TCFs are active in the URL, and are strictly necessary for the emergence and maintenance of the GNP germinative zone. We identify BarH-like 1 (Barhl1), a direct target of Atoh1, as a gate keeper for GNP exit from the URL, through silencing of TCF transcriptional activity. Our transcriptomic and *in silico* analysis identifies Barhl1/TCF URL target genes, and confirms our functional data. Our study provides *in vivo* evidence that inhibition of TCF repressive activity is necessary for maintenance of the URL, a long-lived neural germinative niche.

## INTRODUCTION

The Wnt/β-catenin cell-to-cell signaling pathway coordinates development and is one of the most conserved in the animal kingdom. The large majority of Wnt/β-catenin transcriptional targets are regulated by T-Cell Factor/Lymphoid Enhancer-binding Factor (TCF/LEF) transcription factors (TF) ^1, 2,reviewed^ ^in^ ^3^. Investigation of the developmental fate of Wnt/β-catenin- responsive cells in embryonic and postnatal mouse brains reveals that long-lived NSCs retain neuroepithelial properties, and Wnt/β-catenin responsiveness throughout development ^4^. In the adult mouse ventricular-subventricular zone of the lateral ventricles, WNT signaling promotes both NSC self-renewal and neural progenitor cell proliferation, while TCF/LEF activity is detected in deeply quiescent NSC cells ^4–7^ reviewed in ^8^. Moreover, hippocampal quiescent NSC and progenitors in culture exhibit enhanced TCF/LEF1 driven transcription ^9^. Taken together, these observations suggest contribution(s) of Wnt driven TCF transcriptional activity in adult NSC biology. However, currently, our understanding of such activities in long- lived NSC remains surprisingly fragmented 8,reviewed in ^10–12^.

A crucial component of the central nervous system (CNS) in all jawed vertebrates is the cerebellum, involved in executing motor functions as well as participating in higher cognitive processes such as decision-making, emotional and social behaviour, and expectation of reward ^13–15^. The cerebellum has two major stem cell niches: the ventricular zone (VZ) adjacent to the fourth ventricle, which produces all cerebellar GABAergic inhibitory neurons^16–18;^ and the upper rhombic lip (URL) which is the origin of glutamatergic excitatory neurons, derived from Atonal homologue 1 *(Atoh1)-*expressing progenitors. The URL gives birth first to the deep cerebellar nuclei (DCN), followed by the unipolar brush cells (UBC) and the granular neuron progenitors (GNP) that in turn produce granule neurons, the predominant neuronal population in the entire CNS reviewed in ^19–21^.

While the VZ appears to be TCF inactive, positive TCF transcriptional activity has been documented in the URL of mice, human and *Xenopus* species ^22–25^. Moreover, *in vitro* and *in vivo* studies in mice show that, in contrast to what is observed in NSC and progenitors of the developing CNS, or in the VZ ^26, 27^, constitutive activation of β-catenin in *Atoh1+* URL cells does not promote their proliferation ^26, 28–30^. Taken together, these data indicate that TCF-mediated transcription probably contributes to the URL biology, but its role(s) in this germinative area remains undefined. In addition, they highlight the presence of TCF developmental regulators within this germinative area that have not yet been identified.

The GNP developmental path is marked by expression of specific TF, including ATOH1 which is indispensable for maintaining GNP in an immature state ^31^. In amniotes, *Atoh1* expression initiates in the RL, is maintained in the EGL during GNP proliferation, and is lost in differentiated GN that start expressing the basic Helix Loop Helix (bHLH) neurogenic differentiation factor 1 (*Neurod1*). In addition, the Paired box protein 6 (*Pax6*) and BarH-like 1 homeobox protein (*Barhl1*) expressions are markers of GNP commitment ^25, 32–36^.

Amphibians show a marked development of the cerebellum that displays morphological features resembling those found in higher vertebrates ^37, 38^ reviewed in ^39, 40^. The few studies performed in the amphibian *Xenopus* pre-metamorphosis reveal the presence of an EGL-like structure that is unique compared with other anamniotes. However, unlike in higher vertebrates, the *Xenopus* EGL lacks any cells that undergo proliferation ^41^. These studies also indicate that the developmental processes that lead to the formation of GN, specifically the presence of an *Atoh1*-expressing URL and EGL, and the expression of *Neurod1* are close to those described in higher vertebrates ^41, 42^. Of Note, the amphibian URL maintains itself until post-metamorphic stages ^38, 41^. However, early developmental events leading to GNP induction and EGL formation have not been described in amphibians.

ATOH1 directly induces its own expression as well as the expression of the two homeodomain (HD)-containing TF, BARHL1 and BARHL2, which are mammalian homologues of the *Drosophila* Bar-class HD, BarH1 and BarH2 ^43–46^ reviewed in ^3, 47^. In mice, *Barhl1* cerebellar expression is detected in committed GNP, and persists in the EGL ^33, 48^. In the developing cerebellum, BARHL1 participates in the generation of the EGL ^45^, and is one of the major TF that regulate the radial migration of GNP in a mechanism involving Neurotrophin3 (NT3) ^49^. Furthermore, an impairment in GN survival, and an attenuated cerebellar foliation, are observed in *Barhl1-/-* mice ^50^. On the other hand, we established that BARHL2 dramatically enhances the transcriptional repressor activity of TCF, and prevents β-catenin driven activation of TCF target genes ^51, 52^. Immunoprecipitation assays reveal physical interaction between BARHL2 and the two transcriptional repressors of Wnt target genes TCF7L1 and Groucho/Transducin-like Enhancer of Split (Gro/TLE) ^52^. However, the role of BARHL2 in cerebellar development has not been investigated, and whether BARHL1 similarly interacts with, and regulates, TCF transcriptional activity is unknown.

Here, we investigated the role of both Tcf transcriptional activity, and Barhl1, in GNP development using *Xenopus* as a model system. We establish that markers of GN early progenitors, EGL, commitment, and differentiation, are conserved in *Xenopus* compared to amniotes. Using gain and loss of function experiments (GOF/LOF), immunoprecipitation, and a *X. tropicalis* Wnt reporter transgenic line, we demonstrate that Tcf-mediated transcriptional activation is strictly necessary for both the emergence and maintenance of the cerebellar URL, and furthermore demonstrate that, in this germinative area, Barhl1 is the main repressor of Tcf transcriptional activity. Barhl1 LOF, through Morpholino (MO)-mediated depletion or Crispr/Cas9 gene knock-out, dramatically increases Tcf transcriptional activity in the URL, leading to a major enlargement of the URL, and significant delays in GNP differentiation. Using a transcriptomic approach, we confirm our functional assays and identify direct and indirect target genes repressed by Barhl1 in the cerebellar URL. Indeed, Barhl1 depletion induces an upregulation of TF involved in maintenance of neural stem/progenitor properties, an enhancement of both Wnt/Tcf and Notch signaling activities, associated with a down regulation of genes involved in inhibiting proliferation, and promoting neural differentiation. Together with *in silico* analysis of Barhl1 target genes regulatory regions, our study confirms that in the amphibian, Barhl1 drives URL stem/progenitor cells out of their germinative niche, towards commitment and differentiation, by repressing TCF transcriptional activity.

## RESULTS

### Spatial and temporal expression of key markers of GNP development are conserved in *Xenopus* compared to higher vertebrate

We performed *in situ* hybridization (ISH) on *X. laevis* tadpoles through pre-metamorphic froglets (stages 35 to 50) and assessed the expression patterns of genes known to be involved in the development of *Atoh1* lineage in rodents, focusing on GN. By using *atoh1* to label the URL and EGL, *pax6* and *barhl1* to mark GNP commitment, and *neurod1* to mark post-mitotic GN, we established a fate map that outlines the developmental progression of *atoh1* lineage cells within rhombomere 1 (R1), located caudal to the midbrain-hindbrain boundary (MHB). The proliferation marker *n-myc* helps define the posterior boundary of R1 (Fig. 1A; Fig. S1B).

**Figure 1:**
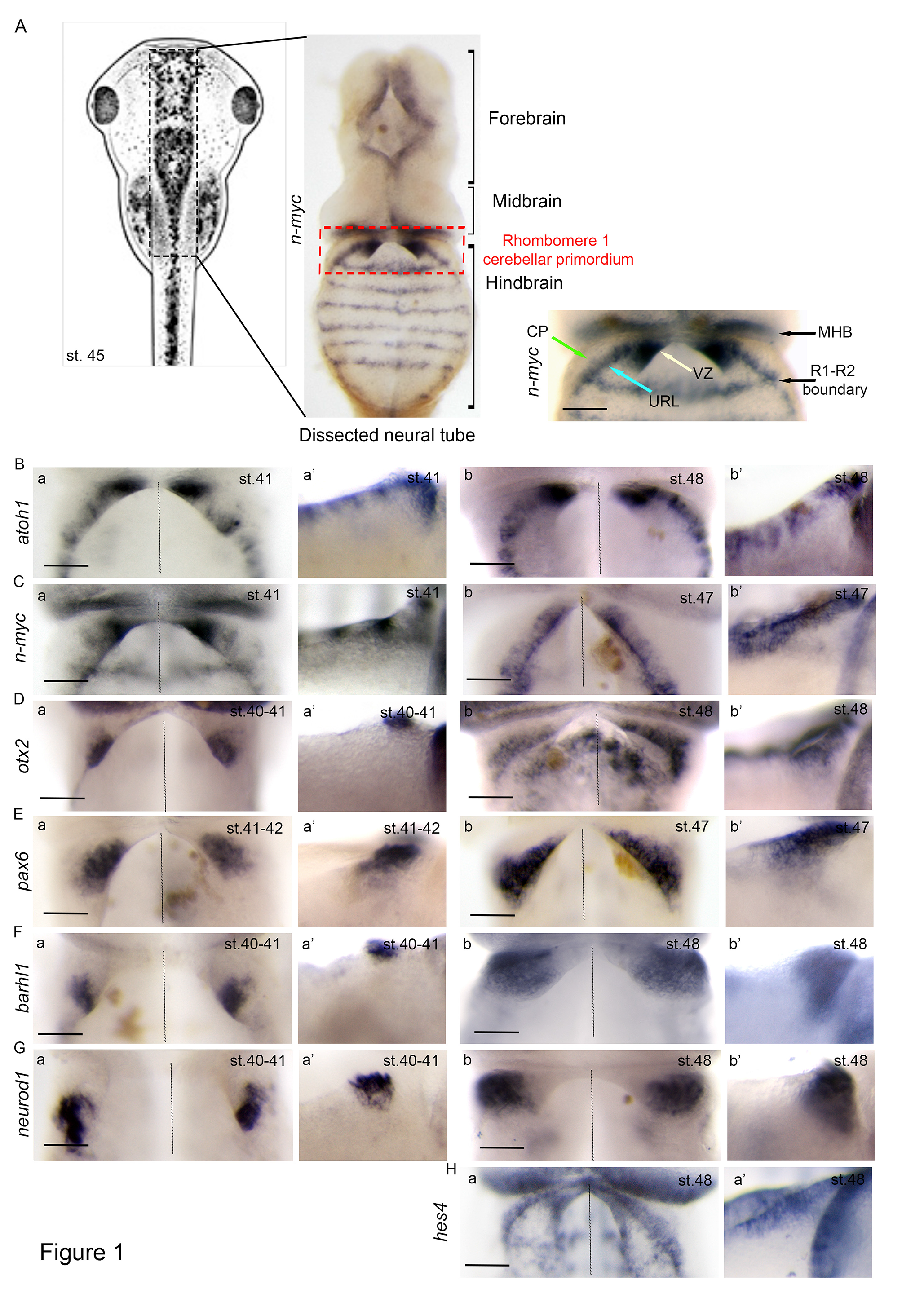
Temporal and spatial expression pattern of genes involved in granule neuron progenitors’ (GNP) development (A) Neural tube dissection and analysis. Shown on the left is a representation of stage (st.) 45 *X. laevis* embryo. Following ISH, neural tubes are dissected as shown on the middle (entire neural tube) and right (a focus on the rhombomere 1 (R1)) panels. The proliferation marker *nmyc* is expressed in the upper rhombic lip (URL) (blue arrow), the ventricular zone VZ (white arrow). Red dotted lines delineate rhombomere 1 (R1) located caudal to the midbrain-hindbrain boundary (MHB). *nmyc* marks proliferating progenitors at the boundary between R1 and R2 and is used as a marker of cerebellar primordium’s caudal limit. (B) ISH analysis of GNP markers in *X. laevis* embryos at the indicated Nieuwkoop and Faber stages. Shown are dorsal and lateral views of the R1. From st. 41 to st. 48 stem/progenitor markers *atoh1* (Ba-b’), *nmyc* (Ca-b’), and *hes4* (Ha, a’) display a strong expression in the URL and in the EGL. *hes4 and nmyc* are also detected in the VZ. (Da, a’) *otx2* expression is first detected in caudal EGL and becomes restricted to the cerebellar plate (CP) (green arrow) at st. 48 (Db, b’). At st. 41 committed GNP markers *pax6* (Ea, a’) and *barhl1* (Fa, a’), together with the differentiation marker *neurod1* (Ga, a’) are detected in the caudal EGL and the cerebellar plate. As development proceeds, transcripts for these markers are detected in the CP and their expression significantly increases in this area (E-G, b, b’). Fully differentiated GNs settling in the internal granule layer (IGL) are stained with *neurod1* as observed in lateral views of st. 48 *X. laevis* embryos. The CP is devoid of *atoh1, hes4,* and *nmyc* expressions. CP: cerebellar plate; VZ: ventricular zone; URL: upper rhombic lip; EGL: external granule layer; R: rhombomere; MHB: midbrain-hindbrain boundary. Scale bar 150μm.

We found that *pax6* expression begins at stage 38 (Fig. S1C), while *barhl1* expression begins at stage 39-40 (Fig. 1F), which we used as a landmark of GNP induction. From stage 38 to stage 48, *atoh1* is expressed in the URL and in a layer of 3 to 4 cells bordering the URL, which we consider to be the EGL (Fig. 1B; Fig. S.1A). Although the *n-myc* expression pattern is similar to that of *atoh1* in the URL, *n-myc* is also strongly detected in the VZ, in agreement with its expression in proliferating cells (Fig. 1C, Fig. S1B).

We investigated the dynamics of *pax6*, *barhl1*, and *neurod1* expressions within R1 from stage 38 to stage 48 (Fig. 1E-G; Fig. S1C-E). All markers are present in the cerebellar primordium and are first detected in the caudal region of the EGL. During this developmental period, cells *expressing barhl1* and *neurod1* migrate from an external layer partially covering the cerebellar plate towards the inner cerebellar tissue, where they undergo their final differentiation (Fig. 1F, G, Fig. S1D, E). Furthermore, while *Orthodenticle Homeobox 2 (otx2)* expression is typically limited to the posterior lobes of the EGL in amniotes, we detected *otx2* expression in the caudal EGL at stage 39, which was subsequently restricted to the cerebellar plate (Fig. 1D). At stage 48, *Hairy and enhancer of split-4 (hes4)*, a known marker of Notch-active cells and of stemness in *Xenopus* ^53, 54^, strongly labels both the VZ and the EGL at stage 48 (Fig. 1H). *hes4* is not expressed in mice, while its expression is found in *Xenopus* and in Human. Additionally, we did not observe expression of *barhl2* in the cerebellar anlage at the analysed developmental stages (Fig. S1F).

These observations indicate striking similarities in GNP development between *Xenopus* and amniotes. Specifically, the expression pattern of genes involved in induction and specification of GNP, including *atoh1, n-myc, pax6, barhl1, otx2, neurod1,* is conserved. As previously reported ^41^, we detect the presence of an EGL along the R1 antero-posterior axis marked by *atoh1* and *hes4*. Our observations also indicate a gradient in GNP differentiation, initiated in the caudal EGL at stage 38 and progressing to the rostral part up to stage 50. Starting at stage 50, we report changes in the shape of the URL. Thus, we focused our analysis on cerebellar anlage development between stage 38 and stage 48.

### In the cerebellar primordium, constitutive Tcf inhibition and Barhl1 overexpression produce similar developmental defects in *atoh1* expression, URL induction and GNP early commitment/differentiation

We focused our study on the role of Tcf and Barhl1 in URL establishment and maintenance, during the time window where GNP are produced. We used *tcf7l1-Δβcat-GR*, a previously described inducible form of *tcf7l1* which lacks its β-catenin binding domain, and thus acts as a dominant negative and constitutive inhibitor of Tcf transcriptional activity ^55^ (Fig. S2). Development of the URL and GNP was investigated using either *atoh1*, or *pax6, barhl1* and *neurod1* as respectively URL/EGL, and GNP commitment /differentiation markers.

At a high dose, *tcf7l1-Δβcat-GR* overexpression induced a dramatic reduction in the size of the URL, associated with the disappearance of the expression of its key marker *atoh1*. Of note, this effect is restricted to the R1 (Fig. 2Aa-a”, c). At a lower dose, *tcf7l1-Δβcat-GR* overexpression induced a decrease in *atoh1* expression (Fig. 2Ab, c). This decrease is associated with both an increase of expression, and a rostral shift, observed with the three commitment/differentiation markers *pax6*, *barhl1*, and *neurod1* within the R1 (Fig.2Ba-c). *tcf7l1-Δβcat-GR* effects on *atoh1, pax6, barhl1* and *neurod1* expression were quantified (Fig.2Ac-Bd).

**Figure 2:**
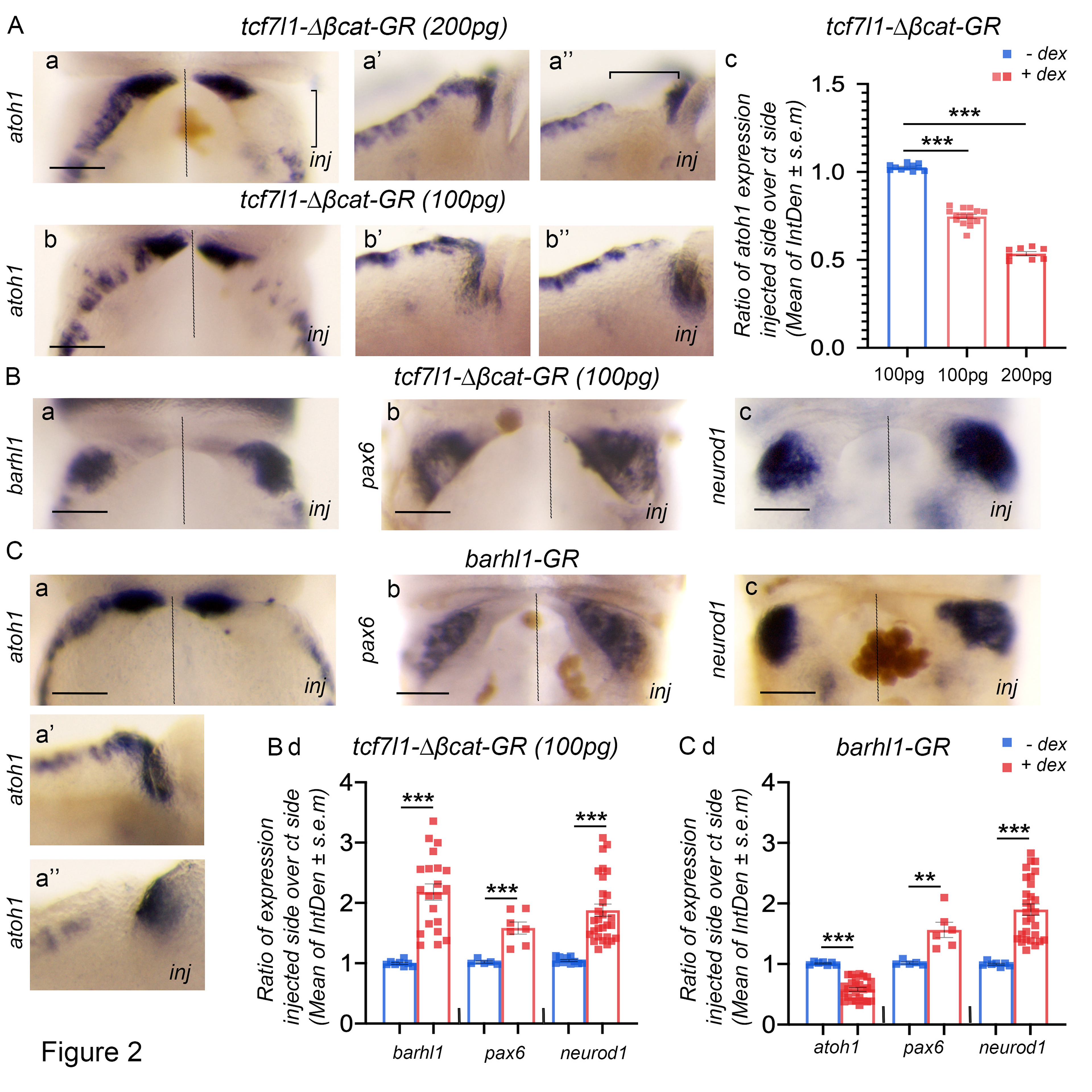
Tcf activity is required for the induction of the URL and its inhibition by Barhl1 is necessary for the proper progression of GNPs development. (A) Overexpression of *tcf7l1* inhibits/abolishes *atoh1* expression in a dose dependent manner. ISH analysis of *atoh1* expression in the rhombomere 1 (R1) showing dorsal views (a, b) and lateral views of control sides (a’, b’) and injected sides (a”, b”) of stage 45 *X. laevis* embryos unilaterally injected with 200pg (a, a’, a”) and 100pg (b, b’, b”) of inducible *tcf7l1-Δβcat-GR.* The non-injected side is an internal control. (B) Forced expression of *tcf7l1* increased GNP differentiation. ISH analysis of the commitment/differentiation markers *barhl1, pax6* and *neurod1* (a-c) in stage 45 *X. laevis* embryos. (C) *barhl1* overexpression phenocopies defects of *tcf7l1* overexpression. Dorsal views showing *atoh1, barhl1* and *neurod1* (a, b, and c respectively) expressions in the R1 primordium of stage 45 *X. laevis* embryos injected with *mBarhl1GR* (200pg). Lateral views of *atoh1* expression in control side (a’) and injected side (a”) are shown. Integrated densities *(IntDen)* of markers’ expressions were measured. Ratio of markers expression in injected side over control side is represented (Ac; Bd; Cd) and indicated as mean ± s.e.m. Dotted lines separate injected and control sides. Scale bar 150μm. Square brackets delineate R1. Dex: dexamethasone; *inj*: injected side. Statistical analysis C: One-way ANOVAone way anova (F(2,31)=437.5; p < 0.001) followed by post hoc Tukey test. Bd, Cd: student’s t-test. Data are presented as means ± SEM.** p ≤ 0.01; *** p ≤ 0.001; **** p ≤ 0.0001.

In amniotes, BARHL1 is a direct target of ATOH1 ^45, 46^. We next asked whether Barhl1 overexpression impacts early development of the URL and GNP (Fig. S2). We observed that *barhl1* overexpression phenocopies that of Tcf inhibition. We observed a strong decrease in *atoh1* transcripts levels within the URL (Fig. 2Ca-a”, d). Loss of *atoh1* and the URL is associated with an increase of *pax6* and *neurod1* expression, and a concomitant rostral shift in both markers’ expression within R1 (Fig. 2Cb-d).

Our data indicate that Tcf transcriptional activity is strictly necessary for the expression of *atoh1* within the URL, and that inhibiting Tcf activity leads to accelerated GNP differentiation. Similarly, overexpression of Barhl1 in the cerebellar primordium results in URL induction defects, associated with premature GNP differentiation.

### In the cerebellar URL, inhibition of Barhl1 maintains GNP in an early progenitor state

To decrease Barhl1 activity within the cerebellar anlage, we designed two morpholinos (MO), *MObarhl1-1* and *MObarhl1-2*, specifically targeting *Xenopus barhl1* mRNA (Fig. S2; S3; Methods). We investigated whether, andby what mechanism(s), Barhl1 Knock-Down (KD) affects the development of the URL, the EGL, and/or GNP development. At stage 42, 45, and 48, depletion of Barhl1 induced an increase in *atoh1* expression. We observed *atoh1* expressing cells spreading across the surface of the cerebellar plate (Fig. 3Aa-a”, Ba-a”; Fig. S3B, C, D). This ectopic expansion within the URL and to the EGL is associated with a major decrease in *pax6* expression (Fig. 3Ab, Bb; Fig. S3B, D). Both MO induced the same phenotype, which was quantified (Fig. 3C). At stage 42, the increase in *atoh1* expression in Barhl1-KD embryos is corroborated by a similar increase in *n-myc* expression (Fig. S3C). We further tested the ability of *mbarhl1*-GR to rescue the Barhl1-KD phenotype. *mbarhl1*-GR was co-injected with *MObarhl1-1,* and *neurod1* was used as a marker of GNP differentiation. *MObarhl1-1* and *MObarhl1-2* induced a strong decrease in *neurod1* expression, which was rescued by co-injection of *mbarhl1*-GR (Fig. 3D).

**Figure 3:**
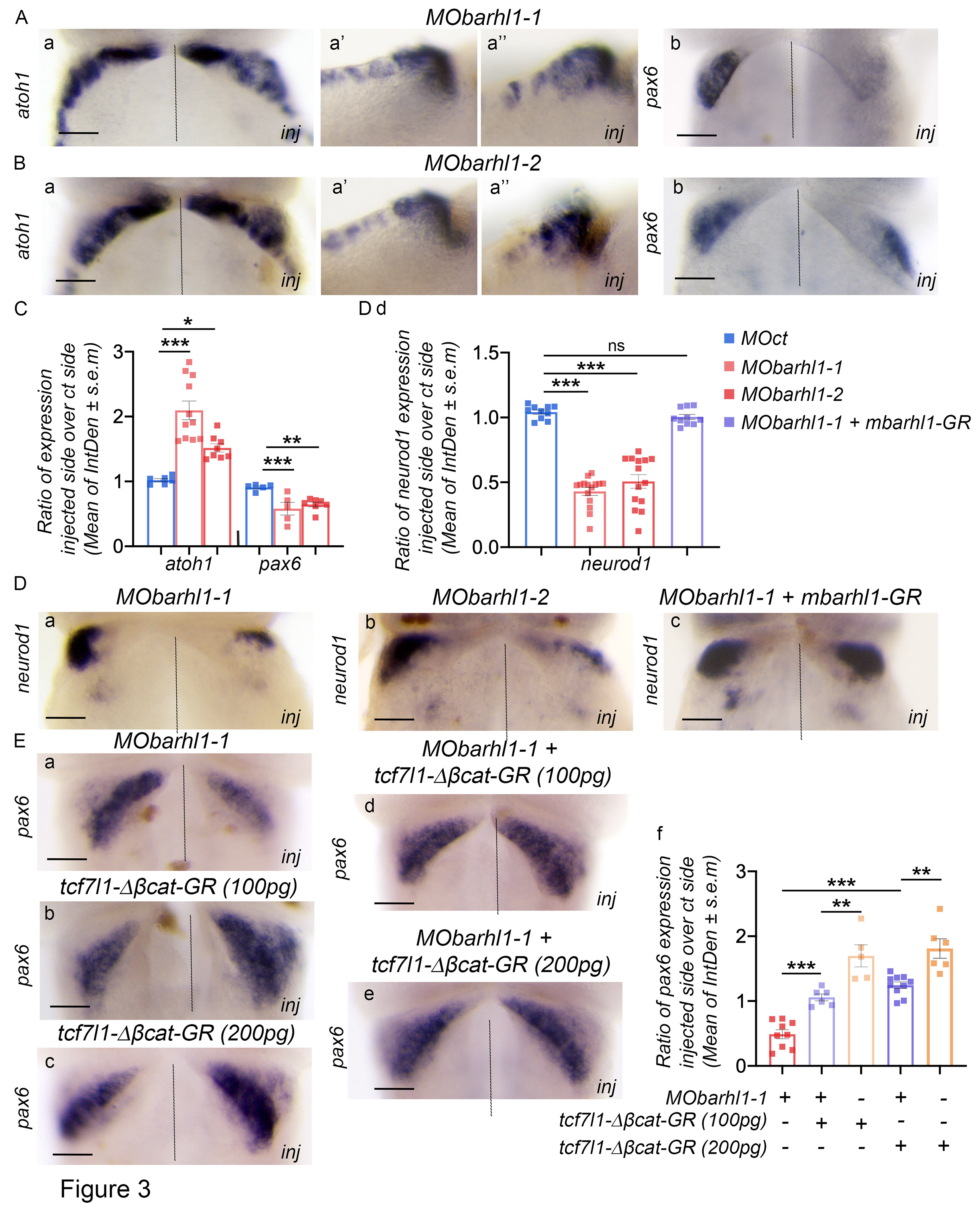
In the cerebellar URL, antagonistic activities of Barhl1 and Tcf are required for the proper development of GNPs. (A-D) Morpholino (MO)-mediated inhibition of Barhl1 induces an ectopic expansion of *atoh1* in the upper rhombic lip (URL) and cerebellar plate and delays GNPs differentiation. *in situ* hybridization (ISH) of stage (st.) 45 *X. laevis* embryos unilaterally injected with (A) *MObarhl1- 1* (15ng) and (B) *MObarhl1-2* (20ng). The non-injected side is an internal control. Shown are dorsal views of *atoh1* (Aa; Ba)*, pax6* (Ab; Bb), and *neurod1* (Da; Db) expressions in the cerebellar anlage. Lateral views of *atoh1* expression in control sides (Aa’, Ba’) and injected sides (Aa”, Ba”) are shown. (C) Quantification of (A) and (B). (Da-c) *MObarhl1* phenotype is rescued by *mBarhl1* overexpression. ISH analysis showing rescue of *neurod1* expression in embryos co-injected with *MObarhl1-1* and *mBarhl1* mRNA. (Dd) Quantification of (D). (E) Inhibition of Tcf activity compensates for Barhl1 depletion. ISH analysis of *pax6* expression in the cerebellar anlage of stage 48 *X. laevis* embryos unilaterally injected with (Ea) *MObarhl1-1* (15ng), (Eb) *tcf7l1-Δβcat-GR* at 100pg and (Ec) *tcf7l1-Δβcat-GR* at 200pg. *pax6* expression was rescued when *MObarhl1-1* (15ng) was co-injected with *tcf7l1-Δβcat-GR* at 100pg (Ed) and at 200pg (Ee). (f) Quantification of (E). Ratio of markers expression in injected side over control side is indicated as mean ± s.e.m. Dotted lines separate injected and control sides. Scale bar 150μm. *inj*: injected side. Statistical analysis was carried out using student’s t-test. C:atoh1:One-way ANOVA(F(2,21)=19.9; p < 0.001) followed by post hoc Tukey test. pax6: One- way ANOVA (F(2,14)=8.63; p = 0.004) followed by post hoc Tukey test. Dd:Kruskal-Wallis test (Chi square=35.6 *p<0.001*, df=3) followed by Nemenyi test post hoc. Ef:One-way ANOVA(F(4,31)=32.9; p < 0.001) followed by post hoc Tukey test.Data are presented as means ± SEM* p ≤ 0.05; ** p ≤ 0.01; *** p ≤ 0.001; **** p ≤ 0.0001.

We next asked whether inhibition of Tcf activity compensates for Barhl1-KD using *pax6* as a marker of GNP commitment. As previously observed, Barhl1-KD delayed GNP differentiation process, while *tcf7l1-Δβcat-GR* overexpression accelerated it (Fig. 3Ea-c). *MObarhl1-1* co- injected with two different doses of *tcf7l1-Δβcat-GR* mRNA rescued the phenotype (Fig. 3Ed- f).

These data provide strong evidence that *MObarhl1* acts by specifically inhibiting endogenous *Xenopus* Barhl1 activity. They also indicate that Barhl1 depletion delays differentiation of GNP. Combined, these observations reveal that Barhl1 and Tcf act in opposing ways within the URL and the EGL, and maintain GNP in an early progenitor state.

### Barhl1 limits Tcf transcriptional activity within the cerebellar primordium

We next asked whether Barhl1 directly controls Tcf transcriptional activity within the cerebellar URL. We investigated interactions between Barhl1, Gro4, and Tcf7l1, by performing co- immunoprecipitation (Co-IP) experiments on protein extracts from HEK293T cells transfected with tagged constructs of Tcf7l1, Gro4, and Barhl1 (Fig. 4A; Fig. S2). In agreement with Barhl1 containing two Engrailed Homology-1 (EH1) motifs known to interact with the WD-repeat domain of Gro, Barhl1 co-immuno-precipitated with Gro4 (Fig. 4Aa). Using Tcf7l1 as bait, we further observed that Tcf7l1 could immuno-precipitate Barhl1, in the presence and absence of Gro4. The presence of Barhl1 does not affect the interaction of Tcf7l1 with Gro4 (Fig. 4Ab).

**Figure 4:**
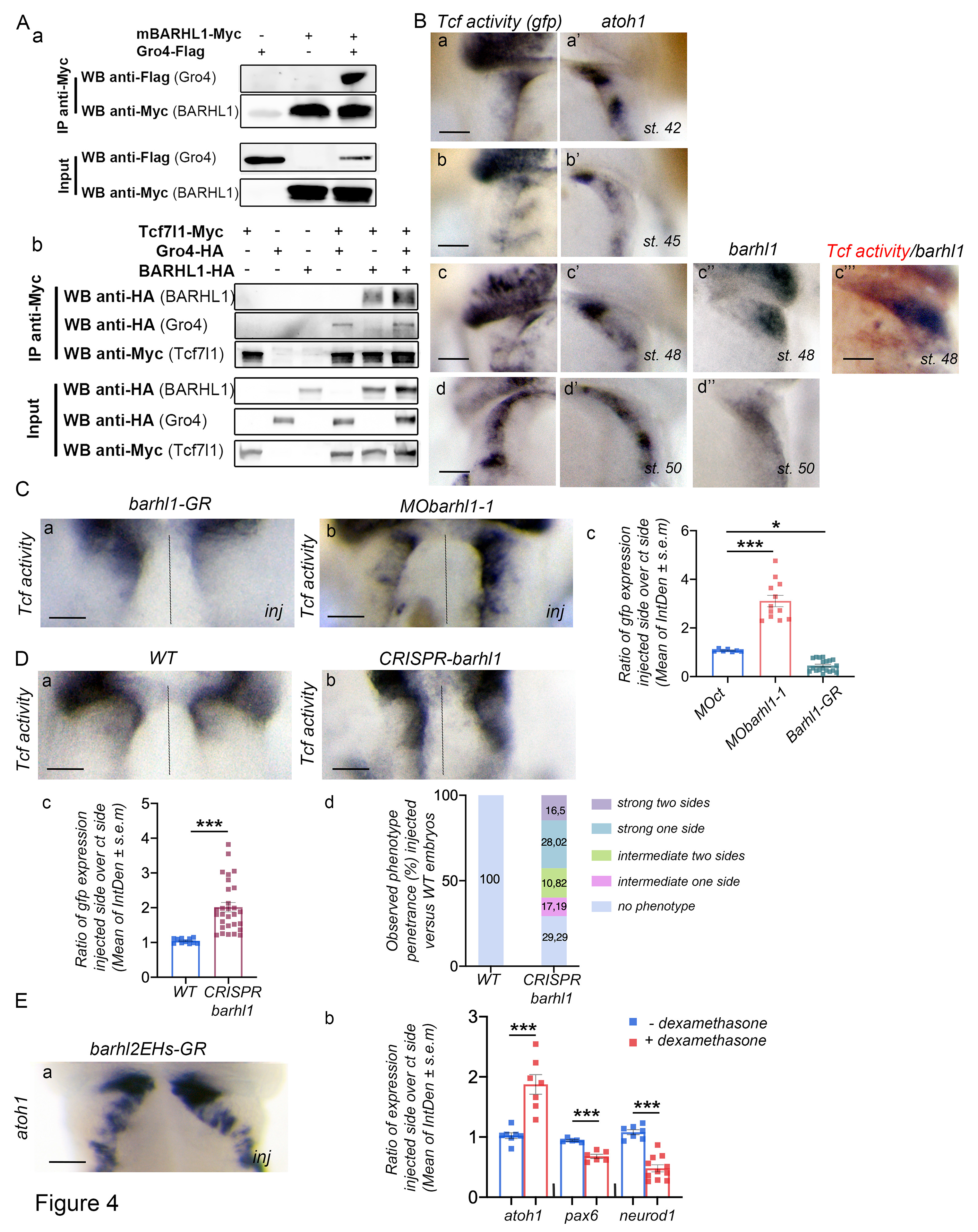
Barhl1 physically interacts with Tcf7l1 and Gro and limits Tcf transcriptional activity. (A) HEK293T cells were transfected with plasmids encoding indicated tagged proteins. Cell lysates were immunoprecipitated (IP) by anti-cMyc antibody. Input and IP samples were subjected to western blot analysis using indicated antibodies. Equal amounts of protein lysates were loaded on SDS-gel. (Aa) Barhl1 interacts with Groucho 4 (Gro4) and Tcf7l1. The interaction between Barhl1 and Tcf7l1 is detected in the presence and in the absence of Gro. (B) Tcf activity is detected in the URL in an area overlapping with that of *atoh1*, and complementary to that of *barhl1*. ISH in *X. tropicalis* pbin7LefdGFP line at indicated stages *(st.)* showing *gfp* (Tcf activity) (Ba, b, c, d), *atoh1* (Ba’, b’, c’, d’) and *barhl1* (Bc”, d”) expression patterns. (Bc’’’) DISH showing expression of *barhl1* (blue) and *gfp* (red). Dorsal views of one side of the embryos are shown. (C, D) Barhl1 limits Tcf transcriptional activity *in vivo*. ISH analysis of *gfp* expression in *X. tropicalis* pbin7LefdGFP embryos injected either unilaterally with (Ca) *mBarhl1GR* (200pg) and (Cb) *MObarhl1-1* or before division with (Db) *Crispr-barhl1*. Embryos injected with *Crispr-barhl1* were compared to their wild-type *(WT)* siblings. (E) Interaction between Barhl1 and Gro is required for Barhl1 function. *mBarhl2EHsGR* only contains the two EH1 motifs of Barhl2 and acts as a dominant negative by capturing Gro. (Ea) ISH showing *atoh1* expression in injected versus control side. Integrated densities *(IntDen)* of markers’ expressions are measured. Ratio of markers expression in injected side over control side is represented (Cc; Dc; Eb) and indicated as mean ± s.e.m. Percentage of phenotype penetrance is quantified in embryos injected with *Crispr/barhl1* versus WT embryos based on indicated criteria. Dotted lines separate injected and control sides. Scale bar 150μm. *inj*: injected side. Statistical analysis Cc: One-way ANOVA(F(2,35)=111,3; p < 0.001) followed by post hoc Tukey test.was carried out using Eb: student’s t-test. Data are presented as means ± SEM * p ≤ 0.05; *** p ≤ 0.001; **** p ≤ 0.0001.

Concatemers of the consensus Tcf binding motif have been used to generate Wnt/Tcf reporter lines, such as *Xenopus tropicalis (X. tropicalis)* transgenic pbin7LefdGFP line ^22, 56, 57^, which contains one copy of a *wnt* reporter gene. We assessed Tcf activity from stage 42 up to stage 50 using this reporter line and observed a positive Tcf activity in the URL. Contrastingly, we did not detect any Tcf activity in the VZ and the EGL at similar developmental stages. Importantly, up to stage 48 we observed that Tcf activity is stronger at the rostral end of the URL. This asymmetry of expression is lost at stage 50 (Fig. 4Ba-d). In addition, we observed a strong correlation between Tcf activity and *atoh1* expression (Fig. 4Ba’-d’), which appears to be complementary to that of *barhl1* (Fig. 4Bc“, c’’’-d”). Taken together, these data are consistent with our previous observations indicating that Tcf activity is strictly necessary for the induction of *atoh1* expression within the URL.

We next assessed the impact of Barhl1 gain of function (GOF) and loss of function (LOF) on Tcf activity (Fig. 4C). Whereas *mBarhl1* overexpression decreased Tcf activity (Fig. 4Ca), we observed a threefold increase in Tcf activity upon Barhl1 downregulation with *MObarhl1-1* (Fig. 4Cb). In contrast, *MOct* had no effect on Tcf activity (Fig. S4A). The effect was quantified (Fig. 4Cc). We further inhibited Barhl1 by selective knock-out (KO) of *xbarhl1* gene in the pbin7LefdGFP line (F0 generation) using Crispr/Cas9 genome editing technology (Fig. S4B). We observed that Barhl1 KO induced an average twofold increase in Tcf transcriptional activity (Fig. 4Da-c). We assessed phenotypic penetrance in Crispr/Cas9 injected embryos based on *gfp* expression. We observed different levels of phenotypic severities in >70% of injected embryos, ranging from a slight increase in *gfp* expression observed in ∼20% of injected embryos to >40% of injected embryos exhibiting strong to complete penetrance as observed by a significant increase in *gfp* expression in the R1 (Fig. 4d).

To determine whether Barhl1 effects were mediated through its interaction with Gro, we used and inducible form of *mBarhl2-EHsGR, which* contains the two EH1 domains of Barhl2 (Fig. S2) and has been previously demonstrated to act as a dominant negative for Tcf repressive activity by competing for Gro binding ^52^. Overexpression of *mBarhl2EHsGR* induced a phenotype similar to that of Barhl1-KD in terms of increasing the URL/EGL size at the expense of GNP commitment/differentiation (Fig. 4Ea, b; S4).

Taken together, our data establish that Barhl1 directly interacts with Tcf7l1 and Gro, limiting their transcriptional activity in the cerebellar URL.

### Through its limiting of Tcf transcriptional activity, Barhl1 allows GNP to leave their early progenitor state and exit the proliferating URL

The URL germinative zone is characterized by its proliferative state, and its bordering of the roof plate. We first asked whether the EGL is proliferative in *Xenopus*, and second whether the enlargement of the URL/EGL territories observed in Barhl1-KD tadpoles was corroborated with an increased proliferation in the URL and/or within the cerebellar plate. Using immunofluorescence staining for Phosphorylated-Histone H3 (PHH3), a marker of cells undergoing mitosis, counterstained with a cell nuclear marker, we measured proliferation in tadpoles injected with either *MOct* or *MObarhl1-1* at stage 45 and 48. In agreement with previously published data ^41^, at both stages, PHH3+ cells were solely detected within the URL (Fig. 5Aa). To investigate the capacity of URL-derived GNP to leave their proliferative state, and to progress along their developmental trajectory, we measured the length of the URL on the injected side compared to the control side in embryos either injected with *MOct* or *MObarhl1-1*. At stage 45 and 48, Barhl1-KD embryos exhibited a 1.2-fold lengthening of the URL on the injected side relative to the control side (Fig. 5Ac-e). Because it was morphologically easier to distinguish the URL from the VZ at stage 48, we performed our following analysis at this later developmental stage (Fig. 5Ab). At stage 48, we measured an average 2-fold increase in the number of PHH3+ cells on the injected side compared to the control side (Fig. 5Ac, d, f). Moreover, we observed and quantified the presence of proliferating cells in the cerebellar plate, which is normally devoid of PHH3+ cells as observed in tadpoles injected with *MOct* (Fig. 5Ac, d, g). Taken together, these results confirm that in *Xenopus,* the EGL is non proliferative, and show that Barhl1-KD cells are compromised in their ability to leave the URL niche and become postmitotic.

**Figure 5:**
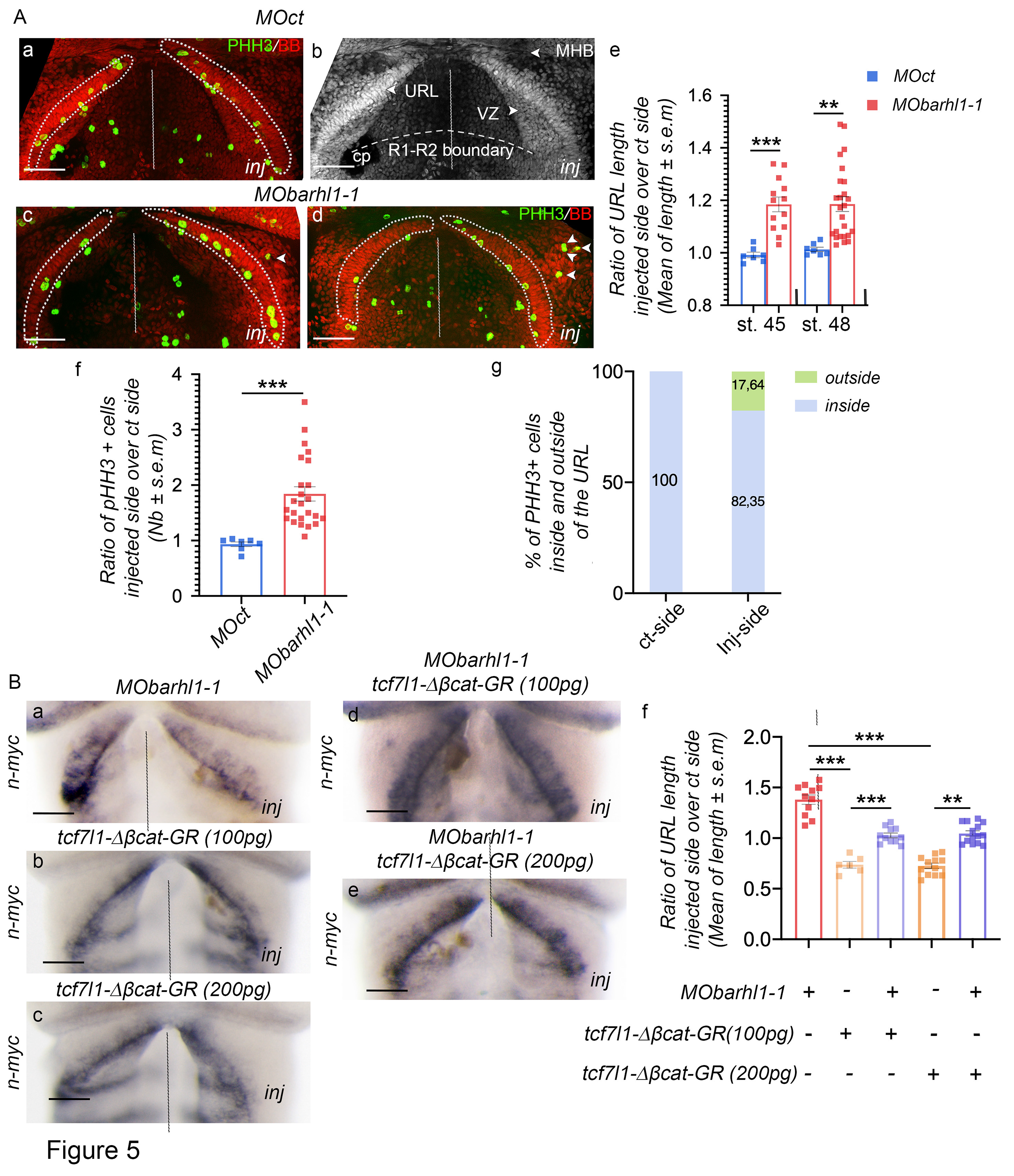
In the URL, Barhl1 activity as an inhibitor of Tcf transcription is required for GNPs to exit their germinative zone and become post-mitotic. (A) Barhl1-KD induces an increase in the URL length associated with increased proliferation within this compartment. (a-d) Imaging of cerebellar anlage of stage 48 *X. laevis* tadpoles unilaterally injected with *MOct* and *MObarhl1-1* (15 ng). Collected neural tubes were stained for the mitotic marker PhosphoHistone-H3 (PHH3) (green) merged with bisbenzimide (BB) (red). In *Xenopus*, the EGL is devoid of proliferating cells. (e-g) Quantifications of (A). The ratio of (e) measured URL length and (f) PHH3+ cells in injected side over control side are represented. (Ac, d) PHH3 positive cells are ectopically detected in the cerebellar plate (Ac, d white arrow heads) of injected embryos. (g) Percentage of PHH3+ cells located inside the URL compared to that located outside the URL were quantified on the injected side and on the control side. (E) The abnormal increase in URL length is rescued upon co-inhibiting Tcf and Barhl1 activities. ISH analysis of *n-myc* expression in the cerebellar anlage of stage 48 *X. laevis* embryos unilaterally injected with (Ba) *MObarhl1-1* (15ng), (Bb) *tcf7l1-Δβcat-GR* at 100pg and (Bc) *tcf7l1-Δβcat-GR* at 200pg. *n-myc* marks the boundaries between different rhombomeres which allows the exact measurement of URL length. URL length was rescued when *MObarhl1-1* (15ng) was co-injected with *tcf7l1-Δβcat-GR* at 100pg (Bd) or at 200pg (Be). Ratio of URL length in injected side over control side is represented (Bf) and indicated as mean ± s.e.m. Dotted lines separate injected and control sides. Scale bar 150μm. *inj*: injected side; cp: choroid plexus; URL: Upper Rhombic Lip; VZ: Ventricular Zone; R1-R2: Rhombomere 1 and 2; MHB: Midbrain-Hindbrain Boundary. Statistical analysis Ae, Af: was carried out using student’s t-test. Bf: One-way ANOVA(F(4,49)=65.1; p < 0.001) followed by post hoc Tukey test.Data are presented as means ± SEM ** p ≤ 0.01; *** p ≤ 0.001; **** p ≤ 0.0001.

We next investigated whether Tcf inhibition could counteract Barhl1-KD effect on URL extension. As previously observed, whereas Barhl1-KD induced an extension of the URL length, *tcf7l1-Δβcat-GR* overexpression reduced it (Fig. 5Ba-c), and co-injection of *MObarhl1- 1* and *tcf7l1-Δβcat-GR* mRNA brought back the URL size to normal (Fig. 5Bd-f).

In conclusion, Barhl1 activity as an inhibitor of Tcf transcription is strictly necessary for URL cells to exit their niche, become postmitotic and enter the EGL.

### Transcriptomic analysis of Barhl1 activity in the developing cerebellum

*barhl1* starts to be significantly expressed in the developing cerebellum at stage 40. To further document Barhl1 activity, we designed an RNA-sequencing experiment allowing the identification of Barhl1 direct and indirect target genes in the early *Xenopus* cerebellum.

We isolated and sequenced RNA from R1 of stage 42 tadpoles previously injected at the 4 cells stage in the 2 dorsal blastomeres with *MObarhl1-1*, *MObarhl1-2* or *MOct* together with *gfp*. Tadpoles were selected for hindbrain injection and R1 were dissected. Samples were compared through differential expression (DE) analysis. Genes with adjusted p*-*value (pAdj) inferior to 0.001 were selected as significant DE genes (DEG) (Table S1). Principal component analysis of these R1 samples demonstrated that they clustered by Barhl1-KD status (Fig. 6A), indicating that changes in gene transcription were consistent across different clutches. At stage 41-42 we identified 1622 and 830 genes differentially expressed between respectively *MObarhl1-1* and *MObarhl1-2* injected R1, compared with *MOct* injected R1. Amongst these DE genes 575 were common between MOs injected samples (Fig. S6B). A selection of significant upregulated (Log2FC>0.4) and downregulated (Log2FC<-0.4) genes are represented in volcano plots for both MOs (Fig. 6B). Furthermore, we generated a heatmap representing upregulated and downregulated common DEG for both MOs (Fig. 6C).

**Figure 6:**
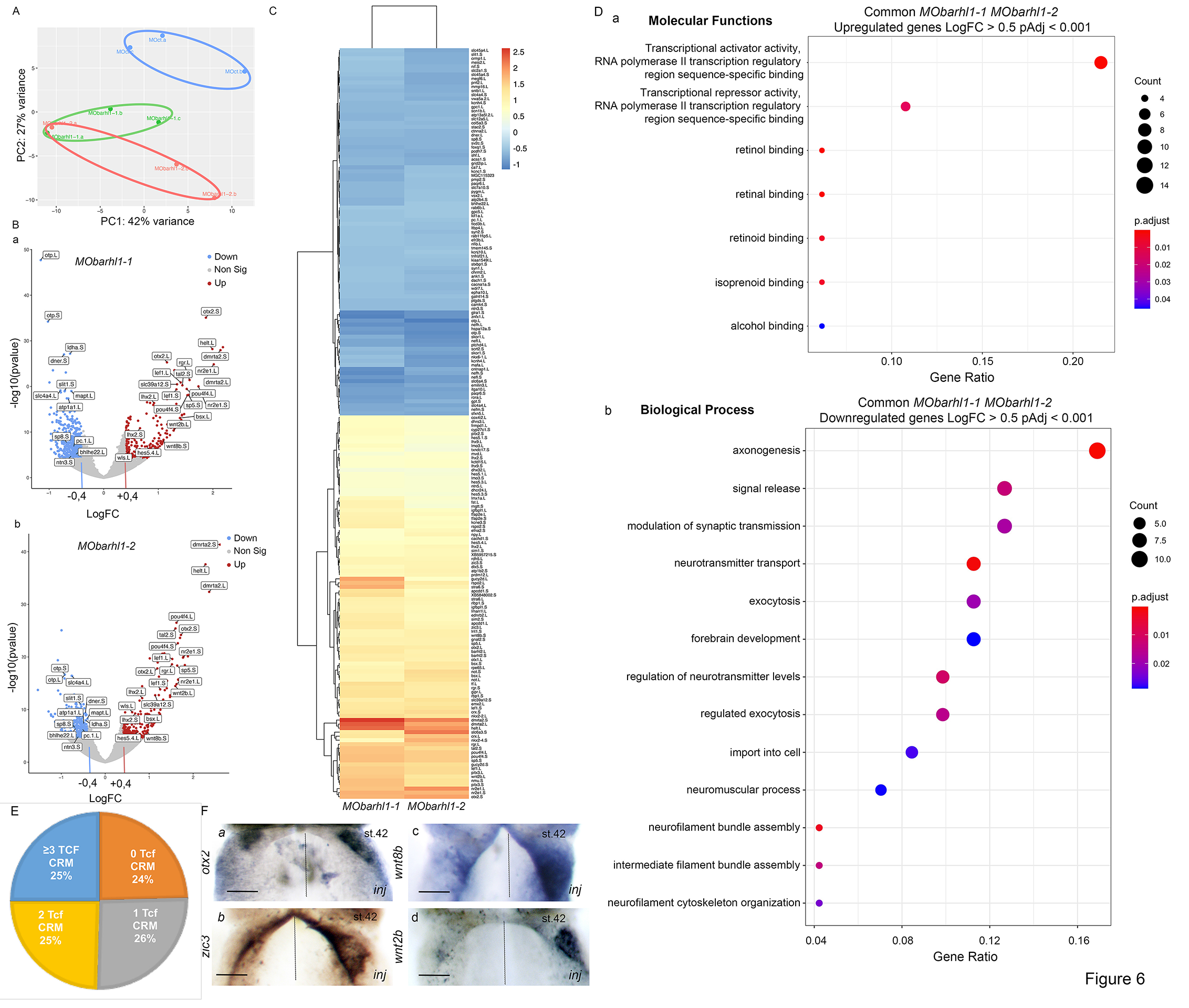
RNA-sequencing data processing and analysis. (A) Principal Component Analysis (PCA) plots were obtained based on RNAseq data aligned with STAR and reads counted using feature-counts. Three samples have been generated for each condition. Sample groups are represented by different colors as indicated. Each dot refers to a sample. Samples showing similar gene expression profiles are clustered together. (B) **Volcano plots** showing a selection of significant DEGs with pAdj < 0.001 in (a) *MObarhl1-1 vs MOct* and (b) *MObarhl1-2 vs MOct*. Upregulated genes with Log2FC>0.4, and downregulated genes with Log2FC<-0.4 are shown. Red and blue dots indicate significant DEGs that are upregulated and downregulated, respectively. Grey dots denote RNAs with non- significant difference. PCA and volcano plots were generated using Galaxy. (C) **Differentially expressed genes (DEGs) visualization** Heatmap displaying expression profiles of most significantly upregulated and downregulated DEGs for each condition (*MObarhl1-1 vs MOct* and *MObarhl1-2 vs MOct*). Each row represents a gene, and each column represents a sample. Results are shown as a gradient from blue (downregulated) to dark orange (upregulated). Heatmap is generated using R package. (D) **Gene ontology enrichment comparison.** Shown on Y axis are the altered molecular functions (a) and biological processes (b) for selected (A) upregulated (Log2FC≥0.4, PAdj<0.001), and (b) downregulated (Log2FC≤- 0.4, PAdj<0.001) DEGs respectively. Enrichment analysis comparing functional profiles among *MObarhl1-1* and *MObarhl1-2* was performed on the DEGs in common between both conditions. Results are visualized as a dot plot based on indicated gene counts and adjusted p-values for enrichment. Dot size corresponds to the count of differentially expressed genes associated with the molecular function or the biological pathway, and dot color refers to the adjusted P-value for enrichment. (E) **TCF Cis Regulatory Motifs (CRM) in regulatory regions of MOBarhl1 DEGs**: pie chart of % of *MObarhl1* DEGs containing either no TCF CRM (orange), one TCF CRM (grey), two TCF CRM (yellow) and three or more TCF CRM (blue) located 5Kb upstream or downstream of their TSS. (F) **ISH analysis of 4 DEGs**: Dorsal views R1 territory of st. 42 *X. laevis* embryos unilaterally injected with *MObarhl1-1* using *wnt8b*, *wnt2b*, *zic3*, *otx2* as ISH probes as indicated*. inj:* injected side.

As a first approach we performed gene ontology analysis (GO) based on pAdj<0.001 DEG using the clusterProfiler algorithm ^58^ and compared altered biological functions between both Barhl1-KD conditions (Fig. 6D). Our GO analysis reveals that the most significantly upregulated genes act as transcriptional activators when bound to DNA (Fig. 6Da). In agreement with a delay of GNP differentiation the downregulated DEG were found to be involved in axon development and axonogenesis, in addition to neuronal differentiation (Fig. 6Db). Indeed, our differential expression dataset reveals that genes that are the most up- regulated in the *MObarhl1-1* and *MObarhl1-2* conditions are involved in adult neural stem cell (NSC) maintenance. For example, *dmrta2* that encode for doublesex and mab-3-related transcription factor a2, also known as *dmrt5,* the orphan nuclear receptor subfamily 2 group E member 1 (*nr2e1*) commonly known as Tailless, that are upregulated with a Log2FC over 1.5. We also observed a down regulation of the Delta/Notch-like epidermal growth factor (EGF)- related receptor (*dner*) (Log2FC ≤ -0.5), which has been suggested to be a neuron-specific Notch ligand ^59^. Indeed, *dner* has been suggested to inhibit neural proliferation and induce neural and glial differentiation ^60^. We also identify Basic Helix-Loop-Helix Family Member E22 (*bhlhe2*), a downstream target of NEUROD1, which is strongly downregulated in Barhl1 depleted R1 (Log2FC ≤ -0.5) ^61^.

Our functional data argue that Barhl1 mostly act through inhibition of TCF transcriptional activity. We first investigated the presence of Barhl1 Cis Regulatory Motifs (CRM) defined as CAATTAC/G and its mirror motif ^62^, within the regulatory sequences - 5Kb upstream or downstream of the Transcription Start Site (TSS) - of previously identified DEG common to *MObarhl1-1* and *MObarhl1-2* conditions. We observed that all DEG regulatory regions contain at least 2 Barhl1 CRM, 87.5% contain 5 or more Barhl1 CRM and 40% 10 or more Barhl1 CRM (Table 1A). Thereby our identified DEG appear to be Barhl1 direct target genes. To investigate which Barhl1 target genes are also regulated by TCF we similarly searched for TCF CRM defined as CTTTGAA/CTTTGAT, within the regulatory sequences of previously identified DEG common to *MObarhl1-1* and *MObarhl1-2* conditions (Table 1B; Fig. 6E) ^63, 64^. We observed that 76% of Barhl1 depleted DEG regulatory regions contain at least one Tcf CRM: 26% contain one CRM, 25% contain two Tcf CRM and 25% contain three and more Tcf CRM (Fig. 6E).

**Table 1:**
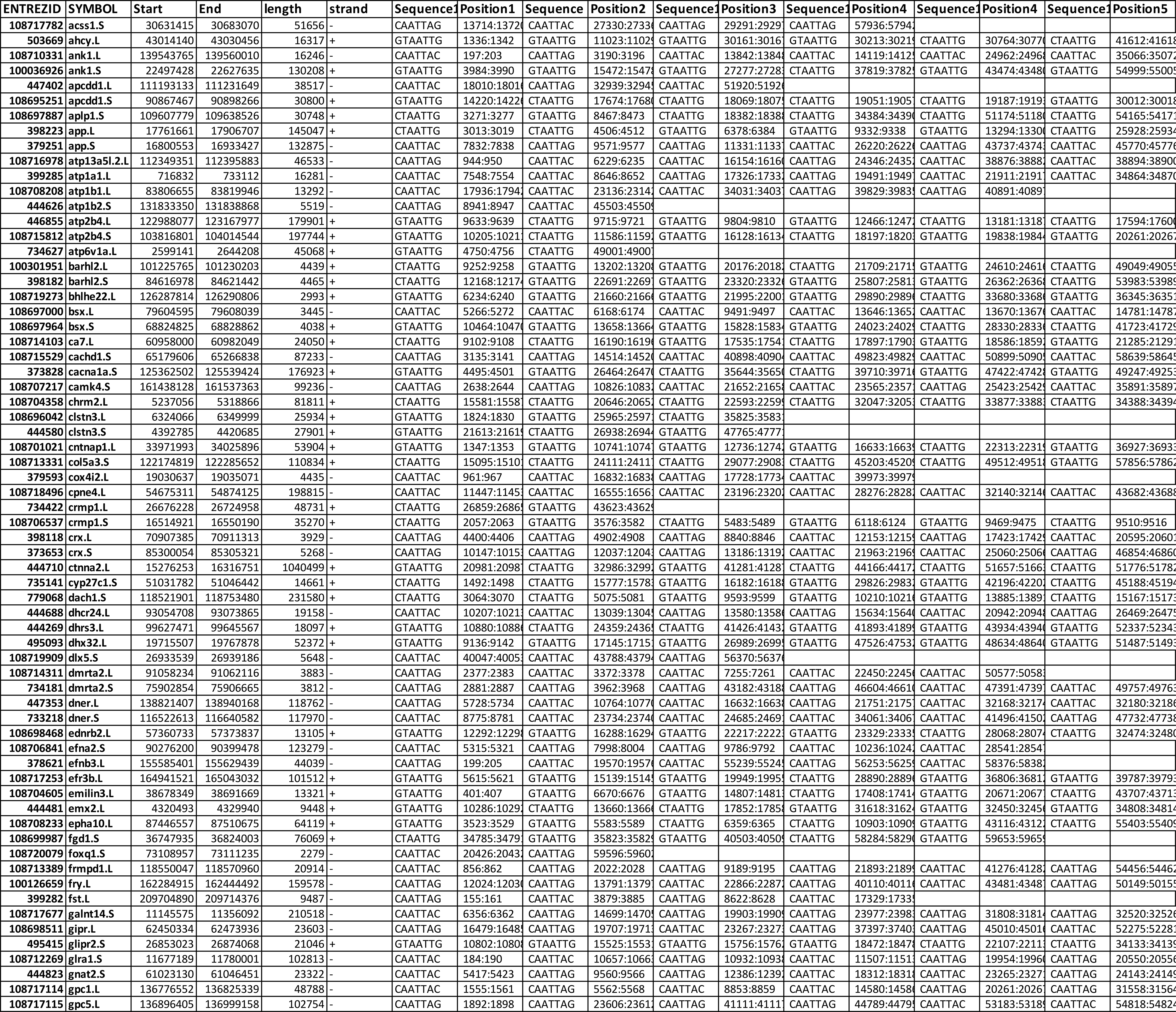
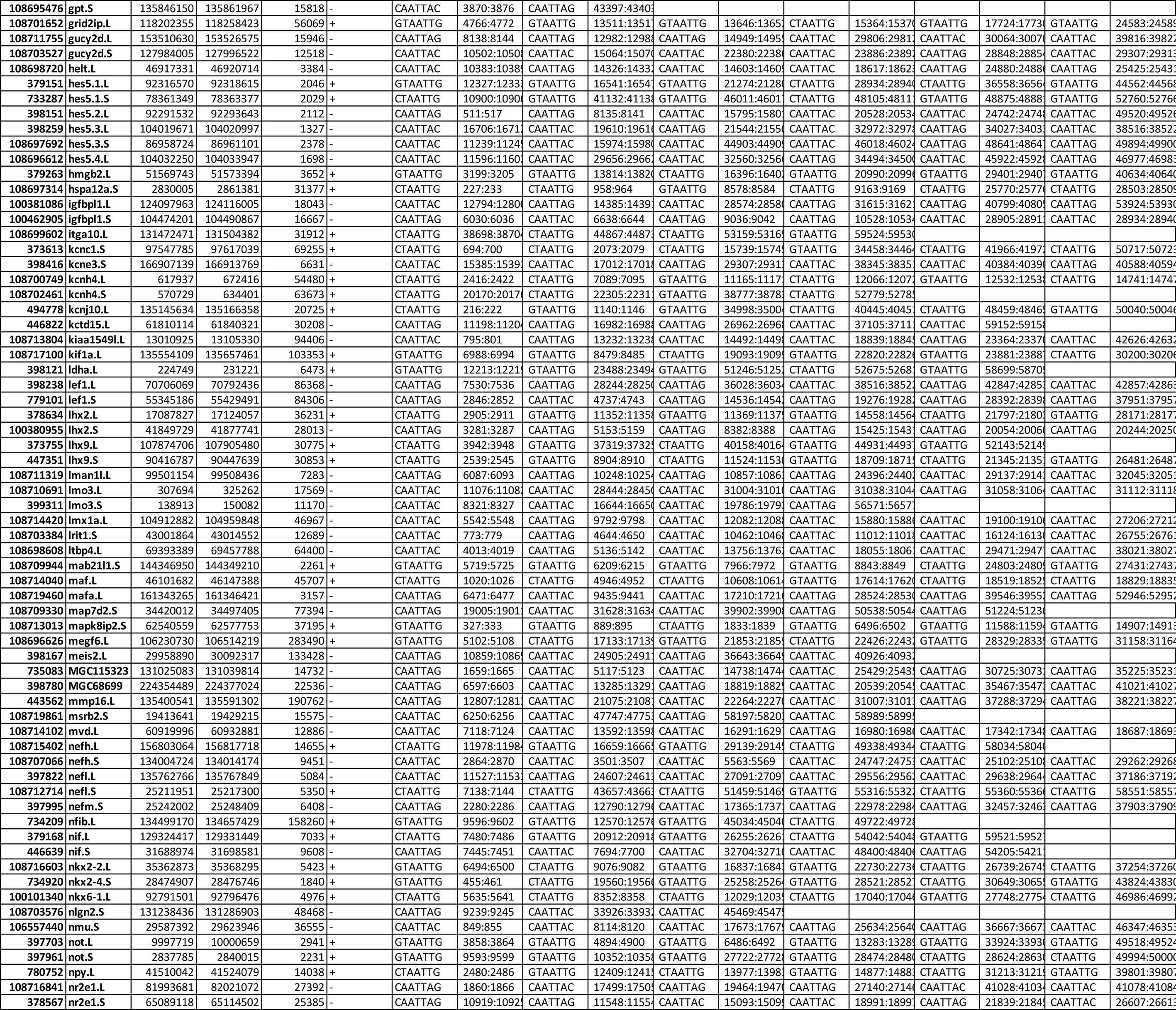
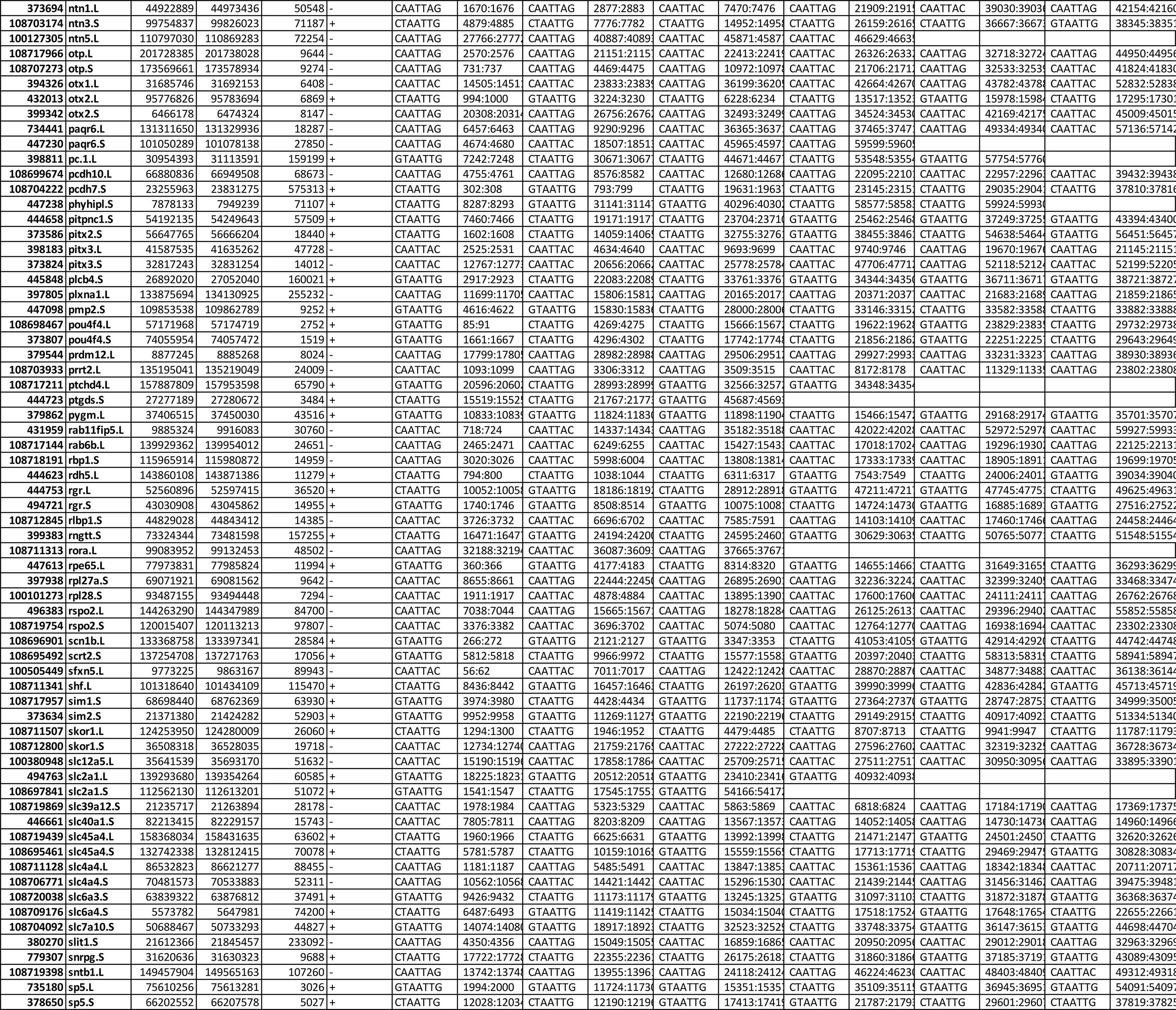
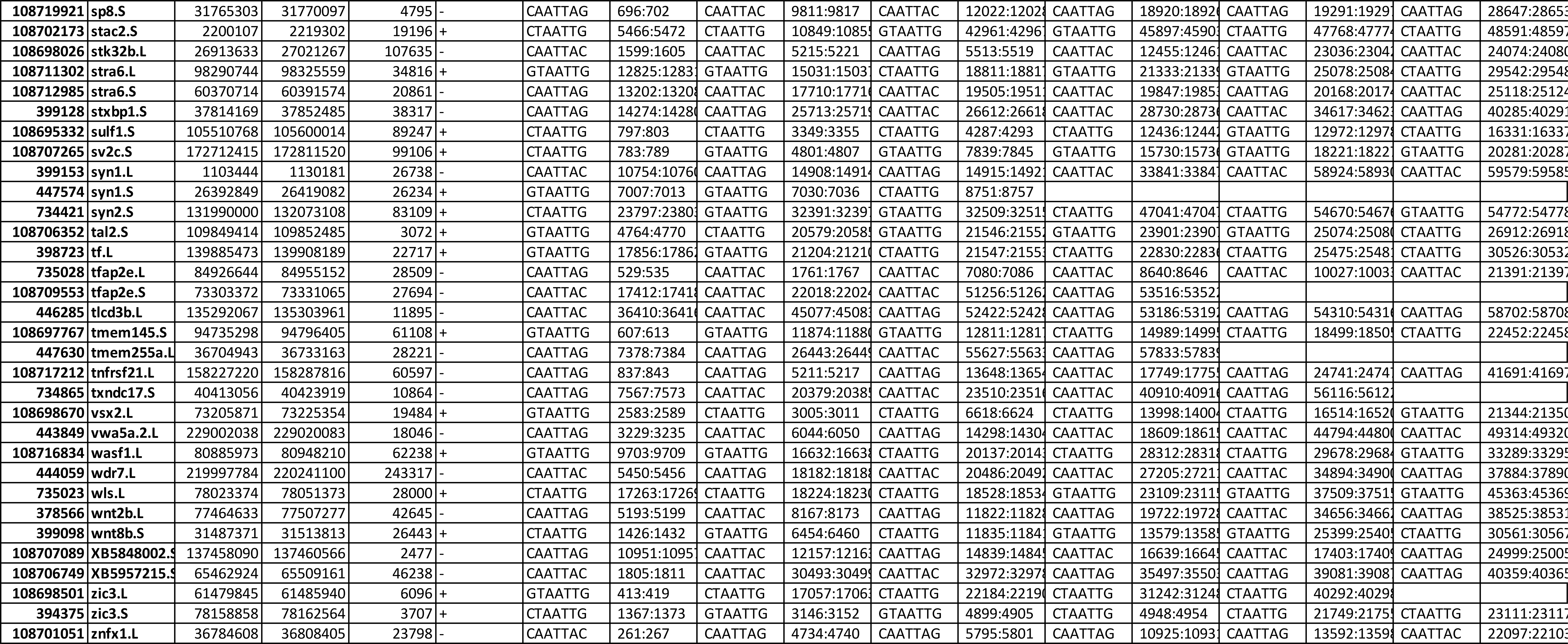
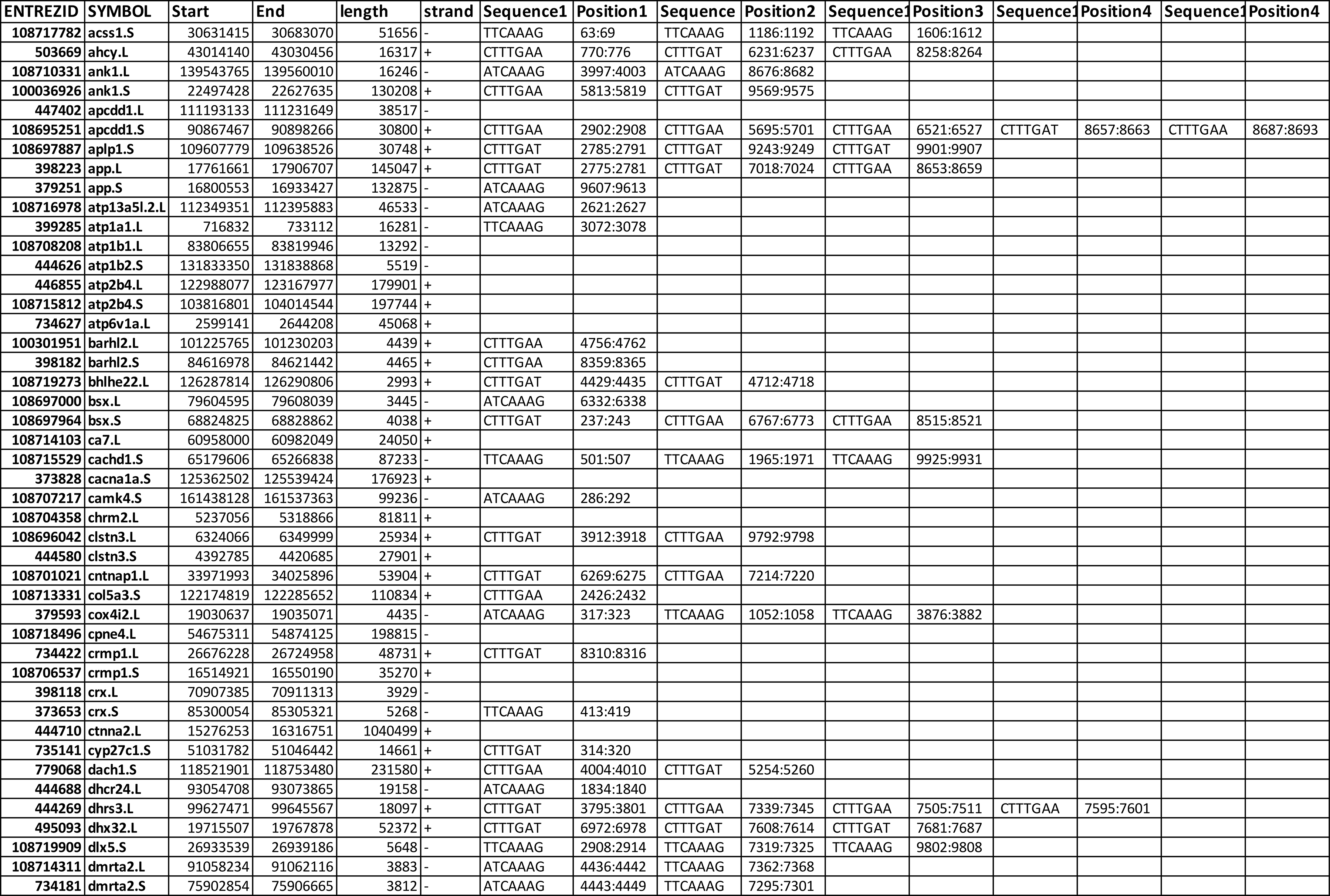
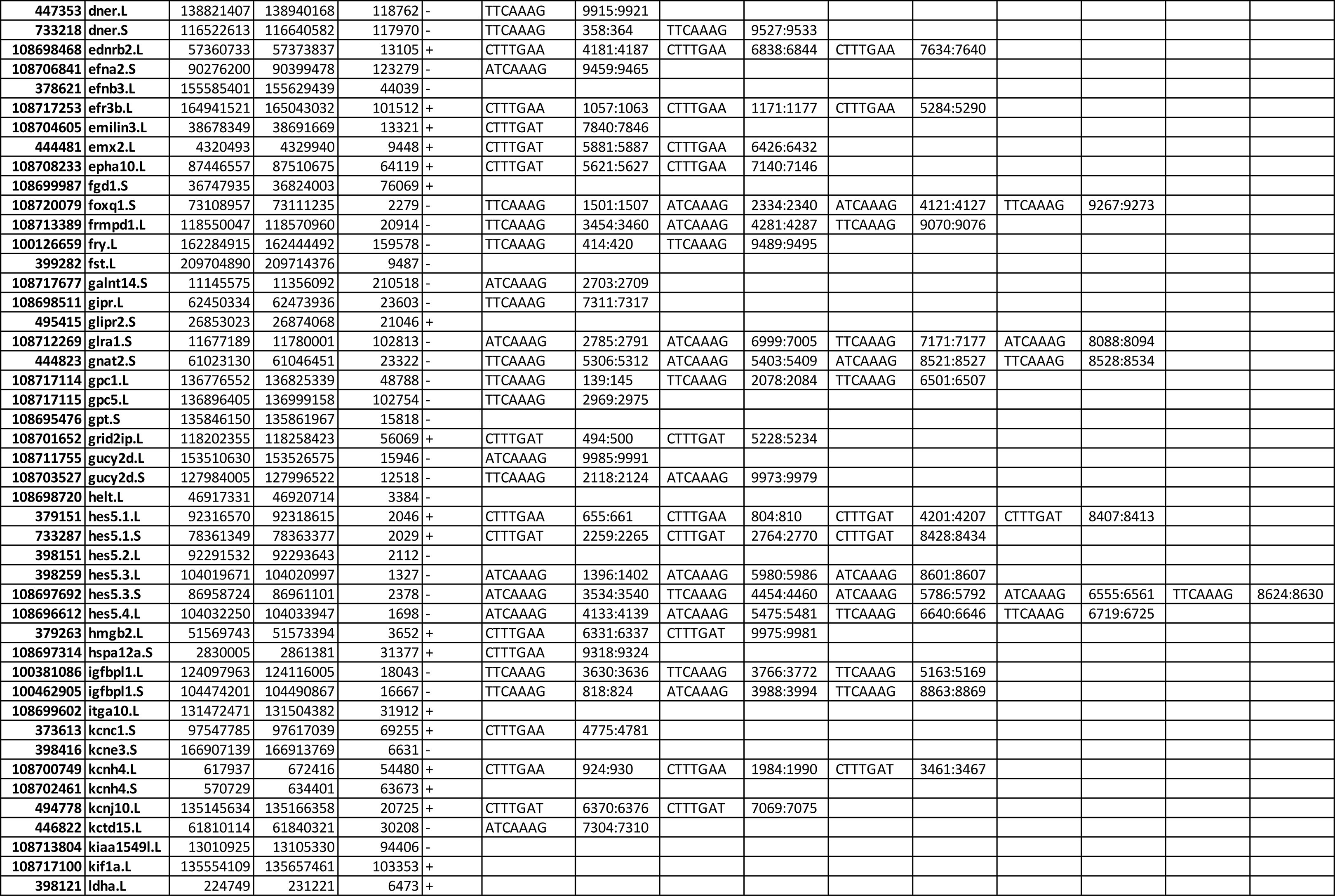
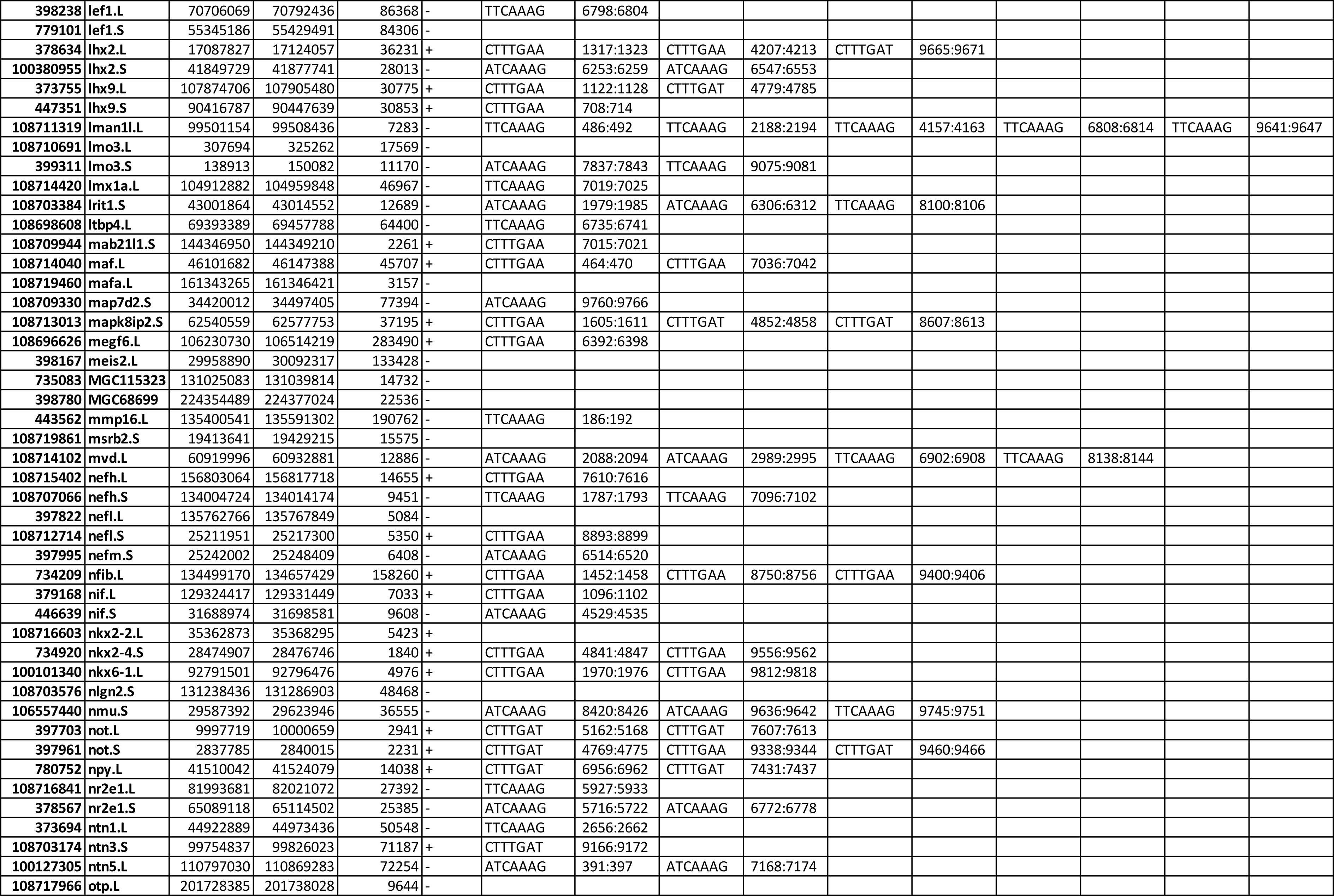
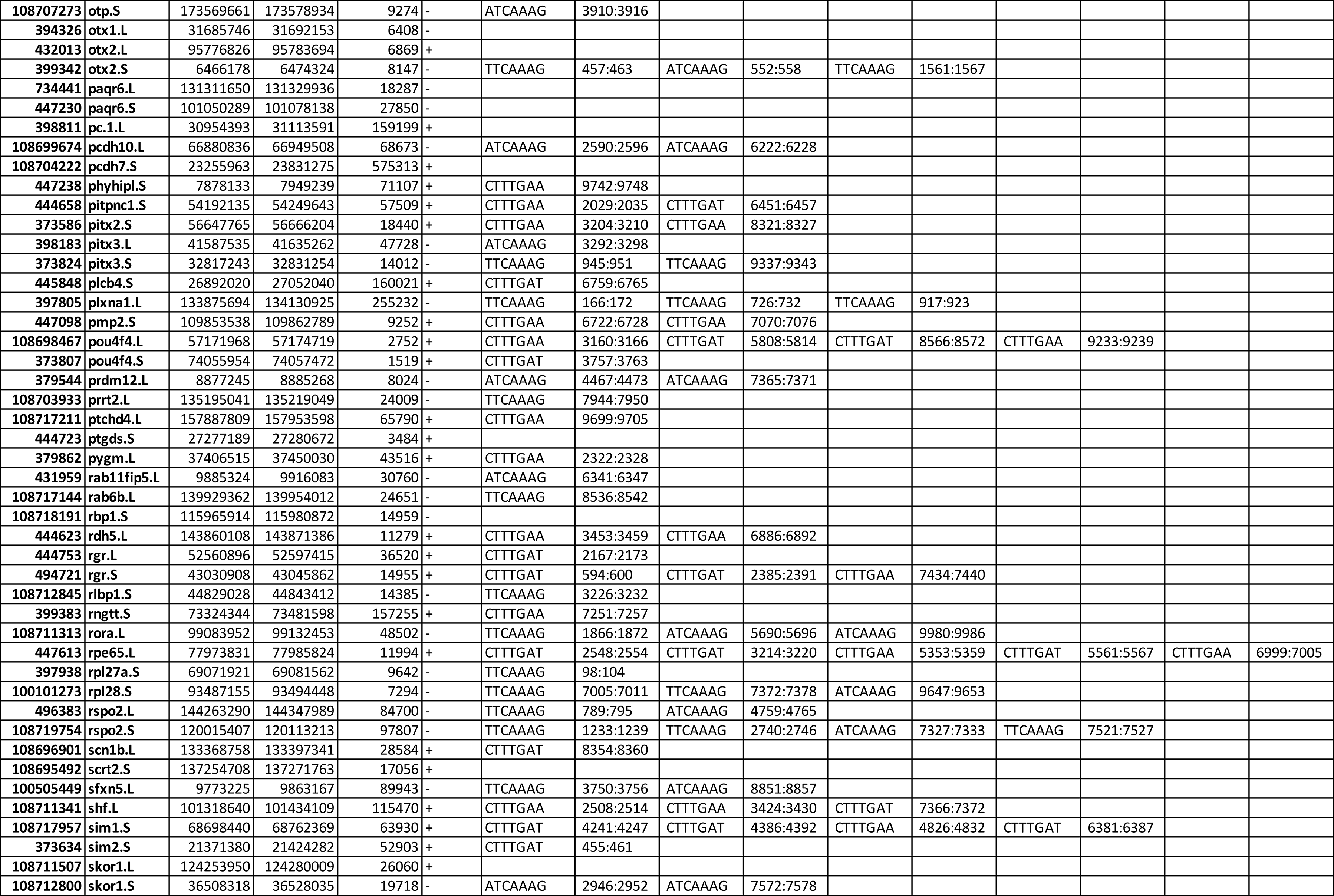
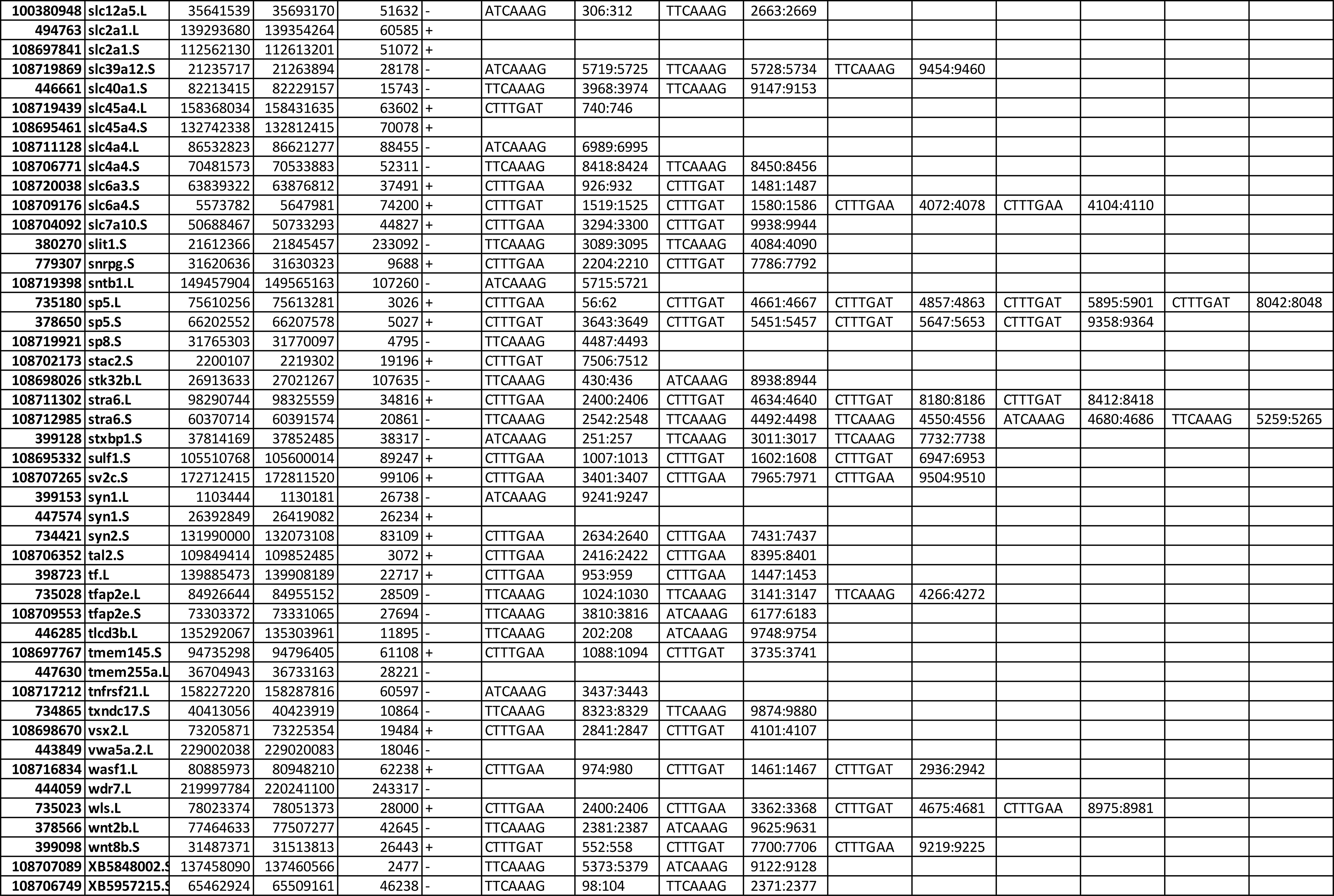
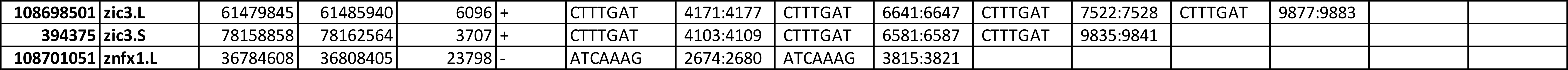
Barhl1 and TCF Cis Regulatory Motif (CRM) on regulatory regions of Barhl1 depleted DEGs. . We explore the putative transcription factor-target relationships of Barhl1 (A) and Tcf (B) on Barhl1 depleted DEGs (PAdj<0.001, Log2FC≥0.45 or Log2FC≤-0.45). We applied R packages Biostrings (v2.64) and GenomicFeatures (v1.48) and determine potential (A) Barhl1 binding sites (5’-C-A-A-T-T-A-C/G-3’) (and the mirror sequence (5’-G/C-T-A-A-T-T- G-3’)) ^60^, or (B) TCF binding sites (5’-C-T-T-T-G-A/T-A-3’) (and the mirror sequence (5’-T-A/T- C-A-A-A-G-3’)) ^61, 62^ 5Kb upstream and downstream of the Transcription Start Site (TSS) of DEGs using *X. laevis* v10.1 genome assembly downloaded with its corresponding annotation file from Xenbase. For each gene identified through its EntrezID and its symbol, is indicated the sequence of the detected putative CRM and its position within the gene locus.

**Table 2:**
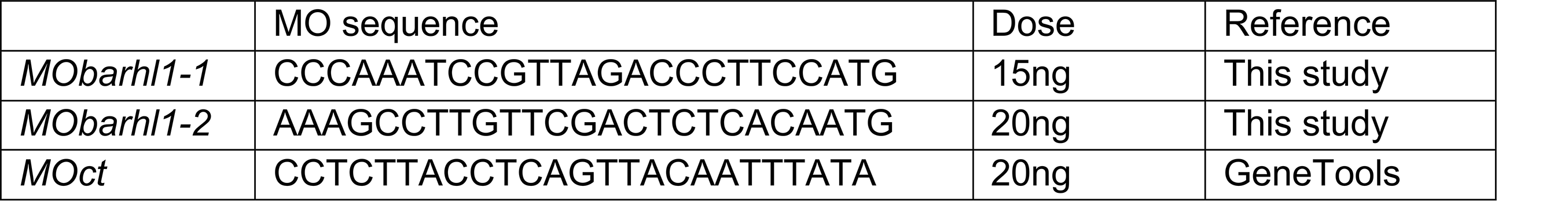
Morpholino (MO) oligonucleotide sequences used in this study.

**Table 3:**
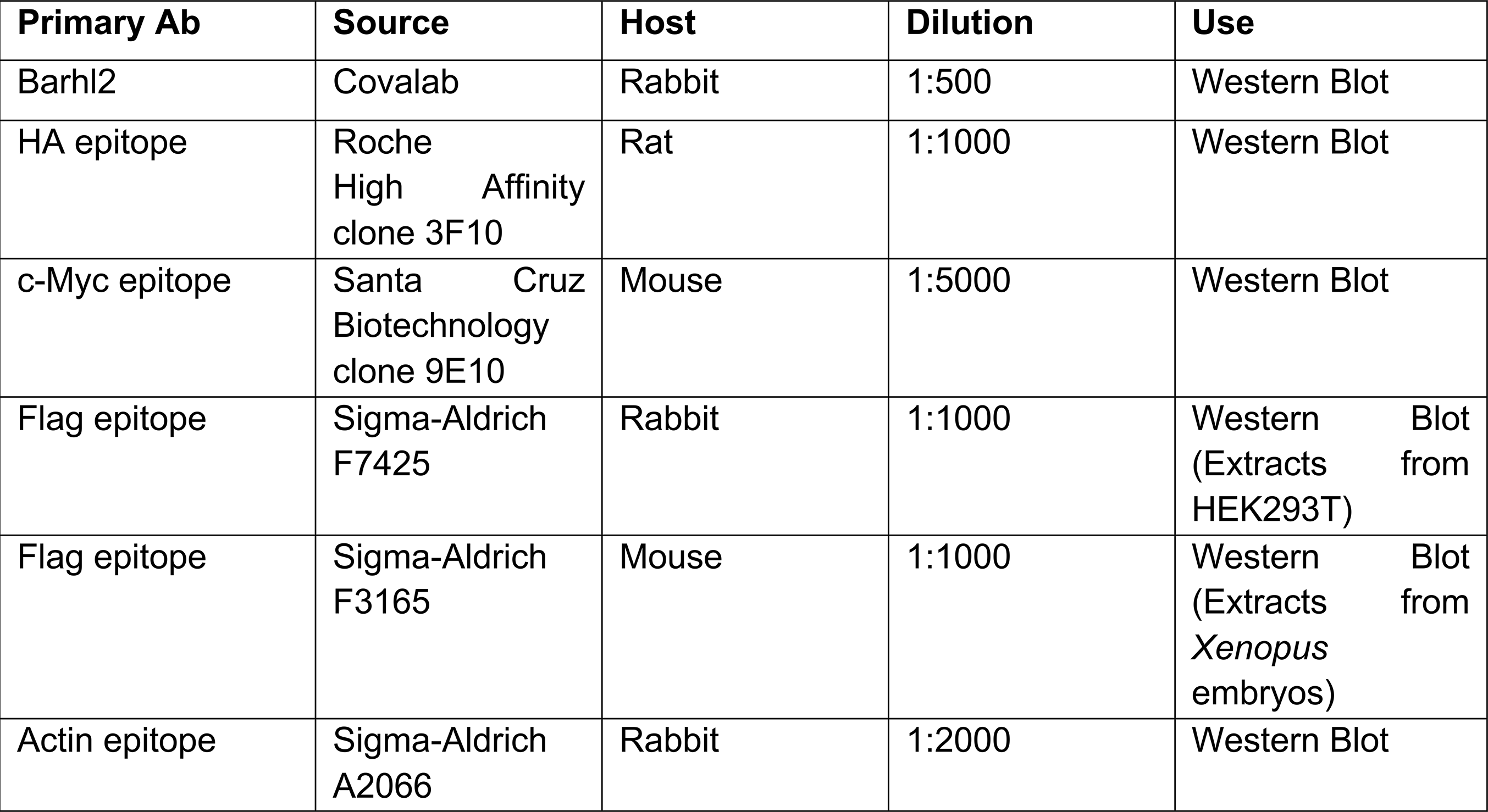
Primary antibodies (Ab) used in this study

**Table 4:**
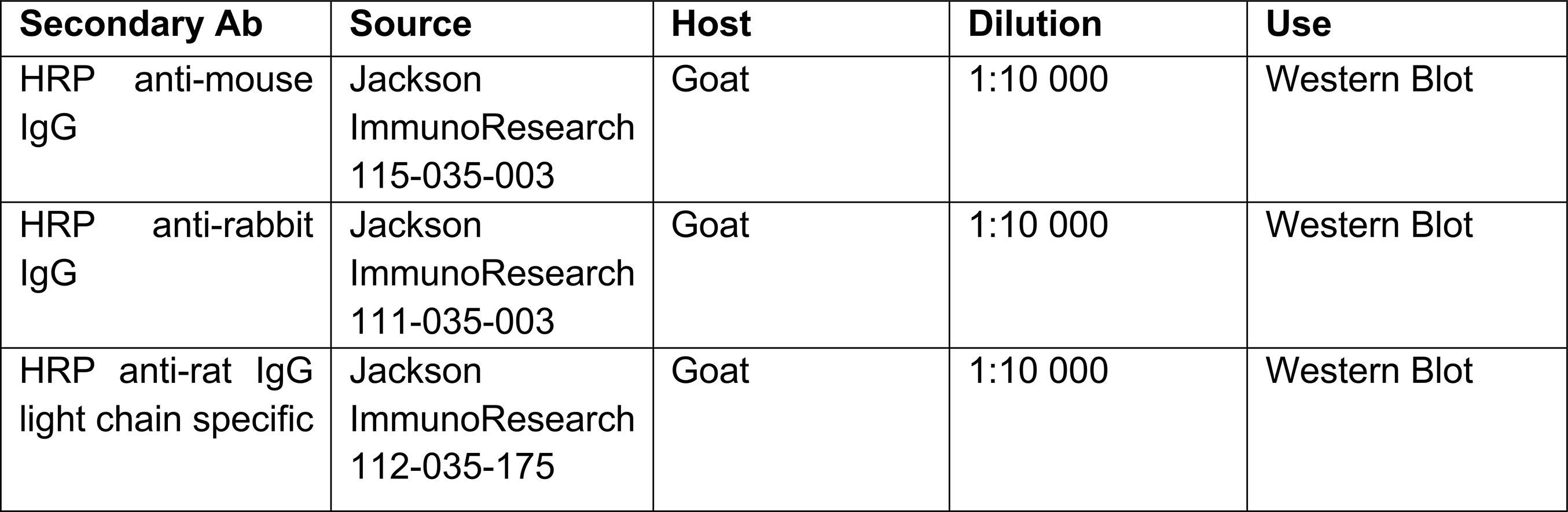
Secondary antibodies (Ab) used in this study.

Using ISH, we explored changes in two up-regulated DEG. *zic3*, a member of the Zinc Finger of the Cerebellum (Zic) family known to be involved in regulation of neuronal progenitor proliferation versus differentiation, and cerebellar patterning reviewed in ^65–67^ (Log2FC≥1) and *otx2* that is detected in a subset of GNP (Log2FC≥1.2). At stage 41-42 we observed a significant expansion of both *otx2*, and *zic3* expression territories within the cerebellar plate (Fig. 6Fa,b). *zic3* transcripts are present in the URL, and *zic3* regulatory regions contain at least three Tcf CRM (Table 1B). *otx2* regulatory regions contain either no Tcf CRM (*otx2.L*), either 3 Tcf CRM (*otx2.S*). Thereby we argue that *zic3* is a direct target of Tcf, whereas *otx2* genes may be indirect targets of Barhl1 depletion effect on GNP development.

Amongst the DEG, we also observed an upregulation of faithful reporters of Notch pathway activation *hes5* family genes (*hes5.1, hes5.2, hes5.3, hes5.4*), which regulatory regions contain between 1 and 5 Tcf CRM (Log2FC ≥ 0.5 ≤ 0.91) and HES/HEY-Like Transcription Factor (*helt*) (Log2FC ≥ 2), which regulatory regions lack Tcf CRM. HELT is closely related to Hairy enhancer of split proteins that act as a major downstream effector in the Notch pathway, that is required for the maintenance of NSC, and a proper control of neurogenesis in both embryonic and adult brains ^68^.

Finally and importantly amongst the DEG, we identified markers of Wnt pathway activity including one of the bona fide direct target gene of Wnt signalling *sp5* (Log2FC ≥ 1.55; 3 Tcf CRM) (Fig.6B,C; Suppl. Table 1), together with *lef1* (Log2FC ≥ 1.55; one Tcf CRM) known to be a target of Wnt/β-catenin signalling ^69^, two Wnt secreted signals *wnt8b* (Log2FC≥1; one Tcf CRM) (Fig. 6Fc), and *wnt2b* (Log2FC≥1.3; one Tcf CRM) (Fig. 6Fd), both of which activate the Wnt canonical pathway, and Wnt Ligand Secretion Mediator (*wls*) (Log2FC≥0,5; one Tcf CRM), which expression in the URL orchestrates cerebellum development in mice ^70, 71^. Thereby Barhl1 depletion activates Wnt/Tcf activity throughout the cerebellar anlage, specifically within the URL and the cerebellar plate.

Taken together, our transcriptomic analysis identified direct and indirect Barhl1 target genes. Within the R1 territory, when upregulated these genes are involved in i) the maintenance of neural stem/progenitor properties, ii) the enhancement of Notch activity, iii) promoting Wnt/Tcf activity. When down regulated they are mostly involved in inhibiting proliferation and promoting neural differentiation. Our analysis of Barhl1 target genes regulatory regions confirms our functional analysis demonstrating that Barhl1 mostly acts by inhibiting Tcf transcriptional activity.

## DISCUSSION

This study conducted in amphibians provides evidence that the development of the *Xenopus Atoh1* lineage is similar to that of higher vertebrates. We show that Tcf transcriptional activity is necessary for inducing the cerebellar URL, as well as *atoh1* expression. Furthermore, we demonstrate that Barhl1 plays a critical role in promoting the exit of URL cells from their niche, and initiating their differentiation trajectory towards mature GN. Most importantly, Barhl1 acts primarily by inhibiting Tcf transcriptional activity. Transcriptomic analysis of Barhl1 depletion in the cerebellar anlage confirms our functional study.

### *Xenopus* represents a novel model for studying cerebellar development

Our ISH experiments provide a developmental map of GNP development in *Xenopus*, revealing that the processes leading to the emergence of URL derivatives and maturation of GN are similar to those seen in higher vertebrate ^37, 38^ reviewed in ^39, 40^. Amongst the important similarities i) the *Xenopus* URL expresses *atoh1*, is proliferative and is Tcf responsive; ii) *pax6* and *barhl1* are early markers of GNP commitment; iii) GNP migrate out of the URL along the cerebellar surface where they appear as the equivalent of the amniote-like EGL; iv) As in amniotes, postmitotic GN express *neurod1*, and migrate inwardly to form the IGL. Amongst the differences we observed i) a GNP caudal-to-rostral temporal differentiation gradient, with caudal URL differentiating first; ii) Although the EGL express stem/progenitor markers such as *atoh1,* the Notch pathway activity marker *hes4*, and *n-myc*, our proliferation analysis confirmed the absence of proliferating cells within the EGL from earlier stages ^41^; iii) Although *Barhl2* is expressed in the amniote EGL ^72^, we could not detect this transcription factor in the *Xenopus* cerebellar anlage. Our observations suggest that the tetrapod vertebrate *Xenopus*, the only described anamniote displaying an EGL, could be an alternative useful model for some clinical evaluation of cerebellar developmental defects, especially those related to early cerebellar development 41 reviewed in ^39, 40^.

### Tcf transcriptional activity is strictly necessary for *atoh1* expression and URL induction

Our data show that within R1, TCF transcriptional activity is strictly necessary for induction of *atoh1* expression and of the URL territory. Moreover, TCF inhibition is associated with an increase and/or an acceleration of GNP commitment/differentiation. Interestingly, studies performed in mouse neuroblastoma, and neural progenitor cells in culture, identified two TCF/LEF binding sites present in the 3’ enhancer region of *Atoh1* that are required for *Atoh1* activation ^73^. In these cells, the concomitant inhibition of Notch signaling and activation of WNT/Tcf, appear to be required for atoh1 expression ^73^. In mice low levels of Notch activity are necessary to induce a glutamatergic cell fate in *Sox2*-expressing cerebellar progenitors ^74^. Whereas it remains to be demonstrated in amniotes, our data argue that concomitant TCF activation and Notch inhibition are responsible for *atoh1* expression in the cerebellar primordium

We did not investigate which Tcf/Lef isoforms are transcriptionally active in the amphibian URL. However, our transcriptomic data reveal that *lef1 is* one of the most up-regulated DEG in the absence of *barhl1*, suggesting that it is present in the URL and could mediates Wnt signaling in this germinative niche. Three of the the four Tcf isoforms (Tcf7l2, Tcf7 and Lef1), mostly act as transcriptional activators, whereas the fourth (Tcf7l1) mostly acts as a transcriptional repressor reviewed in ^2^. Transcriptomic analyses of human cerebellar development reveal that the transcriptional activator TCF7 is active in the human URL ^32^, whereas Tcf7l2 is detected in the mouse URL ^33^. In both species, Tcf7l1 is associated with differentiated GNs ^25, 32^.

It is well documented that inhibition of TCF7l1-mediated repression is at the core of mouse embryonic stem cell (ESC) self-renewal and pluripotency. In contrast, enhancement of TCF7l1 repressive activity blocks mESC self-renewal, and allows mESC to differentiate, even in the presence of Wnt signaling ^75–82^ reviewed in ^3, 12, 83^. In adult mice, canonical WNTs are produced by both NSCs and astrocytes, and WNT/ß-catenin signalling stimulates both NSC self-renewal and neural progenitor cell proliferation 4–7,9 reviewed in 8. At least up to stage 50, we observed that the entire amphibian URL is Tcf active whereas the VZ is not. Early observation using electronic microscopy reported that during the premetamorphic phase, the cerebellum remains in an immature state and that a well-defined EGL up to 8 cells layers is likely to be established by the end of the prometamorphic phase ^38^. Taken together these observations indicate that the cerebellar URL displays features of an adult NSC niche. Our data provide new *in vivo* evidence that Tcf activity is strictly necessary for NSC niche maintenance and function.

### In the URL, *barhl1* promotes GNP exit from their germinative niche, towards commitment and differentiation

R1 territory is correctly established in embryos either lacking or overexpressing *barhl1,* arguing that Barhl1 is not involved in the establishment of the cerebellar anlage. Yet both our functional and transcriptomic data show that Barhl1 activity is strictly necessary for development of URL- derived GNPs. Whereas Barhl1 overexpression decreases the size of the URL, and promotes GNP commitment/differentiation, MO-mediated depletion of Barhl1 induces an enlargement of the URL associated with a marked delay in the GNP differentiation process.

Our transcriptomic analysis is consistent with our phenotypic observations. Barhl1-depleted DEG identification reveals that most significant upregulated genes regulate URL cell behavior either by acting on the fine equilibrium between a proliferative state and commitment and / or in maintenance of their stem/progenitor features. Indeed, *dmrta2* (*dmrt5)* expression is specific to neural stem/progenitor cells and has been shown to maintain NSC self-renewing ability ^84^. In neural progenitor cells derived from mESC, Dmrta2 maintains proliferation by binding to a target of Notch signaling, *Hes1,* and upregulates its expression, which will further inhibit neuronal differentiation through repressing the transcription of proneural genes ^84^. In the rodent developing and adult brain, the primary function of the orphan nuclear receptor *Nr2e1* (also known as *Tlx*) is to maintain NSC pools in an undifferentiated, self-renewing state preventing their premature differentiation ^85–88^ reviewed in ^89, 90^. In mice, *otx2* is expressed in GNPs during their massive expansion in the EGL ^91^, and its expression is associated with the high proliferation rate of GNP ^92^. However, the exact role of *otx2* in GNP development has not yet been elucidated. Finally, another upregulated URL target gene is *zic3*. Although *zic3* activity in the URL has not been described in mice, *zic3* is involved in maintaining pluripotency in both ESC ^93, 94^, and neural progenitor cells ^95^.

On the other hand, most downregulated DEG are involved in terminal neuronal differentiation, including dendrite development, and axonogenesis. One example is *Bhlhe2*, which in mice is expressed in the inner EGL, and is a downstream target of *Neurod1* in migrating and in differentiated GNs ^61^. *In vitro* KD experiments in primary GN culture indicate that *bhlhe22* is a regulator of post-mitotic GN radial migration towards the IGL ^96^.

### In the cerebellar primordium Barhl1 acts through repression of Tcf transcriptional activity

Similar to what we previously described for Barhl2 ^52^, in mammalian cells Barhl1 physically interacts with both Gro/Tle and Tcf7l1. In R1, Barhl1 overexpression phenocopies inhibition of Tcf transcriptional activity, and decreases Tcf activity. Conversely, Barhl1 depletion dramatically increases Tcf transcriptional activity in the URL. Both the increase in URL length, and the delay in GNP commitment/differentiation induced by Barhl1 depletion, are compensated by co-expression of a constitutive inhibitory form of Tcf7l1. Finally, Barhl1-KD embryos display a massive increase of Wnt activity throughout the cerebellar anlage.

Over 75% of Barhl1 depleted DEG regulatory regions contain at least one Tcf CRM; these include markers of the URL and EGL including *zic3*, *hes5* family genes, and wls. In line with our data in the rodent telencephalon, *Dmrta2* is transcriptionally activated by a stabilized form of beta-catenin and inhibited by a dominant-negative form of TCF ^86^. Tcf7l1 directly represses transcription of Lef1, which is stimulated by Wnt/β-catenin activity ^69^. These data argue that Barhl1 drives GNP out of the URL via Tcf-mediated repression and that Barhl1 LOF and GOF phenotypes are, at least partly, the result of alteration of its inhibitory effect on Tcf transcriptional activity.

### Barhl1 activates Notch activity in the cerebellar primordium

*Hes4,* a marker of Notch activity and stemness in *Xenopus* ^53^, is detected at stage 48 in the VZ, the URL, and the EGL. Our RNA-seq analysis reveals that depletion of Barhl1 leads to a significant upregulation of Notch pathway activity. Among upregulated components of Notch signaling, we identified the bHLH TF *helt (*also known as *Heslike and Megane).* In mice *Helt* is expressed in undifferentiated neural progenitors where it acts as transcriptional repressor of proneural genes ^97, 98^. Similarly, Barhl1 depletion in R1 leads to upregulation of *hes5* genes, which are known to inhibit neuronal differentiation by directly repressing proneural genes. In rodents, levels of Notch activity regulate the early progenitor choice between inhibitory (Notch +) and excitatory GN (Notch -) fate in the VZ ^74^. In agreement with its described function in maintaining cells in a primitive state, Notch has been suggested to prevent early GNP differentiation ^99^ reviewed in ^100, 101^, yet its exact function in developing GNP is still debated. Overexpression of a dominant negative form of Barhl2, which binds to Gro/Tle and counters its inhibitory activities, increases the URL/EGL size at the expense of GNP commitment/differentiation. Gro/Tle acts as a corepressor of both TCF and Enhancer of split E(spl), a major transcriptional repressor of Notch target gene activation, including proneural genes ^102^ reviewed in ^3^. Our findings suggest that there may be as yet unknown crosstalk between Wnt/Tcf and Notch signaling pathways in the maintenance of the cerebellar URL/EGL reviewed in ^103^.

### *Barhl* genes in amphibian versus mammalian cerebellar development

In mice, *Barhl*1 and *Barhl2* transcripts are detected in the outer URL and the posterior EGL from E11.5 onwards ^32, 45, 48, 50, 72, 104, 105^. scRNA-sequencing analysis of mouse cerebellar cells reveals that *Barhl1* is associated with early GNP differentiation, whereas *Barhl2* expression is uniquely associated with early fate commitment in the *Atoh1* lineage ^33^. Barhl1 and Barhl2 are highly conserved through evolution reviewed in ^3^, and their functional conservation is evidenced through studies in various species, including mouse, *C. elegans* and the acorn worm *Saccoglossus kowalevskii* ^106, 107^. Our data demonstrate that in the amphibian URL, Barhl1 mostly acts through inhibiting TCF activity. It remains to be investigated which of the *Barhl* TF inhibits TCF activity in mammals.

### Biological significance of our findings

Our study reveals previously undescribed roles for TCF and Barhl1 in the early development of URL-derived GNPs. We show that Barhl1 is the main repressor of Wnt/TCF activity in this germinative area. Our analysis reveals a set of Barhl1 target genes and opens the way for further characterization of relevant targets in order to create a global picture of GNP development and for further investigations of their relevance in adult NSC niche biology.

## ACKNOWLEDGEMENTS

The authors gratefully acknowledge M. Perron, D. Turner, P. Kreig, J. Christian, R. Vignali and G. Schlosser for gifts of materials and S. Authier and M. Abdelkrim for animal care. We thank the IBPS Aquatic platform supported by Sorbonne Université, CNRS, IBISA and the Conseil Régional Ile-de-France, specifically S. Authier and M. Abdelkrim, for Xenopus care. We thank C. Antoniewski (ArtBio platform, FR3631 IBPS) for tutoring in RNAseq data analysis and the team of G. Almouzni (Institut Curie) for providing high sensitivity RNA screentape system. We acknowledge Ann Lohof and Rachel Sherrard for editing work on the manuscript. This work was supported by the Centre National de la Recherche Scientifique (CNRS - UMR7622), donors to B Durand project, Sorbonne-Université, the Ligue Nationale contre le Cancer Comité Ile de France (RS19/75-52 and RS23/75-68). The CNRS supports Béatrice Durand and Mohamed Doulazmi. Alexis Eschstruth is supported by Sorbonne Université, and Johnny Bou- Rouphael by a fellowship from the “Ministère de la Recherche”.

## AUTHOR CONTRIBUTIONS

Conceptualization, J.BR., and B.C.D.; Methodology, J.BR., M.D., A.E., and B.D., Software, M.D.; Investigation, J.BR., M.D., A.E., A.A., and B.C.D.; Data curation, J.BR., and M.D., Writing – Original Draft, J.BR., and B.C.D.; Writing — Review & Editing, J.BR., M.D., A.E., and B.C.D.; Visualisation, J.BR., and B.C.D.; Funding Acquisition, B.C.D.; Resources, J.BR., M.D., A.E., and B.C.D.; Supervision, B.C.D.;

## DECLARATION OF INTERESTS

The authors declare no competing interests.

## STAR METHODS

### EXPERIMENTAL MODEL

#### *Xenopus* embryos care and husbandry

*X. laevis* embryos were obtained by conventional methods of hormone-induced egg laying and *in vitro* fertilization and were staged according to ^108^. *X. tropicalis* transgenic Wnt reporter pbin7LefdGFP has been generated as previously described ^22, 56, 57^. Briefly, the synthetic Wnt- responsive promoter consists of 7 copies of TCF/LEF1 binding sites and a TATA box driving destabilized green fluorescent protein (eGFP) and a polyA sequence. *gfp* expression reveals Wnt/TCF activity. *X. tropicalis* embryos were obtained by *in vitro* fertilization. Experimental procedures were specifically approved by the ethics committee of the Institut de Biologie Paris Seine (IBPS) (Authorization 2020-22727 given by CEEA #005) and have been carried out in strict accordance with the European community directive guidelines (2010/63/UE). B.D. carries the authorization for vertebrates’ experimental use N°75-1548.

### METHOD DETAILS

#### Plasmids design and preparation

*mbarhl1-HA-GR* contains the full-length *mbarhl1* sequence (two Engrailed-Homology (EH1) motifs, Nuclear Localization Signal (NLS); Homeodomain HD); and the C-terminal part), followed by an HA tag at the C-terminal part. This construct is inducible as it contains a glucocorticoid receptor which can be activated by dexamethasone (10uM). Dexamethasone- inducible *mbarhl2-EHs-GR* contains the first 182 amino acids (a.a) of mouse Barhl2 full-length cDNA, which correspond to the N-terminal EHs Gro-binding domains, and has been shown to act as a dominant negative ^52^. The full-length *mbarhl1-HA-GR* and truncated *mbarhl2-EHs-GR* constructs were generated in pCS2+ by Vector Builder. Non-inducible *mbarhl1-Myc* and *xbarhl1-Flag* were generated by GeneScript. Peptide sequences of the tags used are the following: HA (YPYDVPDYA); FLAG (DYKDDDDK) and MYC (EQKLISEEDL). The constitutive repressor pCS2-Tcf7l1-Δβcat-GR was a gift from H. Clevers ^55^, and consists of the full-length Tcf7l1 lacking the β-catenin-binding domain (BCBD), which reinforces its repressive activity. Constructs used for immunoprecipitation assay are *pCS2+ mbarhl1-3xFlag-HA* which was generated by Vector Builder. It contains the full-length *mbarhl1* sequence followed by three Flag tags and one HA tag at the C-terminal part. *pCS2+ Myc-Tcf7l1, pCS2+ Flag-Gro4* and *pCS2+GroHA* have been previously described ^52^. All necessary sequences were obtained from NCBI database. Constructs were validated by western blot on extracts from injected embryos or cell lysates.

#### mRNA synthesis, morpholino oligonucleotides (MOs) and *Xenopus* injection

Capped messenger RNAs (mRNAs) were synthesized using the mMessage mMachine kit (Invitrogen) and resuspended in RNAse-free H2O. Antisense morpholino oligonucleotides (MOs) were generated by Gene Tools. ATG start-site *MObarhl1-1* and *MObarhl1-2* were designed to block initiation of xBarhl1 protein translation. The MO were designed in a region overlapping the translation initiation site, so that they do not recognize mouse *Barhl1* or *xbarhl2* mRNA (Fig. S3). To establish the specificity of the MO effect, we tested the ability of *MObarhl1- 1* and *MObarhl1-2* to specifically inhibit translation of *xbarhl1* mRNA. Flag-tagged *xbarhl1* (*xbarhl1-flag)* or myc-tagged *mBarhl1* (*mBarhl1-myc*) were co-injected with *MObarhl1-1*, or *MObarhl1-2*, or a control MO (*MOct*) (Fig. S2). Western blot analysis on extracts from injected embryos confirmed a *MObarhl1*-mediated dramatic decrease in *Xenopus* Barhl1 protein levels, while MOct had no effect. We also observed that *MObarhl1-1* did not decrease mBarhl1-myc protein levels (Fig. S2; S3). *MObarhl1-1* was used for both *X. laevis* and *X. tropicalis* as the mRNA sequence of *barhl1* is highly conserved between both species, more specifically in the region on which *MObarhl1* is hybridized. Standard control MO from gene tools was used in this study. MO sequences and doses are summarized in table 2.

*Xenopus* embryos were injected unilaterally in one dorsal blastomere at the four and eight-cell stage together with *gfp* as a tracer for phenotype analysis by *in situ* hybridizations (ISH), except for CRISPR/Cas9 genome editing and RNAseq analysis *(see corresponding sections in material and methods)*. MOs were heated for 10 min at 65°C before usage. Injected embryos were transferred into 3% Ficoll in 0.3X Marc’s Modified Ringer’s (MMR) buffer (stock solution: 1M NaCl, 20 mM KCl, 20 mM CaCl2, 10 mM MgCl2, 50 mM HEPES pH 7.4). 10nl of mRNA or MO solution was injected together with a tracer in *X. laevis* while 5nl were injected in *X. tropicalis*. In *X. laevis*, MOs or mRNAs were co-injected with *gfp* mRNA (100 pg). MOs or mRNAs were co-injected with *mcherry* (100 pg) in *X. tropicalis.* Concentration of injected mRNA and MOs per embryo have been optimized in preliminary experiments. The minimal mRNA or MO quantity that induced the specific phenotype without showing toxicity effects was used. For embryos injected with inducible constructs, half of the injected embryos were treated with 10μM dexamethasone at stage 35, while the other half were left untreated and served as control. All necessary *Xenopus* sequences were obtained from https://www.xenbase.org/entry/.

#### *in situ* hybridization

Embryos were staged according to ^108^, and collected at the desired stage, then fixed in PFA4% for 1-2 hours at room temperature and dehydrated in 100% MeOH. ISH were performed using digoxigenin (DIG)-labeled probes. Antisense RNA probes were generated for the following transcripts: *atoh1, barhl1, hes4, neurod1, pax6, n-myc, otx2, zic3, wnt2b, wnt8b* and *gfp* according to the manufacturer’s instructions (RNA Labeling Mix, Roche). pCS2-Gfp is a gift from David Turner (University of Michigan, Ann Arbor, MI, USA). pBSK+xBarhl1 is a gift from Roberto Vignali (Unità di Biologia Cellulare e dello Sviluppo, Pisa Italy). pCS2-Atoh1 is a gift from G. Schlosser (University of Galway, Ireland). pBSK+Wnt2b was a gift from S. Sokol (Icahn School of Medicine at Mount Sinai, NY, USA). pBSK+Wnt8b was a gift from J Christian (University of Utah, USA). ISH was processed following the protocol described by (El Yakoubi et al., 2012; Sena et al., 2019). DISH was processed as described by ^109^. For *X. laevis* embryos, following rehydration, the eyes and ectoderm overlying the anterior neural tube were removed, which allows to skip the further Proteinase K (PK) treatment. Dissections weren’t performed on *X. tropicalis* embryos which were treated with PK. In both cases, bleaching was carried out, and samples were incubated with the probes overnight. Alkaline phosphatase-conjugated anti- DIG or anti-FLUO antibodies (Roche) were incubated 3 hours at room temperature. Enzymatic activity was revealed using NBT/BCIP (blue staining) and INT/BCIP (red staining) substrates (Roche). Following ISH, post-fixation was carried out in PFA 4% and the neural tubes of control and injected *X. laevis* embryos were dissected in PBS-0.1% Tween and stored in 90% glycerol. *X. tropicalis* embryos were stored in PFA 4%. Dissected neural tubes or embryos were photographed on a Leica M165 FC microscope equipped with Leica DFC320 camera using the same settings to allow direct comparison. Dorsal and lateral views of the dissected neural tubes were photographed.

#### Immunofluorescence

Immunofluorescence was carried out as previously described ^51^. The entire brains of wild-type (WT) and MO-injected *X. laevis* embryos were carefully dissected and transferred into a tube containing PBS-0.1% Tween, where they were progressively permeabilized. Samples were incubated with primary antibody (anti-Phospho-Histone H3; Upstate Biotechnology Cat#06– 570; d1:500) at 4°C overnight. Cellular nuclei were stained with bisBenzimide (BB) (Sigma) which was added to the solution containing diluted secondary antibody (Alexa Fluor 488 donkey anti-rabbit IgG; Invitrogen; d1:500) and incubated at 4°C overnight. Neural tubes were captured on a Zeiss Axio Observer.Z1 microscope equipped with apotome. Acquisitions were taken using the Z stack tool from the most superficial layer to deeper layers.

#### Immunoprecipitation in transfected HEK293T cells

HEK293T cells were cultured in supplemented Dulbecco’s modified Eagle’s medium (DMEM) (Gibco). Cells were transfected with expression vectors for *pCS2-mbarhl1-3xFlag-HA; pCS2- mbarhl1-Myc; pCS2-Tcf7l1-Myc; pCS2-Gro-Flag and pCS2-Gro-HA* encoding tagged proteins using the Phosphate Calcium method. Plasmids coding for pCS2+ or pSK+ were used as a supplement to ensure that cells in different dishes were transfected with the same quantity of expression vectors and plasmids (a total of 2 μg). Thirty-six hours post-transfection, cells were harvested and lysed in ice-cold lysis buffer (20 mM Tris pH7.6, 150 mM NaCl, 1% Triton, 1 mM EDTA) supplemented with completeTM protease inhibitor (Roche). Cell lysates were centrifuged 15min at 14,000 rpm. Protein complexes were precipitated from the cell lysates with anti-c-Myc antibody (clone 9E10). Protein complexes were then precipitated with protein A-Sepharose beads (Sigma) pre-washed with lysis buffer. Immunoprecipitated proteins were eluted from protein A beads by heating beads in Laemmli sample loading buffer (BioRad).

#### Western blot

Western blot (WB) analysis was performed on protein extracts from injected/WT *Xenopus* embryos, and on extracts from transfected HEK923T cells. *Xenopus* embryos were injected with *mbarhl1HAGR, xbarhl1Flag, mBarhl1Myc, mBarhl2EHsGR* mRNA at the two-cells stage, targeting both blastomeres. Proteins were extracted at stage 10 with lysis buffer (10 mM Tris- HCl pH 7.5, 100 mM NaCl, 0.5%NP40, 5mM EDTA supplemented with a cocktail of protein inhibitors). WB was carried out using the conventional methods. Proteins were separated by 10% SDS-polyacrylamide gel and transferred to nitrocellulose membrane. Membranes were blocked using 5% milk and incubated with the corresponding primary and secondary antibodies diluted in 5% milk (summarized in tables 3 and 4). Proteins were detected with Western Lightning Plus-ECL (Perkin Elmer Life Sciences). Membrane stripping was carried out between two staining steps using stripping buffer (Thermo Scientific) for the removal of primary and secondary antibodies from the membranes. ChemiDoc MP Imaging System (BioRad) was used for imaging the blots.

#### CRISPR/Cas9

Three CRISPR target sites (*barhl1-1* : GAGTCGGACGAGGCCATGGAAGG), *barhl1-2* : ACCAGCTCTGTGCGACAGAATGG, *barhl1-3* : AGAGTTGGACTCCGGGCTGGAGG) cutting respectively at 2, 37 and 230 bp from the beginning of the coding sequence were selected for their high predicted specificity and efficiency using CRISPOR online tool (http://crispor.tefor.net/). Alt-R crRNA and tracrRNA were purchased from Integrated DNA Technologies (IDT, Coralville, IA, USA) and dissolved in duplex buffer (IDT) at 100µM each. cr:tracrRNA duplexes were obtained by mixing equal amount of crRNA and tracrRNA, heating at 95°C for five minutes and letting cool down to room temperature. gRNA:Cas9 RNP complex was obtained by incubating 1µL 30µM Cas9 protein (kindly provided by TACGENE, Paris, France) with 2µL cr:tracrRNA duplex in a final volume of 10 µL of 20mM Hepes-NaOH ph 7.5, 150mM KCl for 10 min at 28°C. *X. tropicalis* one-cell stage embryos were injected with 2nL of gRNA:Cas9 RNP complex solution and were cultured to the desired stage. For coinjection, the three complexes were mixed at equal quantity.

Single embryo genomic DNA was obtained by digesting for 1h at 55°C in 100 µL lysis buffer (100 mM Tris-Hcl pH 7.5, 1 mM EDTA, 250 mM NaCl, 0.2% SDS, 0.1 µg/µL Proteinase K), precipitating with 1 volume of isopropanol and resuspended in 100µL PCR-grade water. The region surrounding the sgRNA binding sites was amplified by PCR using *X. tropicalis Xt_barhl1_F* (CAGCTCCTCCGACTTTTGTG) as forward primer and *Xt_barhl1_R* (GTTGCCCGTTGCTGGAATAA) as reverse primer. CRISPR efficiency was assessed by T7E1 test ^110^ on mono-injected embryos and by detecting deleted fragments on coinjected embryos.

#### RNA-sequencing and data analysis

*X. laevis* embryos were injected with three different conditions: *MObarhl1-1*; *MObarhl1-2* and *MOct* in the two dorsal blastomeres at four cells stage. At stage 42, neural tubes were extracted in RNAse-free conditions, and the rhombomere 1 which includes the URL was carefully dissected. For each condition, three biological replicates were collected. Each replicate contains three rhombomeres, which was the optimal number to get the minimal RNA concentration required for this experiment (Total RNA concentration was ∼30ng per sample). Briefly, total RNA was extracted using the TRIzol reagent (ambion) according to the manufacturer’s instructions. The overall RNA quality was assessed using Agilent High Sensitivity RNA ScreenTape System. Samples with an RNA Integrity Number (RIN) > 9 were used for subsequent analysis.

Sequencing was performed using Illumina NovaSeq (paired-end sequencing) by Next Generation Sequencing Platform (NGS) (Institut Curie). RNAseq data processing was performed using Galaxy server of ARTBio platform (IBPS).

Data sets were aligned against the *X. laevis* v10.1 genome assembly downloaded with its corresponding annotation file from Xenbase ^111^. Alignment was made using two read mapping programs, STAR v2.7.8a ^112^ and HISAT2 v2.2.1 ^113^. Quality control checks were assessed using FastQC v0.73 ^114^ and summarized in a single report generated by MultiQC v1.9 ^115^. As both alignment programs provided comparable results, we proceeded with STAR alignment tool. The number of aligned reads was counted by featurecounts tool v2.0.1 ^116^. Finally, we used the DESeq2 v2.11.40.6 package ^117^ to determine differentially expressed genes (DEG) from count tables. In the present study, genes with adjusted p value pAdj<0.001 were selected as significant DEG. Venn diagrams were produced with JVenn v2021.05.12 ^118^. Volcano Plots v0.0.5 were generated to show significant upregulated and downregulated genes, only a selection of DEG names were represented.

Further analysis and data visualization were performed using R v4.2.1package. A heatmap was generated to visualize gene expression across the samples. To overcome the lack of

*Xenopus* gene ontology (GO) annotation, we replaced *X. laevis* gene symbols with the Human orthologs. Functional enrichment analysis was performed using the *compareCluster* function of ClusterProfiler v4.8.1 ^56^ to identify GO-term enrichment amongst DEG with pAdj<0.001 as threshold. It provides the biological processes, cellular components, and molecular functions of DEG and compares each of the three subgroups between both knockdown conditions.

### QUANTIFICATION AND STATISTICAL ANALYSIS

#### Image processing and analysis

For ISH performed on embryos injected unilaterally, comparison of the expression levels between injected and control sides was assessed using a specific macro from ImageJ v2.1.0/1.53c ^119, 120^. The macro functions based on the RGB color mode. RGB images are split into three channels (red, green, and blue) and pixel values corresponding only to the blue channel are recorded, excluding the red and green channels, since the signal recorded on the blue channel represents the expression levels. For each image, the region of interest (ROI) was specified, and its dimensions were fixed, such that the same ROI is placed on the control and injected side of the embryo which prevents any subjectivity in ROI determination. Measured are the area corresponding to the blue signal; the mean or average value of signal within the selected ROI; and the integrated density which is the equivalent of the product of area and mean, as it sums the values of pixels in the selection. In this study, ratio of integrated density measured in the injected *versus* control side was assessed. The macro is available from the authors upon request and will be available as a plug-in in ImageJ.

The same macro was used for the analysis of CRISPR/Cas9-injected embryos, except that ROI was placed in all the rhombomere 1 as the entire embryo was targeted. The mean of int. density values of control embryos was compared to each individual int. density value of control and injected embryo. Phenotype penetrance was evaluated by counting and classifying embryos based on the intensity of *gfp* expression increase.

For immunofluorescence, Z-stack images were reconstructed and processed using ImageJ v2.1.0/1.53c. PHH3-positive cells were counted, and the length of the RL was measured on the control and injected side. Ratio of PHH3-positive cells and RL length in the injected *versus* control side was measured.

For the same experiment, all images were acquired using the same magnification and camera settings. In this way, all images were processed in a standardized manner, such that results are objectively analyzed. Final images were processed with Adobe Photoshop (v24.00).

#### Statistical analysis

Three independent experiments were performed for each condition analyzed. Dissected neural tubes and embryos were analyzed individually, and the results were pooled for data representation. Statistical analyses were implemented with R. Normality in the variable distributions was assessed by the Shapiro-Wilk test. Furthermore, the Levene test was performed to probe homogeneity of variances across groups. Variables that failed the Shapiro- Wilk or the Levene test were analyzed with non-parametric statistics using the one-way Kruskal-Wallis analysis of variance on ranks followed by Nemenyi test post hoc and Mann-

Whitney rank sum tests for pairwise multiple comparisons. Variables that passed the normality test were analyzed by means of one-way ANOVA followed by Tukey post hoc test for multiple comparisons or by Student’s *t* test for comparing two groups. A *p-*value of <0.05 was used as a cutoff for statistical significance. Results are presented as the means ± SEM. The statistical tests are described in each figure legend.

## Notes

### Competing Interest Statement

The authors have declared no competing interest.

## REFERENCES

1. Hoppler S, Waterman ML. Evolutionary Diversification of Vertebrate TCF/LEF Structure, Function, and Regulation. In: Hoppler S, Moon RT, éditeurs. Wnt Signaling in Development and Disease [Internet]. Hoboken, NJ, USA: John Wiley & Sons, Inc; 2014 [cité 24 sept 2021]. p. 225-37. Disponible sur: https://onlinelibrary.wiley.com/doi/10.1002/9781118444122.ch17

2. Torres-Aguila NP, Salonna M, Hoppler S, Ferrier DE. Evolutionary diversification of the canonical Wnt signaling effector TCF/LEF in chordates. Development, Growth & Differentiation. 2022;64(3):120.

3. Bou-Rouphael J, Durand BC. T-Cell Factors as Transcriptional Inhibitors: Activities and Regulations in Vertebrate Head Development. Frontiers in Cell and Developmental Biology [Internet]. 2021;9. Disponible sur: https://www.frontiersin.org/article/10.3389/fcell.2021.784998

4. Bowman AN, van Amerongen R, Palmer TD, Nusse R. Lineage tracing with Axin2 reveals distinct developmental and adult populations of Wnt/ -catenin-responsive neural stem cells. Proceedings of the National Academy of Sciences. 30 avr 2013;110(18):7324-9.

5. Kalamakis G, Brüne D, Ravichandran S, Bolz J, Fan W, Ziebell F, et al. Quiescence modulates stem cell maintenance and regenerative capacity in the aging brain. Cell. 2019;176(6):1407–19.

6. Qu Q, Sun G, Murai K, Ye P, Li W, Asuelime G, et al. Wnt7a regulates multiple steps of neurogenesis. Molecular and cellular biology. 2013;33(13):2551–9.

7. Lie DC, Colamarino SA, Song HJ, Désiré L, Mira H, Consiglio A, et al. Wnt signalling regulates adult hippocampal neurogenesis. Nature. 2005;437(7063):1370-5.

8. Urbán N, Blomfield IM, Guillemot F. Quiescence of adult mammalian neural stem cells: a highly regulated rest. Neuron. 2019;104(5):834–48.

9. García-Corzo L, Calatayud-Baselga I, Casares-Crespo L, Mora-Martínez C, Julián Escribano-Saiz J, Hortigüela R, et al. The transcription factor LEF1 interacts with NFIX and switches isoforms during adult hippocampal neural stem cell quiescence. Frontiers in Cell and Developmental Biology. 2022;1480.

10. Nusse R. Wnt signaling and stem cell control. Cell research. 2008;18(5):523–7.

11. Ding WY, Huang J, Wang H. Waking up quiescent neural stem cells: Molecular mechanisms and implications in neurodevelopmental disorders. PLoS genetics. 2020;16(4):e1008653.

12. Sokol SY. Maintaining embryonic stem cell pluripotency with Wnt signaling. Development. 15 oct 2011;138(20):4341–50.

13. Carta I, Chen CH, Schott AL, Dorizan S, Khodakhah K. Cerebellar modulation of the reward circuitry and social behavior. Science. 2019;363(6424):eaav0581.

14. Deverett B, Koay SA, Oostland M, Wang SS. Cerebellar involvement in an evidence- accumulation decision-making task. elife. 2018;7.

15. Wagner MJ, Kim TH, Savall J, Schnitzer MJ, Luo L. Cerebellar granule cells encode the expectation of reward. Nature. 2017;544(7648):96-100.

16. Hoshino M, Nakamura S, Mori K, Kawauchi T, Terao M, Nishimura YV, et al. Ptf1a, a bHLH transcriptional gene, defines GABAergic neuronal fates in cerebellum. Neuron. 2005;47(2):201–13.

17. Pascual M, Abasolo I, Mingorance-Le Meur A, Martínez A, Del Rio JA, Wright CV, et al. Cerebellar GABAergic progenitors adopt an external granule cell-like phenotype in the absence of Ptf1a transcription factor expression. Proceedings of the National Academy of Sciences. 2007;104(12):5193–8.

18. Yamada M, Seto Y, Taya S, Owa T, Inoue YU, Inoue T, et al. Specification of spatial identities of cerebellar neuron progenitors by ptf1a and atoh1 for proper production of GABAergic and glutamatergic neurons. Journal of Neuroscience. 2014;34(14):4786–800.

19. Wingate RJT. The rhombic lip and early cerebellar development. Current Opinion in Neurobiology. 1 févr 2001;11(1):82-8.

20. Wang VY, Rose MF, Zoghbi HY. Math1 expression redefines the rhombic lip derivatives and reveals novel lineages within the brainstem and cerebellum. Neuron. 2005;48(1):31–43.

21. Ben-Arie N, Bellen HJ, Armstrong DL, McCall AE, Gordadze PR, Guo Q, et al. Math1 is essential for genesis of cerebellar granule neurons. Nature. 1 nov 1997;390(6656):169-72.

22. Borday C, Parain K, Thi Tran H, Vleminckx K, Perron M, Monsoro-Burq AH. An atlas of Wnt activity during embryogenesis in Xenopus tropicalis. Schubert M, éditeur. PLoS ONE. 19 avr 2018;13(4):e0193606.

23. Garbe DS, Ring RH. Investigating Tonic Wnt Signaling Throughout the Adult CNS and in the Hippocampal Neurogenic Niche of BatGal and Ins-TopGal Mice. Cell Mol Neurobiol. oct 2012;32(7):1159–74.

24. Selvadurai HJ, Mason JO. Wnt/β-catenin Signalling Is Active in a Highly Dynamic Pattern during Development of the Mouse Cerebellum. Gottardi C, éditeur. PLoS ONE. 8 août 2011;6(8):e23012.

25. Wizeman JW, Guo Q, Wilion EM, Li JY. Specification of diverse cell types during early neurogenesis of the mouse cerebellum. eLife [Internet]. 8 févr 2019 [cité 8 mars 2021];8:e42388. Disponible sur: https://elifesciences.org/articles/42388

26. Gibson P, Tong Y, Robinson G, Thompson MC, Currle DS, Eden C, et al. Subtypes of medulloblastoma have distinct developmental origins. Nature. 1 déc 2010;468(7327):1095–9.

27. Xing L, Anbarchian T, Tsai JM, Plant GW, Nusse R. Wnt/β-catenin signaling regulates ependymal cell development and adult homeostasis. Proceedings of the National Academy of Sciences. 2018;115(26):E5954–62.

28. Lorenz A, Deutschmann M, Ahlfeld J, Prix C, Koch A, Smits R, et al. Severe alterations of cerebellar cortical development after constitutive activation of Wnt signaling in granule neuron precursors. Molecular and cellular biology. 2011;31(16):3326–38.

29. Pei Y, Brun SN, Markant SL, Lento W, Gibson P, Taketo MM, et al. WNT signaling increases proliferation and impairs differentiation of stem cells in the developing cerebellum. Development. 2012;139(10):1724–33.

30. Selvadurai HJ, Mason JO. Activation of Wnt/β-catenin signalling affects differentiation of cells arising from the cerebellar ventricular zone. 2012;

31. Flora A, Klisch TJ, Schuster G, Zoghbi HY. Deletion of Atoh1 disrupts Sonic Hedgehog signaling in the developing cerebellum and prevents medulloblastoma. Science [Internet]. 4 déc 2009 [cité 17 juin 2021];326(5958):1424-7. Disponible sur: https://www.ncbi.nlm.nih.gov/pmc/articles/PMC3638077/

32. Aldinger KA, Thomson Z, Phelps IG, Haldipur P, Deng M, Timms AE, et al. Spatial and cell type transcriptional landscape of human cerebellar development. Nature Neuroscience. 1 août 2021;24(8):1163-75.

33. Carter RA, Bihannic L, Rosencrance C, Hadley JL, Tong Y, Phoenix TN, et al. A Single- Cell Transcriptional Atlas of the Developing Murine Cerebellum. Current Biology. sept 2018;28(18):2910–2920.e2.

34. Hanzel M, Rook V, Wingate RJ. Mitotic granule cell precursors undergo highly dynamic morphological transitions throughout the external germinal layer of the chick cerebellum. Scientific reports. 2019;9(1):1–13.

35. Machold R, Fishell G. Math1 is expressed in temporally discrete pools of cerebellar rhombic-lip neural progenitors. Neuron. 2005;48(1):17–24.

36. Miyata T, Maeda T, Lee JE. NeuroD is required for differentiation of the granule cells in the cerebellum and hippocampus. Genes & development. 1999;13(13):1647–52.

37. Herrick CJ. The cerebellum of Necturus and other urodele Amphibia. Journal of Comparative Neurology. 1914;24(1):1–29.

38. Gona AG. Morphogenesis of the cerebellum of the frog tadpole during spontaneous metamorphosis. Journal of Comparative Neurology. 1972;146(2):133–42.

39. Hibi M, Matsuda K, Takeuchi M, Shimizu T, Murakami Y. Evolutionary mechanisms that generate morphology and neural-circuit diversity of the cerebellum. Development, growth & differentiation. 2017;59(4):228–43.

40. Miyashita S, Hoshino M. Transit Amplifying Progenitors in the Cerebellum: Similarities to and Differences from Transit Amplifying Cells in Other Brain Regions and between Species. Cells. 2022;11(4):726.

41. Butts T, Hanzel M, Wingate RJ. Transit amplification in the amniote cerebellum evolved via a heterochronic shift in NeuroD1 expression. Development. 2014;141(14):2791–5.

42. D’Amico LA, Boujard D, Coumailleau P. The Neurogenic Factor NeuroD1 Is Expressed in Post-Mitotic Cells during Juvenile and Adult Xenopus Neurogenesis and Not in Progenitor or Radial Glial Cells. PLOS ONE. 14 juin 2013;8(6):e66487.

43. Kojima T, Ishimaru S, Higashijima S ichi, Takayama E, Akimaru H, Sone M, et al. Identification of a different-type homeobox gene, BarH1, possibly causing Bar (B) and Om (1D) mutations in Drosophila. Proceedings of the National Academy of Sciences. 1991;88(10):4343-7.

44. Higashijima S ichi, Kojima T, Michiue T, Ishimaru S, Emori Y, Saigo K. Dual Bar homeo box genes of Drosophila required in two photoreceptor cells, R1 and R6, and primary pigment cells for normal eye development. Genes & Development. 1992;6(1):50-60.

45. Kawauchi D, Saito T. Transcriptional cascade from Math1 to Mbh1 and Mbh2 is required for cerebellar granule cell differentiation. Developmental Biology. oct 2008;322(2):345–54.

46. Klisch TJ, Xi Y, Flora A, Wang L, Li W, Zoghbi HY. In vivo Atoh1 targetome reveals how a proneural transcription factor regulates cerebellar development. Proceedings of the National Academy of Sciences. 2011;108(8):3288–93.

47. Reig G, Cabrejos ME, Concha ML. Functions of BarH transcription factors during embryonic development. Developmental Biology. févr 2007;302(2):367–75.

48. Bulfone A. Barhl1, a gene belonging to a new subfamily of mammalian homeobox genes, is expressed in migrating neurons of the CNS. Human Molecular Genetics. 22 mai 2000;9(9):1443-52.

49. Li S, Qiu F, Xu A, Price SM, Xiang M. Barhl1 Regulates Migration and Survival of Cerebellar Granule Cells by Controlling Expression of the Neurotrophin-3 Gene. Journal of Neuroscience. 2004;24(12):3104–14.

50. Li S. Barhl1 Regulates Migration and Survival of Cerebellar Granule Cells by Controlling Expression of the Neurotrophin-3 Gene. Journal of Neuroscience. 24 mars 2004;24(12):3104-14.

51. Juraver-Geslin HA, Ausseil JJ, Wassef M, Durand BC. Barhl2 limits growth of the diencephalic primordium through Caspase3 inhibition of -catenin activation. Proceedings of the National Academy of Sciences. 8 févr 2011;108(6):2288-93.

52. Sena E, Rocques N, Borday C, Amin HSM, Parain K, Sitbon D, et al. Barhl2 maintains T-cell factors as repressors, and thereby switches off the Wnt/β-Catenin response driving Spemann organizer formation. Development. 1 janv 2019;dev.173112.

53. El Yakoubi W, Borday C, Hamdache J, Parain K, Tran HT, Vleminckx K, et al. Hes4 controls proliferative properties of neural stem cells during retinal ontogenesis. Stem Cells. 2012;30(12):2784–95.

54. Grbavec D, Lo R, Liu Y, Stifani S. Transducin-like Enhancer of split 2, a mammalian homologue of Drosophila Groucho, acts as a transcriptional repressor, interacts with Hairy/Enhancer of split proteins, and is expressed during neuronal development. Eur J Biochem. déc 1998;258(2):339–49.

55. Molenaar M, van de Wetering M, Oosterwegel M, Peterson-Maduro J, Godsave S, Korinek V, et al. XTcf-3 Transcription Factor Mediates β-Catenin-Induced Axis Formation in Xenopus Embryos. Cell. 9 août 1996;86(3):391-9.

56. Tran HT, Sekkali B, Van Imschoot G, Janssens S, Vleminckx K. Wnt/β-catenin signaling is involved in the induction and maintenance of primitive hematopoiesis in the vertebrate embryo. Proceedings of the National Academy of Sciences. 2010;107(37):16160–5.

57. Tran HT, Vleminckx K. Design and use of transgenic reporter strains for detecting activity of signaling pathways in Xenopus. Methods. 2014;66(3):422–32.

58. Wu T, Hu E, Xu S, Chen M, Guo P, Dai Z, et al. clusterProfiler 4.0: A universal enrichment tool for interpreting omics data. The Innovation. 28 août 2021;2(3):100141.

59. Eiraku M, Tohgo A, Ono K, Kaneko M, Fujishima K, Hirano T, et al. DNER acts as a neuron-specific Notch ligand during Bergmann glial development. Nature neuroscience. 2005;8(7):873–80.

60. Hsieh FY, Ma TL, Shih HY, Lin SJ, Huang CW, Wang HY, et al. Dner inhibits neural progenitor proliferation and induces neuronal and glial differentiation in zebrafish. Developmental biology. 2013;375(1):1–12.

61. Ma N, Puls B, Chen G. Transcriptomic analyses of NeuroD1-mediated astrocyte-to- neuron conversion. Developmental Neurobiology. 2022;82(5):375–91.

62. Chellappa R, Li S, Pauley S, Jahan I, Jin K, Xiang M. Barhl1 Regulatory Sequences Required for Cell-Specific Gene Expression and Autoregulation in the Inner Ear and Central Nervous System. Mol Cell Biol. 15 mars 2008;28(6):1905-14.

63. Kjolby RA, Truchado-Garcia M, Iruvanti S, Harland RM. Integration of Wnt and FGF signaling in the Xenopus gastrula at TCF and Ets binding sites shows the importance of short-range repression by TCF in patterning the marginal zone. Development. 2019;146(15):dev179580.

64. Nakamura Y, de Paiva Alves E, Veenstra GJ, Hoppler S. Tissue- and stage-specific Wnt target gene expression is controlled subsequent to β-catenin recruitment. Development. 1 janv 2016;dev.131664.

65. Aruga J. The role of Zic genes in neural development. Molecular and Cellular Neuroscience. 1 juin 2004;26(2):205-21.

66. Aruga J, Millen KJ. ZIC1 function in normal cerebellar development and human developmental pathology. Zic family. 2018;249–68.

67. Houtmeyers R, Souopgui J, Tejpar S, Arkell R. The ZIC gene family encodes multi- functional proteins essential for patterning and morphogenesis. Cellular and Molecular Life Sciences. 2013;70:3791–811.

68. Imayoshi I, Sakamoto M, Yamaguchi M, Mori K, Kageyama R. Essential roles of Notch signaling in maintenance of neural stem cells in developing and adult brains. Journal of Neuroscience. 2010;30(9):3489–98.

69. Wu CI, Hoffman JA, Shy BR, Ford EM, Fuchs E, Nguyen H, et al. Function of Wnt/β- catenin in counteracting Tcf3 repression through the Tcf3–β-catenin interaction. Development. 2012;139(12):2118–29.

70. Yeung J, Ha TJ, Swanson DJ, Choi K, Tong Y, Goldowitz D. Wls Provides a New Compartmental View of the Rhombic Lip in Mouse Cerebellar Development. J Neurosci. 10 sept 2014;34(37):12527.

71. Yeung J, Goldowitz D. Wls expression in the rhombic lip orchestrates the embryonic development of the mouse cerebellum. Neuroscience. 2017;354:30–42.

72. Mo Z, Li S, Yang X, Xiang M. Role of the *Barhl2* homeobox gene in the specification of glycinergic amacrine cells. Development. 1 avr 2004;131(7):1607-18.

73. Shi F, Cheng Y fu, Wang XL, Edge AS. β-catenin up-regulates Atoh1 expression in neural progenitor cells by interaction with an Atoh1 3′ enhancer. Journal of Biological Chemistry. 2010;285(1):392–400.

74. Zhang T, Liu T, Mora N, Guegan J, Bertrand M, Contreras X, et al. Generation of excitatory and inhibitory neurons from common progenitors via Notch signaling in the cerebellum. Cell Reports. 2021;35(10):109208.

75. Cole MF, Johnstone SE, Newman JJ, Kagey MH, Young RA. Tcf3 is an integral component of the core regulatory circuitry of embryonic stem cells. Genes & Development. 15 mars 2008;22(6):746-55.

76. Moreira S, Polena E, Gordon V, Abdulla S, Mahendram S, Cao J, et al. A Single TCF Transcription Factor, Regardless of Its Activation Capacity, Is Sufficient for Effective Trilineage Differentiation of ESCs. Cell Reports. sept 2017;20(10):2424–38.

77. Park MS, Kausar R, Kim MW, Cho SY, Lee YS, Lee MA. Tcf7l1-mediated transcriptional regulation of Krüppel-like factor 4 gene. Animal Cells and Systems. 2 janv 2015;19(1):16-29.

78. Pereira L, Yi F, Merrill BJ. Repression of Nanog Gene Transcription by Tcf3 Limits Embryonic Stem Cell Self-Renewal. Mol Cell Biol. 15 oct 2006;26(20):7479–91.

79. Salomonis N, Schlieve CR, Pereira L, Wahlquist C, Colas A, Zambon AC, et al. Alternative splicing regulates mouse embryonic stem cell pluripotency and differentiation. Proceedings of the National Academy of Sciences. 8 juin 2010;107(23):10514-9.

80. Tam WL, Lim CY, Han J, Zhang J, Ang YS, Ng HH, et al. T-Cell Factor 3 Regulates Embryonic Stem Cell Pluripotency and Self-Renewal by the Transcriptional Control of Multiple Lineage Pathways. Stem Cells. août 2008;26(8):2019–31.

81. Wray J, Kalkan T, Gomez-Lopez S, Eckardt D, Cook A, Kemler R, et al. Inhibition of glycogen synthase kinase-3 alleviates Tcf3 repression of the pluripotency network and increases embryonic stem cell resistance to differentiation. Nat Cell Biol. juill 2011;13(7):838–45.

82. Atlasi Y, Noori R, Gaspar C, Franken P, Sacchetti A, Rafati H, et al. Wnt Signaling Regulates the Lineage Differentiation Potential of Mouse Embryonic Stem Cells through Tcf3 Down-Regulation. Cadigan K, éditeur. PLoS Genet. 2 mai 2013;9(5):e1003424.

83. Wray J, Hartmann C. WNTing embryonic stem cells. Trends in Cell Biology. 1 mars 2012;22(3):159-68.

84. Young FI, Keruzore M, Nan X, Gennet N, Bellefroid EJ, Li M. The doublesex-related Dmrta2 safeguards neural progenitor maintenance involving transcriptional regulation of Hes1. Proceedings of the National Academy of Sciences. 2017;114(28):E5599–607.

85. Kandel P, Semerci F, Mishra R, Choi W, Bajic A, Baluya D, et al. Oleic acid is an endogenous ligand of TLX/NR2E1 that triggers hippocampal neurogenesis. Proceedings of the National Academy of Sciences. 29 mars 2022;119(13):e2023784119.

86. Konno D, Iwashita M, Satoh Y, Momiyama A, Abe T, Kiyonari H, et al. The mammalian DM domain transcription factor Dmrta2 is required for early embryonic development of the cerebral cortex. 2012;

87. Saulnier A, Keruzore M, De Clercq S, Bar I, Moers V, Magnani D, et al. The doublesex homolog Dmrt5 is required for the development of the caudomedial cerebral cortex in mammals. Cerebral Cortex. 2013;23(11):2552–67.

88. Urquhart J, Beaman G, Byers H, Roberts N, Chervinsky E, O’Sullivan J, et al. DMRTA2 (DMRT5) is mutated in a novel cortical brain malformation. Clinical Genetics. 2016;89(6):724–7.

89. Islam MM, Zhang CL. TLX: A master regulator for neural stem cell maintenance and neurogenesis. Biochimica et Biophysica Acta (BBA)-Gene Regulatory Mechanisms. 2015;1849(2):210–6.

90. Wang Y, Liu HK, Schütz G. Role of the nuclear receptor Tailless in adult neural stem cells. Mechanisms of Development. 1 juin 2013;130(6):388-90.

91. Fossat N, Chatelain G, Brun G, Lamonerie T. Temporal and spatial delineation of mouse Otx2 functions by conditional self-knockout. EMBO reports. 2006;7(8):824–30.

92. El Nagar S, Chakroun A, Le Greneur C, Figarella-Branger D, Di Meglio T, Lamonerie T, et al. Otx2 promotes granule cell precursor proliferation and Shh-dependent medulloblastoma maintenance in vivo. Oncogenesis. 13 août 2018;7(8):60.

93. Lim LS, Hong FH, Kunarso G, Stanton LW. The pluripotency regulator Zic3 is a direct activator of the Nanog promoter in ESCs. Stem cells. 2010;28(11):1961–9.

94. Lim LS, Loh YH, Zhang W, Li Y, Chen X, Wang Y, et al. Zic3 is required for maintenance of pluripotency in embryonic stem cells. Molecular biology of the cell. 2007;18(4):1348–58.

95. Inoue T, Ota M, Ogawa M, Mikoshiba K, Aruga J. Zic1 and Zic3 regulate medial forebrain development through expansion of neuronal progenitors. Journal of Neuroscience. 2007;27(20):5461–73.

96. Ramirez M, Badayeva Y, Yeung J, Wu J, Yang E, Trost B, et al. Temporal analysis of enhancers during mouse brain development reveals dynamic regulatory function and identifies novel regulators of cerebellar development. bioRxiv. 2021;

97. Nakatani T, Mizuhara E, Minaki Y, Sakamoto Y, Ono Y. Helt, a novel basic-helix-loop- helix transcriptional repressor expressed in the developing central nervous system. Journal of Biological Chemistry. 2004;279(16):16356–67.

98. Nakatani T, Minaki Y, Kumai M, Ono Y. Helt determines GABAergic over glutamatergic neuronal fate by repressing Ngn genes in the developing mesencephalon. 2007;

99. Solecki DJ, Liu X, Tomoda T, Fang Y, Hatten ME. Activated Notch2 signaling inhibits differentiation of cerebellar granule neuron precursors by maintaining proliferation. Neuron. 2001;31(4):557–68.

100. Cavallucci V, Fidaleo M, Pani G. Neural stem cells and nutrients: poised between quiescence and exhaustion. Trends in Endocrinology & Metabolism. 2016;27(11):756–69.

101. Hu N, Zou L. Multiple functions of Hes genes in the proliferation and differentiation of neural stem cells. Annals of Anatomy-Anatomischer Anzeiger. 2022;239:151848.

102. Cinnamon E, Helman A, Ben-Haroush Schyr R, Orian A, Jiménez G, Paroush Z. Multiple RTK pathways downregulate Groucho-mediated repression in *Drosophila* embryogenesis. Development. 1 mars 2008;135(5):829-37.

103. Muñoz Descalzo S, RuÉ P, Garcia-Ojalvo J, Arias AM. Correlations between the levels of Oct4 and Nanog as a signature for naïve pluripotency in mouse embryonic stem cells. Stem cells. 2012;30(12):2683–91.

104. Pöschl J, Lorenz A, Hartmann W, von Bueren AO, Kool M, Li S, et al. Expression of BARHL1 in medulloblastoma is associated with prolonged survival in mice and humans. Oncogene. nov 2011;30(47):4721–30.

105. Rachidi M, Lopes C. Differential transcription of Barhl1 homeobox gene in restricted functional domains of the central nervous system suggests a role in brain patterning. International journal of developmental neuroscience. 2006;24(1):35–44.

106. Schwartz HT, Horvitz HR. The C. elegans protein CEH-30 protects male-specific neurons from apoptosis independently of the Bcl-2 homolog CED-9. Genes & Development. 1 déc 2007;21(23):3181-94.

107. Yao Y, Minor PJ, Zhao YT, Jeong Y, Pani AM, King AN, et al. Cis-regulatory architecture of a brain signaling center predates the origin of chordates. Nat Genet. mai 2016;48(5):575–80.

108. Nieuwkoop P, Faber J. Normal table of Xenopus laevis (Daudin) garland publishing. New York. 1994;252.

109. Juraver-Geslin HA, Gómez-Skarmeta JL, Durand BC. The conserved barH-like homeobox-2 gene barhl2 acts downstream of orthodentricle-2 and together with iroquois-3 in establishment of the caudal forebrain signaling center induced by Sonic Hedgehog. Developmental Biology. 1 déc 2014;396(1):107-20.

110. Mashal RD, Koontz J, Sklar J. Detection of mutations by cleavage of DNA heteroduplexes with bacteriophage resolvases. Nature genetics. 1995;9(2):177–83.

111. Fortriede JD, Pells TJ, Chu S, Chaturvedi P, Wang D, Fisher ME, et al. Xenbase: deep integration of GEO & SRA RNA-seq and ChIP-seq data in a model organism database. Nucleic Acids Research. 2020;48(D1):D776–82.

112. Dobin A, Davis CA, Schlesinger F, Drenkow J, Zaleski C, Jha S, et al. STAR: ultrafast universal RNA-seq aligner. Bioinformatics. 2013;29(1):15–21.

113. Kim D, Langmead B, Salzberg SL. HISAT: a fast spliced aligner with low memory requirements. Nature methods. 2015;12(4):357–60.

114. Andrews S. FastQC: a quality control tool for high throughput sequence data. 2010. 2017;

115. Ewels P, Magnusson M, Lundin S, Käller M. MultiQC: summarize analysis results for multiple tools and samples in a single report. Bioinformatics. 2016;32(19):3047–8.

116. Liao Y, Smyth GK, Shi W. featureCounts: an efficient general purpose program for assigning sequence reads to genomic features. Bioinformatics. 2014;30(7):923–30.

117. Love MI, Huber W, Anders S. Moderated estimation of fold change and dispersion for RNA-seq data with DESeq2. Genome biology. 2014;15(12):1–21.

118. Bardou J. An interactive Venn diagram viewer. BMC Bioinformatics. (15).

119. Abràmoff MD, Magalhães PJ, Ram SJ. Image processing with ImageJ. Biophotonics international. 2004;11(7):36–42.

120. Schneider CA, Rasband WS, Eliceiri KW. NIH Image to ImageJ: 25 years of image analysis. Nature Methods. 1 juill 2012;9(7):671-5.

