## Supplemental Data for "Cerebellar granular neuron progenitors exit their germinative niche via Barhl1 mediated silencing of T-Cell Factor transcriptional activity"

### SUPPLEMENTARY FIGURES

**Figure S1. Granule neuron progenitors' development in *X. laevis***

Spatial and temporal expression of key markers of GNPs development in the cerebellar anlage of *X. laevis* at indicated stages (st.). Shown are (A-Ea) dorsal view of st. 38 and (A-E) dorsal (b) and lateral (b') views of st. 45 *X. laevis* neural tubes stained with indicated markers. (A) *atoh1* expression is detected in the URL (B) the proliferation marker *nmyc* is expressed in the URL, the VZ, and in proliferating progenitors at the boundaries between the rhombomeres (R). (C) *pax6*+ GNPs are first detected around st. 38 in amphibian. (C, D) *pax6* and *barhl1* mark the committed GNPs while (E) *neurod1* is expressed in differentiated GNs. (F) Dorsal views of st. 38, 45 and 48 showing absence of *barhl2* expression in the R1 of amphibian. Scale bar 150μm.

**Figure S2. Experimental procedure and constructs**

(A) Representation of the injection procedure. mRNAs of the different constructs used in this study (B) were co-injected with a tracer - *gfp* mRNA in *X. laevis* and *mcherry* mRNA in *X. tropicalis* - into one dorsal blastomere at the four/eight-cell stage embryo. Unilaterally injected embryos were selected. On the right is the image of a stage 48 *X. laevis* embryo injected with *gfp* as tracer. Injected and wild-type embryos were left to develop at 18°C. For embryos injected with inducible constructs, half of the injected embryos were treated with 10μM dexamethasone at stage 35, while the other half were untreated and used as control. Embryos were fixed at different stages and used for further analysis by ISH. (B) Schematic representation of the different constructs used in this study. Construct organization is indicated in the drawings. Tcf7l1-Δβcat-GR lacks the β-catenin-binding domain (BCBD) at its N-terminal region (yellow). Absence of this domain reinforces its repressive activity. It contains the DNA-binding domain which contains a High-Mobility Group box (HMG-box) (green) and a Nuclear Localization Signal (NLS) (grey). The DNA-binding domain is preceded by a less well-defined binding sequence for the Groucho/Transducin-like enhancer of split (Gro/TLE) (Gro-binding sequence, GBS) (red). The Context-dependent Regulatory Domain (CRD) is encoded by three exons and is flanked by two small motifs LVPQ shown in blue at its N-terminal end, and SxxSS shown in pink at its C terminal end. The long (E) tail of Tcf7l1 (light blue) contains two C-terminal-binding protein (CtBP) motifs (PLDLS) (purple). This construct is inducible and contains at its Carboxyl (C) terminal end a glucocorticoid receptor/ligand binding domain (GR/LBD). xBarhl1 and mBarhl1 are composed of the full length Barhl1 structure including both N-terminal Engrailed Homology motifs (EH1) (grey and orange), the NLS (light green) followed by the homeodomain (HD) (yellow). xBarhl1 is Flag-tagged. mBarhl1 is Myc tagged. mBarhl1-HA-GR is HA tagged and is inducible. Inducible mBarhl2EHs-GR contains the two EH domains and acts as dominant negative. Western blot was carried out on extract from embryos injected with the mBarhl1-HA-GR and mBarhl2EHs-GR constructs and their expression validated. Samples were separated by SDS-page and detected by immunoblotting with anti-Ha, anti-Flag, anti-Myc, or anti-Barhl2. On the right are shown the molecular weights in kDa. n-inj: non-injected.

**Figure S3. Morpholino-mediated depletion of xBarhl1 and its impact on GNPs development**

(Aa-c) *MObarhl1-1* and *MObarhl1-2* specifically block translation of *xbarhl1* mRNA. (a, b) Morpholino (MO) oligonucleotides were designed to target the translation initiation site of *X. laevis* and *X. tropicalis (XL/XT) barhl1* mRNA. *MObarhl1-1* and *MObarhl1-2* do not recognize mouse *Barhl1* and *X. laevis barhl2* mRNAs. *X. tropicalis* were injected with *MObarhl1-1.* Red characters indicate nucleotides that are not recognized by *MObarhl1-1* and *MObarhl1-2*. (c) Western blot on extracts from *X. laevis* embryos injected with flag-tagged *xbarhl1* (*xbarhl1-flag)* together with *MObarhl1-1*, or *MObarhl1-2*, or control MO *(MOct)*. myc-tagged *mBarhl1* (*mBarhl1-myc*) was co-injected with *MObarhl1-1. MObarhl1-1* and *MObarhl1-2* induced a dramatic decrease in xBarhl1 protein levels without affecting those of mBarhl1-myc, while *MOct* had no effect on xBarhl1 expression. Membranes were incubated with stripping buffer to eliminate primary anti-flag and secondary antibodies. Asterisk indicates post-stripping xBarhl1-flag corresponding band. Actin was used as loading control to confirm that levels of proteins loaded are equal across the gel. (B) Injection of *MOct* didn't induce any significant effect. *in situ* hybridization analysis of *atoh1, pax6* and *neurod1* expressions in stage 45 *X. laevis* embryos injected with *MOct (20ng).* (C-D) Analysis of *MObarhl1-1* effect at stage 41 and stage 48. ISH analysis of *X. laevis* embryos injected with *MObarhl1-1 (15ng)* report a significant increase in *atoh1* expression and a dramatic decrease in the commitment/differentiation markers *pax6* and *neurod1* at stage 41 and stage 48. Scale bar 150µm.

**Figure S4** (A) *in situ* hybridization analysis of *gfp* expression (Tcf activity) performed on *X. tropicalis* pbin7LefdGFP line injected with morpholino control (*MOct).* (B) *in situ* hybridization analysis showing decreased expression of *neurod1* and *pax6* in embryos unilaterally injected with *barhl2EHs-GR. inj:* injected side. Scale bar 150µm. (B) Efficiency of CRISPR/Cas9-mediated mutation of Barhl1 in *X. tropicalis* using T7E1 assay. Representative gel images displaying Polymerase Chain Reaction (PCR) products amplified from genomic DNA isolated from non-injected embryos (ninj) and from embryos injected (inj) with *CRISPRbarhl1-1; CRISPRbarhl1-2; CRISPRbarhl1-3* mixed (a) or each alone (b-c-d) and treated (+T7 digestion) or not (ct: control) with T7E1. PCR product size is 430bp in *ct* samples.

**Figure S5. Analysis of count data from RNAseq**
(A) Dispersion plot showing the dispersion estimates for each gene separately (black points), and the dispersions' dependence on the mean of normalized counts (red line). Final estimates are represented by blue points. The blue circles are genes which have high gene-wise dispersion estimates. (B, C, D) p-values plots showing non-significant difference between *MObarhl1-1* versus *MObarhl1-2* and significant difference between (C) *MObarhl1-1* versus *MOct* and (D) *MObarhl1-2* versus *MOct.* Analysis is provided by DESeq2 package using galaxy. (E) Venn-Diagram showing the distribution of DEGs between *MObarhl1-1 vs MOct* and *MObarhl1-2 vs MOct.* The numbers of DEGs with pAdj<0.001 exclusively expressed by each subset and genes overlapping between both conditions are indicated. Green represents *MObarhl1-1 vs MOct* and light red represents *MObarhl1-2 vs MOct.* Venn diagram is generated using Galaxy.

**Table S1. List of differentially expressed genes identified by RNAseq analysis**
The table shows the list of all DEG obtained from the RNAseq experiment for *MObarhl1-1* and *MObarhl1-2* conditions, each compared to *MOcontrol (MOct)*. Gene counts are obtained using FeatureCounts. Statistically significant DEG with pAdj≤0.001 are included in the table. DEG in common between both conditions are also represented.

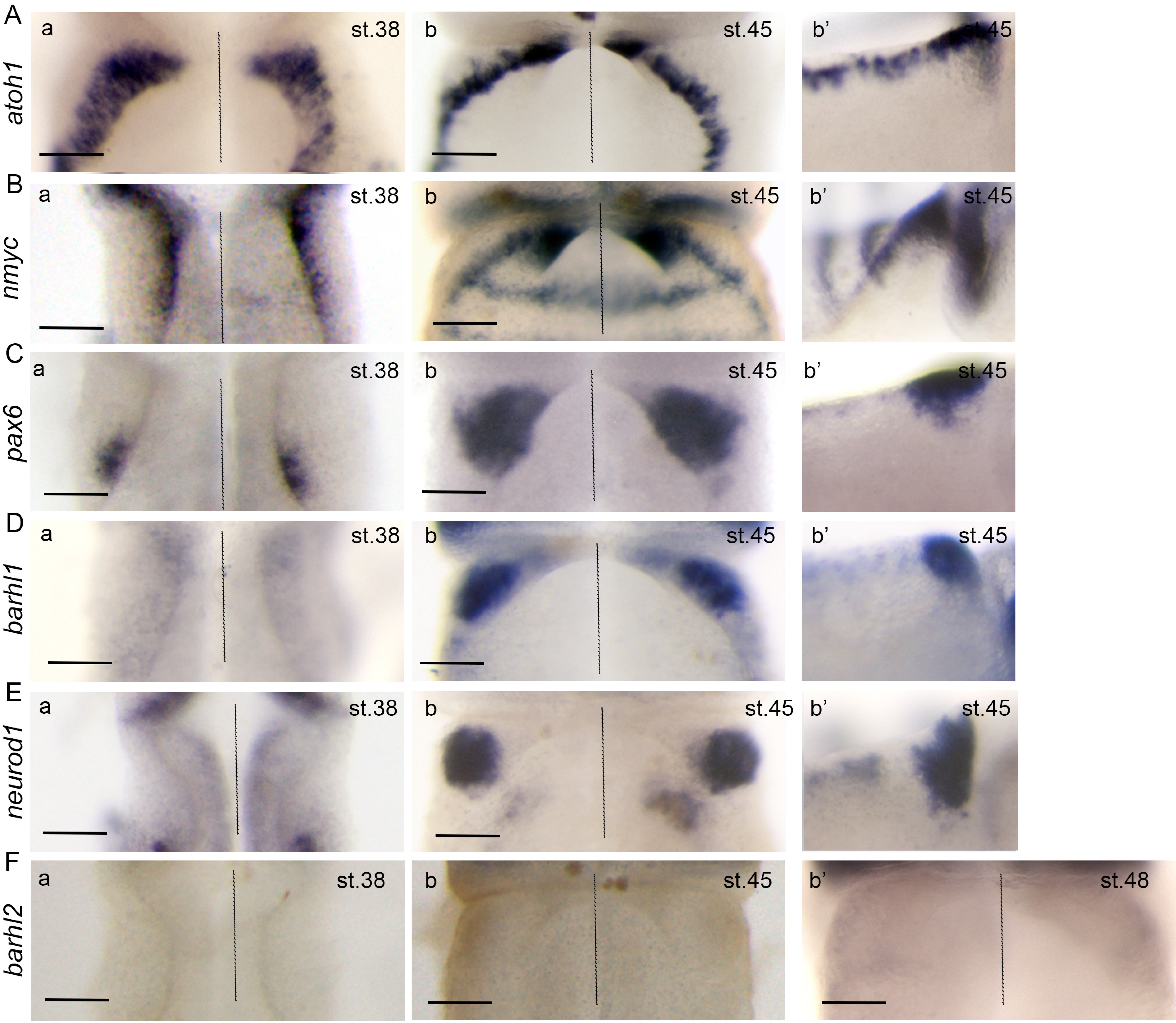


Supplementary Figure 1

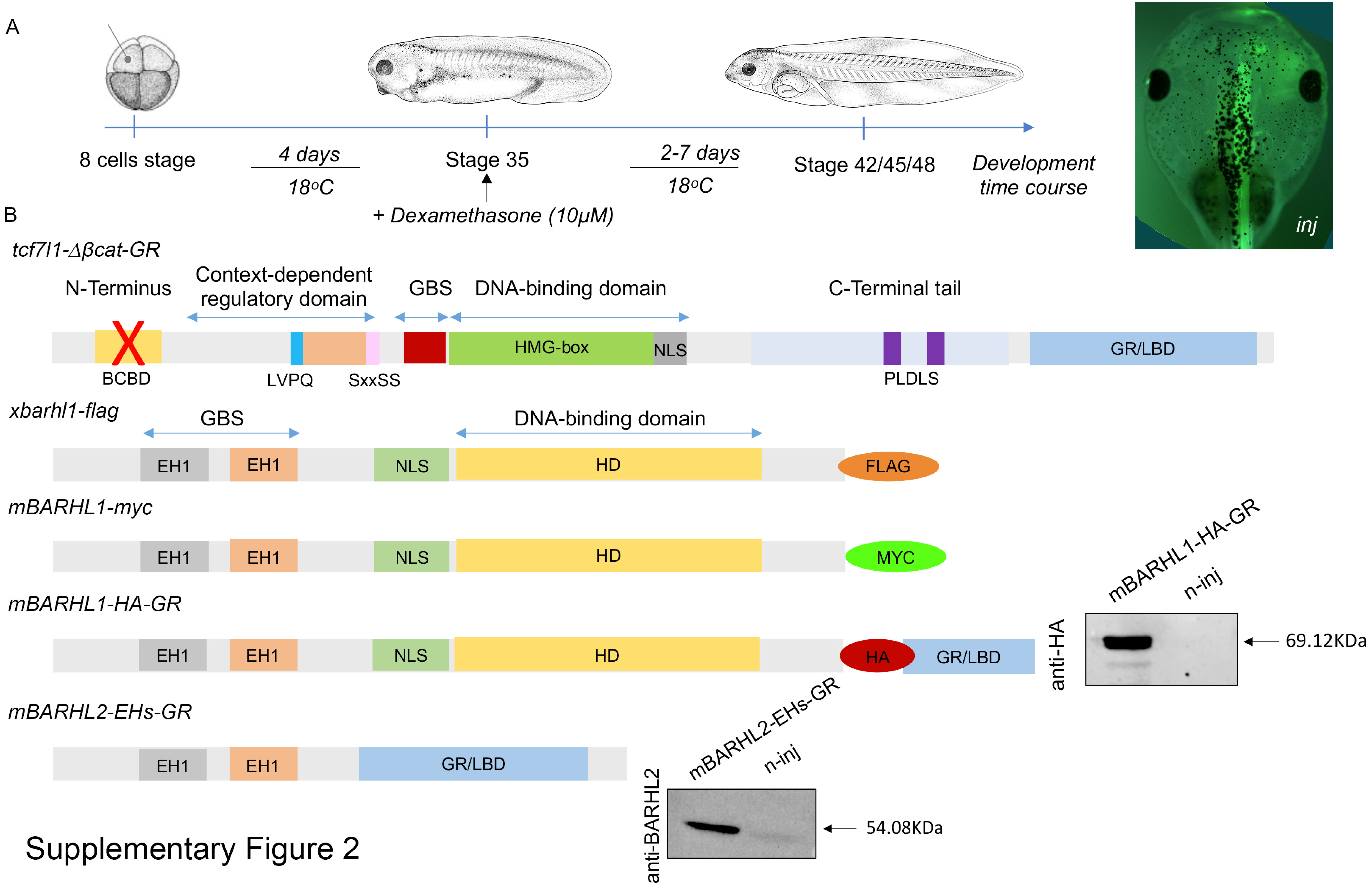


Supplementary Figure 2

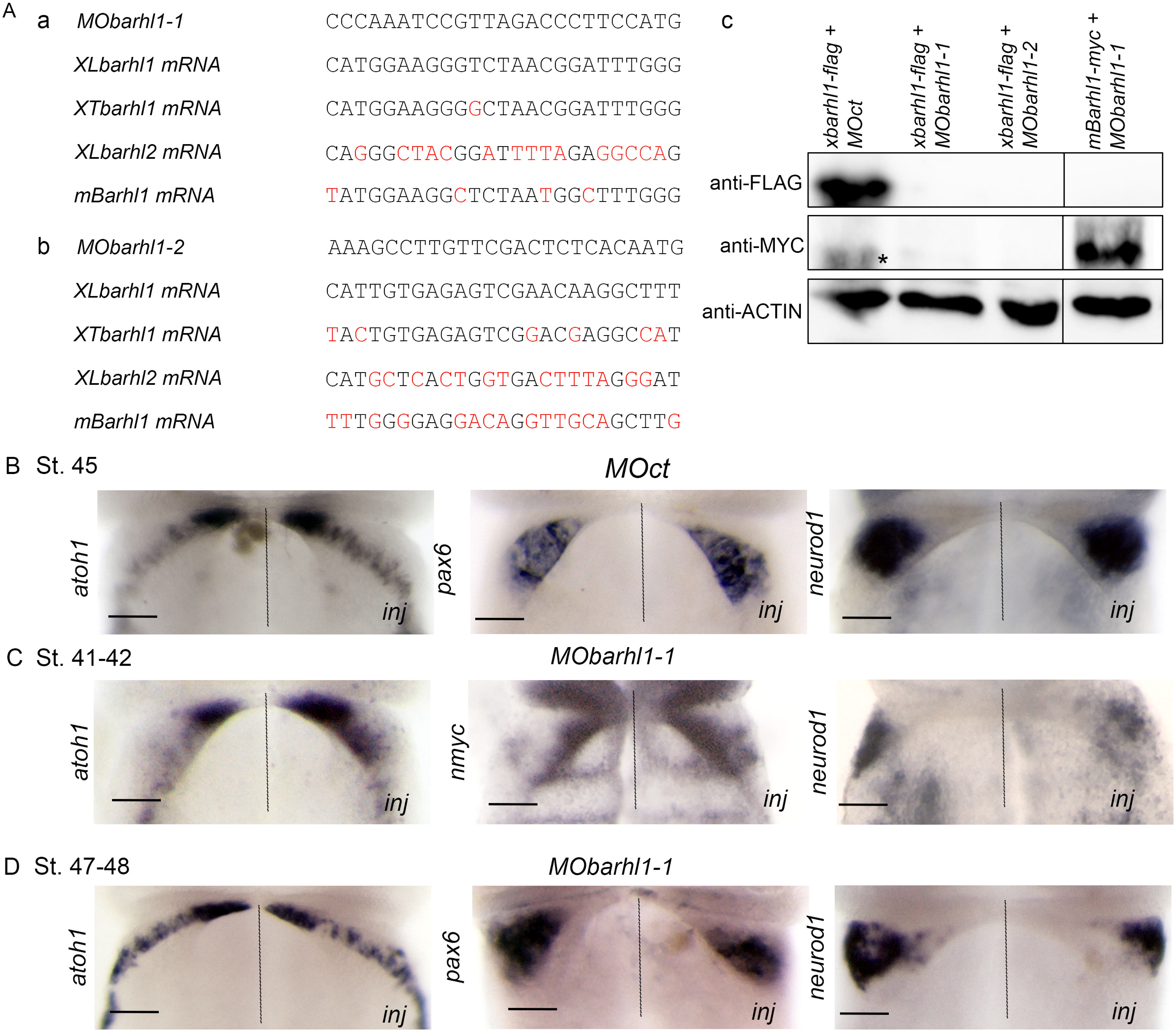


Supplementary Figure 3

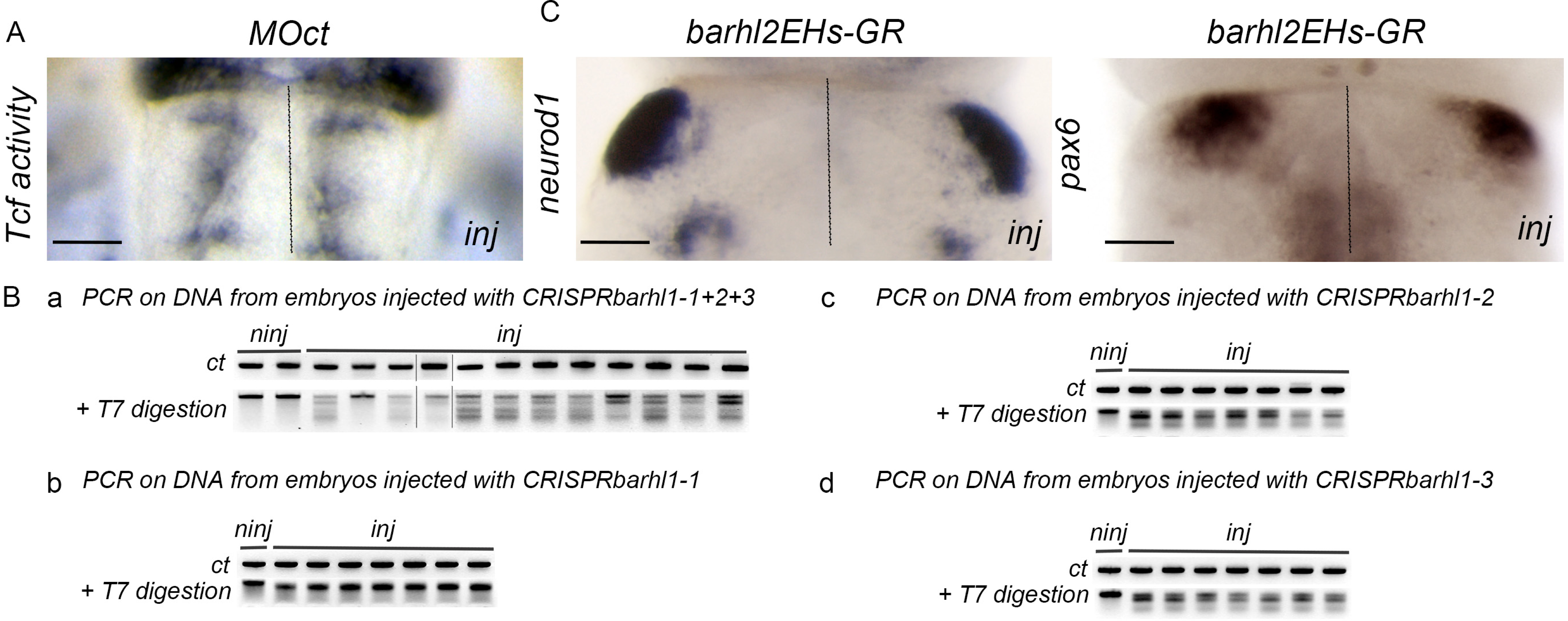


Supplementary Figure 4

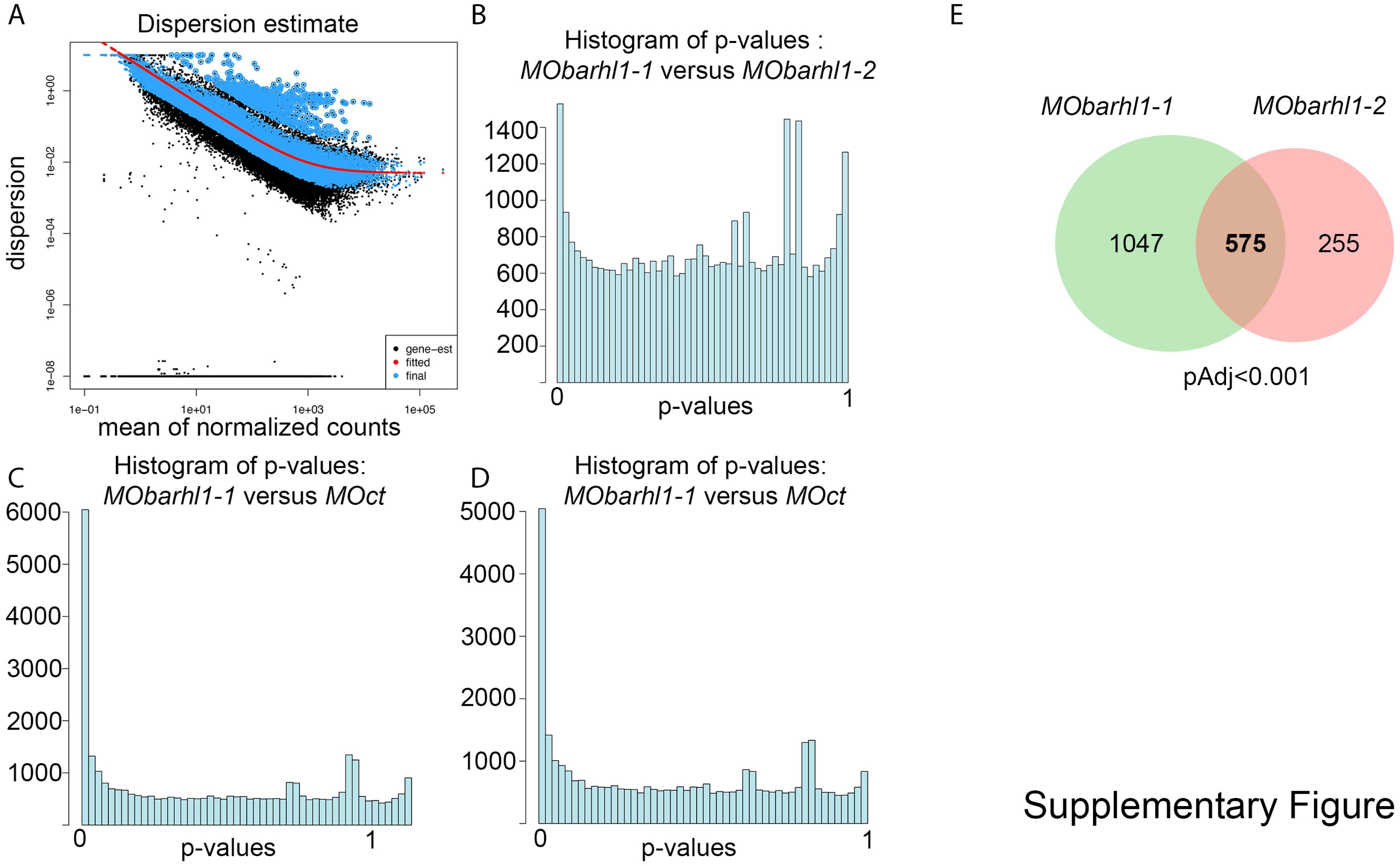


Supplementary Figure 6

**KDBarhl1_A_vs_KDCT pAdj<0.001**

| Gene | meanCounts | Log2FC | SD | Wald | pVal | pAdj |
|---|---|---|---|---|---|---|
| otp.L | 4498.84634473394 | -1.15359174551513 | 0.0788940099340072 | -14.6220447722214 | 2.03204451526619e-48 | 4.93664894538767e-44 |
| otx2.S | 474.089930221584 | 1.87165627384235 | 0.149833551293335 | 12.4915698632687 | 8.29995501355197e-36 | 1.00819553549616e-31 |
| otp.S | 3420.42311010403 | -1.02103163193367 | 0.0827825257365918 | -12.3339028719964 | 5.94937527630556e-35 | 4.81780409875225e-31 |
| slc6a3.S | 98.2928363391053 | 2.18360840231818 | 0.194222747333913 | 11.2428046266077 | 2.51270386555132e-29 | 1.52609069274259e-25 |
| helt.L | 302.019020885779 | 1.98998424091082 | 0.178297298086386 | 11.161045412739 | 6.32434676307051e-29 | 3.0728736052407e-25 |
| dmrta2.S | 101.476947605541 | 2.13854594233106 | 0.192489498202655 | 11.109935670774 | 1.12240343390244e-28 | 4.544611503871e-25 |
| ldha.S | 5459.2709775048 | -0.615728469541824 | 0.0562190771568151 | -10.9523048168211 | 6.47780247797267e-28 | 2.24816761999811e-24 |
| dner.S | 5574.26893781311 | -0.727189341389905 | 0.066619632333731 | -10.9155411988324 | 9.71489440083035e-28 | 2.95017055717216e-24 |
| otx2.L | 1301.45818728385 | 1.15878492959565 | 0.109861019373537 | 10.5477350947899 | 5.20363420538696e-26 | 1.40463432650745e-22 |
| nr2e1.L | 134.855866685233 | 2.01590940459308 | 0.192873937532354 | 10.4519533866773 | 1.43537911256732e-25 | 3.48711001607106e-22 |
| LOC121393901 | 1651.28239302207 | 1.29494458168199 | 0.127378802325441 | 10.1660916733502 | 2.80956810055679e-24 | 5.89223427548713e-21 |
| nefl.L | 4095.08931530961 | -0.930046216438326 | 0.0915160830662501 | -10.1626532220031 | 2.91046395430335e-24 | 5.89223427548713e-21 |
| LOC108704633 | 2141.34855501679 | -0.694201878443451 | 0.0690396906942687 | -10.0551128121013 | 8.72214794050146e-24 | 1.62996816974264e-20 |
| rgr.L | 282.323846705935 | 1.57234999748159 | 0.162528006148918 | 9.6743326564998 | 3.87612162295944e-22 | 6.72617847915547e-19 |
| lef1.L | 520.784379224114 | 1.41056555477071 | 0.147787677992212 | 9.54454101948228 | 1.36711616602004e-21 | 1.95368941984064e-18 |
| skor1.L | 685.480279731138 | -0.99383415535365 | 0.10409715146248 | -9.54717916284059 | 1.3327567984313e-21 | 1.95368941984064e-18 |
| dmrta2.L | 113.472457035016 | 1.85761717264088 | 0.194400418324687 | 9.55562332966945 | 1.22842182106314e-21 | 1.95368941984064e-18 |
| slc39a12.S | 495.19600779649 | 1.34333346764006 | 0.142154355425106 | 9.4498227903245 | 3.39404816559008e-21 | 4.58083367415808e-18 |
| tal2.S | 767.741658266835 | 1.44064369259442 | 0.153310458826451 | 9.39690418789379 | 5.61920605815065e-21 | 7.18489431456378e-18 |
| nr2e1.S | 118.89335064376 | 1.73504856631917 | 0.185772711422724 | 9.33963095565246 | 9.66704728221066e-21 | 1.17425623337012e-17 |
| pou4f4.L | 337.228606988696 | 1.52057717474746 | 0.165256346307002 | 9.20132393537635 | 3.53566252072251e-20 | 4.09025644183013e-17 |
| lef1.S | 429.733279673732 | 1.33199732408724 | 0.145147005193151 | 9.17688465094211 | 4.43743217297532e-20 | 4.9001353277392e-17 |
| slc4a4.L | 1401.50693400372 | -0.793004756678376 | 0.0869578115015006 | -9.11941944013497 | 7.5527831907578e-20 | 7.97770934070739e-17 |
| slit1.S | 1813.49900261657 | -0.710872890736128 | 0.0779923930304521 | -9.11464391736983 | 7.89292365800899e-20 | 7.9896119728196e-17 |
| mapt.L | 17602.41813225 | -0.5922823927577 | 0.0652038072415668 | -9.08355535994109 | 1.05084869930477e-19 | 1.0211727320364e-16 |
| crmp1.L | 5653.92500434536 | -0.715869531023678 | 0.0788853538610895 | -9.07480915005166 | 1.13876895261353e-19 | 1.06404818979973e-16 |
| nkx2-4.S | 58.9493268350775 | 1.75024180971758 | 0.193819340978297 | 9.03027427956002 | 1.71242048688443e-19 | 1.54079790031002e-16 |
| LOC108699226 | 8729.96741449577 | -0.785982622824739 | 0.0872088783900174 | -9.01264455334068 | 2.01146357769942e-19 | 1.74523200559392e-16 |
| sp5.S | 201.620840624883 | 1.55186377952788 | 0.174936436084213 | 8.87101517708313 | 7.24820738755466e-19 | 6.07199828528458e-16 |
| slc6a4.L | 1113.10980162782 | -0.987150375798448 | 0.111350597632511 | -8.86524542110074 | 7.63357410677509e-19 | 6.18166831166647e-16 |
| pou4f4.S | 203.970652047039 | 1.5329329397058 | 0.173112784480412 | 8.85511110174104 | 8.36004997082029e-19 | 6.55158238680994e-16 |
| dner.L | 6391.66455116577 | -0.675687809917366 | 0.0778881261985737 | -8.67510675753988 | 4.13162699455839e-18 | 3.13667956893129e-15 |
| atp1a1.L | 9448.70733264771 | -0.648667823560026 | 0.0748367674570057 | -8.66776914078619 | 4.40669072077842e-18 | 3.2441255869876e-15 |
| rbp1.S | 455.860425898749 | 1.1608482322363 | 0.134386436885938 | 8.63813535901381 | 5.71378970784224e-18 | 4.08267079889174e-15 |
| cox4i2.L | 1319.30308557081 | 0.689332546438432 | 0.0798543715727348 | 8.63237081279336 | 6.0093198065162e-18 | 4.1711547251287e-15 |
| igfbpl1.L | 3039.22136102478 | 0.793767783549317 | 0.093947614293959 | 8.44904673221018 | 2.93690905215525e-17 | 1.98192412536277e-14 |
| crx.L | 73.966489578511 | 1.59895783344645 | 0.189732866907344 | 8.42741618523745 | 3.53381221693911e-17 | 2.32028199995456e-14 |
| efna2.L | 986.166119930827 | 0.849044156491525 | 0.100933252536999 | 8.41193694991941 | 4.03294671984356e-17 | 2.57832651610209e-14 |
| lhx2.L | 711.620417944345 | 0.921913704820772 | 0.109699974354287 | 8.40395551819695 | 4.31686262349003e-17 | 2.68907334807863e-14 |
| app.L | 8825.46075688036 | -0.626822204401027 | 0.0748726122207 01 | -8.37184900873168 | 5.67210943684603e-17 | 3.44495566646844e-14 |
| nkx2-2.L | 471.457869917758 | 1.33056312351714 | 0.159322106814763 | 8.35140301693434 | 6.74568220442386e-17 | 3.99706349937252e-14 |
| LOC108719610 | 72.0829012716346 | 1.60017435789276 | 0.192480690536587 | 8.31342797779813 | 9.29764577546614e-17 | 5.37802396355177e-14 |
| igfbpl1.S | 1122.38106871084 | 0.907853131567451 | 0.109531758953337 | 8.28849221671149 | 1.14693066970386e-16 | 6.47989155576407e-14 |
| LOC108710372 | 3582.80539203409 | -0.66290226649708 | 0.0800750830480818 | -8.2785086354395 | 1.24727455554583e-16 | 6.88665637555238e-14 |
| rora.L | 628.945763780762 | -0.800977432205159 | 0.0971117986910465 | -8.24799296276456 | 1.61076852599393e-16 | 8.69600234899922e-14 |
| app.S | 10664.4445379749 | -0.614023130150492 | 0.0747131095371992 | -8.21841218969442 | 2.06215146085921e-16 | 1.08908494761117e-13 |
| hspa12a.S | 483.598822721795 | -0.96692198200007 | 0.118958669591517 | -8.12821785348063 | 4.35647749994008e-16 | 2.25183541241584e-13 |
| sulf2.L | 3447.00702636397 | -0.550213171524578 | 0.0682902314330663 | -8.05698208921524 | 7.82011032182271e-16 | 3.95795333663252e-13 |
| mapt.S | 10225.4048883714 | -0.501541211327098 | 0.0627338769565395 | -7.9947428033908 | 1.29844633581169e-15 | 6.43764393514475e-13 |
| pacsin2.L | 4377.66197931134 | -0.586558275490463 | 0.0738032486989932 | -7.94759425676155 | 1.9016830628177e-15 | 9.23989766561865e-13 |
| plxna1.L | 6553.39790156272 | -0.606978488563316 | 0.076477917433568 | -7.93665032903862 | 2.07714573226537e-15 | 9.89454478816763e-13 |
| LOC108719750 | 1363.45527026836 | -0.74031181149366 | 0.0936596447841542 | -7.904277377944 | 2.69492336787878e-15 | 1.25904746729321e-12 |
| LOC108697760 | 11408.9440404374 | -0.533305739089508 | 0.0675224680710175 | -7.89819676805353 | 2.82967199712679e-15 | 1.2970575754377e-12 |
| zic3.L | 687.438450672142 | 0.99152431295769 | 0.125732134498838 | 7.886005569762 | 3.12012690496804e-15 | 1.4037104264684e-12 |
| nefl.S | 1947.66469121088 | -0.857192734273325 | 0.108893039101004 | -7.87187814161595 | 3.49356367252148e-15 | 1.54313883382249e-12 |
| slc45a4.L | 877.360392491325 | -0.739928904522114 | 0.0940525228217264 | -7.86718827228724 | 3.62700596968115e-15 | 1.57347291120418e-12 |
| rgma.L | 5038.05291087251 | -0.619646366343524 | 0.0789952575307694 | -7.84409578134696 | 4.36084174992162e-15 | 1.85863665741396e-12 |
| nkx6-1.L | 626.861691066251 | -0.908490476776176 | 0.117131674022422 | -7.75614695477047 | 8.7548570607308e-15 | 3.66707754195507e-12 |
| cachd1.S | 944.022500788257 | 0.783802705176573 | 0.10111181535866 | 7.75184089412593 | 9.05698280821874e-15 | 3.72932780242146e-12 |
| opnpl.S | 64.0845960759803 | 1.47359226567724 | 0.190809589341117 | 7.72284176474408 | 1.13764369273322e-14 | 4.60631931187679e-12 |
| LOC108695757 | 4776.33091855359 | -0.709252987666007 | 0.0920491930226705 | -7.70515160835062 | 1.30688043896478e-14 | 5.20481203019842e-12 |
| nefh.L | 2220.10104305609 | -0.991169266772848 | 0.128724549134588 | -7.69992416704081 | 1.36147003580523e-14 | 5.33476662094393e-12 |
| LOC108703510 | 809.590332718214 | -0.863999163893274 | 0.112546606110081 | -7.67681224477089 | 1.63096203567589e-14 | 6.28930026900158e-12 |
| not.S | 27.9850032000555 | 1.43455559203037 | 0.186938618797404 | 7.67393918527385 | 1.66793182862588e-14 | 6.33136497572455e-12 |
| crx.S | 68.5326858897321 | 1.43249057843058 | 0.18766356145871 | 7.63329101982196 | 2.28835516580323e-14 | 8.55281544584979e-12 |
| npy.L | 845.358066951188 | 0.95873303959195 | 0.125984608619698 | 7.60992195868954 | 2.74261477104276e-14 | 1.00953156435929e-11 |
| nnt.S | 2043.61296470859 | -0.574427224466309 | 0.0757796396255869 | -7.58023167310437 | 3.44938456922107e-14 | 1.24833918407389e-11 |
| ntn1.L | 5319.01266882461 | -0.673993630048701 | 0.0889342800078686 | -7.57855834655735 | 3.4941575910523e-14 | 1.24833918407389e-11 |
| ctnnb1.L | 9699.86497436317 | 0.490809988683321 | 0.0649970312624481 | 7.55126779100887 | 4.31041517962192e-14 | 1.51764096193819e-11 |
| ctnna2.L | 1027.84874334319 | -0.665291706559749 | 0.0882429193195689 | -7.53932113408914 | 4.72424290167871e-14 | 1.63958224361975e-11 |
| otx1.L | 550.606158142413 | 1.03089025865552 | 0.137385927659824 | 7.50360882089805 | 6.20843964443974e-14 | 2.12433567214111e-11 |
| paqr6.S | 591.103323780211 | -0.789936964958847 | 0.105622956490143 | -7.47883785124465 | 7.49827203472145e-14 | 2.5300419557156e-11 |
| LOC121394610 | 7550.90533736695 | 0.476736575001744 | 0.0637900760623 38 | 7.47352259834053 | 7.80760237976015e-14 | 2.59832729060128e-11 |
| hmgb2.L | 24995.9234695256 | 0.452603362485464 | 0.0606216976596498 | 7.46602916049182 | 8.26510380153973e-14 | 2.69739763514529e-11 |
| bsx.L | 31.6339464250892 | 1.39623161404494 | 0.187036023409364 | 7.46504116476546 | 8.32735748069058e-14 | 2.69739763514529e-11 |
| efna2.S | 621.250620245399 | 0.909984335965475 | 0.121961011220734 | 7.46127247435263 | 8.56908352417605e-14 | 2.73917519916228e-11 |
| ctnnb1.S | 15175.6387960335 | 0.470004897218403 | 0.0632689910946533 | 7.42867697250387 | 1.09689253893842e-13 | 3.46076718713894e-11 |
| lrp1.L | 5371.81583492197 | -0.577311549580665 | 0.0777838846033437 | -7.42199431829158 | 1.15369763588544e-13 | 3.59332441517035e-11 |
| rtn4.S | 6515.00857492491 | -0.489942400756993 | 0.0661313351330626 | -7.40862708685953 | 1.2761358675733e-13 | 3.92436009706654e-11 |
| fam219a.S | 2335.21429307898 | -0.658615994689745 | 0.0891076774495595 | -7.39123736069277 | 1.45468668403138e-13 | 4.4175197877323e-11 |
| LOC108696074 | 1871.07661177262 | -0.597553251025306 | 0.0809351794430561 | -7.38310899088979 | 1.54635336358811e-13 | 4.63791464382834e-11 |
| hmgn1.S | 27311.2821214967 | 0.488978325489644 | 0.0662734544423254 | 7.37819281647946 | 1.60452486066436e-13 | 4.69642493553976e-11 |
| LOC108710030 | 2950.41175258469 | -0.593652133913865 | 0.0804537129169222 | -7.37880344350148 | 1.59718419583583e-13 | 4.69642493553976e-11 |
| gpc1.S | 2289.51157411177 | -0.572333394559083 | 0.0776410611761733 | -7.37152977933179 | 1.68681073373608e-13 | 4.87849761492671e-11 |
| ank1.S | 2153.08359288837 | -0.624114234813624 | 0.0849736706245821 | -7.34479551402448 | 2.06073908543267e-13 | 5.88983474605896e-11 |
| pitx3.L | 42.5464940543809 | 1.42033604832274 | 0.193558701482708 | 7.3380118663879 | 2.1678995267823e-13 | 6.12406408181968e-11 |
| LOC108706170 | 950.502770323648 | -0.860732968800999 | 0.117744921786597 | -7.31014939535236 | 2.66845534574186e-13 | 7.4514315137302e-11 |
| fry.L | 3418.13245587727 | -0.524734762539569 | 0.0718635983951769 | -7.30181586029216 | 2.83909654027643e-13 | 7.74977655612084e-11 |
| glipr2.S | 1238.87213421075 | -0.591016916771125 | 0.0809308607473008 | -7.30273855132373 | 2.8196876800812e-13 | 7.74977655612084e-11 |
| cyp27c1.S | 1106.68809416432 | 0.629519021571361 | 0.0864785895677768 | 7.2794783624215 | 3.35113411384454e-13 | 9.04582801797103e-11 |
| col12a1.L | 513.162495920935 | -0.819757385807293 | 0.112731724860223 | -7.27175412976001 | 3.54849038279811e-13 | 9.47329949007662e-11 |
| atp6v1a.L | 2639.43088732933 | -0.588959019896682 | 0.0811235746041694 | -7.26002302993206 | 3.87024423437965e-13 | 1.0219968851089e-10 |
| prdm12.L | 454.263674587309 | 0.988627311230685 | 0.136364493321753 | 7.24988805478866 | 4.17116235467499e-13 | 1.08961524994058e-10 |
| caprin1.L | 2573.98567862467 | -0.539440568951877 | 0.0744882772418739 | -7.24195254509965 | 4.42270614160061e-13 | 1.13967138602415e-10 |
| rplp1.L | 16579.8346101088 | 0.424170338976629 | 0.0585796425860243 | 7.24091715571215 | 4.4560605815105e-13 | 1.13967138602415e-10 |
| znf703.L | 981.926377415962 | -0.79047091320857 | 0.109337290574896 | -7.22965521691888 | 4.84221789725149e-13 | 1.22538376662321e-10 |
| not.L | 32.8874683343444 | 1.36364179544179 | 0.188754255325306 | 7.22442942063289 | 5.03210125853697e-13 | 1.26030791726698e-10 |
| kcnh4.L | 921.997077259224 | -0.862005766333303 | 0.119728160945695 | -7.19969102945612 | 6.03491522695612e-13 | 1.49604316860 89e-10 |
| plxnb1.S | 3341.99150138984 | -0.555805935790047 | 0.0773251243436547 | -7.18790872317112 | 6.57912185281814e-13 | 1.6144766292158e-10 |
| glra1.S | 617.346013216389 | -0.940108841603404 | 0.130894987780873 | -7.18216073465863 | 6.86181795847467e-13 | 1.66701005483184e-10 |
| rplp2.L | 8752.60318216921 | 0.426430944862992 | 0.0594123280261716 | 7.17748252627 4 | 7.10067421367749e-13 | 1.70795821135724e-10 |
| endod1.L | 9817.78353497793 | -0.454150197574216 | 0.0632874337815001 | -7.17599324918386 | 7.17840966169975e-13 | 1.70972827766013e-10 |
| dchs1.S | 4830.52985893699 | -0.473837563285357 | 0.0662000744665851 | -7.15765906765753 | 8.20663405011732e-13 | 1.91703815013029e-10 |
| map4.L | 5194.52685838413 | -0.511705580694262 | 0.0714783987294551 | -7.15888421942777 | 8.13363428179792e-13 | 1.91703815013029e-10 |
| rpe65.L | 454.146017051386 | 1.21497656449879 | 0.169973760144935 | 7.14802428011705 | 8.80357435576425e-13 | 2.03689557522797e-10 |

| | | | | | | |
|---|---|---|---|---|---|---|
| skor1.S | 565.759626398 | -0.820606641216342 | 0.114870056069925 | -7.14378202024045 | 9.07973496067318e-13 | 2.08097246353391e-10 |
| nfib.L | 1594.64941260286 | -0.564582119293252 | 0.0793421162966898 | -7.1157935488142 | 1.11270748077058e-12 | 2.52636593811593e-10 |
| hmgb2.S | 12962.511026317 | 0.455457687664851 | 0.0640608049429906 | 7.10977153768478 | 1.1623520111433e-12 | 2.61464627395512e-10 |
| kalrn.L | 3091.24482714556 | -0.438088156027231 | 0.061750197713704 | -7.09452232134324 | 1.29799025310798e-12 | 2.89297020266103e-10 |
| sulf2.S | 4283.87851341345 | -0.508486131439755 | 0.071775137808605 | -7.08443267354888 | 1.39615066896348e-12 | 3.08346221379988e-10 |
| gucy2d.S | 118.170657897624 | 1.37653042480735 | 0.194720856575446 | 7.06925004858943 | 1.55773238254504e-12 | 3.40932887401343e-10 |
| znf703.S | 1105.5449356608 | -0.792185761353631 | 0.112435610906571 | -7.04568379151602 | 1.84552931831336e-12 | 4.00315082670579e-10 |
| LOC108707869 | 1411.0532119591 | -0.711479848720536 | 0.101175802000404 | -7.03211474140523 | 2.03426399285046e-12 | 4.37348756126628e-10 |
| rgr.S | 611.098053386951 | 1.16328518524841 | 0.165513197301689 | 7.02835305107443 | 2.08985560939714e-12 | 4.45359229602579e-10 |
| barhl2.L | 436.621948637395 | 1.0554058593549 | 0.150637803346508 | 7.00624833812248 | 2.44793260774322e-12 | 5.1267305838374e-10 |
| meis1.S | 1018.06231552231 | -0.774843185827328 | 0.11057414965982 | -7.00745326291112 | 2.42695048554284e-12 | 5.1267305838374e-10 |
| scn1b.L | 2277.76168858732 | -0.639834203037718 | 0.0913769161378147 | -7.00214266448563 | 2.52077193640132e-12 | 5.23415670281485e-10 |
| LOC108716604 | 28.2600460154639 | 1.29452124574962 | 0.185559540381847 | 6.97631198636156 | 3.03029401207867e-12 | 6.18638342264194e-10 |
| nppc.L | 628.385891872387 | -0.742135649857594 | 0.106378671535401 | -6.97635756440732 | 3.02931157876605e-12 | 6.18638342264194e-10 |
| pitx3.S | 36.403007255475 | 1.34762752981699 | 0.193284397175458 | 6.9722520260839 | 3.1190713062774e-12 | 6.31455985595586e-10 |
| zic3.S | 427.116360823445 | 0.975237249860131 | 0.14019418696186 | 6.95633157832318 | 3.49247327905034e-12 | 7.01207816869826e-10 |
| hmgn2.S | 13177.0998214996 | 0.445613286046956 | 0.0641514492447257 | 6.94626997976345 | 3.7507067635041e-12 | 7.46882541906301e-10 |
| wdr7.L | 2045.18309940234 | -0.663711659729695 | 0.0956601216662567 | -6.93822721703493 | 3.97050759120238e-12 | 7.84223670086753e-10 |
| rps7.L | 12102.6664938259 | 0.425368287322251 | 0.0614758300838843 | 6.9192768400497 | 4.53954975005216e-12 | 8.89385658288446e-10 |
| rpl7.L | 20041.6111835862 | 0.417885077831273 | 0.0604802342282749 | 6.90944873417685 | 4.86540654128016e-12 | 9.45601492110881e-10 |
| paqr6.L | 3033.15967644058 | -0.562687392973677 | 0.0814632001174321 | -6.90725864147913 | 4.94108606520766e-12 | 9.52688451334563e-10 |
| cacna1a.S | 3512.87605570904 | -0.638903835457279 | 0.0925366976323227 | -6.9043293288447 | 5.04411593803102e-12 | 9.6489568975217e-10 |
| ppp3cb.L | 1701.06968906993 | -0.586467438212976 | 0.0850068302096365 | -6.89906254317072 | 5.2346823040736e-12 | 9.93526342930969e-10 |
| ap2a2.L | 3110.9747100826 | -0.417092420491619 | 0.0605298357284386 | -6.89069143294664 | 5.55218548086691e-12 | 1.04561855869908e-09 |
| LOC108708507 | 2351.58727490952 | -0.629979644522835 | 0.0914582629189768 | -6.88816542558802 | 5.65164691898524e-12 | 1.05616238653713e-09 |
| gngt1.S | 89.7941460829131 | 1.239836966126 | 0.180125319027437 | 6.88319095183471 | 5.85264897648909e-12 | 1.08537598652539e-09 |
| nmu.S | 60.1247696223422 | 1.33203102295551 | 0.194674935741628 | 6.84233446838463 | 7.79129122832508e-12 | 1.43395173561331e-09 |
| rpl34.S | 6324.80400414156 | 0.413719183956911 | 0.0604783467754886 | 6.84078196602736 | 7.87620556946293e-12 | 1.43868073762806e-09 |
| stxbp1.S | 14500.7663055513 | -0.562540976762014 | 0.0822985210838701 | -6.8353716367361 | 8.17927088962721e-12 | 1.48288960442241e-09 |
| bsx.S | 28.9822538589958 | 1.23128120370261 | 0.181156605909826 | 6.79677783494963 | 1.06984881045998e-11 | 1.92525237046777e-09 |
| LOC108700035 | 1601.13324350975 | -0.650687930218076 | 0.0959360992442373 | -6.78251393734004 | 1.18102464431396e-11 | 2.10969211095319e-09 |
| nme2.S | 3859.51394696091 | 0.411454610539084 | 0.0607382954511057 | 6.77422057176932 | 1.25078724241911e-11 | 2.21800184433064e-09 |
| cntnap1.L | 1170.56058459662 | -0.722822273963238 | 0.106724201828004 | -6.77280562030468 | 1.26308606844827e-11 | 2.22358064832481e-09 |
| nr2f1.L | 3223.09980468629 | -0.490650247918651 | 0.0725203840653617 | -6.76568738903029 | 1.32677746275867e-11 | 2.31890155973088e-09 |
| lrp1.S | 4344.29096307483 | -0.526961527038136 | 0.0779885313474063 | -6.75691050894064 | 1.40965196582166e-11 | 2.446148918405e-09 |
| lhx2.S | 899.233305608489 | 0.664983436259647 | 0.0984873678042876 | 6.75196678604601 | 1.45854270811796e-11 | 2.51303805326367e-09 |
| agpat3.S | 2930.80078956433 | -0.507320490988701 | 0.0753022943429522 | -6.73711864180647 | 1.6155826207566e-11 | 2.71569074074134e-09 |
| nefm.S | 15404.2947008401 | -0.728722443449827 | 0.108142954813282 | -6.73851056416968 | 1.60018425738668e-11 | 2.71569074074134e-09 |
| th.S | 77.9524399838011 | 1.29727937482154 | 0.192566867344879 | 6.7367735307142 | 1.61942286011964e-11 | 2.71569074074134e-09 |
| tmem63b.S | 2269.386406138 | -0.478700790032323 | 0.0710592452859325 | -6.736643319333 | 1.62087411462704e-11 | 2.71569074074134e-09 |
| scrt2.S | 353.094437983648 | -0.814617697887976 | 0.120992673512667 | -6.73278533516072 | 1.66445542754296e-11 | 2.76960822991293e-09 |
| barhl2.S | 414.272266541186 | 1.07211235799577 | 0.159372070888055 | 6.72710313684028 | 1.73074069964703e-11 | 2.84098747008277e-09 |
| sez6l2.L | 6445.04983857967 | -0.51209824919828 | 0.0761203459687131 | -6.72748189306394 | 1.72624307161518e-11 | 2.84098747008277e-09 |
| slc2a1.S | 1176.58317459525 | -0.692642027446262 | 0.103084875487199 | -6.71914307673849 | 1.82796354872871e-11 | 2.98043935924934e-09 |
| atp6ap1.2.S | 3818.88370978691 | -0.446110266712066 | 0.0664929220737083 | -6.70913915044291 | 1.95775911832188e-11 | 3.17078666803411e-09 |
| fn1.S | 3333.66041002239 | -0.601426646132078 | 0.0898640763204277 | -6.69262591636256 | 2.1920098830916e-11 | 3.52666808608128e-09 |
| purb.L | 6378.43477533977 | -0.455904413926598 | 0.0683487429712313 | -6.67026771975152 | 2.55337204869537e-11 | 4.05435428437944e-09 |
| bag6.L | 3682.45540137023 | -0.439863059347232 | 0.0659374490298161 | -6.67091411359168 | 2.54214969631737e-11 | 4.05435428437944e-09 |
| mab21l1.S | 2099.56840030275 | -0.5895027662205 | 0.088513456157289 | -6.66003556761979 | 2.73761325640449e-11 | 4.31867379552536e-09 |
| prrt2.L | 1292.06997769031 | -0.745787952107593 | 0.112401133039704 | -6.63505724487808 | 3.24376621649267e-11 | 5.08413267506276e-09 |
| dnm1l.L | 3910.27916995742 | -0.495438804397123 | 0.075004095286222 | -6.60549004033028 | 3.96204545265408e-11 | 6.17012386069091e-09 |
| LOC121393045 | 19586.6559782129 | 0.410657547967602 | 0.0622417595631265 | 6.59778179232076 | 4.17354918117993e-11 | 6.45810215334938e-09 |
| hmgn1.L | 20689.0148154507 | 0.451302752095097 | 0.0684275618062709 | 6.59533585856537 | 4.2429423238172e-11 | 6.52392663384906e-09 |
| emx2.L | 148.673957134924 | 1.27952615280486 | 0.194295074459951 | 6.58547910368464 | 4.53420864795076e-11 | 6.9279286096425e-09 |
| wnt2b.L | 47.8190854361246 | 1.28173726375021 | 0.19467559400965 | 6.58396482759247 | 4.58065530680729e-11 | 6.95515250147351e-09 |
| nkx1-1.L | 382.58139911905 | -0.89775224280972 | 0.136571999088537 | -6.57347222564797 | 4.91552460987073e-11 | 7.39098200408071e-09 |
| rspo2.L | 128.253449009422 | 1.15444296565613 | 0.175632000477358 | 6.57307872436925 | 4.92853825908074e-11 | 7.39098200408071e-09 |
| gpc1.L | 1038.56511111647 | -0.631175634996507 | 0.0964895030874131 | -6.54139170376599 | 6.09489619373037e-11 | 9.08401276874144e-09 |
| ogdh.S | 1609.81511893786 | -0.508370227956707 | 0.0777799831220685 | -6.53600332053128 | 6.31845131008023e-11 | 9.35978391018836e-09 |
| LOC108718165 | 6252.33958250651 | -0.371936414980999 | 0.0569326164486628 | -6.53292327283748 | 6.44981910013661e-11 | 9.49647910416478e-09 |
| maf.L | 1047.08754968366 | -0.686589775086649 | 0.105356934040541 | -6.51679722212161 | 7.18244154640557e-11 | 1.05114599354444e-08 |
| dach1.S | 947.271067591902 | -0.623581290354189 | 0.0959862798138123 | -6.49656692147846 | 8.21735717419453e-11 | 1.19540404305319e-08 |
| LOC121393057 | 1092.48614652167 | -0.845423313438912 | 0.130362636442824 | -6.48516581520434 | 8.86342514409891e-11 | 1.2817145860163e-08 |
| actl6a.S | 2217.5257722163 | 0.433941501271432 | 0.0669924138307102 | 6.4774722458576 | 9.32719222565916e-11 | 1.34079768006014e-08 |
| sptbn1.L | 12353.6935492804 | -0.526184372058495 | 0.0815282687122528 | -6.45401135544309 | 1.08927643111934e-10 | 1.55664009515371e-08 |
| LOC108718312 | 2782.5664321436 | -0.51893068527954 | 0.0804336583713282 | -6.4516608567505 | 1.10630937836433e-10 | 1.57173567473585e-08 |
| rps7.S | 12297.6491481465 | 0.388940902716077 | 0.0603811425116822 | 6.44142999846064 | 1.18353034084431e-10 | 1.67166779653905e-08 |
| LOC108711739 | 1399.3575195954 | -0.721606378710993 | 0.112270344937146 | -6.42739967633468 | 1.29805207624232e-10 | 1.82282526822145e-08 |
| LOC108695983 | 4349.54964352368 | -0.523577411574269 | 0.0815306810757025 | -6.42184518350975 | 1.34632311584059e-10 | 1.87974561932363e-08 |
| dnajc5.S | 7872.99361108948 | -0.522404308905423 | 0.0816850678930229 | -6.39534644923818 | 1.60183742597106e-10 | 2.22371648151663e-08 |
| sh3glb2.S | 3342.45816836182 | -0.413287563523191 | 0.0646844638649236 | -6.38928637309622 | 1.66661687359857e-10 | 2.30049945040493e-08 |
| kcnf1.L | 309.731270393435 | 1.00887775504585 | 0.157953804131035 | 6.38716972089451 | 1.68984058753146e-10 | 2.31937780980166e-08 |
| vav2.L | 1074.18397487009 | -0.523646421033831 | 0.0820197067591842 | -6.38439762496706 | 1.72073450879323e-10 | 2.3485125930687e-08 |
| nkx2-1.L | 165.870906973826 | 1.13153105890676 | 0.177363844341901 | 6.37971658262848 | 1.77416011886301e-10 | 2.40790200713174e-08 |
| rack1.L | 28359.7478307492 | 0.4001991738962 | 0.0627432249122721 | 6.37836474704258 | 1.78988817385708e-10 | 2.41575240531577e-08 |
| LOC108709778 | 1657.7219135364 | -0.618905637270611 | 0.097104656390301 | -6.37359381390518 | 1.84649237478844e-10 | 2.47838042834864e-08 |
| atp6v1a.S | 6031.77901410661 | -0.529585832249219 | 0.0832658573900684 | -6.36017989664496 | 2.01517595134485e-10 | 2.67522866458862e-08 |
| hmgn3.S | 5069.44803576131 | 0.394050380991664 | 0.0619547409119139 | 6.36029422755423 | 2.01367650860348e-10 | 2.67522866458862e-08 |
| LOC108707564 | 5597.80400520361 | -0.550745316307646 | 0.0866557212853372 | -6.35555631109652 | 2.07673653411074e-10 | 2.74196942172208e-08 |
| gipr.L | 126.162125019071 | 1.15947086338288 | 0.182793712273661 | 6.34305660167913 | 2.25250624152973e-10 | 2.95796684495801e-08 |
| ebf2.L | 1848.58361348298 | -0.455687788587676 | 0.0718691827575512 | -6.34051718836397 | 2.28995122244611e-10 | 2.99097177409172e-08 |
| rpl27a.S | 4743.02656425133 | 0.467959019406531 | 0.0738503347839572 | 6.33658629680562 | 2.34911632970322e-10 | 3.05184128950856e-08 |
| mafa.S | 203.603009756091 | -0.966322444502968 | 0.152656121613811 | -6.33006023137131 | 2.45065517344386e-10 | 3.16682004168325e-08 |
| aldoa.L | 10787.9702958996 | -0.442592557290601 | 0.069940267868077 | -6.32815073178486 | 2.48116712996037e-10 | 3.18928435213002e-08 |
| LOC779056 | 3669.58983192954 | -0.487110715413934 | 0.0769996258923386 | -6.32614392302393 | 2.51363383990647e-10 | 3.21401160561514e-08 |
| grik5.L | 1069.86793866257 | -0.622630379062149 | 0.0985460716367484 | -6.31816538925299 | 2.64686662277576e-10 | 3.36664804888556e-08 |
| nasp.L | 3108.71028941369 | 0.393595134954361 | 0.0624300904115934 | 6.30457416222594 | 2.8898732551345e-10 | 3.65659275313737e-08 |
| pfkp.S | 5822.53549534496 | -0.437738320529226 | 0.0696336213014226 | -6.28630699291642 | 3.25106755354787e-10 | 4.09230233916539e-08 |
| dnmt1.S | 1932.66877599506 | -0.519446002534208 | 0.0827288047555464 | -6.27890133393209 | 3.40973879320237e-10 | 4.26990691969373e-08 |
| eef1b2.L | 15565.0265451971 | 0.421140466697436 | 0.06709726213544 | 6.27656708372598 | 3.46130082727163e-10 | 4.31224832296087e-08 |
| rpl13.S | 15143.860515686 | 0.377642509653453 | 0.0601821369607297 | 6.27499335724609 | 3.49649220886236e-10 | 4.33386641439296e-08 |
| ubap2.S | 4077.8132983354 | -0.450240759238649 | 0.0717715390635255 | -6.27324932854147 | 3.53589997516515e-10 | 4.36046466988133e-08 |
| LOC108697856 | 1753.5950781691 | -0.484824085621188 | 0.0775050652304463 | -6.25538581484522 | 3.96534380502699e-10 | 4.86535668683464e-08 |
| kcnh4.S | 618.475901253844 | -0.626040891333978 | 0.100279410735493 | -6.24296539780519 | 4.29351910912591e-10 | 5.24154538879924e-08 |
| sp5.L | 268.149167564392 | 1.03531200528829 | 0.166009534271015 | 6.23646111552912 | 4.47580503086759e-10 | 5.43676037099486e-08 |
| rab6a.S | 2047.03370230567 | -0.47103626016696 | 0.0755669986059818 | -6.23335938778007 | 4.56537150981783e-10 | 5.51796693828429e-08 |
| tf.L | 139.834262875133 | 1.21335991347789 | 0.194684906455334 | 6.2324292908463 | 4.59256856088096e-10 | 5.52335943653673e-08 |
| shf.L | 417.858873850691 | -0.760886717475915 | 0.122737485083661 | -6.19930184293149 | 5.67141721722736e-10 | 6.78726157021288e-08 |
| rbmx.L | 5408.67588320708 | 0.371503638812373 | 0.0599360388731697 | 6.19833485490273 | 5.70636368542618e-10 | 6.79560781243841e-08 |
| rpl28.S | 5382.49827610562 | 0.46350286844714 | 0.0747969399420126 | 6.19681592330485 | 5.76168177728076e-10 | 6.79933400729752e-08 |
| LOC108700932 | 1000.70101558318 | -0.671972469642711 | 0.108440156252332 | -6.19671248056009 | 5.76546803944714e-10 | 6.79933400729752e-08 |
| eef1g.L | 31945.0754412483 | 0.359585171973202 | 0.0580997032779158 | 6.18910513627156 | 6.05067184248434e-10 | 7.10120877977365e-08 |
| cct5b | 14651.5501274034 | 0.347854757649762 | 0.0562195150204762 | 6.18743789452945 | 6.11499259550072e-10 | 7.14219375553339e-08 |
| rpl22.L | 7823.52572740096 | 0.411530491871729 | 0.0665218337235928 | 6.1863972899347 | 6.1554758516519e-10 | 7.15507800670007e-08 |
| atp6v1b2.S | 6091.84766067152 | -0.350630049911142 | 0.0568316286702311 | -6.16962874574111 | 6.84505213826476e-10 | 7.91874745938115e-08 |
| rpsa.L | 17842.5320831508 | 0.379875899005706 | 0.0616280483017832 | 6.16400988630228 | 7.09255118906941e-10 | 8.13515676278806e-08 |
| odc1.L | 12564.5541406099 | 0.362678912858376 | 0.0588395382840839 | 6.16386401788745 | 7.09909127237617e-10 | 8.13515676278806e-08 |
| MGC68699 | 4147.03528880073 | -0.488319843899461 | 0.0792408013569543 | -6.16247987826041 | 7.16144335274797e-10 | 8.16808003810606e-08 |

| | | | | | | |
|---|---|---|---|---|---|---|
| slc6a8l.S | 4439.76182220023 | -0.440432358065885 | 0.0716051415157437 | -6.15084823160427 | 7.7069641937318e-10 | 8.70851107546607e-08 |
| rpl27a.L | 7930.03284241934 | 0.435829346692557 | 0.0708530741531003 | 6.15117060059823 | 7.69131306758344e-10 | 8.70851107546607e-08 |
| gpt.S | 546.532585746387 | -0.67926466284301 | 0.110588878426411 | -6.1422511242395 | 8.13600934631274e-10 | 9.15075051200564e-08 |
| cirbp.S | 75904.7312791304 | 0.360002178179366 | 0.0586307206058436 | 6.14016294630848 | 8.24368744462213e-10 | 9.22913100367051e-08 |
| kctd15.L | 522.086582847193 | 0.71110107796458 | 0.115829337052972 | 6.13921391641373 | 8.29308292638633e-10 | 9.24184204649676e-08 |
| cdk5r1l.L | 1582.7935893407 | -0.601375023771577 | 0.0979857517292625 | -6.13737215011825 | 8.38976897669106e-10 | 9.30689714701975e-08 |
| ptprg.L | 1528.13796415227 | -0.581654633793285 | 0.0947847793982983 | -6.13658266111582 | 8.43155012814762e-10 | 9.31073085514628e-08 |
| tlcd3b.L | 1078.94188523793 | -0.570295462583554 | 0.0929786658269925 | -6.13361632489668 | 8.59035523557512e-10 | 9.44317149742362e-08 |
| ptchd4.L | 205.736953661931 | -0.924171839947293 | 0.150711980748392 | -6.13203963850864 | 8.67594787273811e-10 | 9.4943007937072e-08 |
| fau.S | 5580.07948869296 | 0.399671582776615 | 0.0651990520125995 | 6.13002137974921 | 8.78672644615067e-10 | 9.57240951940737e-08 |
| LOC108704587 | 1260.35421044252 | -0.684395649743402 | 0.111706055550155 | -6.12675513760412 | 8.96893367070722e-10 | 9.72728904447148e-08 |
| LOC108701185 | 6215.73765961632 | -0.383743166804864 | 0.0627402104933704 | -6.11638315822057 | 9.57230120734291e-10 | 1.03355326902751e-07 |
| rpl18.S | 11702.0828397391 | 0.389519185340697 | 0.0637170530527775 | 6.11326429390974 | 9.76135066450027e-10 | 1.04930200461668e-07 |
| tfap2a.L | 799.56703265523 | -0.601767702129289 | 0.098597002643561 | -6.10330624658789 | 1.03896496411719e-09 | 1.11192135851379e-07 |
| dchs1.L | 2956.79611622866 | -0.513215317816157 | 0.0841741885411409 | -6.09706284920488 | 1.08035052403252e-09 | 1.14611509305004e-07 |
| MGC115323 | 2104.11665755237 | -0.542718386078424 | 0.0890054020345868 | -6.09758928865383 | 1.07679975806442e-09 | 1.14611509305004e-07 |
| nif.S | 6637.456159759 | -0.684167639116293 | 0.112240435803645 | -6.09555401507159 | 1.09059078311568e-09 | 1.15194836891358e-07 |
| crmp1.S | 5967.96218812764 | -0.488447302000128 | 0.0802716488680646 | -6.08492922330455 | 1.16542781650696e-09 | 1.22038376613017e-07 |
| sap130.L | 1823.67372456005 | -0.441331703769953 | 0.0725210547385796 | -6.08556653458564 | 1.16080105509398e-09 | 1.22038376613017e-07 |
| tacc2.L | 2393.78133916612 | -0.414250657994731 | 0.068134976994144 | -6.07985320124691 | 1.2029262663213e-09 | 1.25424423665278e-07 |
| nell1.L | 1504.13607934905 | -0.517690467293292 | 0.0853002244753226 | -6.06903991727548 | 1.28677163474218e-09 | 1.33593291001823e-07 |
| LOC108708617 | 1765.60179817989 | -0.468576564954552 | 0.0773276521026724 | -6.05962488467122 | 1.36439360492107e-09 | 1.4104926909767e-07 |
| rps15.L | 7335.50771594136 | 0.396611283373802 | 0.0655874512944032 | 6.04706045968348 | 1.47512413287207e-09 | 1.50574225563001e-07 |
| rpl5.L | 24314.2463532009 | 0.39046717672138 | 0.0645605742926219 | 6.04807471122516 | 1.46586981627453e-09 | 1.50574225563001e-07 |
| mafa.L | 358.564980671543 | -0.864707446055619 | 0.142991445573414 | -6.04726697172711 | 1.47323525393216e-09 | 1.50574225563001e-07 |
| LOC121402164 | 59.9571557277156 | 1.15738264218028 | 0.191420192191727 | 6.04629338696432 | 1.48216091355618e-09 | 1.50659486334451e-07 |
| LOC108695345 | 15850.9561229299 | 0.399585679027294 | 0.0662345722204624 | 6.03288683887426 | 1.61056202363543e-09 | 1.62764280560923e-07 |
| LOC108703204 | 442.045448641806 | -0.73024998458808 | 0.121053074383637 | -6.0324778061712 | 1.61464524636463e-09 | 1.62764280560923e-07 |
| ptma.S | 17384.0466780053 | 0.380586830803753 | 0.0631073721481734 | 6.03078242443928 | 1.63167744727732e-09 | 1.6380153679403e-07 |
| copa.L | 3982.83438331146 | -0.423358129627803 | 0.0702332204752611 | -6.02789003213838 | 1.66114008893151e-09 | 1.66072993088486e-07 |
| clstn3.S | 3925.21453233741 | -0.55095751533231 | 0.0914356626601529 | -6.02563047396619 | 1.68451660004565e-09 | 1.67719861809463e-07 |
| slc45a4.S | 449.703398084661 | -0.772345180425736 | 0.128296076653262 | -6.02002181651358 | 1.74393563971088e-09 | 1.729272344128e-07 |
| pcdh7.S | 710.14943331988 | -0.702944673238039 | 0.116870313195591 | -6.01474107510614 | 1.80174450645435e-09 | 1.77933256259358e-07 |
| h2az2.S | 4031.28746956935 | 0.414687961440364 | 0.0689648029532628 | 6.01303771898538 | 1.82078649357225e-09 | 1.79085777630948e-07 |
| rps23.S | 8573.13768914038 | 0.39113595121544 | 0.0650751582704568 | 6.010526314663 | 1.84921988753534e-09 | 1.81148983660417e-07 |
| LOC108701719 | 1845.7713291334 | -0.577111851150852 | 0.0960951126544703 | -6.00563166230915 | 1.90588429084018e-09 | 1.85950011894262e-07 |
| uhrf1.L | 1616.73954009076 | 0.518673961226267 | 0.0864529001994917 | 5.99949753020913 | 1.97929039832781e-09 | 1.92339523747903e-07 |
| snca.L | 1563.85246507791 | 0.550716079168696 | 0.0920535454265447 | 5.98256239471132 | 2.19654324953607e-09 | 2.12388053335117e-07 |
| map7d1.S | 1276.69255307011 | -0.542548564933647 | 0.0906956682693521 | -5.98207803400667 | 2.20308674736353e-09 | 2.12388053335117e-07 |
| cck.L | 229.191483771595 | 1.01224324864328 | 0.169311376001176 | 5.97858969993988 | 2.25077665557992e-09 | 2.16127937038176e-07 |
| LOC108698386 | 3767.62224031122 | -0.541883911127763 | 0.0906600510584885 | -5.9770969109445 | 2.27149083778815e-09 | 2.16548391318221e-07 |
| cdca7.L | 1616.41831800724 | 0.433436086259047 | 0.0725174531516315 | 5.97698991652047 | 2.2729826206531e-09 | 2.16548391318221e-07 |
| apcdd1.L | 351.706090933081 | 0.929400858223221 | 0.155638603632979 | 5.97153171853751 | 2.35036352880269e-09 | 2.23045826440362e-07 |
| rps27a.L | 14427.889620882 | 0.365930168552277 | 0.0613620095967361 | 5.96346454356902 | 2.46945057637384e-09 | 2.33435145145627e-07 |
| rpl5.S | 15774.2757514601 | 0.353679013769451 | 0.0593327514528522 | 5.9609407133343 | 2.50789952086464e-09 | 2.36150817673975e-07 |
| LOC108704616 | 1807.82432171125 | -0.600472594494171 | 0.100778212428578 | -5.95835726814196 | 2.54786035752024e-09 | 2.38987334075663e-07 |
| syn1.L | 9925.97837491991 | -0.614355602051786 | 0.103210102223975 | -5.95247547297823 | 2.6411672620905e-09 | 2.46786605635487e-07 |
| per1.L | 453.495133756901 | -0.677217771128265 | 0.113799706214252 | -5.95096238520387 | 2.66570349965720e-09 | 2.48124907358896e-07 |
| atp1b1.L | 6157.47075481706 | -0.49330934956394 | 0.0829861589219495 | -5.94447744024288 | 2.77340160251333e-09 | 2.57164192868164e-07 |
| LOC108696402 | 2419.3474850894 | -0.425492463269874 | 0.0716335917796211 | -5.93984543702473 | 2.85290925776035e-09 | 2.63530712958289e-07 |
| pabpc1.L | 5304.60639615379 | -0.376331646273295 | 0.0634038715403179 | -5.93546793170177 | 2.93008594138057e-09 | 2.69634499469316e-07 |
| rpl30.S | 6467.75535222168 | 0.368612289514371 | 0.0621193518632168 | 5.9339365022036 | 2.95756252777336e-09 | 2.71135939810287e-07 |
| dnm1l.S | 2565.52166527264 | -0.428640981693634 | 0.0722702649345407 | -5.93108358025087 | 3.00941964037601e-09 | 2.74852784749229e-07 |
| LOC108699962 | 9780.41967009096 | -0.452138247720197 | 0.0762534388330564 | -5.92941452398068 | 3.0401671775524e-09 | 2.76621053975498e-07 |
| LOC108709856 | 1958.39995200388 | -0.481546038858379 | 0.081240134922118 | -5.92744016366629 | 3.07693406798747e-09 | 2.7892177704361e-07 |
| rpl12.L | 11668.5553713492 | 0.352030485080583 | 0.0594116384880811 | 5.92527817847014 | 3.11769168449047e-09 | 2.81565805884801e-07 |
| khdrbs1.S | 11001.2083853616 | 0.373545766622064 | 0.0630811354817197 | 5.92167157058093 | 3.1868558820405e-09 | 2.86746210364044e-07 |
| slc4a4.S | 878.305920973088 | -0.683605635604958 | 0.115477501825187 | -5.91981662921509 | 3.22300769525037e-09 | 2.88928962909271e-07 |
| ahcy.L | 1781.43292189467 | 0.459046558108747 | 0.0777094109537071 | 5.90721963369674 | 3.47929755378755e-09 | 3.10757554307774e-07 |
| atp1a1.S | 5253.04099474539 | -0.576902713421233 | 0.0976833661012345 | -5.90584391638755 | 3.50846034591576e-09 | 3.11391133978268e-07 |
| neo1.L | 4232.73438862758 | -0.44280432823471 | 0.0749946678590599 | -5.90447748987819 | 3.53766168510752e-09 | 3.11391133978268e-07 |
| rabl6.L | 2301.7011995014 | -0.444783562592041 | 0.0753251272579576 | -5.90484979957404 | 3.52968183964311e-09 | 3.11391133978268e-07 |
| emilin3.L | 381.855245383817 | -0.775595221528216 | 0.131339309325833 | -5.90527866721213 | 3.52051148463108e-09 | 3.11391133978268e-07 |
| LOC108719231 | 1227.62475483294 | -0.599449978357704 | 0.101665642520768 | -5.89628869197624 | 3.71768309028406e-09 | 3.26055570380364e-07 |
| meis2.L | 3194.59108194245 | -0.709334986800153 | 0.120349525348849 | -5.89395749375873 | 3.77054200506481e-09 | 3.29501969320304e-07 |
| LOC108703011 | 463.527555582088 | -0.878631159258192 | 0.149088556123743 | -5.89335078494508 | 3.78441839912806e-09 | 3.2952924942085e-07 |
| dbi.S | 2398.50732205432 | 0.419292976573887 | 0.071321126240919 | 5.87894497287575 | 4.12889588033166e-09 | 3.58240701845634e-07 |
| rps15.S | 9999.56868322357 | 0.387486158588164 | 0.0659186053146887 | 5.87825177335511 | 4.14621962920443e-09 | 3.58463557551219e-07 |
| tmem255a.L | 857.757093475842 | -0.556512043167136 | 0.0946851183992645 | -5.8775027435722 | 4.16501815032826e-09 | 3.58811882780407e-07 |
| h3c13.S | 6002.12039698048 | 0.425902488883901 | 0.072476159131021 | 5.87644949719208 | 4.19159203780342e-09 | 3.59825218962531e-07 |
| hid1.L | 880.46482782387 | -0.551489953717388 | 0.0939830290658904 | -5.86797381611041 | 4.41152897293625e-09 | 3.77372129818709e-07 |
| ckap5.L | 4779.26142999743 | -0.348009681622441 | 0.0593465283711688 | -5.86402762173209 | 4.51772276066068e-09 | 3.85100199114002e-07 |
| snap91.S | 6378.83134708475 | -0.488280050617853 | 0.0833725013960833 | -5.8566079035838 | 4.72416690712381e-09 | 4.01289898047782e-07 |
| rpl3.L | 47787.1206270069 | 0.331613349294174 | 0.0566725952323072 | 5.85138809921893 | 4.87487214717202e-09 | 4.12648585168631e-07 |
| wnt8b.S | 89.9905817465224 | 1.09528622242775 | 0.187240178372814 | 5.84963244505632 | 4.92660458760878e-09 | 4.15579624483916e-07 |
| coasy.L | 1168.90767663364 | -0.47338472945224 | 0.0809638724024339 | -5.84686373570749 | 5.00927514308495e-09 | 4.2109110839483e-07 |
| LOC108704046 | 1919.72905148548 | -0.594746588967236 | 0.10180465233063 | -5.84203742512358 | 5.15662175173755e-09 | 4.31982651161076e-07 |
| slc6a8l.L | 2939.6606220548 | -0.484454039232798 | 0.0829549659281792 | -5.83996429643801 | 5.22120104331235e-09 | 4.35889546894262e-07 |
| LOC108714021 | 692.265469871481 | 0.68815308038224 | 0.117849816496723 | 5.83923760629167 | 5.24402362512349e-09 | 4.36295582016267e-07 |
| eif3e.L | 6635.59599415033 | 0.341604954254979 | 0.0585183882739097 | 5.83756600841453 | 5.29689121514373e-09 | 4.39190017681576e-07 |
| ldha.L | 1518.02732240304 | -0.473981685630721 | 0.0812438027606992 | -5.83406573208812 | 5.4092795151597e-09 | 4.43962961288141e-07 |
| ptgds.S | 17327.193745525 | -0.590968183366172 | 0.101293095125069 | -5.83423956624576 | 5.40364364020031e-09 | 4.43962961288141e-07 |
| rack1.S | 18191.540109843 | 0.377023729631465 | 0.0646218411176806 | 5.83430807774233 | 5.40142399966641e-09 | 4.43962961288141e-07 |
| copa.S | 2288.9770041978 | -0.412766784802285 | 0.0707845279631194 | -5.83131365963685 | 5.49927005997199e-09 | 4.49829181269224e-07 |
| atp2b1.L | 4435.91019726829 | -0.38112463922334 | 0.0653785195747435 | -5.82950855575159 | 5.55908468912301e-09 | 4.5319598468978e-07 |
| foxd3.L | 220.167374959166 | -0.831968264923368 | 0.142820632383506 | -5.82526663717145 | 5.70214864255703e-09 | 4.63304344890571e-07 |
| celf4.L | 3125.67160532629 | -0.507466578708973 | 0.0871527842605959 | -5.82272365724536 | 5.7896242698621e-09 | 4.68675916637043e-07 |
| rpl39.S | 5592.95924475436 | 0.381816607679896 | 0.0655791289685059 | 5.8222274934951 | 5.80684329084342e-09 | 4.68675916637043e-07 |
| snrpe.S | 2716.41684218184 | 0.410340395058782 | 0.0705013993038246 | 5.82031561232462 | 5.87366085016859e-09 | 4.72499061900648e-07 |
| mab21l1.L | 3048.88824022623 | -0.457637087760482 | 0.0787210073815001 | -5.8134048709853 | 6.12147652072992e-09 | 4.87590657687254e-07 |
| rtn4.L | 2953.35444611283 | -0.418816789712319 | 0.0720405562192939 | -5.81362515355138 | 6.11342265665001e-09 | 4.87590657687254e-07 |
| LOC108701569 | 9444.87898547903 | 0.392007371869919 | 0.0674189748663672 | 5.81449618074031 | 6.08167741915902e-09 | 4.87590657687254e-07 |
| hes5.3.S | 832.521883308395 | 0.569542621969866 | 0.0979828514165464 | 5.81267654223101 | 6.14817886424297e-09 | 4.88117180810192e-07 |
| rpl10a.S | 24159.4309449792 | 0.382126103457832 | 0.0658267796663597 | 5.80502502772614 | 6.43563901288527e-09 | 5.08448231682957e-07 |
| hcfc1.S | 4425.91439486723 | -0.357265107945402 | 0.0615470025436863 | -5.80475235478469 | 6.44612066182394e-09 | 5.08448231682957e-07 |
| ipo7.L | 3440.41456297543 | -0.367885826383025 | 0.0633911890256561 | -5.80342208495462 | 6.4974952916759e-09 | 5.10841911378558e-07 |
| ptmap12.S | 13819.8755718935 | 0.412701027322901 | 0.0711307920645278 | 5.80200241477011 | 6.55276179955006e-09 | 5.13525145671836e-07 |
| ttbk1.L | 2485.8970618217 | -0.499963009488799 | 0.0862614886574848 | -5.79590054924722 | 6.79555268129501e-09 | 5.29138323203144e-07 |
| LOC121397179 | 278.254795621355 | -0.970117550226211 | 0.167365170625404 | -5.7964123993129 | 6.77485469602381e-09 | 5.29138323203144e-07 |
| plxnb1.L | 2853.34623535765 | -0.612466657035479 | 0.105706040019983 | -5.79405544772744 | 6.87067580409039e-09 | 5.33278587810134e-07 |
| nsf.L | 6829.94308180836 | -0.368549356006405 | 0.0636915172549282 | -5.78647474405865 | 7.18788879859867e-09 | 5.56122835901771e-07 |
| LOC108719876 | 184.823871235878 | -0.952881843902417 | 0.16473445645553 | -5.78435055060655 | 7.27930045640426e-09 | 5.61407381866302e-07 |
| rpsa.S | 30766.5969457726 | 0.393995419164802 | 0.0681413541196679 | 5.78203095983209 | 7.38041205589284e-09 | 5.67404210631666e-07 |
| aldh4a1.L | 887.458708778824 | -0.538906219356483 | 0.0932535100432873 | -5.77893764112824 | 7.51737803733554e-09 | 5.76110984350251e-07 |
| LOC100126660 | 961.002374124018 | -0.745920082836038 | 0.129087658033992 | -5.77839968744046 | 7.54144852000604e-09 | 5.7613820863216e-07 |
| caprin1.S | 4307.28145680405 | -0.356103854417476 | 0.0617533890352474 | -5.76654755278646 | 8.09118871737805e-09 | 6.16198553891844e-07 |
| rplp0.S | 8020.45013236367 | 0.421286548775567 | 0.0731291196939116 | 5.76085901948362 | 8.36869195183768e-09 | 6.35340632118577e-07 |
| LOC108703406 | 2665.13508551477 | -0.488366571974223 | 0.0847879481673875 | -5.7598583587622 | 8.41845487622483e-09 | 6.37127547548305e-07 |

| | | | | | | |
|---|---|---|---|---|---|---|
| rpl34.L | 8567.12325492785 | 0.386855180455877 | 0.067173252145989 | 5.75906582005463 | 8.45807192754747e-09 | 6.38137886359746e-07 |
| ptprg.S | 1275.42950185153 | -0.513819960404965 | 0.0892937120338487 | -5.75426811923991 | 8.70179553202283e-09 | 6.54493562399265e-07 |
| rps5.S | 12050.0678788244 | 0.396198699304339 | 0.0689199013645151 | 5.74868349286888 | 8.99410022550214e-09 | 6.74390959501077e-07 |
| pcdh7.L | 997.702206539428 | -0.557071974832535 | 0.0969464817180742 | -5.74618041789832 | 9.12819083612895e-09 | 6.82339286685898e-07 |
| LOC108706667 | 231.305747652099 | 0.902202638502425 | 0.15704688008579 | 5.74479822846262 | 9.2030659846906e-09 | 6.85826027705747e-07 |
| crb1.L | 140.59079979943 | 1.03945588171445 | 0.18118081067558 | 5.7371190571373 | 9.63005640545697e-09 | 7.15451346526519e-07 |
| stra6.S | 42.779526625875 | 1.00150682246257 | 0.174676479962301 | 5.73349556092904 | 9.83816422966752e-09 | 7.22079642886836e-07 |
| qser1.L | 1437.85133703365 | -0.504876924871278 | 0.08805529711461 | -5.73363490232899 | 9.83008124547591e-09 | 7.22079642886836e-07 |
| bzw1.S | 5246.50982577265 | 0.333393493037712 | 0.0581481863197069 | 5.73351490628905 | 9.83704164847283e-09 | 7.22079642886836e-07 |
| crebbp.S | 1675.4467263477 | -0.456570815905398 | 0.0796299446858597 | -5.73365732836524 | 9.82878094817131e-09 | 7.22079642886836e-07 |
| LOC108701252 | 991.963058314587 | -0.563565661893352 | 0.0983161969493798 | -5.73217515912984 | 9.9150801014381e-09 | 7.25532999952823e-07 |
| rrm1.S | 4440.23299471438 | 0.416109250683887 | 0.0726061234093388 | 5.73104899621132 | 9.98114277362735e-09 | 7.28173821448957e-07 |
| rpl28.L | 5877.54966945465 | 0.377279407022079 | 0.0659927806958045 | 5.71697999454151 | 1.08433877128863e-08 | 7.88710362565447e-07 |
| plxna1.S | 3142.90404525285 | -0.512730596456738 | 0.0897088066244627 | -5.71549902121785 | 1.09382550862212e-08 | 7.93235728551218e-07 |
| LOC121396917 | 24.6983791967832 | 1.05648632768624 | 0.184933817038894 | 5.71278062932131 | 1.1114491745962e-08 | 8.01232826339472e-07 |
| atp1a2.S | 15253.600974817 | -0.457426527401408 | 0.0800693496047617 | -5.71287926852606 | 1.11080488701044e-08 | 8.01232826339472e-07 |
| LOC108701642 | 3421.57377566064 | -0.572077751966787 | 0.100196220637734 | -5.7095741568454 | 1.13259200254988e-08 | 8.14058878992512e-07 |
| LOC108700360 | 1459.42010899536 | -0.497080145473269 | 0.0870799228392285 | -5.70832092250476 | 1.14096136855421e-08 | 8.17655324119646e-07 |
| dact1.L | 1483.70647541058 | -0.460234074732923 | 0.0807965306317623 | -5.69621085378632 | 1.22499365309764e-08 | 8.75293994363354e-07 |
| rpl14.S | 6791.43323070543 | 0.384521657399971 | 0.0675262647746972 | 5.69440141081305 | 1.23805499356724e-08 | 8.82032493071045e-07 |
| LOC108700185 | 63.7729736106881 | 1.08495311899496 | 0.190655291916159 | 5.69065305290386 | 1.26554417395685e-08 | 8.98980414096714e-07 |
| rps8.S | 19445.8930650487 | 0.351516626470099 | 0.0618291848805919 | 5.68528644116801 | 1.30593585469205e-08 | 9.24968094865559e-07 |
| vcp.L | 7938.74593843303 | -0.366416425870487 | 0.0644772930522541 | -5.68287545157229 | 1.32448727134894e-08 | 9.32669384642062e-07 |
| rps17.L | 11178.7807188951 | 0.360937514229543 | 0.0635111160684899 | 5.68306048724307 | 1.32305448605501e-08 | 9.32669384642062e-07 |
| adap1.L | 161.973269659226 | -0.87668857050087 | 0.154307810337424 | -5.68142706829423 | 1.33575472775564e-08 | 9.37885125898716e-07 |
| mapk8ip2.S | 2793.90402780464 | -0.490137543545198 | 0.0862945086168448 | -5.67982310115991 | 1.34834119348964e-08 | 9.43994263822399e-07 |
| mafb.L | 600.166803202925 | -0.765932089943113 | 0.134890792906227 | -5.67816433902624 | 1.36147881203044e-08 | 9.50453053432972e-07 |
| rnf157.L | 2657.27106872211 | -0.439688930371055 | 0.0774671396487474 | -5.6758121232396 | 1.38032209810867e-08 | 9.60846563078853e-07 |
| rusc2.L | 2224.48935396068 | -0.514880919346164 | 0.0907814038271578 | -5.67165628245257 | 1.41423503966158e-08 | 9.76063240157339e-07 |
| snrpg.S | 2334.02138016181 | 0.452112101918319 | 0.0797009011686746 | 5.67260966047917 | 1.40638435642112e-08 | 9.76063240157339e-07 |
| nt5dc2.L | 1776.52578497996 | -0.421578815069709 | 0.0743253208698516 | -5.67207527846292 | 1.41077954706306e-08 | 9.76063240157339e-07 |
| mafb.S | 433.896196646311 | -0.715431633095134 | 0.126157037695448 | -5.67096094014382 | 1.41998773392576e-08 | 9.77257280679669e-07 |
| rps20.L | 17266.705943628 | 0.36095163045361 | 0.0637698889898836 | 5.66022046095878 | 1.51178656619988e-08 | 1.03667802840348e-06 |
| hes5.1.S | 833.754109701168 | 0.598633217538423 | 0.105767984673291 | 5.65987164629784 | 1.51486251783664e-08 | 1.03667802840348e-06 |
| pabpc1.S | 24774.3911268508 | -0.342212964208647 | 0.0604769037555757 | -5.65857282627662 | 1.52636944928687e-08 | 1.04161852249931e-06 |
| sfxn5.L | 367.852237517535 | -0.748634438067727 | 0.132378012480295 | -5.65527782175431 | 1.55594384256617e-08 | 1.05718548455177e-06 |
| rpl35a.S | 1942.8049946459 | 0.514092052921448 | 0.0909082681397247 | 5.65506376308144 | 1.5578842655369e-08 | 1.05718548455177e-06 |
| LOC108714603 | 2205.59303406798 | -0.520125235733099 | 0.0920887262523921 | -5.64808806571573 | 1.62242065662737e-08 | 1.09791329894444e-06 |
| rpl18a.L | 11874.2405232799 | 0.371694690972451 | 0.0658253253882709 | 5.64668216647632 | 1.63573830015197e-08 | 1.10169465067848e-06 |
| rps5.L | 12024.906847872 | 0.362465825249704 | 0.0641925390385632 | 5.64654133764604 | 1.63707816290002e-08 | 1.10169465067848e-06 |
| rpl35.L | 8651.71563445942 | 0.360778063831273 | 0.0639279121526867 | 5.64351394692171 | 1.66614022857843e-08 | 1.11815499207416e-06 |
| prrc2c.L | 5299.84020641828 | -0.357861501283538 | 0.0634695393864464 | -5.63831886512726 | 1.71718318028237e-08 | 1.14923548710137e-06 |
| LOC108708523 | 44251.5911427682 | 0.329345373398553 | 0.0584276378309553 | 5.63680794954309 | 1.73231118355409e-08 | 1.15300733954146e-06 |
| spon1.S | 2151.43527486933 | -0.551233444904425 | 0.0977897920248373 | -5.6369221520015 | 1.73116322734361e-08 | 1.15300733954146e-06 |
| hnrnpdl.L | 23541.5208727984 | 0.336882216382206 | 0.0597964129756675 | 5.63381981657146 | 1.76261180802402e-08 | 1.16996970667037e-06 |
| efnb3.L | 1141.62962760704 | -0.462467165101769 | 0.0821047250613437 | -5.63264982321348 | 1.77461559219989e-08 | 1.17472782552872e-06 |
| ncs1.L | 531.737599934871 | -0.64684652559256 | 0.114861735815895 | -5.63152316128457 | 1.78624981625253e-08 | 1.17921611510975e-06 |
| vwa5a.2.L | 1095.76666750352 | -0.702384600322449 | 0.124793191534982 | -5.6283887901493 | 1.81900719924601e-08 | 1.18792905641082e-06 |
| pum2.L | 1954.3756354911 | -0.435724645816221 | 0.0774109919345789 | -5.62871802733722 | 1.81553910255206e-08 | 1.18792905641082e-06 |
| rpl18.L | 12901.1394127313 | 0.375759861112884 | 0.0667485376160334 | 5.62948454802738 | 1.80748963536687e-08 | 1.18792905641082e-06 |
| rpl22.S | 5355.23157879907 | 0.391794470081143 | 0.0696101218520703 | 5.62841235810149 | 1.81875872710973e-08 | 1.18792905641082e-06 |
| slc25a11.S | 1516.60559641231 | -0.556223219037511 | 0.0988630264993037 | -5.6262006002965 | 1.84222100940045e-08 | 1.19986373196715e-06 |
| rps12.L | 23853.0894192628 | 0.401831331711002 | 0.0714841113094518 | 5.62126778035318 | 1.89561179311559e-08 | 1.23133670860829e-06 |
| cpne7.S | 1005.71155268006 | 0.738785574283466 | 0.131458065423401 | 5.61993341301639 | 1.91031070400091e-08 | 1.23428426178186e-06 |
| LOC108701181 | 4444.74046595002 | -0.464732360428737 | 0.0826893632809384 | -5.62021936061838 | 1.90715152083013e-08 | 1.23428426178186e-06 |
| LOC108700022 | 8306.04041578457 | 0.373526494005862 | 0.0664830166472475 | 5.61837462923438 | 1.92762188200887e-08 | 1.24216567643299e-06 |
| rpl10.L | 28852.2121535939 | 0.349810009314083 | 0.062293748534101 | 5.61549140236101 | 1.96004403197314e-08 | 1.25971718816813e-06 |
| MGC85536 | 3803.74064217 | 0.388287687909725 | 0.0692207340113653 | 5.60941303867094 | 2.03013976406801e-08 | 1.30132494533689e-06 |
| kiaa1549.L | 1140.1927550641 | -0.611351348516715 | 0.109070901370882 | -5.60508202309518 | 2.08156365175078e-08 | 1.33077650935878e-06 |
| sp8.S | 654.359337636006 | -0.674490154753022 | 0.120367040773773 | -5.60361167323796 | 2.09930753239133e-08 | 1.33859782655945e-06 |
| zmiz2.S | 2737.45261721985 | -0.402957139908834 | 0.0719327153389814 | -5.60186193458577 | 2.12061440879233e-08 | 1.34512288373892e-06 |
| rps21.S | 4661.8684124698 | 0.38949650954395 | 0.069524509668478 | 5.6022906368018 | 2.11537469156625e-08 | 1.34512288373892e-06 |
| abr.L | 4796.58780671422 | -0.365672700918541 | 0.0653153755686313 | -5.5985699800247 | 2.16127157353551e-08 | 1.3637904313629e-06 |
| abca3.L | 1824.94214150643 | -0.451701603794936 | 0.0806802852600236 | -5.59866146158447 | 2.16013158500494e-08 | 1.3637904313629e-06 |
| pcdh10.L | 2091.49268154722 | -0.492935406457489 | 0.0880830299192754 | -5.59625851777856 | 2.19027034624638e-08 | 1.37850849201132e-06 |
| rps4x.L | 20396.3329923059 | 0.352162625804575 | 0.0629353562100319 | 5.59562457435397 | 2.19828934396441e-08 | 1.37998039592432e-06 |
| hmgn2.L | 19608.6012626183 | 0.39281700447519 | 0.0702360804429772 | 5.59280930823166 | 2.2342463733443e-08 | 1.3989376647945e-06 |
| rpl38.S | 3713.93861247788 | 0.352739483888824 | 0.063109132696076 | 5.58935717889776 | 2.27911728500523e-08 | 1.42336440416239e-06 |
| pfkp.L | 4700.48537482796 | -0.412026750960767 | 0.0737340773991389 | -5.58800985235599 | 2.29686616204768e-08 | 1.43077093694324e-06 |
| ncor1.S | 2751.65340693034 | -0.442758068093466 | 0.0792891636443967 | -5.584093055631 | 2.3492290153055e-08 | 1.45221805846901e-06 |
| LOC108715150 | 1241.78025622478 | -0.577963023527912 | 0.103500858129199 | -5.58413750354078 | 2.34862835136991e-08 | 1.45221805846901e-06 |
| LOC108704466 | 6234.03756151266995 | -0.370375615126995 | 0.0663228625058904 | -5.58443349899291 | 2.34463210292421e-08 | 1.45221805846901e-06 |
| sim1.L | 31.2769383303894 | 1.07411118990579 | 0.192376381219423 | 5.58338390137751 | 2.35883263533933e-08 | 1.45272586292573e-06 |
| trappc9.L | 875.448999481692 | -0.530355421367677 | 0.0949921516744332 | -5.58314989206019 | 2.36201002657308e-08 | 1.45272586292573e-06 |
| pafah1b1.L | 1644.90063635414 | -0.527950082177178 | 0.0946218986722746 | -5.57957607684186 | 2.41105475804088e-08 | 1.47914556292538e-06 |
| nlgn2.S | 4665.7692377746 | -0.497935233000738 | 0.0892903008135011 | -5.57658814523164 | 2.4528165541888e-08 | 1.50097545006203e-06 |
| mtss2.L | 5572.2083234799 | -0.455790965104724 | 0.0817823578167543 | -5.57321869009931 | 2.50075321685454e-08 | 1.52263390639164e-06 |
| LOC108715931 | 3769.93051865929 | -0.394259561607972 | 0.0707409490054009 | -5.57328629529513 | 2.49978253131394e-08 | 1.52263390639164e-06 |
| pygm.S | 3615.36425510715 | -0.441309441573631 | 0.0792054202272822 | -5.57170759661752 | 2.52254543755206e-08 | 1.53206797149725e-06 |
| gnat2.S | 59.0431770066415 | 1.04301208036064 | 0.187241247589469 | 5.57041834418595 | 2.54128398384635e-08 | 1.53959982801903e-06 |
| svil.L | 1023.16151942361 | -0.541753282177585 | 0.0972807632369015 | -5.56896619795518 | 2.56255187872897e-08 | 1.54862276969755e-06 |
| naca.S | 8431.92389235302 | 0.319293990448702 | 0.0573524152102864 | 5.5672283247007 | 2.58823160216987e-08 | 1.56026050975471e-06 |
| lmo3.S | 1303.64636181486 | 0.601391967924575 | 0.108135342912677 | 5.56147464580771 | 2.67504547650716e-08 | 1.60860284173923e-06 |
| rpl11.L | 11220.4958782437 | 0.376052727597831 | 0.067645886653854 | 5.55913664820603 | 2.71112377729351e-08 | 1.6262726184091e-06 |
| trim9.L | 1247.65588700726 | -0.471642165824895 | 0.0848633445851666 | -5.55766648286608 | 2.73405164204891e-08 | 1.63196684501072e-06 |
| snrpa.S | 2295.13843107244 | 0.450646704793283 | 0.0810829080076751 | 5.55785079576359 | 2.73116692028942e-08 | 1.63196684501072e-06 |
| LOC108699470 | 383.043003478842 | -0.664551167251813 | 0.119603153307658 | -5.55630139234181 | 2.75550920599046e-08 | 1.6407436433905e-06 |
| LOC108708306 | 3022.91439894573 | 0.409851956127487 | 0.0737774008115206 | 5.5552506813643 | 2.77213631000497e-08 | 1.64653659666377e-06 |
| snx27.L | 2332.88259823942 | -0.373162507269704 | 0.0671780038119734 | -5.55483173203776 | 2.77879313670925e-08 | 1.64653659666377e-06 |
| slc20a1.S | 3978.71463168777 | -0.4102332787981 | 0.0738648054459454 | -5.55383956298796 | 2.79461996669026e-08 | 1.650457567717e-06 |
| copb1.S | 1945.94433084692 | -0.402416904124715 | 0.0724609913602247 | -5.5535660852911 | 2.79899776858239e-08 | 1.650457567717e-06 |
| rpl7a.S | 19029.3232718484 | 0.338922331440254 | 0.0610398197291503 | 5.55247923313243 | 2.81646184724043e-08 | 1.65673424011765e-06 |
| stac2.L | 761.25664218696 | -0.63069993102818 | 0.113621072313021 | -5.55090634324087 | 2.84192317342649e-08 | 1.66767346800056e-06 |
| pmch.S | 46.9419430802733 | 0.973761068420136 | 0.175448598561141 | 5.5501216675766 | 2.85470855534632e-08 | 1.67113950948394e-06 |
| chrm2.L | 459.613098772266 | -0.650602794664501 | 0.117320106157381 | -5.54553533894472 | 2.93056136753916e-08 | 1.70323104935398e-06 |
| plcb4.S | 813.224002844635 | -0.570818031863902 | 0.102918968509459 | -5.54628597751096 | 2.91801417971958e-08 | 1.70323104935398e-06 |
| asap1.L | 2630.73688311983 | -0.504103770837028 | 0.0908982554698656 | -5.5458024824706 | 2.92608998152304e-08 | 1.70323104935398e-06 |
| fgd1.S | 886.132280581251 | -0.550973824722242 | 0.0994383721893141 | -5.5408572424464 | 3.00994524043053e-08 | 1.74519354823435e-06 |
| fbxl16.L | 1730.01702861097 | -0.404486535362493 | 0.0730472962041561 | -5.53732384881179 | 3.07128336439648e-08 | 1.77651804892019e-06 |
| rpl9.L | 12451.2792063293 | 0.392083695457075 | 0.0708337224199436 | 5.53526882482011 | 3.10751358402178e-08 | 1.78895580592951e-06 |
| lrit1.S | 93.4491673967911 | 1.07317154308283 | 0.19387270164078 | 5.53544430959275 | 3.10440365388917e-08 | 1.78895580592951e-06 |
| LOC108701384 | 230.583793203195 | -0.854382069966279 | 0.154485623452487 | -5.53049565954626 | 3.19327199888423e-08 | 1.83397990404004e-06 |
| nedd4l.S | 1660.41490479838 | -0.431497829755391 | 0.0780443897209573 | -5.52887698011585 | 3.22287239882842e-08 | 1.84027681760025e-06 |
| LOC108696609 | 441.053341633444 | -0.684470061913099 | 0.123786942925601 | -5.52942051670574 | 3.21290331624225e-08 | 1.84027681760025e-06 |
| megf6.L | 291.23163007235 | -0.761084794085624 | 0.13766184706815 | -5.52865452770544 | 3.22696107803452e-08 | 1.84027681760025e-06 |
| rpl15.S | 12253.4383172509 | 0.38233441449086 | 0.0692643511761419 | 5.51993064251145 | 3.39133492666159e-08 | 1.92948690183411e-06 |
| LOC108704607 | 7516.6980193946 | -0.348581436634138 | 0.0631782849846976 | -5.51742480376871 | 3.4400319885748e-08 | 1.95262002641206e-06 |
| agpat3.L | 1004.03937491954 | -0.538292420553485 | 0.0976267762291167 | -5.51377850775474 | 3.51210551075815e-08 | 1.98888324658179e-06 |

| | | | | | | |
|---|---|---|---|---|---|---|
| necap1.L | 1835.80067309597 | -0.408117613356023 | 0.0740975005510688 | -5.50784588307056 | 3.63251202787755e-08 | 2.05228481872691e-06 |
| slmap.L | 1657.55913752151 | -0.466295016484227 | 0.0847424426867867 | -5.50249676195531 | 3.7445010618852e-08 | 2.11064753590346e-06 |
| kars1.S | 3988.79173447696 | 0.363816930834061 | 0.0661646100401323 | 5.49866356974502 | 3.82680460463348e-08 | 2.15204608946686e-06 |
| rps4x.S | 17264.7002112582 | 0.374626155147281 | 0.0681423340335701 | 5.49770066523877 | 3.84775350564099e-08 | 2.15882964586703e-06 |
| ipo5.S | 4902.44882997532 | 0.354777768521507 | 0.0645653709070589 | 5.49486146423918 | 3.91017246749511e-08 | 2.18879562039922e-06 |
| pmel.L | 560.914473688289 | 0.859829577691587 | 0.156496084673501 | 5.49425616292867 | 3.92360625041818e-08 | 2.19126644247492e-06 |
| ilf2.S | 9015.390692995 | 0.334401256299516 | 0.0608740241615722 | 5.49333251588471 | 3.94419154651866e-08 | 2.19771076676891e-06 |
| ncstn.L | 1186.46256759195 | -0.447303987023659 | 0.0814833427882267 | -5.48951444206446 | 4.03040163854815e-08 | 2.24060817864734e-06 |
| rpl27.S | 8967.28553167695 | 0.322052124087103 | 0.0586937333371988 | 5.48699334283091 | 4.08832534389943e-08 | 2.26762045444504e-06 |
| larp1.S | 1662.61644901369 | -0.461263356839885 | 0.0841405697983255 | -5.48205649124407 | 4.20409933489119e-08 | 2.3230420992479e-06 |
| tnfrsf21.L | 1079.08776821014 | -0.522164342436476 | 0.0952521094372065 | -5.48191893619642 | 4.20737022997068e-08 | 2.3230420992479e-06 |
| MGC80700 | 9410.64811497972 | 0.410862219465422 | 0.0749666790421197 | 5.48059784313749 | 4.23891015501923e-08 | 2.33514928131604e-06 |
| rps19.S | 16371.044961116 | 0.37089545665126 | 0.067722614050669 | 5.47668547427543 | 4.33366507059022e-08 | 2.38194704128776e-06 |
| LOC108701091 | 1197.33086816294 | -0.549295524688077 | 0.100314675592737 | -5.47572447842166 | 4.3572520502577e-08 | 2.38950522142123e-06 |
| grid2ip.L | 334.73183718955 | -0.729200012005892 | 0.133195500198799 | -5.47465951115115 | 4.38353633277411e-08 | 2.39850521775708e-06 |
| rpl36.S | 6261.61837742586 | 0.391911560591876 | 0.0716299739849051 | 5.47133467721858 | 4.46658890925037e-08 | 2.43298903500736e-06 |
| iqsec3.S | 1028.02091627964 | -0.502551446395039 | 0.0918461420237258 | -5.47166636857996 | 4.45823541302326e-08 | 2.43298903500736e-06 |
| mast1.L | 2232.21930710052 | -0.478040688444615 | 0.0873923119469972 | -5.47005426214772 | 4.4989781624565e-08 | 2.44514933956864e-06 |
| ythdf1.S | 2089.50121756454 | -0.391746291941879 | 0.0716245211641516 | -5.46944378230552 | 4.51450084979161e-08 | 2.44810900992941e-06 |
| nif.L | 7383.35629123409 | -0.492312261624883 | 0.0900272564914824 | -5.46848011159222 | 4.53910985631855e-08 | 2.45597182292657e-06 |
| rpl12.S | 13533.8238956574 | 0.335652655576516 | 0.0614233943759013 | 5.4645735389089 | 4.64020988704527e-08 | 2.50125527490006e-06 |
| h3-3b.S | 3386.57570527771 | 0.424739799393297 | 0.0777277961520564 | 5.4644518488906 | 4.64339396138934e-08 | 2.50125527490006e-06 |
| rpl11.S | 13262.7218190177 | 0.365349588989885 | 0.0668794813726873 | 5.46280535511282 | 4.6866839736072e-08 | 2.51898894811534e-06 |
| LOC108703524 | 69.7041438330022 | 1.06205482325313 | 0.194556076948911 | 5.45886224634356 | 4.79195356562742e-08 | 2.56988344201661e-06 |
| LOC108713567 | 7346.95307966258 | -0.647537725082379 | 0.118649014043442 | -5.45759044272624 | 4.82639326607707e-08 | 2.57697578035332e-06 |
| wasf1.S | 1850.19204315727 | -0.506671817039582 | 0.0928360875754114 | -5.45770325174473 | 4.82332879232101e-08 | 2.57697578035332e-06 |
| tnfrsf21.S | 479.234682550459 | -0.665513902567021 | 0.122079514640264 | -5.45147893590596 | 4.99526393619677e-08 | 2.66129258916588e-06 |
| adgrl1.L | 4700.44892889827 | -0.517556814406115 | 0.0949850854021078 | -5.44882190940926 | 5.07045524570143e-08 | 2.68369585488171e-06 |
| pax2.L | 610.319784659983 | -0.86017498611307 | 0.157846327470989 | -5.44944567222298 | 5.05270542823545e-08 | 2.68369585488171e-06 |
| syn1.S | 2049.26293600662 | -0.555184819894107 | 0.101887691930745 | -5.44898809045036 | 5.06572049488544e-08 | 2.68369585488171e-06 |
| dlx5.S | 58.8057980113242 | 0.991113602554507 | 0.181950529815405 | 5.44715975029052 | 5.11804931138684e-08 | 2.70299760806156e-06 |
| rpl13a.L | 20048.8360922857 | 0.352940825143078 | 0.0647990265784735 | 5.44669949193846 | 5.13130466057708e-08 | 2.70411964043513e-06 |
| LOC108695960 | 3221.95050362285 | -0.43917052646045 | 0.0806659013780866 | -5.44431437519093 | 5.20053030480182e-08 | 2.73466846807046e-06 |
| evl.L | 1923.01352282759 | -0.422897285689445 | 0.0776846313800536 | -5.44377025644262 | 5.21644914948714e-08 | 2.73711480858835e-06 |
| rps11.L | 9316.3148901869 | 0.362886024987019 | 0.0666812303421711 | 5.44210151979634 | 5.26556510848522e-08 | 2.75693186951595e-06 |
| pccb.L | 1421.56639153872 | -0.450526127107433 | 0.082889478404176 | -5.43526314534916 | 5.4715622656755e-08 | 2.85862653080259e-06 |
| pcna.S | 4334.23561096636 | 0.406380985999937 | 0.0748217055189879 | 5.4313248165241 | 5.59372115377235e-08 | 2.9161772899087e-06 |
| rpl26.S | 9045.8526451899 | 0.354376116760577 | 0.0653003974211477 | 5.42686003080605 | 5.73540696253261e-08 | 2.98363975905283e-06 |
| igf2r.L | 2446.86682591003 | -0.38184464401359 | 0.070375772342341 | -5.42579685173638 | 5.76965516555356e-08 | 2.99504279042646e-06 |
| syn2.S | 4320.68832680675 | -0.593425260767591 | 0.109416473211572 | -5.42354586425136 | 5.8428214502928e-08 | 3.02655659517157e-06 |
| rere.S | 2115.58300605854 | -0.446065646243208 | 0.0822624063504862 | -5.4224726218524 | 5.87802195693129e-08 | 3.03831203024869e-06 |
| LOC121399515 | 1029.42485316246 | -0.684178722673453 | 0.126225399732692 | -5.42029357104308 | 5.95012433614592e-08 | 3.06551514340517e-06 |
| robo2.S | 2122.82401499696 | -0.548839556129772 | 0.101259659010505 | -5.42012052472777 | 5.95588683496847e-08 | 3.06551514340517e-06 |
| LOC108695916 | 1564.58870054505 | -0.485714241090491 | 0.0896635397817395 | -5.41707635314002 | 6.05814745251295e-08 | 3.11155674865433e-06 |
| chrnb2.S | 1251.12021314315 | -0.547334276381797 | 0.101069575948705 | -5.41542072620922 | 6.11447581897694e-08 | 3.1338623532959e-06 |
| rpl26.L | 9855.95864376054 | 0.347046130123562 | 0.0641186174574244 | 5.41256414884498 | 6.21285863380642e-08 | 3.17758289788828e-06 |
| arhgap35.L | 3174.22398000319 | -0.362985782897953 | 0.0671210007871681 | -5.40793162558665 | 6.37567537310926e-08 | 3.25400540996463e-06 |
| kiaa1549l.L | 1123.68608565971 | -0.552601543705447 | 0.102321834170101 | -5.40062195119369 | 6.64102468427094e-08 | 3.38232816938529e-06 |
| srrm2.S | 2197.99209894662 | -0.382707653766894 | 0.0709594746077899 | -5.39332704874453 | 6.91648971504809e-08 | 3.51525525392005e-06 |
| slc6a4.S | 200.942882189406 | -0.867514784651919 | 0.160943961526433 | -5.39016671656512 | 7.03923495554188e-08 | 3.57017064738903e-06 |
| znf385a.L | 421.25321559891 | -0.666328643826493 | 0.123734329361359 | -5.38515581945343 | 7.23819155485711e-08 | 3.66342970070206e-06 |
| rpl29.L | 4413.73930920797 | 0.352123059117524 | 0.0654215728946739 | 5.38236920234901 | 7.35117907955619e-08 | 3.71288034425651e-06 |
| LOC108704606 | 2911.59458884729 | -0.471072012500493 | 0.0876091599875273 | -5.37697214044237 | 7.57488889607187e-08 | 3.81793259006577e-06 |
| LOC108711293 | 2686.04851400128 | -0.379389486913601 | 0.0706043415634411 | -5.37345832441059 | 7.72406641200315e-08 | 3.88506147853426e-06 |
| ubap2l.L | 3340.09998575746 | -0.371657671101507 | 0.0691760807277156 | -5.37263266712653 | 7.75952997345085e-08 | 3.89483514824411e-06 |
| h1-0.S | 5832.63050985148 | 0.507891958966449 | 0.0945694389345885 | 5.37057176914993 | 7.84873888363037e-08 | 3.92853295162775e-06 |
| rpl15.L | 11347.0809681432 | 0.365845622541103 | 0.0681234136370672 | 5.37033602705487 | 7.85900639866258e-08 | 3.92853295162775e-06 |
| XB5957215.S | 181.850164374887 | 0.897036425251217 | 0.16706413927002 | 5.36941354482644 | 7.89930983633974e-08 | 3.94057129839297e-06 |
| apcdd1.S | 441.123123896056 | 0.840608183155295 | 0.156572966100986 | 5.36879516361158 | 7.92643826767327e-08 | 3.94600187038636e-06 |
| rps6.L | 19736.7943447492 | 0.379604961260228 | 0.0707114869918919 | 5.36836343582698 | 7.94543197519177e-08 | 3.94736859724558e-06 |
| hspa12a.L | 439.356957878487 | -0.648721948036439 | 0.12087757113483 | -5.36676855719444 | 8.01598098312279e-08 | 3.97429065314255e-06 |
| ubap2.L | 1986.30517825412 | -0.43654637009105 | 0.0813781582404133 | -5.36441693361225 | 8.12111217487564e-08 | 4.01821383251382e-06 |
| uba2.L | 3362.12142491265 | 0.369754307133808 | 0.0689431550703993 | 5.36317647134817 | 8.177104674467e-08 | 4.03769473498986e-06 |
| LOC108697808 | 856.144612622466 | -0.524268773553229 | 0.0977887621283028 | -5.36123744838254 | 8.26537872775243e-08 | 4.07300427610583e-06 |
| pygm.L | 7990.20282252568 | -0.538302920431119 | 0.100430673149738 | -5.35994535881019 | 8.3247126954992e-08 | 4.09393866851128e-06 |
| rps10.S | 12618.0482759542 | 0.398746648215699 | 0.0744113636511385 | 5.35867948993833 | 8.38324244437971e-08 | 4.11439377664163e-06 |
| naca.L | 9410.46633497473 | 0.346061007757533 | 0.0646023842731218 | 5.35678383470304 | 8.4716373028858e-08 | 4.14939428702233e-06 |
| abca2.L | 2967.35877580043 | -0.422750556972554 | 0.07893632354635 | -5.35558964466242 | 8.52778529980011e-08 | 4.16849126908137e-06 |
| eif2s2.S | 3609.62365218269 | 0.356327847382011 | 0.0665488685672855 | 5.35437874532366 | 8.58508678577069e-08 | 4.18807426452838e-06 |
| hid1.S | 999.474170872167 | -0.444939299750098 | 0.0831368005795415 | -5.35189346533008 | 8.70386390085829e-08 | 4.23750840896696e-06 |
| vac14.L | 944.992104120365 | -0.508614169819519 | 0.0950429182115874 | -5.35141575395683 | 8.72687640147033e-08 | 4.24021470594641e-06 |
| mrc2.L | 424.549091399259 | -0.588954099915259 | 0.110084976619918 | -5.34999523094506 | 8.79565497071505e-08 | 4.26510263190721e-06 |
| LOC108708151 | 1391.81629210279 | -0.649676273944827 | 0.121448491207072 | -5.34939765399899 | 8.824744886151e-08 | 4.2668829629868e-06 |
| prrc2a.S | 5690.41616043014 | -0.405971244329987 | 0.0758938434756234 | -5.34919864034008 | 8.83445348803144e-08 | 4.2668829629868e-06 |
| per2.L | 622.188113783776 | -0.538907041469714 | 0.100795327384091 | -5.34655069562387 | 8.96461804627983e-08 | 4.32115934159369e-06 |
| ppia.L | 24058.7682223214 | 0.359131343004562 | 0.0672208501405606 | 5.34255877831966 | 9.16436406793454e-08 | 4.40869427062181e-06 |
| LOC108704574 | 4036.94243010751 | -0.556521078256666 | 0.104183157364428 | -5.34175669403051 | 9.2050150873064e-08 | 4.41949874567237e-06 |
| rpl6.S | 17116.8173435853 | 0.342508188306604 | 0.0641717572131524 | 5.33736651731278 | 9.43062825714239e-08 | 4.51888920865912e-06 |
| LOC443585 | 1770.57019075079 | -0.475587472447926 | 0.0891286562911771 | -5.33596591980715 | 9.50372526316021e-08 | 4.54495081777981e-06 |
| aplp1.S | 3791.37703517572 | -0.519177756214199 | 0.0973285392093336 | -5.33428078168886 | 9.59239951705637e-08 | 4.57834486969288e-06 |
| LOC108696012 | 1274.07826655781 | -0.406563627040557 | 0.076243452808106 | -5.33243986291938 | 9.6901865402624e-08 | 4.61594885900264e-06 |
| rps12.S | 10745.8369727216 | 0.360523428621647 | 0.0677098764816213 | 5.3245323630077 | 1.01213081877292e-07 | 4.81187986521904e-06 |
| acbd3.L | 2262.33423158863 | -0.407911327224686 | 0.0766184500036856 | -5.32393081829591 | 1.01548540785171e-07 | 4.8183989254589e-06 |
| rps16.S | 6143.56004401508 | 0.404071347408408 | 0.0759204978670627 | 5.32229580627803 | 1.02465772397098e-07 | 4.85244342030234e-06 |
| slc12a5.L | 4059.89472870863 | -0.609011576321206 | 0.114439411911853 | -5.32169438960679 | 1.02805176586657e-07 | 4.85904466925341e-06 |
| rps25.L | 8910.45530286037 | 0.305707423801466 | 0.0574668492198124 | 5.31971785389045 | 1.03928298811779e-07 | 4.90259046860844e-06 |
| amigo2.L | 879.686029648747 | 0.578396136371078 | 0.10876678835222 | 5.31776422870975 | 1.05050068061849e-07 | 4.94590378584219e-06 |
| rpl31.S | 7351.397234504 | 0.330763132786156 | 0.0622328742892729 | 5.31492617950911 | 1.06700574388399e-07 | 5.0138950758061e-06 |
| lhx9.S | 752.542952329793 | 0.697624740765779 | 0.131276657969473 | 5.31415677056619 | 1.07152341562636e-07 | 5.02540344772719e-06 |
| XB5969698.L | 3554.25287622562 | -0.346931977141238 | 0.0653402980092937 | -5.30961730679423 | 1.09855658490057e-07 | 5.13237186030278e-06 |
| LOC108696525 | 4866.38543365197 | -0.5032586454086 | 0.094779102310694 | -5.30980599245259 | 1.09741990390825e-07 | 5.13237186030278e-06 |
| smn1.S | 400.682584771201 | 0.616624221114777 | 0.116169329811042 | 5.30797777793639 | 1.10848152299036e-07 | 5.16442819898047e-06 |
| sp8.L | 661.413333426378 | -0.629878295988463 | 0.118670706876805 | -5.30778245588741 | 1.1096696797019e-07 | 5.16442819898047e-06 |
| LOC108712711 | 11561.6788760659 | -0.430322629011802 | 0.0810860733430343 | -5.30698566683978 | 1.11452938069193e-07 | 5.17712749034986e-06 |
| fzd3.L | 2064.61268906054 | -0.388402502345233 | 0.0732000481519122 | -5.30604162362271 | 1.12031386052576e-07 | 5.19406582580032e-06 |
| XB5848002.S | 132.994203150466 | 0.915562393425183 | 0.172873985899494 | 5.29612589575782 | 1.18285302418599e-07 | 5.46316185733353e-06 |
| atp1b2.S | 77.9412738901165 | 1.02710605072707 | 0.193933698400417 | 5.29617111001716 | 1.18256033175692e-07 | 5.46316185733353e-06 |
| LOC108696253 | 2874.74970231935 | -0.383078608671533 | 0.0723424463633072 | -5.29535048825546 | 1.18788351168841e-07 | 5.47598520549491e-06 |
| stmn1.L | 1162.5112187811 | 0.571146842277223 | 0.107950560694143 | 5.29081867296137 | 1.21770048527163e-07 | 5.58166331871492e-06 |
| rps20.S | 10165.773705291 | 0.358111353653938 | 0.0676786179038468 | 5.29135145996507 | 1.21415781599897e-07 | 5.58166331871492e-06 |
| plec.S | 1860.12971469686 | -0.572034656383825 | 0.108117965633728 | -5.29083814175447 | 1.21757085523637e-07 | 5.58166331871492e-06 |
| grin1.L | 5278.81702408999 | -0.445911841115637 | 0.0843381442631925 | -5.28719057090097 | 1.24209220721313e-07 | 5.68274728475251e-06 |
| LOC108717002 | 1735.40244895728 | 0.424148478639417 | 0.0802344294261145 | 5.28636498910985 | 1.24770825668778e-07 | 5.69771135112272e-06 |
| LOC108700787 | 6586.69820703563 | 0.376248922038939 | 0.0711881887273491 | 5.28527173910792 | 1.25518294681461e-07 | 5.72109090242293e-06 |
| LOC108712865 | 5169.12355058539 | 0.356454623440282 | 0.0674520052537717 | 5.28456673896067 | 1.2600260802108e-07 | 5.73241078513882e-06 |
| gorasp2.S | 2643.60034569543 | -0.353468801566142 | 0.066941886407869 | -5.28023365539026 | 1.29019248416948e-07 | 5.8586796654978e-06 |
| mmp16.L | 418.137249299582 | -0.702657805453126 | 0.133093179469009 | -5.27944263001654 | 1.29577441285237e-07 | 5.87170927185153e-06 |
| psmc6.L | 4077.98940417025 | 0.348852608651673 | 0.0660812970317365 | 5.27914287887133 | 1.29789572692199e-07 | 5.87170927185153e-06 |

| | | | | | | |
|---|---|---|---|---|---|---|
| LOC108716062 | 1308.39294175412 | -0.506768966355819 | 0.0960635056143994 | -5.27535366437697 | 1.32500306424171e-07 | 5.98320156927287e-06 |
| hes5.4.L | 198.063894210673 | 0.792459174132164 | 0.150341123637905 | 5.2710739081663 | 1.35627837481137e-07 | 6.11306620364889e-06 |
| ddi2.L | 1496.98684516199 | -0.396947824069832 | 0.0753352318766637 | -5.26908611258676 | 1.37104638435764e-07 | 6.16818534473788e-06 |
| LOC108713861 | 982.511289175085 | -0.587384716589641 | 0.111619688013658 | -5.26237554541245 | 1.42205922305768e-07 | 6.38586076986381e-06 |
| LOC108704021 | 410.06130630106 | -0.660490718687307 | 0.125572246285125 | -5.25984632932019 | 1.44175839367947e-07 | 6.46237610362416e-06 |
| cdk5.L | 1350.99360107954 | -0.404145798124163 | 0.0768442404902402 | -5.25928547859735 | 1.44616228266882e-07 | 6.4701779917415e-06 |
| rps3a.S | 24160.3661554951 | 0.357681461780427 | 0.0680810956176311 | 5.2537559587657 | 1.49028320832391e-07 | 6.63094144011375e-06 |
| LOC108712537 | 1140.97061375925 | -0.433351275880047 | 0.0824789200763529 | -5.25408523146135 | 1.48761983203563e-07 | 6.63094144011375e-06 |
| rps27.S | 8010.10019915717 | 0.366294534435175 | 0.0697121552653569 | 5.25438545173201 | 1.48519546360965e-07 | 6.63094144011375e-06 |
| rpl29.S | 5692.16203186022 | 0.33784374056227 | 0.0643266290123224 | 5.25200443035112 | 1.50452842694315e-07 | 6.67180941821376e-06 |
| rps24.L | 4353.97543525385 | 0.356012445642818 | 0.0677866967470693 | 5.2519515292388 | 1.50496071506592e-07 | 6.67180941821376e-06 |
| rpl27.L | 7844.14580482644 | 0.337994904303471 | 0.0643754446469328 | 5.25037001541822 | 1.51793987309041e-07 | 6.71709130726017e-06 |
| aacs.S | 602.285860353545 | 0.757147120715194 | 0.144236203976788 | 5.24935556981963 | 1.5263221368933e-07 | 6.73824408718412e-06 |
| uhrf1bp1l.L | 1785.99992523867 | -0.450867085455998 | 0.0858938269898854 | -5.2491209351877 | 1.52826726436093e-07 | 6.73824408718412e-06 |
| rpl4.L | 32947.6450691034 | 0.279261826265297 | 0.0532394661254979 | 5.24539118418302 | 1.55951081816373e-07 | 6.86354272037495e-06 |
| hsp90ab1.S | 36981.6585506926 | -0.304369175840821 | 0.0580499034261482 | -5.24323311283443 | 1.57786980892802e-07 | 6.91927240759881e-06 |
| sptbn1.S | 8048.02830327646 | -0.408708209951096 | 0.0779457952966808 | -5.24349271689964 | 1.57565031455629e-07 | 6.91927240759881e-06 |
| znf326.S | 6806.5994573521 | 0.339439843252155 | 0.0647492788695539 | 5.24237256658877 | 1.58524872228041e-07 | 6.93910494758205e-06 |
| grp.L | 406.203539319134 | -0.711849940671473 | 0.135821634050036 | -5.24106447143191 | 1.5965291782709e-07 | 6.97591364332971e-06 |
| LOC108698154 | 91.7226538578926 | 0.94427005927159 | 0.180217173164654 | 5.23962307866459 | 1.60904904339335e-07 | 7.0179959533569e-06 |
| LOC108708001 | 1342.23420742354 | -0.434887111552686 | 0.0830278560172392 | -5.23784585576182 | 1.62461660029808e-07 | 7.07319635979238e-06 |
| pcna.L | 5745.84192776488 | 0.370201060220093 | 0.0706971078045057 | 5.23643854348026 | 1.63704717291347e-07 | 7.1145660140894e-06 |
| rps28p9.S | 4648.19961676381 | 0.367365114037935 | 0.0701609321491352 | 5.2360352518843 | 1.64062630607523e-07 | 7.11738847853421e-06 |
| ltbp4.L | 2888.84462561031 | -0.587841943967893 | 0.11229326550104 | -5.23488155184559 | 1.6509069986277e-07 | 7.14922185822841e-06 |
| tbx20.L | 463.125082389948 | -0.848541929123821 | 0.162166664552005 | -5.2325299497771 | 1.67205550460312e-07 | 7.22792107274524e-06 |
| npm1.L | 10414.0678103342 | 0.309874746912529 | 0.0592887358985673 | 5.22653657926981 | 1.72714662744318e-07 | 7.45280642399725e-06 |
| pc.1.L | 702.094111654957 | -0.548904345417597 | 0.105057051739789 | -5.22482152628036 | 1.74323168467897e-07 | 7.50887775666506e-06 |
| rpl18a.S | 9612.75181949482 | 0.350552180368871 | 0.0671094295460534 | 5.22359052580393 | 1.75486611567884e-07 | 7.54561370164634e-06 |
| LOC121398252 | 37.0883652788104 | 0.890349092437396 | 0.170765105298617 | 5.21388190450528 | 1.8492922398824e-07 | 7.93758050807475e-06 |
| LOC108706905 | 5963.96432485054 | 0.374083603982248 | 0.0717605096830574 | 5.21294519275925 | 1.85865823099493e-07 | 7.96371129872855e-06 |
| sec23ip.L | 1250.03809643587 | -0.439884102953337 | 0.0843995863699415 | -5.21192249716995 | 1.86893631663305e-07 | 7.99365121061324e-06 |
| LOC108699273 | 3480.66264153006 | -0.597748392968591 | 0.114809074300017 | -5.2064559932481 | 1.92481314746516e-07 | 8.21817409569749e-06 |
| XB5843130.S | 6014.53419578814 | 0.341782340715943 | 0.0656501304015568 | 5.2061182304649 | 1.92831811136956e-07 | 8.2186947715109e-06 |
| vamp2.S | 3835.15205835979 | -0.49449009939973 | 0.0950746257789783 | -5.20107331844019 | 1.98140912482876e-07 | 8.43018446209983e-06 |
| myg1.S | 1104.31383654522 | 0.464638120861588 | 0.0893513851823833 | 5.20012219075477 | 1.99157554161715e-07 | 8.45862521119705e-06 |
| eef1a1.S | 99429.9721336802 | 0.296071594241415 | 0.0569466116533423 | 5.1991081759829 | 2.00246966886226e-07 | 8.49005203060032e-06 |
| rpl23a.L | 13223.2929926161 | 0.349354684609218 | 0.0672077866457648 | 5.19812810456887 | 2.0130538465733e-07 | 8.5200575171867e-06 |
| arid1b.L | 1435.46076788566 | -0.396488925162134 | 0.0763108936051623 | -5.19570544165809 | 2.03944967535218e-07 | 8.61266701644315e-06 |
| stxbp1.L | 16760.3794926492 | -0.424966520016082 | 0.0817955767176325 | -5.19547067298167 | 2.04202527433574e-07 | 8.61266701644315e-06 |
| lman1l.L | 56.5610962121983 | 0.905251675310142 | 0.17430484591441 | 5.19349689081309 | 2.06380390448032e-07 | 8.68738819115622e-06 |
| qars1.S | 2650.69831372117 | 0.367898148075236 | 0.0708420308546748 | 5.19321854041624 | 2.06689321416329e-07 | 8.68738819115622e-06 |
| hspa1l.S | 94312.2445885215 | 0.250392066446415 | 0.0482201039715678 | 5.19269030597807 | 2.07276818853274e-07 | 8.69703460660005e-06 |
| c11orf87.S | 880.99605615702 | -0.528287412615442 | 0.101840135046567 | -5.18741861814971 | 2.13229056125731e-07 | 8.9159839750749e-06 |
| nell1.S | 1420.53838620552 | -0.469817052763783 | 0.0905678167461295 | -5.18746139239204 | 2.13180101766994e-07 | 8.9159839750749e-06 |
| eif4g1.S | 7351.54951492036 | -0.345590980041374 | 0.0666527928723255 | -5.18494372326391 | 2.16080102772819e-07 | 9.01967356832109e-06 |
| cttn.S | 2412.63403776349 | -0.357409073270133 | 0.0689818617201852 | -5.18120364335644 | 2.20458640662625e-07 | 9.18665903303227e-06 |
| stac2.S | 424.550289017476 | -0.620904401896098 | 0.119848184708015 | -5.18075766778445 | 2.20986433219592e-07 | 9.19288426136431e-06 |
| uba52.L | 8419.95728908677 | 0.367274123125 | 0.0709063667287364 | 5.17970585815044 | 2.22236043814678e-07 | 9.22906401441672e-06 |
| sdhb.S | 1728.08907692962 | -0.440654533415066 | 0.0850954439172083 | -5.17835636234286 | 2.23849325778769e-07 | 9.28019713390685e-06 |
| rlbp1.S | 208.255401044617 | 0.950570788156867 | 0.183632800033323 | 5.17647603251908 | 2.26116084570014e-07 | 9.35604883780733e-06 |
| rab6b.L | 1064.90748840862 | -0.506646852639271 | 0.0978850690374414 | -5.17593599944724 | 2.26771189578198e-07 | 9.35604883780733e-06 |
| LOC100335021 | 65.0183938016937 | 0.994749777583991 | 0.192189347556173 | 5.17588404473483 | 2.26834311577695e-07 | 9.35604883780733e-06 |
| hes5.3.L | 435.550452692112 | 0.602123949399431 | 0.116352434313819 | 5.17500087514642 | 2.27909912688116e-07 | 9.38448037092339e-06 |
| relt.L | 259.038514968689 | -0.698237194206771 | 0.134997562110979 | -5.17222076671848 | 2.31328040472376e-07 | 9.47703779972326e-06 |
| kansl3.L | 1557.27305464202 | -0.376457488056202 | 0.0727819268100548 | -5.17240343249877 | 2.31101941878755e-07 | 9.47703779972326e-06 |
| LOC108717266 | 1269.62680103178 | -0.534332061151681 | 0.103295814213148 | -5.17283362565981 | 2.3057030404187e-07 | 9.47703779972326e-06 |
| hnrnpk.L | 14545.4713536128 | 0.342340167640537 | 0.0662072405391865 | 5.17073608343356 | 2.33173684886844e-07 | 9.53656818289729e-06 |
| LOC108709952 | 6767.34554822591 | 0.336676938758105 | 0.0651288162491531 | 5.16940055336079 | 2.34846062242456e-07 | 9.55670056301211e-06 |
| atp2b1.S | 3837.26092801456 | -0.401287422418222 | 0.0776241859150318 | -5.16961843384065 | 2.34572439115945e-07 | 9.55670056301211e-06 |
| LOC108713446 | 2770.94450038902 | -0.415784990159917 | 0.0804300367414191 | -5.16952381231228 | 2.34691230804095e-07 | 9.55670056301211e-06 |
| gpc5.L | 1158.3213522706 | -0.508588135609665 | 0.0984210295226765 | -5.16747424891024 | 2.37278646009135e-07 | 9.63954419087949e-06 |
| nap1l1.L | 8914.2215603075 | 0.306960161858471 | 0.0594066094078552 | 5.16710455146566 | 2.37748285064708e-07 | 9.64249889375963e-06 |
| LOC108707429 | 587.411995027337 | -0.552163357082141 | 0.107045702781012 | -5.15820198977747 | 2.49332560203304e-07 | 1.00954753626318e-05 |
| tfap2e.L | 185.864498601023 | 0.849915722773631 | 0.164851684756026 | 5.15563868231905 | 2.52767883145944e-07 | 1.0217542351327e-05 |
| rpl30.L | 7888.74193965717 | 0.330154095841154 | 0.0640833910214277 | 5.15194484216292 | 2.57798867103864e-07 | 1.04035974707994e-05 |
| LOC108697493 | 480.919655931801 | -0.84011968560321 | 0.163158047004462 | -5.14911584760564 | 2.61717206274744e-07 | 1.054420863887e-05 |
| dek.L | 2714.2955744377 | 0.392706519340488 | 0.076318176140437 | 5.14564864099803 | 2.66598005966073e-07 | 1.07230661538738e-05 |
| tacc2.S | 1495.25636773113 | -0.403821725820357 | 0.0785293893378384 | -5.14230059886357 | 2.71394423964788e-07 | 1.08979440261166e-05 |
| zmiz2.L | 4487.42833588267 | -0.331640820007474 | 0.0645240484064819 | -5.13980179790081 | 2.75028420843982e-07 | 1.10256443168048e-05 |
| palld.L | 733.760534819524 | -0.5414865210508 | 0.105365538677233 | -5.13912354882499 | 2.76022876985166e-07 | 1.10472813401608e-05 |
| sec24a.L | 1174.32195821933 | -0.429856273504995 | 0.0836627144940547 | -5.13796708730437 | 2.77726507477996e-07 | 1.1097183836629e-05 |
| LOC108696915 | 4256.13956676654 | -0.327629209222434 | 0.0638102745070195 | -5.13442720241538 | 2.83004607089152e-07 | 1.12895138335367e-05 |
| dlx6.S | 39.9451608055836 | 0.868505005309092 | 0.169168382929898 | 5.13396764967005 | 2.83696880892842e-07 | 1.12985770891979e-05 |
| sntb1.L | 307.943459752267 | -0.708953662713208 | 0.138151406058377 | -5.13171514456853 | 2.87113792605 05e-07 | 1.14159451351016e-05 |
| rpl13a.S | 12012.6230148591 | 0.296721427013072 | 0.0578426339434635 | 5.1298048996713 | 2.90042629281244e-07 | 1.15135549604373e-05 |
| srrm1.S | 2814.78858833245 | -0.335385665361089 | 0.0654275881267439 | -5.12605882263904 | 2.95870202041503e-07 | 1.17066297856617e-05 |
| sgo1.L | 853.44349607093 | 0.454944333413256 | 0.0887465438356266 | 5.12633296746619 | 2.9543992312551e-07 | 1.17066297856617e-05 |
| maf.S | 338.305308238753 | -0.630274753241965 | 0.122968969156483 | -5.1254780581263 | 2.96783729897708e-07 | 1.17236811937153e-05 |
| ptmap12.L | 9630.65574306162 | 0.353229575922134 | 0.0689217021961266 | 5.12508491036638 | 2.97403686663069e-07 | 1.17290992918711e-05 |
| hipk2.S | 2013.92212709853 | -0.446774545628838 | 0.0871868668326222 | -5.12433307744086 | 2.98592740042366e-07 | 1.17569076605985e-05 |
| klc4.L | 2659.42841673617 | -0.371617413061357 | 0.0725702010670313 | -5.12079900010341 | 3.04243800320123e-07 | 1.19600305582153e-05 |
| myh10.S | 7374.79930118695 | -0.316832029549048 | 0.0618853306077417 | -5.11966287386066 | 3.06082323410121e-07 | 1.20128658560993e-05 |
| dpysl5.L | 1488.10291879413 | -0.483329582810507 | 0.0944468881263894 | -5.11747493642895 | 3.09653194973413e-07 | 1.21334108365873e-05 |
| nsa2.L | 4993.70676846824 | 0.35375957227581 | 0.0691395952593906 | 5.11659883093923 | 3.11094311246619e-07 | 1.21557617838325e-05 |
| rpl32.L | 8697.76784653158 | 0.372771241253714 | 0.0728564029020261 | 5.11651998184703 | 3.11224328210414e-07 | 1.21557617838325e-05 |
| col5a3.S | 970.917839807593 | -0.632387277285708 | 0.123768931740151 | -5.10941856243365 | 3.23151740776007e-07 | 1.26013617823633e-05 |
| fjx1.S | 959.498885449547 | -0.544716929469721 | 0.106653307855692 | -5.10736085379323 | 3.26689501563318e-07 | 1.2718901844518e-05 |
| r3hdm1.L | 2238.84355095468 | -0.460970719684153 | 0.0902623912309882 | -5.10700761853843 | 3.27300557316337e-07 | 1.27223035831089e-05 |
| gars1.S | 2964.75303448859 | -0.327915603283639 | 0.0642276436589499 | -5.10552130831512 | 3.29883811244056e-07 | 1.28022321251807e-05 |
| rps9 | 12560.9488372551 | 0.39543891019836 | 0.0774933997850254 | 5.10287213227639 | 3.34537048281247e-07 | 1.29621101290982e-05 |
| thoc1.L | 1928.76820043487 | 0.416752327886341 | 0.0816921591225504 | 5.1014972839823 | 3.36976858293289e-07 | 1.30358531773522e-05 |
| emc1.S | 2040.6273514698 | -0.346507919846744 | 0.0679944711902672 | -5.09611904881383 | 3.46687159081951e-07 | 1.33901714510921e-05 |
| rab11fip5.L | 567.808703009492 | -0.571223487149814 | 0.112187028984758 | -5.09170705668142 | 3.54854069401104e-07 | 1.36621628558327e-05 |
| eif3m.S | 3082.99286218013 | 0.388363227411458 | 0.0762734478948448 | 5.09172245559003 | 3.54825244597155e-07 | 1.36621628558327e-05 |
| vcp.S | 7849.5018754614 | -0.416157708989409 | 0.0817666011816238 | -5.08958062308374 | 3.58856263583022e-07 | 1.37943893472879e-05 |
| cyc1.S | 2593.39509032483 | -0.427740629528153 | 0.0840570812761271 | -5.08869238658225 | 3.60540896354869e-07 | 1.3837252031667e-05 |
| git1.L | 1056.81240581029 | -0.51823229076567 | 0.101865898419608 | -5.08739724290219 | 3.63010956659647e-07 | 1.38663336180652e-05 |
| selenoi.L | 1120.97647897205 | -0.491160415200868 | 0.0965399453610964 | -5.08763924988501 | 3.62548170590931e-07 | 1.38663336180652e-05 |
| dpysl5.S | 3141.44619964584 | -0.432816453350827 | 0.0850761417433245 | -5.08740105606386 | 3.63003660392766e-07 | 1.38663336180652e-05 |
| LOC108702869 | 13441.2340091501 | 0.35898244919539 | 0.0706238948854358 | 5.08301687095737 | 3.71486691138383e-07 | 1.41678142457078e-05 |
| ubr4.S | 3213.73540774377 | -0.378576391785221 | 0.0745285283415448 | -5.07961716418543 | 3.78196221956745e-07 | 1.44010956366704e-05 |
| plxna2.L | 2821.66833052054 | -0.571825239964963 | 0.112579332813839 | -5.07930919177263 | 3.78809768684161e-07 | 1.44018850084711e-05 |
| rps14.S | 9394.68180370658 | 0.363419923735711 | 0.0715603568000276 | 5.07850910737174 | 3.80408200024592e-07 | 1.44400575178085e-05 |
| epg5.L | 1963.97852701715 | -0.52307614717592 | 0.10301421309431 | -5.07770855766226 | 3.82014156894154e-07 | 1.44783961428808e-05 |
| rpl8.S | 17062.2648076491 | 0.340350935075286 | 0.0670360053853925 | 5.07713628099699 | 3.83166049379412e-07 | 1.44994330274508e-05 |
| chd4.L | 14021.166208164 | -0.256623879526713 | 0.0505735127065655 | -5.07436335331711 | 3.88795504871092e-07 | 1.46895769756428e-05 |
| stim1.L | 1586.61740184625 | -0.384053300568708 | 0.0757111002128291 | -5.07261550141403 | 3.92384823615665e-07 | 1.48021691070 17e-05 |
| LOC108709337 | 1752.76416304444 | -0.534053644048198 | 0.105290167285032 | -5.07220814458824 | 3.93225938326175e-07 | 1.48109006910017e-05 |

| | | | | | | |
|---|---|---|---|---|---|---|
| LOC108696016 | 865.024441020829 | -0.629888515047915 | 0.124233336008094 | -5.07020527088538 | 3.97386864140752e-07 | 1.49309182864449e-05 |
| LOC108696239 | 29.3631766535534 | 0.934794109300576 | 0.184374506025678 | 5.07008332903893 | 3.9764156299209e-07 | 1.49309182864449e-05 |
| galnt14.S | 427.737491914241 | -0.581371106294641 | 0.114772725595944 | -5.06541169320443 | 4.07518658854194e-07 | 1.52781763861 17e-05 |
| nap1l1.S | 6541.94112022883 | 0.311400562723312 | 0.0615009126886824 | 5.06334864166361 | 4.11955472863797e-07 | 1.54207184248892e-05 |
| LOC108703516 | 671.618102988367 | -0.504763564092463 | 0.0997251695650678 | -5.06154631066452 | 4.15869693118348e-07 | 1.55432897301802e-05 |
| mcm7.L | 4937.64854754719 | 0.334695413743186 | 0.0661434765552726 | 5.06014245355699 | 4.18943361686712e-07 | 1.56341167877373e-05 |
| cadps.L | 5343.53436584962 | -0.39018204166107 | 0.0771273165976699 | -5.05893448486523 | 4.21605676891486e-07 | 1.57069141593477e-05 |
| ado.L | 1947.84101104016 | -0.420842465075815 | 0.0831922876373737 | -5.05867162723331 | 4.22187163334735e-07 | 1.57069141593477e-05 |
| trib2.L | 877.695092162515 | -0.511059186412518 | 0.10107170120287 | -5.05640233943152 | 4.27239497506 94e-07 | 1.58705754624367e-05 |
| dct.L | 127.793589130659 | 0.930483911433607 | 0.184257169593633 | 5.04991970453973 | 4.41995811447918e-07 | 1.63936583867415e-05 |
| snrpd1.L | 3794.7808807206 | 0.370329503223854 | 0.0733479463191091 | 5.04894167878526 | 4.44264327171166e-07 | 1.64526792138663e-05 |
| LOC108710730 | 1878.70694442447 | -0.450623646176523 | 0.0892724957273835 | -5.0477321430849 | 4.47085363985424e-07 | 1.65319510390592e-05 |
| psmc3.L | 4831.37520636516 | 0.323106430077351 | 0.0640461206330577 | 5.04490243723798 | 4.53752837711871e-07 | 1.67529961084684e-05 |
| LOC108702189 | 1007.49537680848 | -0.574755292302693 | 0.113995772704045 | -5.0419000518104 | 4.60932034109616e-07 | 1.69922349536761e-05 |
| LOC108702465 | 392.480939713886 | -0.627236836896912 | 0.124433234246511 | -5.04075009136476 | 4.63710694190421e-07 | 1.70687690979729e-05 |
| mtss1.1.L | 1905.00951850862 | -0.479316590403529 | 0.0951017679881559 | -5.04003869268995 | 4.65437735336571e-07 | 1.71064210926878e-05 |
| LOC108718696 | 679.774413729353 | -0.657879908604062 | 0.130542096896359 | -5.03959967125677 | 4.66506628475464e-07 | 1.71198066951404e-05 |
| stra6.L | 88.8946693673077 | 0.974828136006502 | 0.193475459357818 | 5.03851051312731 | 4.69168654082525e-07 | 1.7145694257467e-05 |
| arpp19.S | 2364.11065239171 | 0.401459010958375 | 0.0796791497655043 | 5.03844496508645 | 4.6932932745598e-07 | 1.7145694257467e-05 |
| stmn4.L | 1995.02920078602 | 0.392758481551604 | 0.0779435594664022 | 5.03901135950692 | 4.67942714297767e-07 | 1.7145694257467e-05 |
| tgm1.L | 238.816252217446 | -0.707501765938251 | 0.140456315485325 | -5.03716592232672 | 4.72475199014912e-07 | 1.72347034307331e-05 |
| LOC108711609 | 4387.60501299124 | -0.313284037032518 | 0.0622102610895625 | -5.03589008542322 | 4.75633442611527e-07 | 1.73073445164093e-05 |
| evx1.S | 309.758385352386 | -0.639726257566337 | 0.127036023397194 | -5.03578623180098 | 4.75891419155405e-07 | 1.73073445164093e-05 |
| phr.L | 1293.25135942355 | 0.392099752161059 | 0.0779019702652018 | 5.03324563969606 | 4.82244548905116e-07 | 1.7512180973245e-05 |
| LOC108717861 | 2824.360726774 | 0.359969234149845 | 0.0715568082274872 | 5.03053787705934 | 4.89105746476585e-07 | 1.77246304790733e-05 |
| ubtf.S | 2901.37679181432 | -0.354744621679068 | 0.0705206914351736 | -5.03036221653 | 4.89554089547139e-07 | 1.77246304790733e-05 |
| nup214.L | 1399.67316960712 | -0.387810757231552 | 0.0771125817196787 | -5.02915021884924 | 4.926583231617e-07 | 1.78104781293011e-05 |
| sf3a3.S | 1543.53232666975 | 0.405340176535198 | 0.0806302379396 93 | 5.02714846058585 | 4.97826946086887e-07 | 1.79705911266491e-05 |
| cry2.L | 186.959138981368 | -0.793576142794536 | 0.157889616780368 | -5.02614522079966 | 5.00436986073813e-07 | 1.80380061419543e-05 |
| sox8.S | 390.132289126518 | -0.686393902052784 | 0.136612608697422 | -5.02438177996478 | 5.05056791755765e-07 | 1.8177555109503e-05 |
| adar-a | 1470.27156354382 | -0.385600551073933 | 0.0767937607375759 | -5.02124843698852 | 5.1336707578789e-07 | 1.84493191408151e-05 |
| LOC108698099 | 2247.74973556903 | -0.485415222911073 | 0.0966963416840861 | -5.01999573569143 | 5.16726266140562e-07 | 1.85426113879155e-05 |
| nup98l.L | 2347.32468265543 | -0.367884686850909 | 0.0733026363880934 | -5.01871017166669 | 5.20195613121604e-07 | 1.86220651232454e-05 |
| nkx6-2.L | 596.719396422661 | -0.784286484914215 | 0.156275714602717 | -5.01860757385126 | 5.20473459236175e-07 | 1.86220651232454e-05 |
| rps23.L | 10903.3386188471 | 0.375873366365875 | 0.0749055461895024 | 5.01796442969602 | 5.22218426919873e-07 | 1.86570212699873e-05 |
| prr12.L | 2251.13529764092 | -0.490869365898665 | 0.0978659973996986 | -5.01572945600181 | 5.2832629060468e-07 | 1.88475167458885e-05 |
| rnf10.S | 2560.72804229153 | -0.334394109201945 | 0.0667021475595465 | -5.01324352268304 | 5.35200909555373e-07 | 1.90647667107599e-05 |
| LOC108702825 | 1284.01831246528 | 0.441158169882519 | 0.0880109168648689 | 5.01253918942656 | 5.37164306623688e-07 | 1.91066905785006e-05 |
| cnih2.S | 1205.72701482241 | 0.432657033198619 | 0.0863282899330704 | 5.01176420306778 | 5.39332680768076e-07 | 1.91557721441223e-05 |
| btg1.S | 2444.10780897301 | 0.441742080147982 | 0.0881616892651553 | 5.01059001738723 | 5.42634083488437e-07 | 1.92168402686124e-05 |
| znfx1.L | 105.500633167793 | -0.964529331079404 | 0.19249212776727 | -5.01074689270178 | 5.42191879467312e-07 | 1.92168402686124e-05 |
| psma4.L | 2718.97427737049 | 0.379689727627953 | 0.0758276243933213 | 5.00727446845078 | 5.52061785083483e-07 | 1.94938793703752e-05 |
| dach2.L | 1576.87082312094 | -0.466085112394535 | 0.0930814630600929 | -5.00728176235942 | 5.52040872695511e-07 | 1.94938793703752e-05 |
| clstn3.L | 5840.59955572796 | -0.452332835854101 | 0.0903435720009979 | -5.00680707919214 | 5.53403432039377e-07 | 1.95128925659864e-05 |
| LOC108711102 | 2151.43926310796 | -0.413215178679364 | 0.0825637645449361 | -5.00480060419808 | 5.5919884443566e-07 | 1.96687007087839e-05 |
| rps13.L | 10347.0631874077 | 0.283414052008198 | 0.0566293859562309 | 5.00471702496033 | 5.59441516002704e-07 | 1.96687007087839e-05 |
| hmgb3.L | 14782.3203465006 | 0.315453839019896 | 0.0630525985225816 | 5.00302678099649 | 5.64370968526563e-07 | 1.98133357072028e-05 |
| LOC108705545 | 234.218648300268 | -0.722352482822484 | 0.144423784617675 | -5.0016171833104 | 5.68513938528068e-07 | 1.99299821393952e-05 |
| pmch.L | 39.6135077401135 | 0.828602025051383 | 0.165686006561956 | 5.00103806136181 | 5.70224528533895e-07 | 1.99611450953926e-05 |
| madd.S | 3250.17206982961 | -0.427203296770195 | 0.085453551860275 | -4.99924564245983 | 5.75550415148637e-07 | 2.01185924972964e-05 |
| MGC81403 | 1109.4838843856 | 0.660096701252305 | 0.132048543577371 | 4.99889422003006 | 5.76600218607385e-07 | 2.01263300443216e-05 |
| madd.L | 3077.05487785014 | -0.372598139338411 | 0.0745442574655938 | -4.99834798824557 | 5.78235641061267e-07 | 2.01544571936046e-05 |
| LOC108718198 | 1726.76672670636 | -0.51034646527116 | 0.102164635488364 | -4.99533388272389 | 5.87340622922595e-07 | 2.04424829416641e-05 |
| rps17.S | 5962.94508929743 | 0.344724974493213 | 0.0690333060688423 | 4.9936037273 | 5.92629331625895e-07 | 2.05327182290229e-05 |
| nup93.L | 2310.36388194367 | 0.335373666579554 | 0.067163851073663 | 4.99336564563172 | 5.9336067886312e-07 | 2.05327182290229e-05 |
| ctsb.S | 1409.04158833328 | -0.408174826829403 | 0.0817476781641765 | -4.99310605506927 | 5.94159089281432e-07 | 2.05327182290229e-05 |
| dnajc9.L | 1504.66222269181 | 0.390759624757035 | 0.0782430162033403 | 4.99417895319266 | 5.90865917599619e-07 | 2.05327182290229e-05 |
| LOC108698868 | 6877.37111274072 | -0.408101002817599 | 0.0817321862268059 | -4.99314923113794 | 5.9402622292915e-07 | 2.05327182290229e-05 |
| armc6.L | 1390.42262172067 | 0.377975914682208 | 0.0757253276360018 | 4.99140679191341 | 5.99411066947267e-07 | 2.06847904267286e-05 |
| pmel.S | 517.072828954625 | 0.818187397732766 | 0.163960045063046 | 4.99016328897783 | 6.03282734981269e-07 | 2.07888663313971e-05 |
| sbf2.L | 2636.8913212575 | -0.365833688159615 | 0.0733188628814698 | -4.98962577680776 | 6.04963740085903e-07 | 2.08172650164971e-05 |
| abca3.S | 3103.3976517227 | -0.422909954770024 | 0.0847773786252423 | -4.98847642647096 | 6.08573358019393e-07 | 2.09118545399196e-05 |
| ubqln4.S | 1930.98118738478 | -0.413082755209465 | 0.0828532898082599 | -4.98571337559953 | 6.17336061555935e-07 | 2.11829975698304e-05 |
| LOC108702627 | 968.584541295722 | -0.523557127487842 | 0.105026573975101 | -4.98499672675187 | 6.19628618839154e-07 | 2.12316751284604e-05 |
| h3-3b.L | 8045.95340010858 | 0.366328185709052 | 0.0734966714912558 | 4.98428266581617 | 6.21921057767187e-07 | 2.12802115174592e-05 |
| LOC108698757 | 7805.13164656805 | 0.360042725344086 | 0.0722527215437616 | 4.98310261055035 | 6.25727464788916e-07 | 2.13803418137 58e-05 |
| atp2b4.S | 3689.30998342273 | -0.524140429153388 | 0.10519345629261 | -4.982633403502 | 6.27247177780831e-07 | 2.14021670463589e-05 |
| rdh5.L | 110.550081827544 | 0.939942293421325 | 0.188680040572277 | 4.98167315721594 | 6.30368413498021e-07 | 2.14341104544477e-05 |
| bag6.S | 2938.46955335237 | -0.37372449872843 | 0.0750212193901407 | -4.98158390074829 | 6.30659296664733e-07 | 2.14341104544477e-05 |
| stau1.S | 2157.6493588342 | -0.353598215980437 | 0.0709818291054061 | -4.98153147695523 | 6.3083020395695e-07 | 2.14341104544477e-05 |
| setdb1.L | 1917.91644580409 | -0.441535651807983 | 0.0887367073956215 | -4.97579485161 03 | 6.49804391188532e-07 | 2.20479718987908e-05 |
| cck.S | 89.3112092198944 | 0.924398103813185 | 0.185837946483404 | 4.97421609152218 | 6.55121989389638e-07 | 2.21973969459301e-05 |
| plod3.L | 1528.87990718228 | -0.426015575714712 | 0.0856641967618986 | -4.97308784554195 | 6.5894782808179e-07 | 2.22763266290258e-05 |
| LOC108697213 | 2037.39508132069 | -0.423303209034155 | 0.0851204864005095 | -4.97298860632006 | 6.59285372778033e-07 | 2.22763266290258e-05 |
| opn1lw.L | 36.4455510620009 | 0.828629052483511 | 0.166756731731385 | 4.96908906693064 | 6.72681680314e-07 | 2.26974010299282e-05 |
| eif3k.S | 2793.99375865322 | 0.342383132445052 | 0.0689589875214347 | 4.96502551373203 | 6.86920388606057e-07 | 2.31456919844599e-05 |
| rab10.S | 2690.09130614045 | -0.37865615270532 | 0.0762817070600538 | -4.96391818299528 | 6.90850557594005e-07 | 2.32458773492919e-05 |
| robo4.L | 1662.96069055601 | 0.472747515991577 | 0.095270197007019 | 4.96217632421551 | 6.97076693990593e-07 | 2.34229338918499e-05 |
| plekhh2.S | 922.859543938209 | -0.536390065710701 | 0.10811654341761 | -4.9612210005535 | 7.00514344382752e-07 | 2.34735110102546e-05 |
| LOC108719848 | 361.44591799775 | -0.65001396953773 | 0.131017237220664 | -4.96128588365021 | 7.00280352017089e-07 | 2.34735110102546e-05 |
| npm1.S | 7410.50183592574 | 0.324982530316689 | 0.0655440781312478 | 4.95822871542922 | 7.11387859503716e-07 | 2.38050367200872e-05 |
| eif3e.S | 1477.57761804476 | 0.367580641895032 | 0.0741470964685973 | 4.95745159826602 | 7.14238278892603e-07 | 2.38675443568 32e-05 |
| r3hdm2.L | 1871.72244885679 | -0.405802998569017 | 0.0818695437907585 | -4.95670281986871 | 7.16995160872531e-07 | 2.39267588437325e-05 |
| evx1.L | 462.829364400545 | -0.632019906333433 | 0.127716528311865 | -4.94861483229589 | 7.47434887644996e-07 | 2.48742235074624e-05 |
| rpl38.L | 4185.42228870551 | 0.356416718113895 | 0.0720214698649468 | 4.94875651360963 | 7.46891102319815e-07 | 2.48742235074624e-05 |
| eef1a1.L | 100865.669707805 | 0.294957494184667 | 0.0596087801255875 | 4.94822228475792 | 7.4894351259633e-07 | 2.48903333721139e-05 |
| kctd15.S | 195.286111983182 | 0.761728222430818 | 0.153955873427361 | 4.94770485512017 | 7.50936562327348e-07 | 2.49224765644544e-05 |
| efr3b.L | 930.399048485149 | -0.571182368135061 | 0.115460548261547 | -4.9469916498334 | 7.53692082835549e-07 | 2.49798028109234e-05 |
| alas1.L | 887.71231466773 | -0.46790162942758 | 0.0947044564213307 | -4.94065059986123 | 7.78623261769191e-07 | 2.57709448520718e-05 |
| nfia.S | 1009.88959987695 | -0.434503541101154 | 0.0879751572160051 | -4.93893452255298 | 7.85505802351105e-07 | 2.59280950574969e-05 |
| kif1a.L | 5478.26651165776 | -0.511536795786753 | 0.103572221919514 | -4.93893812748622 | 7.85491283085248e-07 | 2.59280950574969e-05 |
| clasp1.S | 3012.23987642848 | -0.453400680383639 | 0.0918431284825866 | -4.93668593257473 | 7.94612795105634e-07 | 2.61931115933464e-05 |
| LOC108716425 | 20507.2291815905 | 0.303242612902547 | 0.0614322754127716 | 4.93621001118744 | 7.9655331814598e-07 | 2.62084535386623e-05 |
| rps9.S | 8153.21806089745 | 0.320058513395425 | 0.0648411127328602 | 4.93604288862282 | 7.97235826338 66e-07 | 2.62084535386623e-05 |
| sarnp.L | 1635.18635403777 | 0.397658676191995 | 0.0805670738932235 | 4.93574678806155 | 7.98446448263147e-07 | 2.62127811001418e-05 |
| LOC108704501 | 2570.50363582212 | -0.327138322658484 | 0.066293088044414 | -4.93472747022003 | 8.02627534494715e-07 | 2.63144849163 49e-05 |
| eif5a.L | 10859.2337415277 | 0.3430563707381 | 0.0695550487108839 | 4.9321562862254 | 8.13268037336822e-07 | 2.662740390709e-05 |
| rpl31.L | 8144.3325400629 | 0.351316507322602 | 0.0712491313888791 | 4.93081810927785 | 8.1885952932837e-07 | 2.67743922012159e-05 |
| cyp19a1.L | 30.5274561900233 | 0.915416488491534 | 0.185671951203558 | 4.93028959171079 | 8.2107809214322e-07 | 2.68108483474831e-05 |
| cacnb1.L | 1761.79226720771 | -0.438731353459569 | 0.0889992914407175 | -4.92960501547149 | 8.23960350652379e-07 | 2.68688493406025e-05 |
| ywhaq.L | 13349.2935630685 | 0.304057250105373 | 0.061692856366884 | 4.92856495891779 | 8.28357937470482e-07 | 2.697604253741e-05 |
| odc1.S | 3342.08293359133 | 0.409321883157414 | 0.0830617259565368 | 4.92792412442282 | 8.31078771383 46e-07 | 2.70284172315794e-05 |
| preb.L | 1703.02919510334 | -0.360076477261318 | 0.0743055115824355 | -4.92663961885504 | 8.36558422074949e-07 | 2.71243584163144e-05 |
| dnajc5g.L | 888.908120756477 | -0.457458240857057 | 0.0928576186249859 | -4.92644812166188 | 8.37378316137146e-07 | 2.71243584163144e-05 |
| fzd5.L | 46.6069483687495 | 0.956437842941432 | 0.194136105887066 | 4.92663556102138 | 8.36575787644485e-07 | 2.71243584163144e-05 |
| rhob.S | 1964.00415428284 | -0.344598429746752 | 0.0699533482852142 | -4.92611773694882 | 8.38794675931392e-07 | 2.71340583982387e-05 |
| LOC121393051 | 2775.15918648316 | 0.350380467245973 | 0.0711344832125355 | 4.92560642071591 | 8.4099123882989e-07 | 2.71689377076241e-05 |
| smarca4.S | 6926.4008928144 | -0.350003673443087 | 0.0710662839210698 | -4.92503131065389 | 8.43468473082767e-07 | 2.72127796614512e-05 |

| | | | | | | |
|---|---|---|---|---|---|---|
| uba1.S | 12546.521720989 | -0.324903637545942 | 0.066000436697275 | -4.92274981506837 | 8.53365221848322e-07 | 2.74955632620466e-05 |
| syf2.S | 1468.46982773714 | 0.387132033487101 | 0.0786654269780864 | 4.92124746993292 | 8.59943107617677e-07 | 2.76708051078991e-05 |
| sulf1.S | 1927.0688968126 | -0.468103574749416 | 0.0952511521141247 | -4.91441378250795 | 8.9048501333977e-07 | 2.86156652302598e-05 |
| rps19.L | 2400.08680698245 | 0.332887160785372 | 0.0677420478967983 | 4.91404040947373 | 8.9218348376631e-07 | 2.86269603754088e-05 |
| csnk2b.L | 2840.7608743662 | 0.329762390968476 | 0.0671091887395113 | 4.91381876554148 | 8.93193214973239e-07 | 2.86269603754088e-05 |
| erh.L | 2555.07516346082 | 0.338653407243374 | 0.0689412896911212 | 4.9122000583489 | 9.00600906025212e-07 | 2.88263348367558e-05 |
| LOC108716096 | 276.648078303971 | -0.777974582381399 | 0.158458068045194 | -4.90965586024633 | 9.12363614664376e-07 | 2.9164423229811e-05 |
| cry2.S | 253.017538340015 | -0.671906999962943 | 0.136887022786235 | -4.90847843927619 | 9.17857176477624e-07 | 2.92630213193535e-05 |
| LOC108695484 | 4128.82021776533 | -0.527437503111009 | 0.107451327196594 | -4.90861785397964 | 9.17205042688066e-07 | 2.92630213193535e-05 |
| darmin.L | 5832.97798256106 | -0.421098927637027 | 0.085807940425954 | -4.90745874503776 | 9.22640545769171e-07 | 2.93769717154865e-05 |
| taldo1.L | 3670.10900034419 | 0.311712319494749 | 0.0635522263324782 | 4.90482139624823 | 9.3512388459445e-07 | 2.97354707491329e-05 |
| mcm7.S | 2335.57351728525 | 0.380170843546509 | 0.0775733050098033 | 4.90079471924608 | 9.54497513647606e-07 | 3.03118465314444e-05 |
| nlk.2.L | 2475.00075173084 | -0.38227974990688 | 0.0780554359825036 | -4.89754166503624 | 9.70430621066785e-07 | 3.07775998801521e-05 |
| ints9.S | 710.144301741339 | -0.503769854389091 | 0.102905581318751 | -4.89545705814203 | 9.8077507644523e-07 | 3.1065123477393e-05 |
| neo1.S | 5083.96535762659 | -0.356608973909411 | 0.0728578324949325 | -4.89458664494603 | 9.85125674086486e-07 | 3.11622957373139e-05 |
| epn1.L | 1792.7565352439 | -0.364884871974818 | 0.074564624524764 | -4.89353864919731 | 9.90388537154865e-07 | 3.1288035268713e-05 |
| ezh2.L | 3251.29033638381 | 0.318793311780351 | 0.0651502443243753 | 4.89320209155543 | 9.920844095083e-07 | 3.13009073306424e-05 |
| wasf1.L | 1414.05117327029 | -0.451782722820419 | 0.0923463988946251 | -4.89226140085809 | 9.96839268209247e-07 | 3.1405436837519e-05 |
| LOC121393802 | 78.4086585555899 | 0.903951309184747 | 0.184780191451234 | 4.89203578633218 | 9.97982927412723e-07 | 3.1405436837519e-05 |
| rpl39.L | 5364.07493050776 | 0.326650881562109 | 0.0667924580176314 | 4.89053541757487 | 1.00562061474171e-06 | 3.16048476255305e-05 |
| stmn1.S | 8085.25369564888 | 0.387969600727465 | 0.0793360905001806 | 4.89020316329529 | 1.00731956185058e-06 | 3.16173403560696e-05 |
| adipor2.S | 881.349126256244 | -0.454864499276563 | 0.0930505178283913 | -4.88836075168812 | 1.01679079186888e-06 | 3.17913970368888e-05 |
| LOC108715857 | 12737.1727775167 | 0.335434371500558 | 0.0686188561692602 | 4.88837019773619 | 1.01674201495335e-06 | 3.17913970368888e-05 |
| stau1.L | 3677.72528718954 | -0.368234614050169 | 0.0753262829427312 | -4.88852761167212 | 1.01592950208469e-06 | 3.17913970368888e-05 |
| ppp1cc.S | 3455.12658919705 | 0.323178872738527 | 0.0661314871291056 | 4.88691373456639 | 1.02428946862303e-06 | 3.19846893968225e-05 |
| lrrc8a.L | 958.628398936231 | -0.472511747653713 | 0.0967164127462719 | -4.88553839246821 | 1.03146603418897e-06 | 3.21674400957468e-05 |
| rcvrn.2.S | 47.9547052189453 | 0.841958394852642 | 0.172443875083457 | 4.88250681240585 | 1.0474561940215e-06 | 3.25824593822769e-05 |
| raph1.L | 903.035369040597 | -0.433818613061273 | 0.0888488601896927 | -4.88265816955972 | 1.04665222910027e-06 | 3.25824593822769e-05 |
| slc4a2.S | 439.997628132166 | -0.562160361869072 | 0.115192542018577 | -4.88018019238099 | 1.05988956096398e-06 | 3.2927054979615e-05 |
| LOC108705063 | 2080.03328633562 | -0.358823811481267 | 0.0735671261011082 | -4.87750209228116 | 1.07437708979805e-06 | 3.33345044949603e-05 |
| jpt1.L | 2892.2693353991 | 0.375766981395629 | 0.077049393667295 | 4.87696221229487 | 1.07732063391319e-06 | 3.33831983166927e-05 |
| psmd13.S | 4953.34959462103 | 0.312395882519215 | 0.064068638716511 | 4.87595629901698 | 1.08282580909944e-06 | 3.35110448487412e-05 |
| mtss1.1.S | 3147.3338846638 | -0.366519307215398 | 0.0752197649258763 | -4.87264627291213 | 1.10113272960003e-06 | 3.40342474973322e-05 |
| kif11.S | 1733.78602746623 | 0.367501504117776 | 0.0754851664862908 | 4.86852611214045 | 1.12433667983418e-06 | 3.47072875475116e-05 |
| lhx9.L | 1300.47501254521 | 0.611165041363688 | 0.125556979987609 | 4.8676309467144 | 1.12943993668761e-06 | 3.48205759161024e-05 |
| sim2.L | 31.869168996073 | 0.93468755760986 | 0.192046829133574 | 4.86697729833257 | 1.13318039887808e-06 | 3.48916154757211e-05 |
| ank1.L | 1934.29530730978 | -0.504847736937986 | 0.103746849853367 | -4.86615003396752 | 1.13793145971066e-06 | 3.49935530154568e-05 |
| eif3l.L | 4967.11956406627 | 0.333329630637255 | 0.0685058161718888 | 4.8657128586118 | 1.14044993600418e-06 | 3.50266633947984e-05 |
| hnrnpa2b1.S | 17364.6512635636 | 0.301170057539343 | 0.0619033566049261 | 4.86516521973828 | 1.14361233769439e-06 | 3.50794420857924e-05 |
| hectd1.L | 2236.51518070957 | -0.408617330035486 | 0.0839999149727548 | -4.86449695201946 | 1.14748275553734e-06 | 3.51095038577131e-05 |
| tox2.L | 2798.18073086654 | -0.394009159198196 | 0.0809965182249105 | -4.86451970816961 | 1.14735075130251e-06 | 3.51095038577131e-05 |
| ncbp2.L | 940.408218993738 | 0.429897416815801 | 0.0884015878500861 | 4.86300560058755 | 1.1561657441202e-06 | 3.53306799844733e-05 |
| LOC108707553 | 3568.89095979227 | -0.447365264220589 | 0.0920025305702644 | -4.86253211566609 | 1.15893538591029e-06 | 3.53708244539003e-05 |
| itga10.L | 150.394073432781 | -0.820881558511764 | 0.16887045397598 | -4.86101351174508 | 1.16786281249778e-06 | 3.55985685907416e-05 |
| rpl6.L | 16841.2929817822 | 0.342469827283076 | 0.0704571189305279 | 4.86068451962615 | 1.16980549790726e-06 | 3.56131012107254e-05 |
| LOC108701370 | 174.827305886665 | 0.942046094973223 | 0.193859951266499 | 4.85941572160097 | 1.17732685924568e-06 | 3.57972199230471e-05 |
| fzd7.L | 971.759601742765 | -0.42641379317693 | 0.0877847891863038 | -4.85749065560733 | 1.18882745975673e-06 | 3.61017178841624e-05 |
| garnl3.S | 2344.17469228631 | -0.437921877682292 | 0.0901875659206834 | -4.85567908626592 | 1.19974867771314e-06 | 3.6387883116558e-05 |
| msrb2.S | 662.17567491607 | -0.482569803781926 | 0.0993971634473494 | -4.85496554474155 | 1.20407677337741e-06 | 3.64736173721082e-05 |
| pbrm1.L | 3959.23076830997 | -0.315319620249902 | 0.0649529936511335 | -4.85458179100262 | 1.20641069843349e-06 | 3.6498806360826e-05 |
| sv2c.L | 995.143060798047 | -0.616581109794933 | 0.127037590035217 | -4.85353279784358 | 1.21281272722608e-06 | 3.66468562129731e-05 |
| trappc10.S | 1319.36756687097 | -0.401589790973617 | 0.0828165307157014 | -4.84915013347062 | 1.23991552811815e-06 | 3.74192643976428e-05 |
| pknox2.S | 272.62408204725 | 0.664292188768693 | 0.137022796773146 | 4.8480413800668 | 1.24686396188296e-06 | 3.75375275136056e-05 |
| LOC108704224 | 175.116562779537 | -0.745582241901372 | 0.153790709681467 | -4.84803174031534 | 1.24692453706593e-06 | 3.75375275136056e-05 |
| LOC108709943 | 3033.5784838942 | -0.510278567599753 | 0.105291800756925 | -4.8463276715893 | 1.25767732577756e-06 | 3.7814372465891e-05 |
| plppr3.L | 1562.80500231777 | -0.422199606402337 | 0.0871332034952622 | -4.84545029295628 | 1.2632483742343e-06 | 3.79349270749669e-05 |
| hnrnpk.S | 13187.5394158316 | 0.30145561989083 | 0.0622316258838466 | 4.84409037381552 | 1.27193032623246e-06 | 3.81484880808535e-05 |
| dynll1.S | 3472.65228696307 | 0.363173094479403 | 0.0750082347790124 | 4.8417763136191 | 1.28683576511234e-06 | 3.85004779281272e-05 |
| tln2.S | 1496.35288675772 | -0.46679232431474 | 0.0964085749871592 | -4.84181333845991 | 1.28659596164581e-06 | 3.85004779281272e-05 |
| LOC108712783 | 199.199154707802 | -0.736829882306842 | 0.152235567088287 | -4.84006396402444 | 1.29797345130549e-06 | 3.87859373013721e-05 |
| LOC108717189 | 852.929399863468 | -0.499067512513903 | 0.1031376287286 | -4.83884997809256 | 1.30592572429993e-06 | 3.89612745068208e-05 |
| rps29.S | 3831.48006031218 | 0.346053950478291 | 0.0715182678777075 | 4.8386791339805 | 1.30704860142665e-06 | 3.89612745068208e-05 |
| trim3.S | 1902.33084666753 | -0.370576581909034 | 0.0766874199898979 | -4.83229950828767 | 1.34965004332447e-06 | 4.01818604810352e-05 |
| XB5796436.L | 1412.42828876761 | 0.425064507507856 | 0.0880002404722753 | 4.83026529503376 | 1.36351254358928e-06 | 4.05448882912582e-05 |
| LOC108697444 | 2326.52745196411 | -0.389839820675562 | 0.0807323764601611 | -4.82879159227941 | 1.37364077633618e-06 | 4.07961234967136e-05 |
| rps10.L | 12517.3777361212 | 0.289101300390927 | 0.0599934376359121 | 4.81888206082512 | 1.44364856647827e-06 | 4.28229527155535e-05 |
| rps3a.L | 12039.3506658333 | 0.324061382397622 | 0.0672532492888812 | 4.81852380106782 | 1.44624275401377e-06 | 4.28231996707251e-05 |
| pdhx.S | 762.741024547018 | -0.462710386002216 | 0.0960299981167338 | -4.81839420052624 | 1.44718230549376e-06 | 4.28231996707251e-05 |
| phox2a.S | 237.348727655472 | -0.843412405773976 | 0.17513915142814 | -4.81567027644318 | 1.46706599543447e-06 | 4.33587606971838e-05 |
| gsk3a.L | 1438.45888488832 | -0.402135465930075 | 0.0835126973736805 | -4.81526137433576 | 1.47007342065553e-06 | 4.33948525898e-05 |
| cdr2l.L | 1412.48212328721 | -0.399341952659495 | 0.0829400435972826 | -4.81482689590217 | 1.47327545198138e-06 | 4.34365944544121e-05 |
| LOC108696848 | 1843.06159341067 | -0.371461804849379 | 0.0771765244729348 | -4.81314502546235 | 1.48573388737511e-06 | 4.37508109816858e-05 |
| LOC108698084 | 1029.69337939988 | -0.505439985009488 | 0.105105839372945 | -4.80886683389726 | 1.51788276228915e-06 | 4.46433944637442e-05 |
| eprs1.L | 6236.48349748412 | -0.349500089209673 | 0.0726836181000072 | -4.80851254169527 | 1.52057490597990e-06 | 4.46684966939246e-05 |
| LOC108699406 | 1011.2628557711 | -0.412952351572812 | 0.0858865077451355 | -4.80811669276658 | 1.52358825468467e-06 | 4.47029626320162e-05 |
| sap18.L | 1369.26309750779 | 0.381732644670006 | 0.0794049018611384 | 4.80741913562933 | 1.52891229591025e-06 | 4.48050606958307e-05 |
| LOC108701018 | 1145.86106305175 | -0.54104814765579 | 0.112578200084448 | -4.80597617700348 | 1.53998237122768e-06 | 4.50750984657895e-05 |
| gramd1a.S | 938.467430199417 | -0.445299223192301 | 0.0926621502066517 | -4.80562152075266 | 1.5427149937173e-06 | 4.51007437513456e-05 |
| LOC495017 | 1057.34011008572 | -0.403815298824388 | 0.0840387613242288 | -4.80510769627402 | 1.54668227634967e-06 | 4.51302779145654e-05 |
| tox2.S | 2386.78023680272 | -0.365241590305823 | 0.0760126656484676 | -4.80500962819722 | 1.54744058215333e-06 | 4.51302779145654e-05 |
| psmc1.L | 4329.48529614858 | 0.298874896481486 | 0.0622057224673004 | 4.80467848659096 | 1.55000375764614e-06 | 4.51508288828e-05 |
| LOC108700443 | 299.683008554729 | -0.709826805285293 | 0.14777983346839 | -4.80327246705906 | 1.56093247034134e-06 | 4.54147226760149e-05 |
| LOC108706519 | 834.251407657918 | -0.494638927351031 | 0.102999657156196 | -4.80233566798114 | 1.56825510438848e-06 | 4.55187449295265e-05 |
| LOC121396584 | 155.254680414944 | -0.825542429677603 | 0.171897743093809 | -4.80252046838732 | 1.56680797387449e-06 | 4.55187449295265e-05 |
| fnbp1.L | 2327.75557352671 | -0.347165277275764 | 0.0722977071067505 | -4.80188502746236 | 1.57178935810009e-06 | 4.55668862358992e-05 |
| LOC108702751 | 2508.00774842765 | -0.408950144848687 | 0.0851865144104776 | -4.80064418269458 | 1.58156058589505e-06 | 4.57955099806131e-05 |
| acss1.S | 210.748687472666 | -0.737149771981327 | 0.153585978213178 | -4.79959030477483 | 1.58990534814234e-06 | 4.59823339616309e-05 |
| LOC443608 | 744.869480877313 | -0.496077194431498 | 0.103369932102327 | -4.7990473084613 | 1.59422137911703e-06 | 4.60523355342082e-05 |
| LOC108711545 | 1289.50404076646 | -0.446797188510519 | 0.0931662144642369 | -4.79569971882917 | 1.62107964812507e-06 | 4.67725759757131e-05 |
| ttbk1.S | 1707.45334367528 | -0.416434999545233 | 0.0868936440993192 | -4.79246789407576 | 1.64742141879085e-06 | 4.74762229514887e-05 |
| rdh8.L | 22.9553452048884 | 0.782954684744231 | 0.163390285669563 | 4.79192922355042 | 1.65185179627867e-06 | 4.75474970838792e-05 |
| wls.L | 671.001843062324 | 0.462684977100309 | 0.0965682931844466 | 4.79127218513198 | 1.65727121281072e-06 | 4.76470376852351e-05 |
| sspo.L | 1894.92314999629 | 0.78946710746947 | 0.164790209212493 | 4.79074036766027 | 1.66167028629504e-06 | 4.77170424766569e-05 |
| sfrp1.L | 785.907244491445 | -0.487129166448912 | 0.101745387787164 | -4.78772725765134 | 1.68680676229728e-06 | 4.83516685056803e-05 |
| gal.2.L | 34.3413413654246 | 0.779283130609647 | 0.162770635941139 | 4.78761495342102 | 1.68775067476813e-06 | 4.83516685056803e-05 |
| rps8.L | 15962.0479543478 | 0.317955429319718 | 0.0664154790021166 | 4.78736936173547 | 1.689816632154e-06 | 4.8353834230329e-05 |
| eif3i.L | 5324.66505551441 | 0.292977317931968 | 0.061212072003689 | 4.78626696241081 | 1.69912017487393e-06 | 4.85628535628086e-05 |
| LOC108700413 | 3391.53144284427 | -0.342315900190219 | 0.0715339809624216 | -4.78536068571446 | 1.70680542067056e-06 | 4.87251831842192e-05 |
| rnasek.S | 747.584280843867 | -0.462315694520869 | 0.0967569397274216 | -4.77811406420335 | 1.76946999260445e-06 | 5.04548169017989e-05 |
| dhcr24.L | 704.080074137255 | 0.611412820353503 | 0.128052229284941 | 4.77471437840409 | 1.79962485908506e-06 | 5.12544974520661e-05 |
| rgma.S | 1682.23356291509 | -0.469911251146834 | 0.0984567167963019 | -4.77276986717968 | 1.81709383246331e-06 | 5.16914257211457e-05 |
| snrpf.L | 2357.78798569515 | 0.399152964376491 | 0.0836723768295657 | 4.77042698559329 | 1.8383581414222e-06 | 5.22351727341649e-05 |
| igsf3.S | 3757.25889633152 | -0.355131726195948 | 0.0744958578210225 | -4.76713385929668 | 1.86865163712833e-06 | 5.30339052247612e-05 |
| LOC108715323 | 757.436001842169 | -0.485400813916236 | 0.101891916440704 | -4.763879519321 | 1.8990591972359e-06 | 5.38340071617841e-05 |
| sbf1.S | 1688.84643806849 | -0.377372723761975 | 0.0792243737882615 | -4.76334120065825 | 1.90413468713508e-06 | 5.39149744630064e-05 |
| gpam.S | 1052.7158504523 | -0.463480719696339 | 0.0973145478914738 | -4.76270742390149 | 1.91012689995757e-06 | 5.40216797526999e-05 |
| cherp.L | 1605.04658132015 | -0.348323542601608 | 0.0731576269033318 | -4.76127448833013 | 1.92374183443988e-06 | 5.43434699138169e-05 |
| LOC108719989 | 398.915792163535 | 0.5559318301975 | 0.116768294924924 | 4.76098268416897 | 1.92652579393198e-06 | 5.43589055026523e-05 |

| | | | | | | |
|---|---|---|---|---|---|---|
| rps27a.S | 10219.5168792256 | 0.304012545576648 | 0.0638622966947559 | 4.76043865177201 | 1.93172647691453e-06 | 5.44424165083081e-05 |
| vstm2b.L | 1155.63927400199 | -0.409974529243527 | 0.0861371727003155 | -4.75955405072201 | 1.94021163689445e-06 | 5.46181940981621e-05 |
| slc7a10.S | 1666.90208743986 | -0.57182762602552 | 0.120299482897269 | -4.75336728183477 | 2.00056445352262e-06 | 5.61996519973551e-05 |
| LOC108700447 | 750.671004245146 | -0.540745238563944 | 0.113761551070552 | -4.75332160536902 | 2.00101666986549e-06 | 5.61996519973551e-05 |
| rpl8.L | 13332.5300326795 | 0.352636203156937 | 0.0741954671181487 | 4.7527998252966 | 2.00618948612546e-06 | 5.62798699491131e-05 |
| stk32b.L | 1040.56718316325 | -0.539863374045536 | 0.113594508176918 | -4.75254818837479 | 2.00868874974874e-06 | 5.62849878735825e-05 |
| stard7.L | 2252.42945543143 | -0.385226851334083 | 0.0810654714547133 | -4.7520460242964 | 2.0136851973259e-06 | 5.6306106543306e-05 |
| LOC108703551 | 1438.91867195836 | -0.466390708737804 | 0.0981460562989415 | -4.75200661468487 | 2.01407782111356e-06 | 5.6306106543306e-05 |
| apex1.L | 2922.0091487786 | 0.342977943910467 | 0.0721928162659584 | 4.75085973439428 | 2.02553603963191e-06 | 5.65613477549628e-05 |
| wdr7.S | 2093.12758081584 | -0.461333965232935 | 0.0971132236654819 | -4.75047524755284 | 2.0293913550344e-06 | 5.66039421116023e-05 |
| rps26.L | 6592.68391809757 | 0.381822510568716 | 0.0804717391630823 | 4.74480251750149 | 2.08709850601221e-06 | 5.81467558544273e-05 |
| sez6l2.S | 6019.86285792659 | -0.38832554815983 | 0.0818762830423516 | -4.74283313470597 | 2.10749859144368e-06 | 5.86478474003810e-05 |
| carm1.S | 1462.97296016704 | -0.388902505305338 | 0.0820379291716128 | -4.74052074732169 | 2.13169621373323e-06 | 5.9253349904388e-05 |
| fbxo11.S | 1925.72850340444 | -0.336647862211925 | 0.0710353484867673 | -4.7391597195393 | 2.14606298917254e-06 | 5.95592612512649e-05 |
| sim1.S | 27.0457533738117 | 0.908037511810592 | 0.191608946335064 | 4.73901417015633 | 2.14760487594089e-06 | 5.95592612512649e-05 |
| psmb7.S | 2280.1755773441 | 0.35063460919863 | 0.074033763563641 | 4.73614459566434 | 2.17822205064903e-06 | 6.03394828944897e-05 |
| unc5c.S | 405.731776575574 | -0.612517029030092 | 0.12934146390032 | -4.73565870185405 | 2.18344766320441e-06 | 6.04153502618313e-05 |
| bcl9.S | 1605.59946819568 | -0.345372363105964 | 0.0729399092926468 | -4.73502594745867 | 2.19027075808525e-06 | 6.05351965835302e-05 |
| LOC108718675 | 4020.44994771409 | -0.433171803930101 | 0.091516068504652 | -4.7332868534238 | 2.20912933463421e-06 | 6.09870318813676e-05 |
| shf.S | 553.939356699449 | -0.484353443097381 | 0.102340012025586 | -4.732786654122 | 2.21458226047293e-06 | 6.10681741611002e-05 |
| lrp8.L | 777.169061417781 | -0.487239323361901 | 0.102998471939463 | -4.73054904783706 | 2.2391347793483e-06 | 6.16751992665393e-05 |
| ncaph.L | 886.595164102357 | 0.408282991064631 | 0.0863234537533549 | 4.72968785784667 | 2.24865298553552e-06 | 6.18672430697620e-05 |
| LOC108708803 | 2338.67205439798 | -0.461799248600928 | 0.0976548243684236 | -4.72889334026849 | 2.25746930786588e-06 | 6.19972403496934e-05 |
| pmp2.S | 5154.91820032581 | -0.564253196732558 | 0.119324847526677 | -4.72871500302071 | 2.25945277405736e-06 | 6.19972403496934e-05 |
| rapgef1.S | 1622.96733334742 | -0.398609075082312 | 0.0842979644386014 | -4.72857295827872 | 2.26103379228733e-06 | 6.19972403496934e-05 |
| rora.S | 279.16130452835 | -0.602654762593521 | 0.127477284620042 | -4.72754627924333 | 2.27249279426039e-06 | 6.22411949760563e-05 |
| ahdc1.L | 1862.03229549779 | -0.355777960206155 | 0.0752731284007248 | -4.72649360754786 | 2.28429979717249e-06 | 6.24941208023744e-05 |
| trabd2b.L | 465.108687082717 | -0.527796228158093 | 0.111728796782011 | -4.72390505724188 | 2.31358460232172e-06 | 6.31530610435999e-05 |
| hes5.1.L | 731.479665217605 | 0.575806190384327 | 0.121891970930378 | 4.72390581585734 | 2.31357596737416e-06 | 6.31530610435999e-05 |
| sim2.S | 25.9526101159261 | 0.871958436627861 | 0.184638374751525 | 4.72252010342536 | 2.3294005560847e-06 | 6.35134198760064e-05 |
| dach1.L | 1140.65566809552 | -0.441779062066326 | 0.0935964673141879 | -4.72003991970494 | 2.35798363300463e-06 | 6.42206887670565e-05 |
| c9orf72.L | 975.538091943099 | 0.39199108426037 | 0.0830976387490962 | 4.71723493183653 | 2.39071571876883e-06 | 6.50392471128442e-05 |
| atp1a3.L | 13953.9697488814 | -0.339051882590953 | 0.0718821907709859 | -4.71677169204761 | 2.39616317534547e-06 | 6.51145281676093e-05 |
| kansl1.S | 1266.21461038753 | -0.375883066492031 | 0.079703010366157 | -4.71604604098663 | 2.4047204123773e-06 | 6.52740532941834e-05 |
| cass4.L | 344.530444535924 | -0.624641965348126 | 0.132503060886318 | -4.71417008158055 | 2.42697883575412e-06 | 6.58047141024671e-05 |
| skp1.S | 4623.89605135017 | 0.333829850668566 | 0.0708372271876967 | 4.71263294628996 | 2.44536438265513e-06 | 6.62293002365928e-05 |
| lamc1.L | 2371.21839489729 | -0.371399894764753 | 0.0789013832808577 | -4.7071404748725 | 2.51215801418556e-06 | 6.78869486058109e-05 |
| tmem145.S | 612.88361613502 | -0.531454638068522 | 0.112898563837568 | -4.7073640266421 | 2.50940555403263e-06 | 6.78869486058109e-05 |
| nckipsd.S | 1404.48768165767 | -0.396028638169693 | 0.0841484566888337 | -4.70630898952951 | 2.52242105687932e-06 | 6.80885523953625e-05 |
| rpl37.S | 8211.24692701746 | 0.327380388265192 | 0.0695686146944169 | 4.70586326468084 | 2.52793920662966e-06 | 6.81021887977228e-05 |
| wipf2.S | 744.870814795377 | -0.532111184677106 | 0.113075236041739 | -4.70581537836178 | 2.52853273629481e-06 | 6.81021887977228e-05 |
| foxd3.S | 254.55574497797 | -0.648534676545492 | 0.137822428886764 | -4.70558153548674 | 2.53143303695167e-06 | 6.8104799778188e-05 |
| rps6.S | 14812.166075961 | 0.334908291946105 | 0.0712279349030522 | 4.70192337320949 | 2.57722223990283e-06 | 6.92238692907667e-05 |
| megf8.L | 1921.63169244772 | -0.354979611264286 | 0.0754985951783841 | -4.70180419152911 | 2.57872732807047e-06 | 6.92238692907667e-05 |
| wdr26.L | 2155.58594900581 | -0.330968336223121 | 0.0704246228258455 | -4.69961105861608 | 2.6065744166211e-06 | 6.989417094635e-05 |
| rnf145.L | 708.148518913106 | -0.456564098189064 | 0.0971643500487711 | -4.69888490953621 | 2.61585805855454e-06 | 7.00657725187696e-05 |
| LOC108715250 | 4751.25386582157 | -0.351329450165781 | 0.0747927562457074 | -4.69737268421559 | 2.63529346360549e-06 | 7.05086116793301e-05 |
| ddx39a.S | 6032.84943174927 | 0.319725565767767 | 0.0680689527055769 | 4.69708366383566 | 2.63902374974848e-06 | 7.05307403480633e-05 |
| per3.L | 813.216293730453 | -0.535239882941767 | 0.114009590102014 | -4.69469175762182 | 2.67009037670474e-06 | 7.12826105622692e-05 |
| sf3a3.L | 1846.76888862422 | 0.342404444415072 | 0.0729542268511284 | 4.69341475050908 | 2.68681984621254e-06 | 7.16504954378566e-05 |
| gnas.S | 1084.24551393494 | -0.383198524081097 | 0.0816618532760231 | -4.69250339918025 | 2.69882049470762e-06 | 7.18916064675734e-05 |
| mtmr9.S | 578.257753247388 | -0.540677089548908 | 0.115231258388493 | -4.69210435701448 | 2.704091248113e-06 | 7.19531136710375e-05 |
| ccnb1.2.L | 1414.8134721341 | 0.461603252629589 | 0.0984264045963443 | 4.6898314992066 | 2.73430120780635e-06 | 7.26773671142752e-05 |
| wbp2nl.L | 572.437965572178 | -0.476015513824118 | 0.10153141679978 | -4.68835685374925 | 2.75407458807003e-06 | 7.31229377514463e-05 |
| LOC108704366 | 801.108317484801 | -0.447060811311234 | 0.0953946398843727 | -4.68643533696561 | 2.78004599553245e-06 | 7.37319185758354e-05 |
| naa40.L | 1424.35477697322 | 0.379098449105629 | 0.0808996256664432 | 4.68603465074965 | 2.78549124447533e-06 | 7.3795773493221e-05 |
| tuft1.L | 1799.54281842577 | -0.340827258242525 | 0.0727624963400003 | -4.68410617263498 | 2.81184237611859e-06 | 7.44127436660405e-05 |
| dusp8.L | 1774.14656087128 | -0.425968756595384 | 0.0909719927552075 | -4.68241646351096 | 2.83512738614527e-06 | 7.49473174309175e-05 |
| dcbld2.L | 1244.86084821566 | -0.452805825422791 | 0.0967196388455846 | -4.68163271521006 | 2.84599052452664e-06 | 7.50458469033063e-05 |
| ina.L | 2748.1054992632 | -0.343666628763041 | 0.0734098364511854 | -4.68147928638371 | 2.84812179323489e-06 | 7.50458469033063e-05 |
| grid2ip.S | 300.898743224939 | -0.664487600144513 | 0.141938564648687 | -4.68151556829667 | 2.84761766560153e-06 | 7.50458469033063e-05 |
| hspa4.S | 4131.64839254721 | -0.339790095396763 | 0.0725855224941868 | -4.68123785185917 | 2.85147864401695e-06 | 7.50528951004852e-05 |
| h2az1.S | 2757.33272308149 | 0.409601888546339 | 0.0875042562500635 | 4.68093674638875 | 2.85567046590581e-06 | 7.50818812756665e-05 |
| rps3.S | 9233.93776675157 | 0.316887054962203 | 0.0677097641097188 | 4.68007914558247 | 2.86764193186237e-06 | 7.52337938365708e-05 |
| vapbl.S | 1128.18405960477 | -0.417470951312425 | 0.0892001310840492 | -4.68016073787001 | 2.86650089504306e-06 | 7.52337938365708e-05 |
| uhrf1.S | 3199.86228465941 | 0.362550838656751 | 0.0774906799794628 | 4.67863798269465 | 2.88786801622495e-06 | 7.56827028977012e-05 |
| rps25.S | 10073.3404133685 | 0.308583324587314 | 0.0659679765893866 | 4.67777458914756 | 2.90005085321904e-06 | 7.59200812802838e-05 |
| rnf44.S | 1700.75372730009 | -0.380720260775086 | 0.0813965089822235 | -4.67735367936029 | 2.90600792765513e-06 | 7.59941405753e-05 |
| XB5909790.S | 2789.73236200481 | 0.339493122308579 | 0.0725864879698762 | 4.67708428667152 | 2.90982676267157e-06 | 7.60121842713367e-05 |
| LOC398653 | 6342.49683952781 | 0.351772068257432 | 0.0752214127868996 | 4.67648845221763 | 2.91829025360951e-06 | 7.61513892816213e-05 |
| LOC108708766 | 944.623995125253 | -0.45358152807901 | 0.0970021325053069 | -4.67599542777263 | 2.92531123883705e-06 | 7.62526944595571e-05 |
| LOC121398039 | 732.32136781901 | -0.490301578952255 | 0.104871113677989 | -4.67527769808707 | 2.93556114709873e-06 | 7.64378590649695e-05 |
| kalrn.S | 3940.69840184057 | -0.350257343799512 | 0.0749243039693081 | -4.6748161176511 | 2.94217118010547e-06 | 7.65279514448419e-05 |
| pitpnc1.S | 877.420575468831 | -0.49272590158154 | 0.105423467059407 | -4.67377819498088 | 2.95708688192598e-06 | 7.68336563738072e-05 |
| LOC108706899 | 1743.4799389383 | -0.42320602999313 | 0.0905647185305593 | -4.67296798201087 | 2.9687806215447e-06 | 7.70491436499133e-05 |
| irf2bpl.S | 1479.69591323331 | -0.352062972502444 | 0.0753440273484906 | -4.67273896674037 | 2.97209401494298e-06 | 7.70491436499133e-05 |
| LOC108702731 | 3590.40100778701 | -0.412410861597068 | 0.0882625660428363 | -4.67254556588479 | 2.97489490177075e-06 | 7.70491436499133e-05 |
| btbd10.L | 1024.50150040893 | -0.435501742532569 | 0.0932493644638117 | -4.6702918034533 | 3.00772180369637e-06 | 7.7816393502662e-05 |
| ddx21.S | 9985.07019067684 | 0.312329583043613 | 0.0668882975671376 | 4.66942042784269 | 3.02050663787625e-06 | 7.80640300644314e-05 |
| thdl20.L | 128.274603119844 | -0.903952274877712 | 0.193625463264003 | -4.66856094048537 | 3.03316810982334e-06 | 7.83079554304444e-05 |
| rpl36a.L | 14723.2701094976 | 0.321131606595772 | 0.0687920894152125 | 4.66814730190123 | 3.03927972801659e-06 | 7.83824434314597e-05 |
| LOC108697935 | 2435.47459359263 | -0.332261097391693 | 0.0712051589674925 | -4.66625034210486 | 3.06745944312755e-06 | 7.90253019208278e-05 |
| LOC108697348 | 4289.11937138473 | -0.28713512471035 | 0.0615653142813479 | -4.66391064615001 | 3.10256140139154e-06 | 7.98449435226759e-05 |
| mcm4.S | 1591.78407380936 | 0.389929054747327 | 0.0836978634914764 | 4.65876951311949 | 3.18105129347242e-06 | 8.17782646810784e-05 |
| tfap2e.S | 280.376485501759 | 0.734473427556486 | 0.157728771690244 | 4.65655960980211 | 3.21537212831144e-06 | 8.25732034727254e-05 |
| herc1.L | 5304.71523427648 | -0.327325967992395 | 0.0703244468192554 | -4.65451180630931 | 3.24749229905832e-06 | 8.33100083562016e-05 |
| rab10.L | 3505.48901072445 | -0.366921372345585 | 0.0788462369817615 | -4.653632010751 | 3.26138636013489e-06 | 8.34354273895924e-05 |
| rps27.L | 14846.9758057403 | 0.323074733075948 | 0.0694254350483201 | 4.65354999721771 | 3.26268444966299e-06 | 8.34354273895924e-05 |
| eif2s2.L | 5550.58167834514 | 0.290202227226435 | 0.0623593987387663 | 4.6537047036348 | 3.26023620959645e-06 | 8.34354273895924e-05 |
| LOC121393629 | 85.1259271488625 | 0.880065495725324 | 0.189166795721948 | 4.65232543780523 | 3.28212550160938e-06 | 8.37562572858176e-05 |
| LOC108717874 | 431.674430552926 | -0.572562366566631 | 0.123068503210222 | -4.65238750477528 | 3.28113746198291e-06 | 8.37562572858176e-05 |
| ubc.S | 13255.8481254704 | 0.28903039006491 | 0.0621379041468877 | 4.65143448323702 | 3.29634002849747e-06 | 8.40307289111413e-05 |
| rcan2.S | 635.536644413267 | -0.502791346808957 | 0.108163862634318 | -4.64842262992033 | 3.34483037893962e-06 | 8.51774729831857e-05 |
| ppm1a.S | 2225.85181865348 | -0.364367546609619 | 0.0783912059140391 | -4.64806660850671 | 3.3506072713614e-06 | 8.52352387962866e-05 |
| st13.S | 5301.27866869921 | 0.296021519998256 | 0.0636970206670085 | 4.64733698528507 | 3.36247623212187e-06 | 8.53886991534655e-05 |
| LOC108701216 | 1876.0565874444 | -0.336642702178766 | 0.0724389057021268 | -4.6472637723582 | 3.36366942824839e-06 | 8.53886991534655e-05 |
| aplp1.L | 4734.32935243784 | -0.48824799640718 | 0.105079069649432 | -4.64648191153661 | 3.37643724378732e-06 | 8.56233469734542e-05 |
| rpl37.L | 8182.99791593827 | 0.359695205808424 | 0.07741664466349 | 4.64622572280064 | 3.38063091488887e-06 | 8.56402997354643e-05 |
| LOC108703931 | 2226.09852964322 | -0.378494495404929 | 0.0815502804449103 | -4.64124088035005 | 3.46323117311635e-06 | 8.76413938746757e-05 |
| prrt1.S | 5357.34774804899 | -0.321117769543885 | 0.0692406136017489 | -4.63770831655042 | 3.52293472407289e-06 | 8.9059496552161e-05 |
| LOC108709624 | 4368.19743309447 | 0.359256416373927 | 0.0774704578342504 | 4.63733436443057 | 3.52931230366375e-06 | 8.9127976200839e-05 |
| MGC114789 | 7442.12789101575 | 0.348891720565899 | 0.0752615797335011 | 4.63572146374437 | 3.55694667542301e-06 | 8.97325675313879e-05 |
| ptn.S | 9428.90086096958 | -0.326902785849514 | 0.0705462771560474 | -4.63387720838067 | 3.58879915606930e-06 | 9.04422061177878e-05 |
| LOC108709000 | 1823.11765773358 | -0.39104780968118 | 0.0844008221275305 | -4.63322275570139 | 3.60016793440585e-06 | 9.05408693565795e-05 |
| mapk8ip2.L | 4217.85138045751 | -0.42735517553331 | 0.0922335400166039 | -4.63340315742383 | 3.59703065613024e-06 | 9.05408693565795e-05 |
| atp2b3.L | 2557.97285740786 | -0.380325728504654 | 0.0820924529168607 | -4.63289516869266 | 3.60587154857993e-06 | 9.05905309216141e-05 |
| immt.L | 2868.8311105526 | -0.348255040723507 | 0.075195178970152 | -4.63134798657429 | 3.63292680689743e-06 | 9.0987962728625e-05 |
| psmd14.L | 1971.61851093511 | 0.365729600356365 | 0.0789623745990816 | 4.63169455342873 | 3.62684960039498e-06 | 9.0987962728625e-05 |

| | | | | | | |
|---|---|---|---|---|---|---|
| LOC108702162 | 1093.78296885636 | -0.476970765829174 | 0.102987325712407 | -4.63135402856384 | 3.6328207743094e-06 | 9.0987962728625e-05 |
| gtf2ird1.S | 1009.49849005505 | 0.443510419491897 | 0.0957880349942886 | 4.63012337102794 | 3.65447928869092e-06 | 9.14334910808004e-05 |
| slc40a1.S | 2348.85806777804 | -0.486167512615675 | 0.10500725453403 | -4.62984690698794 | 3.65936181882675e-06 | 9.14614568174466e-05 |
| eef2.1.L | 73697.5130124261 | 0.269809923405136 | 0.0583023256470177 | 4.62916222311865 | 3.67148070763027e-06 | 9.16700434852721e-05 |
| dbi.L | 3699.23539712383 | 0.309809167260767 | 0.0669359180902189 | 4.62844428074018 | 3.68422959376083e-06 | 9.18939155552624e-05 |
| sgip1.L | 2796.61216802382 | -0.384637265536433 | 0.0831405787329428 | -4.62634818518563 | 3.72169437588082e-06 | 9.27331724796396e-05 |
| LOC108719777 | 887.470423644235 | -0.508804300949679 | 0.110042484123808 | -4.62370787974238 | 3.76940612051428e-06 | 9.38257707907519e-05 |
| robo3.S | 2297.56739659268 | -0.417862107330999 | 0.0903903232222952 | -4.62286329371076 | 3.78479163313314e-06 | 9.41123110904162e-05 |
| rps15a.S | 7176.18031084698 | 0.336092274888076 | 0.0727415463955706 | 4.62036197389036 | 3.83071120208764e-06 | 9.51567463635144e-05 |
| LOC108709237 | 365.532888575465 | -0.684787433247987 | 0.148223670499404 | -4.61996003027561 | 3.83813976213687e-06 | 9.51580103378007e-05 |
| ier5l.S | 1518.07751312061 | -0.342760099581993 | 0.0741915356082104 | -4.61993537095708 | 3.83859595501131e-06 | 9.51580103378007e-05 |
| sap130.S | 1095.36574930539 | -0.400478943221106 | 0.0867067168983026 | -4.61877646331396 | 3.8600942509512e-06 | 9.55934044165224e-05 |
| stt3a.S | 2709.1836858539 | 0.334572107356455 | 0.0724900685920909 | 4.61541993068218 | 3.92301269500542e-06 | 9.7052617527965e-05 |
| adar.S | 1968.62115679184 | -0.392534916817553 | 0.0850618417619431 | -4.61470041897416 | 3.93662735229379e-06 | 9.72903610342068e-05 |
| LOC108698041 | 860.377389610197 | -0.504871917561448 | 0.109466497653826 | -4.61211355421311 | 3.98595128799573e-06 | 9.84092485676507e-05 |
| dlx6.L | 29.2539406441083 | 0.741258943971801 | 0.16076653626257 | 4.61077884243987 | 4.01163143126663e-06 | 9.89427147118696e-05 |
| srf.S | 562.809380009428 | -0.511145831452919 | 0.110873936821207 | -4.61015317131907 | 4.02372399532852e-06 | 9.91403151546766e-05 |
| kif2c.S | 728.204094747841 | 0.424540887420483 | 0.0921004078748915 | 4.60954405323781 | 4.03553018884371e-06 | 9.93304664719038e-05 |
| MGC114621 | 15701.2619735032 | 0.317010084682094 | 0.0688191359919854 | 4.60642349126575 | 4.096536864621e-06 | 0.000100730026912047 |
| LOC108711497 | 23.9582806152359 | 0.745281956705542 | 0.161843288918725 | 4.60496052499161 | 4.12544108563158e-06 | 0.00010133818577789 |
| ckmt1b.L | 1164.09805007721 | 0.379056117181339 | 0.0823217846162625 | 4.60456632406943 | 4.13326278384077e-06 | 0.000101427763707705 |
| galnt11.S | 770.784505433202 | -0.456646081122275 | 0.0992006906357761 | -4.60325505997625 | 4.15938315334502e-06 | 0.000101965746041739 |
| LOC108713772 | 1256.26598161878 | -0.376304071234182 | 0.0817559606011081 | -4.60277230513125 | 4.16903940551662e-06 | 0.000102099438828247 |
| snrpf.S | 2179.94347995433 | 0.389757101866476 | 0.0847013048218309 | 4.60154778826997 | 4.19362912099687e-06 | 0.000102598213359011 |
| lcat.L | 1729.58718926513 | -0.494168300306366 | 0.10743747187118 | -4.59958980511833 | 4.23323670990869e-06 | 0.00010345324467905 |
| fscn1.L | 3988.25246737186 | -0.328553760469912 | 0.0714340476066971 | -4.59940002670532 | 4.23709469233781e-06 | 0.00010345324467905 |
| slc2a1.L | 2750.23774197773 | -0.481966851051315 | 0.104802987544486 | -4.59878923629666 | 4.24953426077466e-06 | 0.000103652796517329 |
| ppm1e.S | 3388.57831122116 | -0.373686377187137 | 0.0812953551791745 | -4.59665101854263 | 4.29335821261405e-06 | 0.000104616694500748 |
| smg1.S | 2745.66293384536 | -0.414309589016651 | 0.0901373997213154 | -4.5964226868936 | 4.29806350363138e-06 | 0.000104626407572365 |
| ahcy.S | 2725.91621240208 | 0.341652207824645 | 0.0743559777470576 | 4.59481830750541 | 4.33126498497884e-06 | 0.000105329080625702 |
| wdr47.L | 820.595067005571 | -0.430581948476969 | 0.0937322878887274 | -4.5937420090303 | 4.35367578319723e-06 | 0.000105768199476994 |
| vstm2l.L | 2183.99218950465 | -0.401397984247369 | 0.0874053994355807 | -4.59237057252059 | 4.38239298970007e-06 | 0.000106359495795978 |
| polr2b.S | 2558.30386546109 | 0.330913891723334 | 0.0720638970974798 | 4.59195110244609 | 4.39121267052932e-06 | 0.000106467186245349 |
| sumo2.L | 2232.26279390147 | 0.322767744442979 | 0.0702957577136626 | 4.59156789723397 | 4.39928471703678e-06 | 0.000106556553255924 |
| rusc2.S | 1659.13024190392 | -0.390039985223577 | 0.0849555865128444 | -4.59110461399271 | 4.40906255526047e-06 | 0.00010668701764691 |
| LOC108701130 | 3280.25762547745 | -0.351736771747937 | 0.0766481524415089 | -4.5889791279229 | 4.45418958108788e-06 | 0.000107671723067611 |
| thumpd1.S | 862.885828700435 | 0.438157842678837 | 0.0955002112958912 | 4.5880300863553 | 4.4744816479197e-06 | 0.00010805472878187 |
| ncor2.S | 1200.38661198884 | -0.512427645308268 | 0.111704320867691 | -4.58735742116206 | 4.48891793592347e-06 | 0.000108295702418396 |
| sema5b.S | 1588.69417773047 | -0.384212726122928 | 0.0837603779831213 | -4.5870462308605 | 4.49561157937896e-06 | 0.00010834959098158 |
| rab3d.L | 377.10510665182 | -0.538238964674259 | 0.11743641758454 | -4.58323725931772 | 4.57832055387744e-06 | 0.000110233616983051 |
| en1.L | 278.613755040432 | -0.728827069608907 | 0.159032379078668 | -4.58288478001313 | 4.58604763961741e-06 | 0.000110310337977094 |
| taok2.L | 2869.77937499882 | -0.382446580076555 | 0.0834715739493567 | -4.58175833977433 | 4.61082541884414e-06 | 0.000110796629797626 |
| map7d1.L | 1747.90617122244 | -0.45287427040987 | 0.0989444536750566 | -4.57705564676878 | 4.71566052806485e-06 | 0.000113203811135185 |
| atp2b4.L | 3838.48729647992 | -0.467359282177673 | 0.102116431755083 | -4.57672946601375 | 4.72301598861499e-06 | 0.000113268460441671 |
| psma3.L | 3320.77649949744 | 0.342472069023584 | 0.0748358918701433 | 4.57630771098243 | 4.73254296629327e-06 | 0.00011338500870131 |
| sf3b1.S | 4863.9919615415 | 0.285975189416576 | 0.062506748198685 | 4.57510905074713 | 4.75971997283358e-06 | 0.00011392378031529 |
| LOC108718546 | 2083.73907651635 | -0.416735900645678 | 0.0911053737316493 | -4.57421866105475 | 4.78000431184133e-06 | 0.000114296677905387 |
| h2az1.L | 3165.85844173402 | 0.397753687119964 | 0.0869805761499847 | 4.57290241943326 | 4.81014189655696e-06 | 0.00011490421557026 |
| dctn1.L | 7030.88798072378 | -0.370821991628104 | 0.0811716245686504 | -4.56836971784991 | 4.91532443841437e-06 | 0.000117301465527347 |
| gpaa1.L | 1002.8607052149 | -0.421064664193901 | 0.092176454882077 | -4.56802840522101 | 4.92333324330233e-06 | 0.000117377289315787 |
| pik3c2a.L | 1768.44681289557 | -0.344319393214563 | 0.0754081179726245 | -4.56607859301783 | 4.96932520248937e-06 | 0.000118297203120898 |
| rnf208.S | 915.60643384436 | -0.464667634300102 | 0.101767334889007 | -4.56598018221557 | 4.97165737986485e-06 | 0.000118297203120898 |
| sf3b5.S | 2003.7229492583 | 0.381811582683458 | 0.0836461072127266 | 4.56460671519889 | 5.00431593479495e-06 | 0.000118957780156466 |
| ogdh.L | 2123.60376305366 | -0.31515238177774 | 0.0690486849798421 | -4.56420541346652 | 5.01389688040232e-06 | 0.00011906902327712 |
| acp6.S | 881.710332479569 | -0.425562905317733 | 0.0933090249073901 | -4.5607904030731 | 5.09614296539577e-06 | 0.000120881457822835 |
| grina.L | 939.118780169987 | -0.444035809632838 | 0.0973629376233826 | -4.56062461211317 | 5.1001685300241e-06 | 0.000120881457822835 |
| LOC121393195 | 35.2088621243108 | 0.877930782380631 | 0.192540489545514 | 4.55972031884287 | 5.1221792997197e-06 | 0.000121284818623188 |
| diras3.S | 44.2523626798249 | 0.886194014389613 | 0.194437724648161 | 4.55772672712149 | 5.17102568105712e-06 | 0.000122322198535153 |
| itsn1.L | 1952.54030035339 | -0.38485589294434 | 0.0844932917127943 | -4.55486921083066 | 5.24181818756431e-06 | 0.000123876197518178 |
| camk2g.S | 6092.23209415602 | -0.357930876362103 | 0.0785904000218722 | -4.55438420293685 | 5.25392559036249e-06 | 0.000124041660147975 |
| soga1.L | 1460.81444233079 | -0.421251250955412 | 0.0925467534093631 | -4.55176692252063 | 5.31972507897711e-06 | 0.000125473204921039 |
| rpl7a.L | 21605.0562815206 | 0.293198374820542 | 0.0644284373786009 | 4.55076029700394 | 5.3452415324751e-06 | 0.0001259527621629 |
| lmx1a.L | 277.169372539184 | 0.631644010226461 | 0.138846124339395 | 4.54923760552704 | 5.384062300351e-06 | 0.000126744582872798 |
| plekhm2.L | 1588.7248310748 | -0.350391913965528 | 0.0770511037555255 | -4.54752621165925 | 5.42801617499749e-06 | 0.000127655590469883 |
| LOC108704253 | 576.552365717589 | 0.518447661514547 | 0.114021378182896 | 4.5469338274699 | 5.44331030632606e-06 | 0.000127891470582094 |
| ift172.L | 985.564185954864 | -0.433024625590836 | 0.0952620496605784 | -4.54561524902851 | 5.47750152004424e-06 | 0.000128570456015044 |
| psmc2.S | 2886.35643703657 | 0.281687035354034 | 0.0619861101501801 | 4.54435735153508 | 5.51031081957069e-06 | 0.000129215724952365 |
| LOC108698377 | 574.734173636258 | -0.559511061286509 | 0.123171090375632 | -4.54255182429724 | 5.55773274695896e-06 | 0.000130202082309181 |
| maz.S | 1308.62998867224 | -0.408846666024242 | 0.0900097908870224 | -4.54224659334465 | 5.56578810470091e-06 | 0.000130265179398462 |
| neurog1.L | 462.102477174475 | 0.610954157735691 | 0.134540674825079 | 4.54103681678431 | 5.59782542220226e-06 | 0.000130888903567836 |
| LOC108706535 | 729.596894108094 | -0.567365579839833 | 0.124957127664821 | -4.54048192722313 | 5.61257899749136e-06 | 0.00013098174271379 |
| mmp24.S | 759.818459181352 | -0.531253697140309 | 0.117001558089186 | -4.54056942331788 | 5.61025015492946e-06 | 0.00013098174271379 |
| rplp2.S | 4934.42778651098 | 0.364235948211621 | 0.080225625601194 | 4.5401446917006 | 5.62156369731807e-06 | 0.000131065516758777 |
| LOC108709287 | 1351.68894731903 | -0.400558385462296 | 0.0882734899404448 | -4.53769739626857 | 5.68717877319442e-06 | 0.000132468188989439 |
| LOC108706570 | 9654.57949209433 | 0.30489069357594 | 0.0671959798636823 | 4.53733533158471 | 5.6969482170178e-06 | 0.000132568639831638 |
| rpl23a.S | 9550.042338859 | 0.312658101538138 | 0.0689215880265875 | 4.53643205982899 | 5.72139090750584e-06 | 0.000132882859184462 |
| cdc42bpa.L | 1607.67376925788 | -0.448616553937877 | 0.0988915450047426 | -4.53645004652686 | 5.72090320628383e-06 | 0.000132882859184462 |
| prss35.L | 2023.86793392779 | -0.437373299580001 | 0.0964447193951325 | -4.53496367995109 | 5.7613398838862e-06 | 0.000133682895070804 |
| hadha.L | 1768.95901999269 | -0.465639738524375 | 0.1027198737159 | -4.53310271595767 | 5.81235327335517e-06 | 0.000134737891624895 |
| psmb5.L | 3291.64622162977 | 0.346451108036941 | 0.0764510384050478 | 4.53167301929107 | 5.85183797429625e-06 | 0.000135394811188146 |
| nlk.2.S | 2827.63309536587 | -0.338606965327082 | 0.0747180541857474 | -4.53179581584543 | 5.84843658324989e-06 | 0.000135394811188146 |
| LOC108718876 | 457.340931830347 | 0.526205194373711 | 0.116127822640747 | 4.53125859426108 | 5.86333128257068e-06 | 0.000135531655736225 |
| pax8.S | 120.700205682241 | -0.880856961997873 | 0.19442772821784 | -4.53051100309596 | 5.88411899595542e-06 | 0.000135882877269716 |
| lss.S | 1052.01407471978 | 0.637398841013604 | 0.140748858209516 | 4.52862530554092 | 5.93686700734268e-06 | 0.000136970794944333 |
| hmgn3.L | 3461.43160988108 | 0.385956841751747 | 0.0853173268946876 | 4.5237802894148 | 6.07447829528493e-06 | 0.00014001269042282 |
| nsa2.S | 2928.45362263665 | 0.304800370918676 | 0.0673864580058388 | 4.52316949040809 | 6.0920417382244e-06 | 0.000140284418946373 |
| prdx1.L | 3464.89141380809 | 0.33546524081312 | 0.0741821154478813 | 4.52218487957263 | 6.12045644128425e-06 | 0.000140805273470227 |
| camk4.S | 306.092210593155 | -0.604133379117692 | 0.133622057427104 | -4.52120997648364 | 6.14871592057015e-06 | 0.000141321574810153 |
| camsap1.S | 2486.20015609345 | -0.345600542301612 | 0.076451124704583 | -4.52054228943076 | 6.16814213401863e-06 | 0.000141634069001747 |
| ahnak.S | 1053.15254292052 | -0.549488751528317 | 0.12162314701108 | -4.51796195898679 | 6.24376980107159e-06 | 0.000143235263028549 |
| tmem63b.L | 1550.95618301345 | -0.380583282671193 | 0.084258998185038 | -4.51682658076955 | 6.27732730758256e-06 | 0.000143598295301705 |
| pgd.L | 1826.38911595293 | 0.360955586081796 | 0.0799104269453671 | 4.51700234724779 | 6.27212104575078e-06 | 0.000143598295301705 |
| LOC108700350 | 3898.65119002432 | -0.450534166900769 | 0.0997441668409247 | -4.51689739029347 | 6.27522940863556e-06 | 0.000143598295301705 |
| erh.S | 1814.71308745943 | 0.348182845792283 | 0.0770964401714731 | 4.51619873781302 | 6.29595801228601e-06 | 0.00014361878305209 |
| snrpb.S | 4844.42952412123 | 0.375190606649406 | 0.0830727087280099 | 4.51641233798967 | 6.28961367885451e-06 | 0.00014361878305209 |
| stx1b.L | 8034.86157254458 | -0.399438849591669 | 0.088444937786485 | -4.51624320835586 | 6.29463664767274e-06 | 0.00014361878305209 |
| smad4.L | 1462.57012055775 | -0.380183585192923 | 0.0842127866444973 | -4.51455889706941 | 6.34486878019037e-06 | 0.000144598726215708 |
| hnrnpc.L | 3259.84288557986 | 0.334178268597955 | 0.0740388414585931 | 4.51355345403192 | 6.37503723154383e-06 | 0.0001451194052933 |
| fjx1.L | 725.997739122912 | -0.422284681104954 | 0.0935624382952225 | -4.51339970183865 | 6.37966266787044e-06 | 0.0001451194052933 |
| kat6a.L | 2569.37119470515 | -0.349647785662893 | 0.0774933462499946 | -4.51197170573748 | 6.42277576607826e-06 | 0.000145963437288218 |
| rnmt.L | 1795.71805863113 | 0.355682695069379 | 0.0788678400275876 | 4.50985718570412 | 6.48712828605562e-06 | 0.000147288125777042 |
| hpf1.L | 646.480557477847 | 0.464840842372585 | 0.103134516076096 | 4.50713165735524 | 6.57098642185767e-06 | 0.000148776935774905 |
| sv2c.S | 523.695088389542 | -0.67418956943528 | 0.149582936543083 | -4.50712885450741 | 6.57107319035454e-06 | 0.000148776935774905 |
| LOC108699714 | 1197.15150725383 | -0.563097746308839 | 0.124933135156969 | -4.50719295246492 | 6.56908916711146e-06 | 0.000148776935774905 |
| fau.L | 6367.23316787536 | 0.335569298337829 | 0.0744598510414386 | 4.5067146071925 | 6.58390921309542e-06 | 0.000148928762032533 |
| col5a2.L | 2679.25955344951 | -0.370067668409052 | 0.0821599562971841 | -4.50423399776973 | 6.66127766298479e-06 | 0.000150538678646095 |
| aifm3.L | 901.348279349753 | -0.580529402883157 | 0.128904150645526 | -4.50357416713101 | 6.68200330085428e-06 | 0.000150547581161487 |
| magoh.L | 1480.74178906778 | 0.392893348481554 | 0.0872403061589549 | 4.50357599348274 | 6.68194584911136e-06 | 0.000150547581161487 |

| | | | | | | |
|---|---|---|---|---|---|---|
| psmb1.S | 2178.24555545053 | 0.32364595573317 | 0.0718665043251945 | 4.50343256252842 | 6.68645921104985e-06 | 0.000150547581161487 |
| cbx3.S | 3823.43716569097 | 0.286646464423806 | 0.0636493289590044 | 4.50352688884483 | 6.68349070344815e-06 | 0.000150547581161487 |
| taok2.S | 3202.07159207752 | -0.339516950326857 | 0.0753953327164782 | -4.50315607205555 | 6.69516780630113e-06 | 0.000150604080265074 |
| per3.S | 1486.15550074406 | -0.400877461982832 | 0.0890255539201762 | -4.50294824722207 | 6.70172078797182e-06 | 0.000150612030363541 |
| map4.S | 2294.72614547086 | -0.38381133840222 | 0.0852502120335132 | -4.50217458991583 | 6.72616916322133e-06 | 0.000151021768624121 |
| rnf10.L | 2941.67219401787 | -0.287070608044661 | 0.063779105075695 | -4.50101342287505 | 6.76302343581187e-06 | 0.000151585408835572 |
| sdhb.L | 1125.91287140509 | -0.446624410526023 | 0.0992280269476864 | -4.5009905393109 | 6.76375167439532e-06 | 0.000151585408835572 |
| rps26.S | 9383.5560566633 | 0.349129689501126 | 0.0775774684632969 | 4.50040065004576 | 6.78255001538348e-06 | 0.000151866608362881 |
| hsf2.2.L | 996.484339077289 | 0.410736932064461 | 0.0913103732470867 | 4.49825049945862 | 6.85149407728354e-06 | 0.000153144813052931 |
| ednrb2.L | 183.70715294466 | 0.871273612769369 | 0.193698042971248 | 4.49810230090295 | 6.85627064122524e-06 | 0.000153144813052931 |
| evl.S | 1593.32267923996 | -0.368191502308129 | 0.0818561379251724 | -4.49803168877458 | 6.85854764969084e-06 | 0.000153144813052931 |
| erbb2.L | 977.766202128714 | -0.540049435009405 | 0.120093754787788 | -4.49689857698013 | 6.89518585553863e-06 | 0.000153821529085818 |
| epha4.L | 4230.0913103956 | -0.354774309450527 | 0.0789302834829617 | -4.49478063165845 | 6.96417039082326e-06 | 0.000155075669545976 |
| psmb3.S | 1437.35348623896 | 0.350088794922463 | 0.0778863236496137 | 4.49486865624064 | 6.96129020227212e-06 | 0.000155075669545976 |
| cs.L | 5609.31905546079 | -0.304912328797706 | 0.0678521905753986 | -4.4937728054466 | 6.99722807868375e-06 | 0.000155669101596651 |
| matr3.S | 2182.04271232181 | 0.332340356030586 | 0.0739651530605149 | 4.49320176162795 | 7.0160254333434e-06 | 0.00015594448479199 |
| spon1.L | 1166.12380495656 | -0.484879089827386 | 0.107929197412135 | -4.49256643664124 | 7.03699552347322e-06 | 0.000156267613571534 |
| dennd2a.L | 731.63537990412 | -0.454005226753078 | 0.101094687213396 | -4.49089105735832 | 7.09258241763093e-06 | 0.000157214596034604 |
| taf9b.S | 787.251058875635 | 0.431114396530173 | 0.0959957732255146 | 4.49097269644772 | 7.08986403371774e-06 | 0.000157214596034604 |
| xnf7.L | 9230.09768575046 | -0.277094707404338 | 0.0617060932980069 | -4.49055664674996 | 7.10372788902284e-06 | 0.000157318108783884 |
| mcm5.L | 2144.4266975408 | 0.323401227169674 | 0.0720549641164324 | 4.48825741758882 | 7.18081295721162e-06 | 0.00015888039160519 |
| stt3a.L | 2716.08107488297 | 0.349208307301776 | 0.0778959184012272 | 4.48301161946727 | 7.35969203074231e-06 | 0.00016269004385337 |
| purb.S | 907.268055804074 | -0.449588068448483 | 0.100334054178993 | -4.48091201065623 | 7.43247506648299e-06 | 0.000164149590241034 |
| rfc2.L | 957.756641026837 | 0.42040705101857 | 0.0938799709930521 | 4.47813358452874 | 7.52984794498711e-06 | 0.000166149069914184 |
| psma4.S | 1771.46458036413 | 0.345952059319937 | 0.0772694293314742 | 4.47721773427179 | 7.5622113339322e-06 | 0.000166711762383438 |
| hes5.2.L | 1068.08810402639 | 0.475466314691629 | 0.106225268089558 | 4.47601896651149 | 7.60477319536035e-06 | 0.000167498059844138 |
| sh3d19.L | 349.643620327646 | -0.557337090509793 | 0.124588414297258 | -4.47342631057197 | 7.69760933473445e-06 | 0.000169389240197499 |
| LOC108717850 | 1654.87604534288 | -0.480611261176021 | 0.10744443108356 | -4.47311467266506 | 7.70884093790864e-06 | 0.00016948287940774 |
| LOC108699394 | 3682.67794371258 | 0.310386397041617 | 0.0694037495001213 | 4.47218484991326 | 7.74244546006494e-06 | 0.000170067784816291 |
| map7d2.S | 1988.31316952149 | -0.499171190734828 | 0.111628650507139 | -4.47171213185009 | 7.75958351799594e-06 | 0.000170290263763499 |
| bcl9l.L | 1018.96058119726 | -0.406689383847038 | 0.0909591962837301 | -4.47111892434105 | 7.78114115331821e-06 | 0.00017060924474613 |
| dhrs3.L | 258.07123792248 | 0.63285461012311 | 0.141549762313791 | 4.47089843019432 | 7.78916867475809e-06 | 0.000170631256794024 |
| bhlhe22.L | 571.848483395406 | -0.510617533381453 | 0.11426342573791 | -4.46877493899644 | 7.86688491933495e-06 | 0.000172178470477769 |
| hectd4.L | 3888.17072664961 | -0.350980516072804 | 0.0785528464928457 | -4.46808144762482 | 7.89242573648267e-06 | 0.000172581989956895 |
| LOC108703986 | 516.072394475058 | -0.706898674613818 | 0.158234308302791 | -4.46741722573289 | 7.91696289089501e-06 | 0.000172962856539032 |
| cad.L | 2106.68259540513 | -0.33631640439781 | 0.0753304269120667 | -4.46454929547117 | 8.02374742052626e-06 | 0.000175138292753158 |
| gpx3.L | 1104.43703368584 | 0.409581404737508 | 0.0917554923715003 | 4.46383528823748 | 8.05054605711331e-06 | 0.000175528469600337 |
| srsf7.L | 1885.8751576437 | 0.334884992637936 | 0.0750242780332545 | 4.46368830753024 | 8.05607325283511e-06 | 0.000175528469600337 |
| phc1.L | 1509.83333100743 | -0.331290210474615 | 0.0742465600623731 | -4.46202773833973 | 8.11877134014923e-06 | 0.000176736049227227 |
| LOC108711439 | 5994.1847223163 | 0.34824313090234 | 0.0780583664764664 | 4.46131717357077 | 8.14574238493037e-06 | 0.000177164427483884 |
| LOC108714594 | 1163.85491565212 | -0.437294573237167 | 0.0980234897901856 | -4.46111022917337 | 8.15323298753099e-06 | 0.000177168731841751 |
| map3k9.S | 340.745393778083 | -0.569424257949516 | 0.127652012546739 | -4.46075425360822 | 8.1671700278251e-06 | 0.000177312983606777 |
| LOC108719847 | 685.496812908187 | -0.533180216867824 | 0.119535644991064 | -4.46042866048604 | 8.17958836825578e-06 | 0.000177424035552148 |
| tulp1.S | 22.7951898014828 | 0.757116749024457 | 0.169761327417322 | 4.45988942560073 | 8.20019485808411e-06 | 0.000177712340662172 |
| LOC108718304 | 964.461823975316 | -0.43038913711727 | 0.0965310308566669 | -4.45855734987777 | 8.25131212835771e-06 | 0.000178660763677649 |
| cth.L | 1693.73686297231 | 0.461970369830876 | 0.103625240083036 | 4.45808732950286 | 8.26942134657554e-06 | 0.000178893430270442 |
| rpl24.L | 9204.49898319377 | 0.288640554112629 | 0.0647850751466017 | 4.45535570437275 | 8.37542127806968e-06 | 0.000181025342108029 |
| egr2.L | 109.194830768808 | -0.818060689416004 | 0.183641144158578 | -4.4546699660594 | 8.40223438150478e-06 | 0.000181443450723802 |
| erc1.L | 1156.67095252193 | -0.503169693810428 | 0.112966156308976 | -4.45416268504397 | 8.42212238775008e-06 | 0.000181550169732032 |
| lipe.L | 999.297309920261 | -0.391565709581308 | 0.0879090423977561 | -4.45421425260916 | 8.42009862330515e-06 | 0.000181550169732032 |
| tubb2b.L | 19908.3149053559 | 0.289085503982899 | 0.0649649985711915 | 4.44986547126768 | 8.59240949244399e-06 | 0.000185056734228222 |
| hmga2.L | 4537.49414706631 | 0.373723543161377 | 0.0840014887691323 | 4.44901094775249 | 8.62666195047814e-06 | 0.000185629871944124 |
| hmgb3.S | 14846.5138257461 | 0.280784201447718 | 0.0631237759491342 | 4.44815281129533 | 8.66119052614068e-06 | 0.000186207931541647 |
| gngt1.L | 170.125617713471 | 0.846833884187026 | 0.19044104762332 | 4.44669830771992 | 8.72001679263015e-06 | 0.000187306885906416 |
| rai1.S | 1002.48479013959 | -0.439483094173862 | 0.0988476245777901 | -4.44606631723257 | 8.74569597722247e-06 | 0.000187692524808118 |
| calr.S | 13727.1814137748 | -0.230932603019443 | 0.051950576044917 | -4.44523661912384 | 8.77951819450084e-06 | 0.000188252087393825 |
| clasp1.L | 4512.20316231686 | -0.435466271149934 | 0.0979865534087366 | -4.44414316047479 | 8.82428342694599e-06 | 0.00018904509838997 |
| camta2.L | 1351.56082721316 | -0.464997343325151 | 0.104641240709122 | -4.44372926175191 | 8.84128491459707e-06 | 0.000189075858904244 |
| phactr3.L | 1427.24261535139 | -0.360333677576845 | 0.0810874637246326 | -4.44376554679911 | 8.83979320302856e-06 | 0.000189075858904244 |
| atrn.S | 593.640341884469 | -0.48317908643277 | 0.108751995755798 | -4.44294454621085 | 8.87360420474393e-06 | 0.000189600123614819 |
| prrc2b.S | 6625.25336973992 | -0.330784149019283 | 0.0744563491625621 | -4.44265872205304 | 8.88540418193963e-06 | 0.000189685421086152 |
| nr2f2.L | 3765.76846273252 | -0.334903047576689 | 0.0753964502880501 | -4.44189409842507 | 8.91704469167997e-06 | 0.000190193752185841 |
| LOC108713037 | 816.910789145377 | -0.495044390568355 | 0.111483024401081 | -4.4405360657183 | 8.97350636473297e-06 | 0.000191230143530546 |
| snap91.L | 5077.53211398434 | -0.402018997881044 | 0.0905456724200043 | -4.43995816847262 | 8.99763653735779e-06 | 0.000191408656570803 |
| ier5l.L | 1812.4738285821 | -0.343876127749909 | 0.0774487279789213 | -4.44004874868313 | 8.99385025610053e-06 | 0.000191408656570803 |
| LOC108718817 | 167.312571783671 | -0.811779590907701 | 0.18284657907123 | -4.43976554022078 | 9.00569352292626e-06 | 0.000191412352096212 |
| LOC108696020 | 253.8202673027 | -0.676820529012784 | 0.15246335787421 | -4.43923404580396 | 9.02795989660485e-06 | 0.000191717882629474 |
| slc5a6.L | 1752.76328348525 | -0.389333821518367 | 0.0877387702312421 | -4.43742054387414 | 9.10433119296462e-06 | 0.000193170848909941 |
| phyhipl.S | 1737.96467114518 | -0.494962766518786 | 0.111627462980387 | -4.43405908638946 | 9.24752640226582e-06 | 0.000196037876454316 |
| atf4.S | 6872.96561044367 | 0.330275062426043 | 0.0745044949149884 | 4.43295485464193 | 9.29503344491723e-06 | 0.000196726135634879 |
| ythdf1.L | 1286.40896549844 | -0.37013527476361 | 0.0834967923199195 | -4.4329280739963 | 9.29618851193057e-06 | 0.000196726135634879 |
| LOC398642 | 1187.14506986203 | -0.468533612481509 | 0.105698621433031 | -4.43273153546629 | 9.30466954622547e-06 | 0.000196734240170584 |
| kcnab3.L | 531.547524051971 | -0.526460121788012 | 0.118820497759715 | -4.43071803025644 | 9.39198322760193e-06 | 0.000198407687418575 |
| prdx1.S | 3685.88345082541 | 0.31686810230299 | 0.0715379137412027 | 4.42937298184708 | 9.45074544111315e-06 | 0.000199475594914338 |
| nol6.L | 1289.26203481478 | -0.37105648730362 | 0.0838063615598507 | -4.42754559913246 | 9.53114274835737e-06 | 0.000200966099034914 |
| fscn2.L | 54.2932677676189 | 0.859320540871584 | 0.194091790173893 | 4.42739252444264 | 9.53790698062304e-06 | 0.000200966099034914 |
| rpl21.L | 11774.0958756473 | 0.3437967079522 | 0.0776759546195435 | 4.42603775693668 | 9.59797316434409e-06 | 0.000201881523856775 |
| LOC108698179 | 1447.53297321274 | -0.392604428309487 | 0.0887011331451881 | -4.42614896099313 | 9.59302914029354e-06 | 0.000201881523856775 |
| msmo1.S | 721.147036342445 | 0.655846187251587 | 0.148224614373599 | 4.42467797958666 | 9.65862473178384e-06 | 0.000202981513178163 |
| rplp0.L | 37859.7007517507 | 0.318797352627558 | 0.0720930428942339 | 4.42202658993405 | 9.77794197147651e-06 | 0.000205311428051038 |
| ppa2.S | 2481.42148244226 | 0.299075694610677 | 0.0676654733706339 | 4.41991579623638 | 9.87393672506197e-06 | 0.000207148030050652 |
| asap1.S | 1827.15683607393 | -0.434236657172259 | 0.0982537251509454 | -4.41954395627391 | 9.89094026402806e-06 | 0.000207325714214234 |
| smarca4.L | 3469.38636817756 | -0.316646160283009 | 0.0716533464036479 | -4.41913987518786 | 9.90944983844416e-06 | 0.000207355878014783 |
| mical3.S | 872.315439938356 | -0.4733116684941279 | 0.107101362829493 | -4.41928722881705 | 9.90269624265459e-06 | 0.000207355878014783 |
| tardbp.L | 4715.06689537773 | 0.280349166953139 | 0.0634684215209472 | 4.41714415192462 | 1.0001353362515e-05 | 0.000209098862813201 |
| ccni.S | 875.218367591604 | 0.437022144784637 | 0.0989555288302921 | 4.41634893927075 | 1.00381994322797e-05 | 0.000209688750651594 |
| ntn5.L | 408.307083147717 | 0.597985920397482 | 0.135429110410084 | 4.41549027817401 | 1.00781309452592e-05 | 0.000210342021635849 |
| LOC108708564 | 15956.6261875903 | 0.328727361440953 | 0.0744608584916469 | 4.41476727639166 | 1.01118713123316e-05 | 0.000210865065804107 |
| grin2c.S | 237.926402722512 | -0.650763360866978 | 0.147463004817268 | -4.41306185014597 | 1.01918864338836e-05 | 0.000212351362799974 |
| scube1.S | 392.950304230569 | -0.533666562781874 | 0.120977736444095 | -4.41127912017504 | 1.02761747567531e-05 | 0.000213558074884995 |
| appl1.L | 1425.99568141718 | -0.40728073049068 | 0.0923217698950348 | -4.41153512279647 | 1.02640300275475e-05 | 0.000213558074884995 |
| mark4.L | 2009.81570709388 | -0.329077005619269 | 0.0745973528819458 | -4.41137644843847 | 1.02715559012511e-05 | 0.000213558074884995 |
| zdhhc9.L | 2937.79433155224 | -0.292535131296282 | 0.0663540971492118 | -4.40869733542531 | 1.039942381446e-05 | 0.000215934702691018 |
| nnt.L | 9152.16865477075 | -0.254429251667405 | 0.0577150773820653 | -4.40836715825781 | 1.04152872229778e-05 | 0.000216079408877047 |
| prune1.L | 498.372430580624 | -0.496461149315793 | 0.112625511282559 | -4.40807010474077 | 1.04295789459587e-05 | 0.000216191289175017 |
| fry.S | 2076.29846790907 | -0.37665486811528 | 0.085496260041331 | -4.40551280176696 | 1.05533918943033e-05 | 0.000218571272532143 |
| foxq1.S | 153.360993429823 | -0.688424717291632 | 0.156299433306147 | -4.40452471726624 | 1.06016053314075e-05 | 0.000219382793800012 |
| zc3h18.L | 1251.95820403078 | -0.384461635665381 | 0.0872993759269508 | -4.40394483446348 | 1.06299984900291e-05 | 0.000219783134737675 |
| lpcat3.L | 712.375330469055 | -0.41821185635412 | 0.0949710717072419 | -4.40357099099925 | 1.0648341697143e-05 | 0.000219975181284347 |
| prex1.S | 1054.14937791001 | -0.386427365323611 | 0.0877598164135941 | -4.40323807769216 | 1.06647020415338e-05 | 0.000220125973999169 |
| MGC68655 | 2476.84458755471 | 0.370470218404102 | 0.0841882850476465 | 4.4004960808316 | 1.08003678412692e-05 | 0.000222736957840233 |
| dnah9.S | 755.542092904801 | -0.529945378704044 | 0.120463762628726 | -4.3992099129209 | 1.08645698474506e-05 | 0.000223870958332455 |
| psmd7.S | 2379.81844245165 | 0.296505907292837 | 0.0674145972914413 | 4.39824487879097 | 1.0912980832671e-05 | 0.000224677929109245 |
| LOC108710643 | 280.006349141677 | 0.820103081333073 | 0.186486635565875 | 4.39765068871854 | 1.09428907795701e-05 | 0.000225102953936389 |
| LOC121402892 | 50.8841364053735 | 0.854998621100666 | 0.194496136400267 | 4.39596712266359 | 1.10280626558949e-05 | 0.000226663074587404 |
| hnrnph3.L | 5076.42002585643 | 0.334217397981034 | 0.0760433614562724 | 4.3950897432805 | 1.10726999143729e-05 | 0.000227388141774958 |
| prpf19.L | 2170.91361436533 | 0.333493170431014 | 0.0759047332514809 | 4.3935754220506 | 1.11501480765003e-05 | 0.000228785217373732 |
| kcne3.S | 59.526910750794 | 0.834478299678816 | 0.190007280842794 | 4.39182275530397 | 1.12404318174688e-05 | 0.000230330318115255 |

| | | | | | | |
|---|---|---|---|---|---|---|
| LOC108712602 | 475.579503387247 | -0.519618952616818 | 0.118317174453865 | -4.39174578851554 | 1.1244412500399e-05 | 0.000230330318115255 |
| ppia.S | 23331.5446542454 | 0.286566098938609 | 0.0652794070086314 | 4.38983918620337 | 1.13434516679979e-05 | 0.000232087977649906 |
| mvd.L | 872.126785503272 | 0.669845722145209 | 0.152593949983246 | 4.38972660592869 | 1.13493256544039e-05 | 0.000232087977649906 |
| smox.L | 1685.2572619142 | -0.496098879596297 | 0.113066573947614 | -4.38767057562165 | 1.14571131709305e-05 | 0.000234095128153563 |
| rps14.L | 10476.5928061344 | 0.320574578598317 | 0.073102632071529 | 4.3852672539156 | 1.1584346247093e-05 | 0.000236495888846114 |
| LOC108704039 | 2622.96364006878 | -0.326412784071757 | 0.0744417840488421 | -4.38480603658819 | 1.16089170841884e-05 | 0.000236798515233646 |
| scoc.S | 3888.61092565359 | 0.33073444574831 | 0.0754625494633483 | 4.38276268295104 | 1.17183741211775e-05 | 0.000238749191129899 |
| LOC108709730 | 725.532556395625 | -0.506194421812441 | 0.115499507291571 | -4.38265438253849 | 1.1724202890342e-05 | 0.000238749191129899 |
| LOC108697549 | 645.705121798277 | 0.439796009126072 | 0.100357150289137 | 4.38230866319921 | 1.17428281583826e-05 | 0.000238928197051715 |
| LOC108704215 | 1547.82360748581 | -0.401368789861276 | 0.0916628468413324 | -4.37875108282467 | 1.19361368707245e-05 | 0.00024265816664216 |
| npdc1.1.L | 893.87105862831 | -0.438931119450166 | 0.100247293309793 | -4.37848349774136 | 1.19507988379484e-05 | 0.000242753099472506 |
| arpc1a.S | 4187.63562015042 | 0.301620167213907 | 0.0689071933919814 | 4.3771941994231 | 1.20216854865332e-05 | 0.000243988995162771 |
| LOC108709437 | 1481.93007684716 | -0.474420226440803 | 0.108401012273643 | -4.37652948519701 | 1.20583886034244e-05 | 0.000244529626654084 |
| pi4ka.S | 2692.27985350955 | -0.406741820152525 | 0.0929413216950966 | -4.37632920141681 | 1.20694684997019e-05 | 0.000244550181594461 |
| LOC108703048 | 481.304053789073 | -0.506469554349925 | 0.115739735931631 | -4.37593493948441 | 1.20913078540223e-05 | 0.000244785527504682 |
| LOC121395467 | 378.173709188038 | -0.55971744845977 | 0.127938985100803 | -4.37487797811408 | 1.21500423587989e-05 | 0.000245272623678189 |
| LOC108718028 | 639.027101669353 | -0.435476614540173 | 0.099542003382072 | -4.37480259332016 | 1.21542418179789e-05 | 0.000245272623678189 |
| rps9.L | 4777.25974213972 | 0.31666319630941 | 0.0723838296456406 | 4.37477814947971 | 1.2155603807876e-05 | 0.000245272623678189 |
| rbmx.S | 4364.20131842464 | 0.334365135659162 | 0.0764293168795169 | 4.37482826369173 | 1.2152811643601e-05 | 0.000245272623678189 |
| sult4a1.L | 1706.13717402275 | -0.396655294191937 | 0.090694984371028 | -4.37350860075396 | 1.22265425553674e-05 | 0.000246499273726221 |
| rsl24d1.L | 2908.26739005145 | 0.324841127346635 | 0.0743154136673253 | 4.37111376114778 | 1.23614362824388e-05 | 0.000249012216455695 |
| cbl.L | 1413.33233555086 | -0.319684620943174 | 0.073162881322599 | -4.36949194952534 | 1.24535930825322e-05 | 0.000250660803932922 |
| srsf9.L | 2225.77801031031 | 0.371151748445233 | 0.0849699942109108 | 4.36803311441881 | 1.25370487885063e-05 | 0.000252131674890706 |
| slc7a2.1.L | 687.614043867728 | -0.624564133549172 | 0.143007421102005 | -4.3673547060448 | 1.25760400927988e-05 | 0.00025249778348302 |
| LOC108703265 | 1116.61343563796 | -0.420341563863384 | 0.0962438978589825 | -4.36746197124386 | 1.25698673677549e-05 | 0.00025249778348302 |
| mif.L | 2842.3182126329 | 0.326819622546002 | 0.0748641411107367 | 4.36550286555188 | 1.26830639336726e-05 | 0.000254436296618201 |
| sub1.L | 3513.93047533375 | 0.308581486309081 | 0.0707024459928199 | 4.36450934583535 | 1.27408402780335e-05 | 0.000255384466761177 |
| rps13.S | 6934.85755483106 | 0.304067176081425 | 0.0696986349675883 | 4.36259872553923 | 1.28526555318522e-05 | 0.000257413366439257 |
| atp13a5l.2.L | 725.943215610621 | -0.640126394436438 | 0.146773450052856 | -4.36132280195032 | 1.29278471584049e-05 | 0.000258493101947563 |
| eef1a2.L | 2115.06380288312 | -0.376516212337228 | 0.0863291661305259 | -4.36140216816125 | 1.29231578022594e-05 | 0.000258493101947563 |
| LOC108719760 | 381.592566909241 | -0.546115974145095 | 0.125245659476511 | -4.36035848609599 | 1.29849536588757e-05 | 0.000259421434365728 |
| LOC121401019 | 302.084622438887 | -0.643227792825004 | 0.147524548412148 | -4.36014073419145 | 1.29978821306392e-05 | 0.000259466350436934 |
| cst3.L | 2323.42024357006 | 0.350458870630448 | 0.0804184332126772 | 4.35794203679165 | 1.31291141668397e-05 | 0.000261870853505094 |
| LOC108696803 | 111.879756100241 | 0.848342705720156 | 0.194717422325275 | 4.35678890768697 | 1.31984444973663e-05 | 0.000262748034131152 |
| LOC121393327 | 510.437276909584 | 0.544863734255194 | 0.125064229906272 | 4.35667124535557 | 1.3205538391131e-05 | 0.000262748034131152 |
| med25.S | 1269.64625086934 | -0.34297432193892 | 0.0787179477905599 | -4.35700283817676 | 1.31855558833779e-05 | 0.000262748034131152 |
| slc37a3.L | 1784.92878564108 | -0.346036369914387 | 0.0794317934626895 | -4.35639628452965 | 1.32221300384763e-05 | 0.000262862870012066 |
| tjp1.S | 1347.94989404452 | -0.414505993773781 | 0.0951531254791165 | -4.35619945941506 | 1.32340190428638e-05 | 0.000262884103538293 |
| trap1.L | 2093.08976677001 | -0.379606931609943 | 0.0871549911893831 | -4.35553863788567 | 1.32740098846016e-05 | 0.00026346306873898 |
| clcn5.L | 490.88192346524 | -0.463725807861945 | 0.106507281482908 | -4.35393525593237 | 1.33715212861832e-05 | 0.000265181827042069 |
| bzw1.L | 5007.37744018208 | 0.293137214115578 | 0.067334159621946 | 4.35346955782065 | 1.33999710358091e-05 | 0.000265529279236498 |
| snrnp27.S | 2837.37918563342 | 0.294638312434025 | 0.0676896168031924 | 4.3527844645758 | 1.34419287305436e-05 | 0.000266143615794479 |
| sri.S | 4598.71889703362 | 0.328655806505597 | 0.0755387965599722 | 4.35082131927623 | 1.35628543706914e-05 | 0.000268306960952291 |
| LOC108702512 | 3778.62965917848 | -0.401453995790806 | 0.0922744261976042 | -4.35065285511609 | 1.35732796167928e-05 | 0.000268306960952291 |
| oaz1.L | 12691.0117700909 | 0.267873678092316 | 0.0616311755053784 | 4.34639897577387 | 1.38390745923871e-05 | 0.000273338600119879 |
| ptpro.L | 4577.25679882097 | -0.337349657064151 | 0.0776221754383104 | -4.3460474427473 | 1.3861260107241e-05 | 0.000273554389151351 |
| npepps.L | 4094.73391119259 | -0.327955362697201 | 0.0754752380536104 | -4.34520474734048 | 1.3914581464437e-05 | 0.000274383800403435 |
| zfhx2.L | 2226.0910146757 | -0.423133779586586 | 0.0974042500513451 | -4.34409976323967 | 1.39847955859189e-05 | 0.000275544707189224 |
| rpl17.S | 12366.9599012985 | 0.330210081010168 | 0.07602026688604 | 4.34371115146408 | 1.40095694087982e-05 | 0.000275809140370619 |
| hsbp1.L | 2501.37059518157 | 0.348203614895537 | 0.0801738578678536 | 4.34310664542877 | 1.40481896075287e-05 | 0.000276345520911177 |
| rpl17.L | 10478.3937465203 | 0.33188820888948 | 0.0764429636622353 | 4.3416449727922 | 1.41419918143463e-05 | 0.000277740945139636 |
| cdc42bpa.S | 1949.90502428189 | -0.424561303741235 | 0.0977851640498159 | -4.34177625887036 | 1.41335422308624e-05 | 0.000277740945139636 |
| atoh1.S | 501.515751885783 | -0.485413841700321 | 0.111817532645485 | -4.3411246001941 | 1.41755304569735e-05 | 0.000277950231575234 |
| eef1d.L | 8666.90172386233 | 0.244666323477349 | 0.0563582146923019 | 4.34127171723146 | 1.41660409025587e-05 | 0.000277950231575234 |
| psma6.S | 2683.58419037431 | 0.349467802619733 | 0.0805082244481438 | 4.34077145552786 | 1.41983342508703e-05 | 0.000278172848621487 |
| lrrn3.S | 1719.20586424922 | 0.39988126141375 | 0.092180144798898 | 4.33804115068532 | 1.43758242880456e-05 | 0.000281423267730684 |
| atp6ap1l.L | 35.9155149729459 | 0.839697135974103 | 0.193610593527526 | 4.3370412779336 | 1.44413511915015e-05 | 0.000282251155145886 |
| eif3c.S | 6912.16578168564 | -0.275454608089267 | 0.0635105702382389 | -4.33714588069341 | 1.44344827068149e-05 | 0.000282251155145886 |
| pcdhac2.S | 1548.78032824746 | -0.434635454210707 | 0.100224110744267 | -4.33663567561828 | 1.44680135618798e-05 | 0.000282544952952016 |
| septin7.L | 6777.23612753563 | 0.239068633059101 | 0.0551492490945378 | 4.33493904240265 | 1.4580051904197e-05 | 0.000284504241735391 |
| dlgap5.S | 1297.63397612681 | 0.373020174810039 | 0.0860862254380878 | 4.33309943503459 | 1.47024663634275e-05 | 0.000286662694890134 |
| ppp1cc.L | 4041.27448132307 | 0.327915429578841 | 0.0757114769807935 | 4.33111917314764 | 1.48353356316550e-05 | 0.000289021366347575 |
| nr2f1.S | 1906.81207042403 | -0.357139760619693 | 0.0825378027584999 | -4.32698410526782 | 1.51164865332683e-05 | 0.000294208239833243 |
| agrp.L | 37.3754295227 | 0.722325274350759 | 0.166940276336535 | 4.32684844066404 | 1.51257961452096e-05 | 0.000294208239833243 |
| rpl22l1.L | 1853.05404380236 | 0.428009820783835 | 0.0989310249177973 | 4.32634576604733 | 1.51603384894603e-05 | 0.000294644210610358 |
| rab3gap2.L | 1843.55328733406 | -0.3758258976448 | 0.0868754002237408 | -4.32603356781971 | 1.51818296951155e-05 | 0.000294826035661969 |
| polr2e.L | 1349.01518321625 | 0.384757093027615 | 0.0889920465329169 | 4.32349977349154 | 1.53573295958024e-05 | 0.000297995978594588 |
| islr2.S | 1142.40434322102 | 0.554755934901457 | 0.128317056934057 | 4.32332184166716 | 1.53697262165109e-05 | 0.000297998506547419 |
| nkx6-2.S | 318.904846033883 | -0.761580625275413 | 0.176208995302746 | -4.32203034792256 | 1.54599917423797e-05 | 0.000299509600788973 |
| rnf2.S | 3060.62285771363 | 0.310120198899424 | 0.0717851088513573 | 4.32011880822773 | 1.55945218713858e-05 | 0.000301875150871272 |
| luc7l.L | 731.400720616054 | 0.405951249821614 | 0.0939806608891885 | 4.31951899444787 | 1.56369650310473e-05 | 0.000302455755146706 |
| sos1.L | 789.071451720248 | -0.403585413889203 | 0.0934768027038472 | -4.31749270637594 | 1.5781162174922e-05 | 0.000305002031724387 |
| pcbp3.L | 5317.6507858876 | -0.338537535566942 | 0.0784443906005394 | -4.31563726832768 | 1.59143121203496e-05 | 0.000307330921026846 |
| celf2.L | 5254.61930842103 | 0.338460907215592 | 0.0784305569118013 | 4.31542144468267 | 1.59298694013881e-05 | 0.000307387011308437 |
| LOC108700431 | 3439.28492050537 | -0.310653438843653 | 0.0719939585856395 | -4.31499315979575 | 1.59607845289174e-05 | 0.000307739126464698 |
| gabarapl2.S | 1084.94750931718 | 0.392171015049322 | 0.0909055800881585 | 4.31404776988389 | 1.60292287148412e-05 | 0.000308742869542821 |
| col12a1.S | 451.166618316737 | -0.668319507827382 | 0.154921508939245 | -4.31392330479735 | 1.60382605319437e-05 | 0.000308742869542821 |
| pkp3.S | 509.301274762841 | -0.482761275931824 | 0.111918990294819 | -4.31348848537791 | 1.6069851321475e-05 | 0.000309106071262007 |
| tmsb4x.L | 29817.716663834 | 0.310282012371556 | 0.0719475623353548 | 4.31261327416905 | 1.61336176816409e-05 | 0.000310087110726096 |
| slc30a3.L | 902.74115747319 | -0.820632435610571 | 0.190323076102748 | -4.31178631837341 | 1.61940897832549e-05 | 0.000311003333750509 |
| camk4.L | 760.936323507565 | -0.493048277332823 | 0.114431410274431 | -4.30867955005002 | 1.6423211955072e-05 | 0.000314971482848696 |
| mapre3.S | 1226.04211402708 | -0.455373703252433 | 0.105688664446448 | -4.30863333960635 | 1.64266431534246e-05 | 0.000314971482848696 |
| rpl36.L | 5430.90427091994 | 0.352751543211707 | 0.0818769794093828 | 4.30831163724248 | 1.64505490032744e-05 | 0.000315037085609838 |
| LOC108696466 | 768.022362962181 | -0.495715819883763 | 0.115062301573687 | -4.30823834656482 | 1.64559999028107e-05 | 0.000315037085609838 |
| mn1.L | 1835.10212771779 | -0.35374600880273 | 0.0821137725031877 | -4.30799850036116 | 1.64738501917105e-05 | 0.00031513048547828 |
| prrt1.L | 2595.55582337694 | -0.338111445737175 | 0.0784896612346086 | -4.30771951896374 | 1.64946362910727e-05 | 0.000315279853702061 |
| sec14l5.S | 1036.69451194411 | -0.391467405958777 | 0.0908981812998745 | -4.30665828909514 | 1.65739340914607e-05 | 0.000316546505360021 |
| LOC108717635 | 922.042515126691 | -0.410325845254608 | 0.0952968202110123 | -4.3057663870215 | 1.6640860156368e-05 | 0.000317575064131033 |
| map1b.S | 6314.48942181127 | -0.450657463815485 | 0.104676326887017 | -4.3052472055301 | 1.66799367341339e-05 | 0.000317821476877685 |
| naa40.S | 844.663438970405 | 0.381122422774281 | 0.0885216791876017 | 4.30541338881042 | 1.66674192580167e-05 | 0.000317821476877685 |
| sgip1.S | 1109.26318090192 | -0.438738375016279 | 0.101922252676002 | -4.30463773608862 | 1.67259204885768e-05 | 0.000318447893690819 |
| prrc2a.L | 5907.61635480662 | -0.362197699376346 | 0.0841481482257128 | -4.3042860361563 | 1.67525107835663e-05 | 0.000318704382909913 |
| capzb.S | 2275.12899266449 | -0.379930820769201 | 0.0882747146065832 | -4.303959774466 | 1.67772138356556e-05 | 0.000318924595401735 |
| LOC108703540 | 2395.8837140033 | -0.291715830787029 | 0.067785073763642 | -4.30354080315978 | 1.68089873549495e-05 | 0.000319278763722552 |
| LOC121398856 | 854.056717451474 | -0.519853655874148 | 0.12088211832094 | -4.30050087717644 | 1.70412496835041e-05 | 0.000323395638204402 |
| ntn3.S | 326.430978324398 | -0.597272491114076 | 0.138889062145895 | -4.30035657153962 | 1.70523508907483e-05 | 0.000323395638204402 |
| pde6b.L | 27.7684886824603 | 0.664840838722538 | 0.154639944947763 | 4.29928269146191 | 1.71351795348064e-05 | 0.000324712988782049 |
| LOC108717975 | 1432.97346684182 | 0.345934448995115 | 0.0804750477654225 | 4.29865478307615 | 1.71837877417971e-05 | 0.000325126899843629 |
| psmg2.L | 1083.62436058677 | 0.409253559202676 | 0.095203330704404 | 4.29873152729671 | 1.71778397087131e-05 | 0.000325126899843629 |
| yjefn3.L | 389.92027864773 | 0.574778177261062 | 0.133719776986915 | 4.29837822207338 | 1.72052387863671e-05 | 0.000325279432743971 |
| slc35e1.S | 1382.65739834819 | -0.378957060234635 | 0.0882023267092606 | -4.29645196870807 | 1.73553551137697e-05 | 0.000327862361690452 |
| rbfox2.S | 2989.90598966125 | -0.338841637068823 | 0.0788826332502135 | -4.29551630197267 | 1.74287228353699e-05 | 0.000328992535013358 |
| LOC108706545 | 544.828559881416 | -0.520255260046516 | 0.121153973164965 | -4.29416589861339 | 1.75351322253789e-05 | 0.000330744178791424 |
| foxp4.S | 3101.22405635714 | 0.266688707518073 | 0.0621490149056028 | 4.29111721116003 | 1.77776442632013e-05 | 0.000333449069033473 |
| sox14.L | 1348.8895257485 | 0.529360862683632 | 0.123359414013405 | 4.29120766272535 | 1.77704033964246e-05 | 0.000334490690334373 |
| camk2g.L | 5338.49806545006 | -0.337734013066971 | 0.0787079434996762 | -4.29097748016196 | 1.7788835593645e-05 | 0.000334490690334373 |
| LOC108704479 | 7090.95667165059 | -0.351151982779892 | 0.0818281845292125 | -4.29133292887039 | 1.77603801784815e-05 | 0.000334490690334373 |
| map3k12.L | 860.071861536555 | -0.460260305984899 | 0.107281348721477 | -4.29021737207857 | 1.78498318107196e-05 | 0.000335203891872054 |

| | | | | | | |
|---|---|---|---|---|---|---|
| bckdk.L | 828.911478169247 | -0.389482729076871 | 0.0907851071046799 | -4.29016103519906 | 1.78543605862533e-05 | 0.000335203891872054 |
| jmjd1c.S | 1878.2877773938 | -0.353814455153538 | 0.0825192009593899 | -4.28766215668594 | 1.80563437666766e-05 | 0.000338734220438332 |
| txndc17.S | 463.285569070333 | 0.502304244831563 | 0.117162616703429 | 4.28723989754372 | 1.80906890640888e-05 | 0.000339116666761554 |
| acvr2b.S | 1136.23510976368 | -0.361202473788308 | 0.0842646650425077 | -4.28652358145728 | 1.81490944613658e-05 | 0.000339949191090533 |
| kiaa1549l.S | 899.503232561239 | -0.451814863339228 | 0.105466920300003 | -4.28394857889119 | 1.83605363920998e-05 | 0.000343644738913461 |
| tra2b.S | 3757.5389324765 | 0.312821169830499 | 0.0730308125506203 | 4.28341351965202 | 1.84047653590616e-05 | 0.000344207366923049 |
| ube2d4.L | 1287.64413950248 | -0.342820079116062 | 0.0801177943844245 | -4.27895053464813 | 1.87776584046245e-05 | 0.000350911102524575 |
| alpl.L | 659.333549527385 | -0.44264102336164 | 0.103518980624584 | -4.27594070856334 | 1.90331878142345e-05 | 0.000355412962920072 |
| ctbp1.S | 1681.63332731565 | -0.326141170108378 | 0.0762828703367997 | -4.2754181727617 | 1.90778863294618e-05 | 0.000355974017271847 |
| LOC108719375 | 1340.33302380603 | -0.439950230440632 | 0.10293215439102 | -4.27417684049771 | 1.91844730213465e-05 | 0.000357688094843124 |
| slc38a3.S | 567.365740272204 | -0.458752661054363 | 0.107357665008106 | -4.27312442963178 | 1.92752820505995e-05 | 0.000359105599798515 |
| fam53a.S | 1159.79207242721 | 0.378785886342929 | 0.0886648377097155 | 4.27210939677187 | 1.93632535849486e-05 | 0.000360192099994442 |
| nasp.S | 1241.84201280123 | 0.327055340334769 | 0.0765538432877182 | 4.27222626962792 | 1.93531049259123e-05 | 0.000360192099994442 |
| caskin1.L | 2376.68515441439 | -0.410141070105793 | 0.0960085364859864 | -4.27192294682731 | 1.93794544667758e-05 | 0.000360217648673183 |
| gnao1.S | 11028.9081526785 | -0.293221689456571 | 0.0686427613778628 | -4.27170591000051 | 1.93983293432878e-05 | 0.000360292823444826 |
| col18a1.S | 1135.26293724905 | -0.349653326507721 | 0.0818660789952459 | -4.27104034783474 | 1.94563200104735e-05 | 0.000361093841355572 |
| slc39a7.L | 1746.75944802782 | 0.340420730624809 | 0.0797175998563729 | 4.27033341744038 | 1.9518095907305e-05 | 0.000361710513965184 |
| spred1.L | 884.029489354499 | -0.381929522761738 | 0.0894381636144066 | -4.27031937292837 | 1.95193250929594e-05 | 0.000361710513965184 |
| cdc20.L | 603.125769440973 | 0.451782870550204 | 0.105829277013725 | 4.26897814384211 | 1.963705071252e-05 | 0.000363614717995396 |
| rps28p9.L | 4422.15144909726 | 0.309148750403457 | 0.0724788029230697 | 4.26536777561838 | 1.99573179568161e-05 | 0.000368989355395554 |
| tp53bp2.S | 1439.00425307167 | -0.37371968964762 | 0.0876173098327492 | -4.26536366342456 | 1.99576855597991e-05 | 0.000368989355395554 |
| LOC108712551 | 1260.39655142235 | -0.433430720021226 | 0.101627516628825 | -4.26489531968245 | 1.99995946105347e-05 | 0.000369483004918882 |
| cbx3.L | 2650.13296235012 | 0.341692683727923 | 0.080130709524313 | 4.26419141620415 | 2.0062740039012e-05 | 0.000370367938075803 |
| hoxb1.S | 128.602542000148 | -0.769835643486427 | 0.180614103485781 | -4.26232297826638 | 2.0231275047406e-05 | 0.000373195593015704 |
| znf706l.L | 1418.34425849738 | 0.32310909425093 | 0.0758093800517995 | 4.26212553156553 | 2.02491634898629e-05 | 0.000373242168303152 |
| LOC108707903 | 3766.3588537816 | -0.455984967512099 | 0.107001262713575 | -4.26149146232692 | 2.03067113262878e-05 | 0.000373735791636997 |
| jmjd1c.L | 1788.76112630586 | -0.389071805571252 | 0.0912992541090213 | -4.26150037443523 | 2.03059013894 14e-05 | 0.000373735791636997 |
| extl3.S | 1011.80714405103 | -0.417867998094238 | 0.098061035357523 | -4.26130518172405 | 2.03236476437102e-05 | 0.000373764342056243 |
| tll1.L | 626.989628338666 | -0.542978314327698 | 0.127432850482784 | -4.26089750225789 | 2.03607601112441e-05 | 0.000374163620380154 |
| LOC108708041 | 1727.48266299832 | -0.382807646341205 | 0.0898535786746932 | -4.26034947063295 | 2.04107510062717e-05 | 0.000374798779248953 |
| aldh18a1.L | 2008.19663709792 | 0.433501757357866 | 0.1017798286689 | 4.25921091661581 | 2.05149826258033e-05 | 0.000376428238603674 |
| aipl1.S | 37.9777563245854 | 0.817369667808696 | 0.1920656760067 | 4.2556779785066 | 2.08416496410854e-05 | 0.000382133612362664 |
| pcdh10.S | 2040.91179690639 | -0.323562844546079 | 0.0760348822949673 | -4.25545269197444 | 2.08626474662657e-05 | 0.000382230133895519 |
| rtf2.S | 1084.85960133605 | 0.345629027658067 | 0.0812600578307951 | 4.25336920603426 | 2.10577955082922e-05 | 0.000385514758160099 |
| kcnq2.L | 1372.27794772463 | -0.472166087294123 | 0.111018786902549 | -4.25302870322826 | 2.10898531857654e-05 | 0.000385810913625741 |
| rhoa.2.S | 1467.81206345551 | -0.314101750641688 | 0.0739036586562566 | -4.25015156695623 | 2.13625910708602e-05 | 0.000390506235873195 |
| rab3il1.L | 347.810881281003 | -0.550834150372995 | 0.129633198036377 | -4.24917504710809 | 2.14559209746301e-05 | 0.000391917401622304 |
| eif3a.S | 12741.2631479958 | -0.221921488872684 | 0.0522352216152254 | -4.24850286857018 | 2.1520389168843e-05 | 0.000392799650238821 |
| slbp.L | 3377.37680635892 | 0.351363418433696 | 0.0827228376645864 | 4.24747782297263 | 2.16190557263885e-05 | 0.000394304309171834 |
| kctd8.L | 253.422672979255 | 0.658356474185925 | 0.155008520953764 | 4.24722767313294 | 2.16432028523326e-05 | 0.000394448589718357 |
| LOC108703926 | 236.015565735663 | 0.665059588558216 | 0.156600917047288 | 4.24684351214491 | 2.16803273265399e-05 | 0.000394828989558441 |
| tmem178b.S | 609.492794949056 | -0.475448541554465 | 0.111966000608352 | -4.24636531599933 | 2.17266281876097e-05 | 0.000395375809130929 |
| snrpc.S | 2596.61697737166 | 0.299301538490226 | 0.0705019610377249 | 4.24529380580028 | 2.18307180663044e-05 | 0.000396972653220658 |
| atrn.L | 833.041816928528 | -0.479107689756031 | 0.112865496623656 | -4.24494379671754 | 2.18648217805203e-05 | 0.000397295422839162 |
| LOC108697619 | 1893.93294437799 | -0.403892409909695 | 0.0951652528496272 | -4.2441163955913 | 2.19456427260137e-05 | 0.000398465952455738 |
| srsf3.S | 10928.92254825 | 0.295567125264741 | 0.0696571540529161 | 4.24316970228512 | 2.20384649951833e-05 | 0.000399852478411489 |
| psma5.S | 3165.89655902523 | 0.371023963467973 | 0.08745684786212 | 4.2423660643808 | 2.21175539736122e-05 | 0.000400688930823963 |
| synpr.L | 1049.43925685835 | 0.591181284295673 | 0.139351269772684 | 4.24238175410986 | 2.2116007307017e-05 | 0.000400688930823963 |
| eif3m.L | 2770.74424583226 | 0.301783651800236 | 0.0711475502813441 | 4.24165906776649 | 2.2187355428415e-05 | 0.000401653958850904 |
| frmpd1.L | 451.285514783768 | 0.623275644568091 | 0.146955985092793 | 4.24124028820285 | 2.22288001713902e-05 | 0.00040210459520756 |
| kcnh3.L | 52.1819205615788 | 0.825742481351525 | 0.194717446872924 | 4.24072159229994 | 2.22802353440544e-05 | 0.000402735146911055 |
| opn7a.L | 27.5339476057509 | 0.809987925491485 | 0.191071434584375 | 4.23918900935349 | 2.24328727026297e-05 | 0.000405192720771514 |
| LOC108702891 | 701.313754921509 | -0.464258830389099 | 0.109567505397572 | -4.23719449214902 | 2.26330071499097e-05 | 0.000408503919539305 |
| homer1.L | 477.575956991336 | -0.492921618802688 | 0.116362059189953 | -4.23610257702667 | 2.27432910324812e-05 | 0.000410189689935485 |
| LOC108704147 | 71.2525379610992 | 0.82406187094278 | 0.194601089187341 | 4.23462106190712 | 2.2893742593252e-05 | 0.000412596871335656 |
| inip.S | 1191.5251069506 | 0.381707528106199 | 0.0901475057348725 | 4.23425501343117 | 2.29310613867176e-05 | 0.000412963087716025 |
| LOC108698982 | 44.6041876049019 | 0.740618341950192 | 0.174932024941893 | 4.23374932175056 | 2.2982712104154e-05 | 0.000413586672487642 |
| pdcd4.L | 692.084303252349 | 0.466070569305252 | 0.11009794448119 | 4.23323588375329 | 2.30352672796833e-05 | 0.000414225598292099 |
| hnrnpab.L | 16877.0810382242 | 0.262160177209 | 0.0619360694669792 | 4.23275450743236 | 2.30846445035695e-05 | 0.000414806474533815 |
| eif3k.L | 1152.20742008934 | 0.350878539117689 | 0.082947279005559 | 4.23013923210397 | 2.33546715422607e-05 | 0.000419348403878553 |
| rprm.S | 310.058781012441 | 0.542992472196969 | 0.128413961382951 | 4.22845356026107 | 2.35303071442436e-05 | 0.000422190016072567 |
| ptma.L | 25006.5935160784 | 0.262581671377706 | 0.0621216542043063 | 4.22689438555717 | 2.36938808926389e-05 | 0.000424811175207211 |
| capzb.L | 2433.19160583829 | -0.344044576791755 | 0.0814171636502447 | -4.22570083956395 | 2.38198268813054e-05 | 0.000426754332046041 |
| LOC108699444 | 134.1982855136 | -0.810718506582783 | 0.191879398040431 | -4.2251461848549 | 2.38785719955572e-05 | 0.000427491546101744 |
| otx1.S | 51.4627086458308 | 0.819776782507345 | 0.194154997633432 | 4.22228009837326 | 2.41843306628137e-05 | 0.000432646634110748 |
| rps16.L | 8467.1483051529 | 0.328450669065693 | 0.0778143016310082 | 4.22095504529767 | 2.43269452062603e-05 | 0.000434877709228027 |
| dhcr24.S | 733.840904410896 | 0.551113442294874 | 0.13058425510579 | 4.22036670384498 | 2.43905241625999e-05 | 0.000435693672063383 |
| rpl24.S | 7820.73425932264 | 0.280629089994661 | 0.066516662006449 | 4.21892923561698 | 2.45465293952262e-05 | 0.00043811189806764 |
| pomp.L | 1860.50557777057 | 0.330070777333812 | 0.078238304160482 | 4.21878747086302 | 2.45619661302431e-05 | 0.00043811189806764 |
| endod1.S | 1832.95246663858 | -0.435040080467642 | 0.103136229697031 | -4.2181111501322 | 2.46357377698976e-05 | 0.000438783440895815 |
| LOC108717042 | 169.80837578663 | -0.643159381881988 | 0.152473176121163 | -4.2181805235755 | 2.46281609701611e-05 | 0.000438783440895815 |
| copb1.L | 1500.02000883299 | -0.37023029484645 | 0.0877848182139542 | -4.21747521244623 | 2.47052967571253e-05 | 0.000439699984921321 |
| mab21l2.L | 3091.61780903308 | -0.753566093746151 | 0.178697028475429 | -4.21700405527318 | 2.47569524678869e-05 | 0.000439974691481232 |
| fbn3.L | 3295.19552912085 | -0.428440394818273 | 0.101595181511538 | -4.2171330219004 | 2.4742802900142e-05 | 0.000439974691481232 |
| scaf4.L | 1009.28197014701 | -0.396953217048624 | 0.0941484608816903 | -4.21624754490089 | 2.48401081053059e-05 | 0.00044105112018097 |
| pomp.S | 1010.80854848487 | 0.410162076921986 | 0.0972841835969605 | 4.21612292725043 | 2.48538315439099e-05 | 0.00044105112018097 |
| hnrnpc.S | 5129.08151598266 | 0.272632438930419 | 0.0646733129402671 | 4.21553228890911 | 2.4918973370868e-05 | 0.000441562027040 02 |
| nae1.S | 1541.88527334139 | 0.350833661822363 | 0.0832217598659157 | 4.21564819570766 | 2.49061771510892e-05 | 0.000441562027040 02 |
| stab2.L | 377.789680092082 | -0.679051106167793 | 0.161204590686628 | -4.21235588437941 | 2.52720954298965e-05 | 0.000447492920097599 |
| vsx2.L | 449.500642799965 | -0.532961643928567 | 0.126552978874077 | -4.21137178018445 | 2.53824609659442e-05 | 0.000449119815518316 |
| ogn.L | 712.704975759048 | -0.609219063393957 | 0.144692977501059 | -4.21042592332787 | 2.54889690856257e-05 | 0.000450676138985583 |
| prim2.S | 584.575474678754 | 0.423384451868177 | 0.100616736341536 | 4.20789291386903 | 2.57762958942964e-05 | 0.000455424969058935 |
| khdrbs3.L | 2069.75927244432 | -0.29439873852094 | 0.069977868882533 | -4.20702635307638 | 2.58752977338044e-05 | 0.000456841920890293 |
| LOC108714595 | 1414.53827969064 | -0.349322532048518 | 0.0830388668076828 | -4.20673529731018 | 2.59086309949542e-05 | 0.000457098243566752 |
| LOC108700962 | 441.864258002973 | -0.509869132893554 | 0.121247672352839 | -4.20518697802132 | 2.60866403916851e-05 | 0.000459904819793612 |
| h3-5.L | 7630.82879529395 | 0.271841997730631 | 0.0646759903564966 | 4.20313622152869 | 2.63242059201376e-05 | 0.000463756532722133 |
| epha10.L | 898.303813871738 | -0.594346681189774 | 0.141462032729871 | -4.20145723711401 | 2.65202346750105e-05 | 0.000466871435648337 |
| dynll1.L | 3450.8388481867 | 0.336151458749045 | 0.0800308711173887 | 4.2002723955857 | 2.66594047311355e-05 | 0.000468981591990011 |
| vrk3.L | 1302.16891231184 | 0.321927639265681 | 0.0766980279474501 | 4.19733919998895 | 2.70069283321183e-05 | 0.000474751314689205 |
| klhdc2.L | 629.812312878133 | 0.405179943616926 | 0.0965452133142648 | 4.19678956322798 | 2.70725264529208e-05 | 0.000475560345370396 |
| gucy2d.L | 183.160943117396 | 0.725155740907382 | 0.17285185622113 | 4.19524416318497 | 2.72577799059887e-05 | 0.000478468573002955 |
| tcea1.S | 2159.81993108752 | 0.3035570887731 | 0.0723661264151332 | 4.19474005049992 | 2.73184702457646e-05 | 0.00047918766509069 |
| wnk1.L | 1913.32735138123 | -0.377284688046957 | 0.0899703058984184 | -4.19343564834527 | 2.7476104697184e-05 | 0.000481257741538131 |
| xrcc6.L | 1007.011135356 | 0.347506305145921 | 0.082867988151338 | 4.19349272135443 | 2.74691894804689e-05 | 0.000481257741538131 |
| kif5c.L | 5976.57008091193 | -0.316693152536236 | 0.0755356379576199 | -4.19263226073393 | 2.75736224851285e-05 | 0.000482617856378754 |
| spcs1.L | 1486.32924427488 | 0.343574693486753 | 0.0819531069935227 | 4.19233273869541 | 2.76100635841731e-05 | 0.000482626997097357 |
| XB5969698.S | 1373.59261265504 | -0.434776924036187 | 0.103708412340129 | -4.19230141726847 | 2.76138769229162e-05 | 0.000482626997097357 |
| dusp8.S | 455.010359741033 | -0.496564909016812 | 0.118460020320041 | -4.1918354198763 | 2.7670670617074e-05 | 0.000483271942466712 |
| dusp14.S | 144.030859166281 | -0.657453352531639 | 0.156932323237729 | -4.18940686639614 | 2.79684539933647e-05 | 0.000488121854392818 |
| letm1.L | 959.250471183594 | -0.396072292680168 | 0.0945561810416438 | -4.18875094485608 | 2.80494025915098e-05 | 0.000489183192073324 |
| LOC108713907 | 1378.5507112902 | -0.455776597279439 | 0.108823516612526 | -4.18821783178736 | 2.81153591898454e-05 | 0.000489981733255455 |
| LOC108708999 | 3561.16995886473 | -0.405077104351247 | 0.0967298652457961 | -4.18771496602 34 | 2.81777087167928e-05 | 0.000490716312233522 |
| mab21l2.S | 2595.20765331129 | -0.802825279577633 | 0.191738648417778 | -4.1870811451031 | 2.82564824352941e-05 | 0.000491383668062301 |
| hmgb1.L | 2499.86677921285 | 0.315588703186757 | 0.0753707740637562 | 4.18714955640232 | 2.82479699564614e-05 | 0.000491383668062301 |
| clns1a.L | 1562.37430258044 | 0.378913379730882 | 0.0905113868386759 | 4.18636143987323 | 2.83461840184003e-05 | 0.000492590983221043 |
| LOC108702499 | 14912.0793096108 | 0.25910849177246 | 0.0619008812240717 | 4.18586111616936 | 2.84087020849247e-05 | 0.000493324523553367 |
| emc3.L | 1426.09741122096 | 0.365651073551375 | 0.0873966414006243 | 4.1838115022663 | 2.86661831190911e-05 | 0.000497440180496571 |
| zeb2.S | 1219.28270596899 | 0.345904309586234 | 0.0826943595854411 | 4.18292506671921 | 2.87782266856361e-05 | 0.000499028007923514 |

| | | | | | | |
|---|---|---|---|---|---|---|
| ncapg2.L | 853.970347935608 | 0.385917121691994 | 0.0922712808872448 | 4.18241860285415 | 2.88424293841401e-05 | 0.000499784578786233 |
| znf638.S | 4712.5411880319 | -0.262835126489687 | 0.0628454691029916 | -4.18224464295033 | 2.88645130863523e-05 | 0.00049981074905192 |
| jazf1.S | 362.130590456374 | -0.547236006096632 | 0.130856329540816 | -4.18196053654357 | 2.89006141303994e-05 | 0.000500079429974305 |
| gba2.S | 745.969568687221 | -0.478092789151306 | 0.114338271501741 | -4.18138898613688 | 2.89733703886168e-05 | 0.000500842084022937 |
| ret.S | 466.23873930068 | -0.502524282392958 | 0.120184014705751 | -4.18129052872212 | 2.89859212207232e-05 | 0.000500842084022937 |
| crata.L | 1079.7233282906 | -0.365714809936152 | 0.0874729419445674 | -4.18089070524128 | 2.90369417361045e-05 | 0.000501010981915429 |
| psmd12.S | 3718.98073498046 | 0.315397344955512 | 0.0754376264120976 | 4.18090228916498 | 2.90354623393365e-05 | 0.000501010981915429 |
| wdr11.L | 783.005007933624 | -0.379303767896401 | 0.0907656452277357 | -4.17893539945326 | 2.9287685576344e-05 | 0.000504978732002626 |
| pmpca.S | 927.782489173224 | -0.378822397553825 | 0.0906609548749453 | -4.17845144115634 | 2.93500642517173e-05 | 0.000505695362362568 |
| LOC108698339 | 983.448146862338 | 0.347320436291407 | 0.0831259913871616 | 4.17824113126967 | 2.93772109982655e-05 | 0.000505804368527189 |
| mcm4.L | 2443.25042803731 | 0.33811427847265 | 0.0809299296780266 | 4.17786447878814 | 2.94258888611318e-05 | 0.000506283671382675 |
| mmp1.S | 20.9102195987632 | 0.688709350270575 | 0.164864469548671 | 4.17742738721065 | 2.94824738660403e-05 | 0.000506898244940965 |
| LOC108719591 | 897.161660145771 | -0.384819237611069 | 0.0921538139919364 | -4.17583625616126 | 2.96893332728144e-05 | 0.0005100938207424 |
| srsf12.L | 970.112207471647 | 0.360052711554789 | 0.086239665933635 | 4.1750244235856 | 2.97954085757947e-05 | 0.00051155452716633 |
| hmg20a.L | 1294.45092581919 | 0.379484696166509 | 0.0909196962348301 | 4.17384474301767 | 2.99501896678481e-05 | 0.000513848804936936 |
| relt.S | 164.928035335992 | -0.722073206078801 | 0.173010077727807 | -4.17359043797913 | 2.99836559755858e-05 | 0.000514059942322429 |
| synj1.S | 796.461953208627 | -0.397434885904719 | 0.0952964229922798 | -4.17051210764665 | 3.0391590679304e-05 | 0.000520686392075467 |
| znf593.L | 2038.42430500115 | 0.313666387653158 | 0.0752156970323553 | 4.17022509966541 | 3.042989209512e-05 | 0.000520975192782836 |
| apbb2.L | 961.389014008914 | -0.440762882394412 | 0.105716621607374 | -4.16928649149781 | 3.05554705793794e-05 | 0.00052275676215172 |
| dhx32.L | 394.220717421469 | 0.506964781037517 | 0.121640492753241 | 4.16773041248643 | 3.0764747573116e-05 | 0.000525966768150092 |
| npepps.S | 2524.90308591446 | -0.287915553229797 | 0.0690987045317731 | -4.16672866996236 | 3.09001917900779e-05 | 0.000527910871552851 |
| dpysl2.L | 8010.3489146861 | -0.419793297748435 | 0.100760306889721 | -4.16625664119785 | 3.0964210378111e-05 | 0.000528632836912037 |
| htt.L | 2707.58340652395 | -0.337730390152948 | 0.0810937619441977 | -4.16469013221199 | 3.11775715083476e-05 | 0.000531155625682887 |
| rfc5.S | 704.713327989632 | 0.437195200536844 | 0.104974276678481 | 4.16478412016975 | 3.11647308967115e-05 | 0.000531155625682887 |
| slc5a7.S | 380.219180841542 | -0.59928739776189 | 0.14388948760613 | -4.16491439181663 | 3.11469415258372e-05 | 0.000531155625682887 |
| MGC130950 | 770.179300161819 | -0.439957959512899 | 0.105651084143188 | -4.16425409242966 | 3.12372089475687e-05 | 0.000531205351495948 |
| arhgap23.S | 1226.5291311431 | -0.398904608827689 | 0.095786820123314 | -4.1645041386085 | 3.12029967280484e-05 | 0.000531205351495948 |
| LOC108702718 | 1050.47013441146 | -0.439975245251467 | 0.105656880450315 | -4.16418924518943 | 3.12460873996752e-05 | 0.000531205351495948 |
| kars1.L | 3139.89311409937 | 0.355940837055028 | 0.0855027052380092 | 4.16291901015547 | 3.14204837816922e-05 | 0.000532723027436554 |
| acat2.L | 682.310833670678 | 0.650026215291598 | 0.156145488621111 | 4.1629522635066 | 3.14159065109309e-05 | 0.000532723027436554 |
| lhx6.S | 31.8764083938923 | 0.695839595150153 | 0.167152600244876 | 4.16290021292374 | 3.14230714710045e-05 | 0.000532723027436554 |
| LOC108700953 | 654.386651506498 | -0.491129009744402 | 0.117973409644605 | -4.16304836169378 | 3.14026823096252e-05 | 0.000532723027436554 |
| calb1.L | 2737.27151277242 | -0.347795968599371 | 0.0835673993892877 | -4.16186181622344 | 3.15663355986199e-05 | 0.000534778631124736 |
| grik5.S | 1035.11975097876 | -0.395276553646034 | 0.0949964366965982 | -4.16096189911288 | 3.16909959452622e-05 | 0.000536516414978536 |
| phox2b.S | 706.495709616696 | -0.65289053458562 | 0.156937319507939 | -4.1601993498595 | 3.17969934820598e-05 | 0.000537936044326714 |
| helz.L | 1874.30003172758 | -0.291567986550524 | 0.0700996376795092 | -4.15933657009118 | 3.19173296417326e-05 | 0.000539596107387788 |
| XB22065601.L | 1992.15240830086 | -0.318951344681333 | 0.0767069394122375 | -4.15805072038174 | 3.20974767279267e-05 | 0.000542264325193498 |
| rars1.S | 2934.80006869646 | 0.311629752519289 | 0.074956891266866 | 4.15745300068284 | 3.21815455534674e-05 | 0.000543306787822055 |
| mcm2.L | 2924.28238206636 | 0.345790716132381 | 0.0832053284414773 | 4.15587225733498 | 3.24048852698463e-05 | 0.000546697418573366 |
| LOC108713150 | 284.198182202754 | -0.70420364515029 | 0.169478607075582 | -4.15511820224151 | 3.25119420600203e-05 | 0.00054812291492445 |
| cga.S | 22.2887293809742 | 0.667032008528309 | 0.160608823195736 | 4.15314672790664 | 3.279343158221e-05 | 0.000551887702887097 |
| irs4.L | 922.156699051441 | -0.378035168364268 | 0.0910225776389075 | -4.15320218532987 | 3.2785481752788e-05 | 0.000551887702887097 |
| MGC84235 | 23.1861486645131 | 0.641945272772158 | 0.154576888634852 | 4.1529188382656 | 3.28261188224193e-05 | 0.000551887702887097 |
| vapb.S | 2052.5119824908 | -0.304271797071539 | 0.0732643154684197 | -4.15306954178388 | 3.28044992718955e-05 | 0.000551887702887097 |
| ssbp3.L | 1884.85927498563 | -0.293980221145779 | 0.0707938252065679 | -4.15262518006309 | 3.28682851854737e-05 | 0.000552214467701174 |
| ppp1r9b.S | 2730.68133869804 | -0.274534262044323 | 0.0661152725122878 | -4.15235771724755 | 3.29067349244978e-05 | 0.000552478381655667 |
| apex1.S | 1576.68797907746 | 0.31363400163822 | 0.0755491724865311 | 4.15138897377247 | 3.30463568542557e-05 | 0.000554439360094813 |
| LOC108703350 | 667.459264838394 | -0.426793865771037 | 0.102861860361397 | -4.14919450485855 | 3.33647228862335e-05 | 0.000559394463628817 |
| kmt2c.S | 1748.14492948857 | -0.354829996393308 | 0.0855711480949322 | -4.14660787301418 | 3.37437236307669e-05 | 0.000565358635783346 |
| LOC108718251 | 1034.35203202416 | -0.493387923181236 | 0.11900435818441 | -4.14596516218909 | 3.38385277781798e-05 | 0.000566556301752654 |
| gtf2ird1.L | 614.341926046374 | 0.443700745430513 | 0.107045521132078 | 4.14497253820693 | 3.39854437085041e-05 | 0.000568624221387327 |
| gabbr1.S | 5181.43646132505 | 0.275565340037663 | 0.0664971291879479 | 4.1440185975368 | 3.41272049525494e-05 | 0.000570603108821221 |
| lamtor1.S | 993.658980115515 | -0.352119772278058 | 0.0849786835186742 | -4.1436247032549 | 3.4185903656594e-05 | 0.000570798861466182 |
| nmt1.L | 1183.27029565283 | -0.353759372267168 | 0.0853739428710198 | -4.14364571168531 | 3.4182770530163e-05 | 0.000570798861466182 |
| rlbp1.L | 565.563616082995 | 0.567287025276271 | 0.136932898843922 | 4.14281031122312 | 3.43075697774173e-05 | 0.000572436881986659 |
| LOC108704339 | 1612.16961485956 | -0.35290117365108 | 0.0852305374735425 | -4.14054849484703 | 3.46476347538606e-05 | 0.000577714233843713 |
| fryl.L | 3455.16017094444 | -0.342640568128518 | 0.0827557288897984 | -4.14038487395591 | 3.46723589873202e-05 | 0.000577729965183784 |
| sltm.L | 2635.8940034355 | 0.335637154430044 | 0.0810677643271564 | 4.14020489174402 | 3.46995748805573e-05 | 0.000577787163912446 |
| LOC108703993 | 4249.55619605892 | -0.28634934318265 | 0.0691715391697694 | -4.13969887933036 | 3.47761999647601e-05 | 0.000578666439687591 |
| arhgef11.L | 1914.78282955332 | -0.334248108132204 | 0.0807633198071498 | -4.13861278771522 | 3.49412089732409e-05 | 0.000581014189456477 |
| npas3.S | 1170.52066983189 | -0.462513295120195 | 0.11183012879905 | -4.13585587432609 | 3.53634102126812e-05 | 0.000587632481331654 |
| LOC108716640 | 930.28866488752 | -0.436576286040199 | 0.105613389765521 | -4.13372098944527 | 3.56936758522825e-05 | 0.000592715079395319 |
| pimreg.L | 1214.54985235994 | 0.366902187388411 | 0.0887666190788716 | 4.13333515679366 | 3.5753675614003e-05 | 0.000593305871152042 |
| rab2b.L | 1029.93209138506 | 0.371031082550621 | 0.0897845299743439 | 4.13246115624422 | 3.58899433852744e-05 | 0.000595160603823792 |
| ep400.L | 3974.76944696616 | -0.384535860346947 | 0.093086698307412 | -4.13094316737979 | 3.61277899345126e-05 | 0.000598696131424999 |
| natd1.S | 905.370298526327 | 0.371550737407058 | 0.0899631851539548 | 4.13003093177748 | 3.62714430024796e-05 | 0.000600666964077872 |
| tcp1.L | 9397.19143416421 | 0.30008164988632 | 0.0726613622232851 | 4.12986545674966 | 3.62975590035725e-05 | 0.00060068998530844 |
| sult1c2.S | 539.585281922507 | 0.548562248055345 | 0.132869341270851 | 4.12858408725844 | 3.65003956820092e-05 | 0.000603635543021601 |
| rspo2.S | 26.8624021592287 | 0.771111366574236 | 0.186796973148593 | 4.12807206442704 | 3.65817478080097e-05 | 0.000604569374998495 |
| maea.L | 1510.19169423945 | -0.318421280284145 | 0.0771513940850186 | -4.1272265272777 | 3.67164669463294e-05 | 0.000606383309309399 |
| LOC121399028 | 177.193960112518 | -0.786190800395236 | 0.190507987714242 | -4.12681279051935 | 3.67825589872819e-05 | 0.000607062152199067 |
| glis2.S | 534.162922981615 | -0.465065410421329 | 0.11276277255452 | -4.12428144400648 | 3.71893930336492e-05 | 0.000613359887548862 |
| LOC108701478 | 1138.38078613203 | 0.375577491646146 | 0.0910827582670063 | 4.12347516469749 | 3.73198715314054e-05 | 0.000615094273394819 |
| LOC108713086 | 28.8294058102937 | 0.747579473668189 | 0.181308949679086 | 4.12323536698764 | 3.73587612663563e-05 | 0.000615317794037194 |
| ezh2.S | 812.55512002941 | 0.370548601149974 | 0.0898782228504367 | 4.12278513524452 | 3.74318825681857e-05 | 0.000616104441132454 |
| proca1.L | 1253.91112127919 | -0.367016572371253 | 0.0890465964106593 | -4.12162381455513 | 3.76211182288926e-05 | 0.000618799895905698 |
| gphn.S | 4911.06217096329 | -0.303361645499716 | 0.0736371354839432 | -4.11968286797176 | 3.7939421788648e-05 | 0.000623613202255354 |
| lmo3.L | 506.471049094262 | 0.523157774567626 | 0.127015830266023 | 4.11883915156024 | 3.80785819068261e-05 | 0.000625430907545936 |
| rngtt.S | 550.045215287478 | 0.557862340898502 | 0.135446202039194 | 4.11870050617642 | 3.81014959729968e-05 | 0.000625430907545936 |
| cnnm4.L | 259.866414836816 | -0.587090958316181 | 0.142612445372763 | -4.11668811078607 | 3.84355639304064e-05 | 0.000630488582123763 |
| fzd7.S | 527.323920046147 | -0.56591955466851 | 0.137534459134661 | -4.11474737479728 | 3.8760368187289e-05 | 0.000635387574049932 |
| znf852.L | 1209.75655905956 | 0.359504868193323 | 0.0873799518995682 | 4.11427175659844 | 3.88403648583817e-05 | 0.000636269604767043 |
| plrg1.S | 1948.0045664745 | 0.334859832803598 | 0.0813945132799894 | 4.11403446386751 | 3.88803349149115e-05 | 0.000636495186268774 |
| cdr2l.S | 925.986896565875 | -0.39888941477574 | 0.0969629412002933 | -4.11383369602792 | 3.89141831331024e-05 | 0.000636620313155279 |
| LOC108702158 | 797.830567418936 | 0.465395154729682 | 0.113159880127087 | 4.11272223165143 | 3.91020757522288e-05 | 0.000639263679895455 |
| lmnb1.L | 6240.56531269916 | 0.249776520743435 | 0.0607600977075924 | 4.11086436933468 | 3.94180706144765e-05 | 0.00064399637357639 |
| fgd4.S | 944.598517316492 | -0.443661048163965 | 0.107993034129168 | -4.10823764459949 | 3.98689756512488e-05 | 0.000650925332306074 |
| LOC108716094 | 881.440936992518 | -0.41984504656679 | 0.102285618875916 | -4.10463417223989 | 4.04955188833442e-05 | 0.000660710635159143 |
| ppib.L | 6157.88130245977 | 0.24800516571087 | 0.0604238741288797 | 4.10442344663126 | 4.0532445862859e-05 | 0.000660869288451206 |
| kpna1.S | 547.045748138477 | -0.444765664306967 | 0.108419210146837 | -4.10227729665806 | 4.09103556978104e-05 | 0.000666583622617442 |
| erfl.L | 224.319449380434 | -0.694670269117134 | 0.169363698747331 | -4.10164795794578 | 4.10218067047495e-05 | 0.000667951589869427 |
| fst.L | 453.48104633733 | 0.603867479432497 | 0.147238050932429 | 4.10130041526853 | 4.10834772372694e-05 | 0.000668507699934509 |
| otub1.L | 1037.36234366299 | 0.3401837775664 | 0.0829763433265632 | 4.09976824632494 | 4.13564069214493e-05 | 0.000672498359939551 |
| ptpre.L | 1388.01050467192 | -0.369235751303091 | 0.0900669494473306 | -4.09956985962995 | 4.13918716595999e-05 | 0.000672624836186167 |
| pitx2.S | 340.644066478001 | 0.67488538489625 | 0.164666587316064 | 4.09849621526962 | 4.15843036604258e-05 | 0.000675300182571112 |
| LOC108712511 | 125.633525230048 | -0.68686120928067 | 0.167677625115993 | -4.09631999979439 | 4.19769593298939e-05 | 0.000681221275858678 |
| bmpr1b.L | 268.347904456978 | 0.55207585308697 | 0.134781954650315 | 4.09606652848523 | 4.20229213303372e-05 | 0.00068151191642137 |
| ubr7.L | 1418.88801961678 | 0.3230394933174 | 0.0788736789293413 | 4.09565646870351 | 4.20973786661111e-05 | 0.000682263987534692 |
| tspan18.L | 435.545687942325 | -0.475710736368782 | 0.116156853207425 | -4.09541687152364 | 4.21409418658795e-05 | 0.000682514694459784 |
| LOC108713695 | 952.707718613064 | -0.473655472522723 | 0.115671736690742 | -4.09482459651384 | 4.22488120294061e-05 | 0.000683805889035571 |
| pex5l.L | 146.679055598784 | -0.682884045616096 | 0.166812232534329 | -4.09372883056141 | 4.24490732683818e-05 | 0.000686658973767115 |
| prdx2.L | 3721.58570022773 | 0.32052197962159 | 0.0783257131264802 | 4.09216803559785 | 4.27358782992018e-05 | 0.000690768747438995 |
| arhgap44.L | 1110.52047343612 | -0.426935311231067 | 0.104360457046796 | -4.09096820110346 | 4.29576035357041e-05 | 0.000693890970941752 |
| mtmr14.S | 576.133762445333 | -0.447184296324231 | 0.109367409128537 | -4.08882591154194 | 4.33562075135888e-05 | 0.000699864256036629 |
| chat.S | 204.073318037494 | -0.643222697962477 | 0.157320085526364 | -4.08862413092628 | 4.33939319817446e-05 | 0.000700008090016271 |
| ptk2.L | 1823.83432216052 | -0.344540419379093 | 0.0842770591046543 | -4.08818749775365 | 4.34756705874638e-05 | 0.000700861274885101 |
| rnf44.L | 678.69012318082 | -0.419224027952722 | 0.102551624161117 | -4.08793162840684 | 4.35236376905583e-05 | 0.00070116926661434 |
| myef2.L | 8338.59959660308 | 0.260512128686143 | 0.0637332749161675 | 4.08753714647194 | 4.35976884772111e-05 | 0.000701431949579713 |

| | | | | | | |
|---|---|---|---|---|---|---|
| LOC108699878 | 1704.35894599808 | -0.324484179901017 | 0.079381762257773 | -4.08764142634342 | 4.35781018077961e-05 | 0.000701431949579713 |
| ube2r2 | 708.697175706175 | -0.422454398960215 | 0.103369213959397 | -4.08684929273191 | 4.37270959664363e-05 | 0.000703048358311452 |
| LOC108703378 | 21.2120552185113 | 0.643001614171903 | 0.157345837320562 | 4.08654988985767 | 4.37835370475657e-05 | 0.000703490244069816 |
| adcyap1r1.S | 5149.13282933889 | 0.326674938785982 | 0.0799502462467904 | 4.08597789402176 | 4.38915573888394e-05 | 0.000704759745673803 |
| LOC108717735 | 626.753728647867 | -0.561249988659027 | 0.137391842507505 | -4.08503138480268 | 4.40708593186389e-05 | 0.000707171371391686 |
| LOC108718965 | 686.022179136094 | 0.437121257194916 | 0.107052592654446 | 4.08323840045514 | 4.44124194897148e-05 | 0.000712181728767744 |
| dazap2.S | 534.55563032397 | -0.4624040370277 | 0.113360059921697 | -4.07907368209848 | 4.52155055286397e-05 | 0.000724581458649587 |
| slco3a1.S | 423.805801852242 | -0.519218126680613 | 0.127321015216698 | -4.078023771621 | 4.54201235571899e-05 | 0.000726901503095107 |
| cpne4.L | 1162.04476481356 | -0.482764034928396 | 0.118379782403394 | -4.07809530586326 | 4.54061543506154e-05 | 0.000726901503095107 |
| nefh.S | 1764.77627828041 | -0.766841088101758 | 0.188079685840234 | -4.07721378667742 | 4.55785820301628e-05 | 0.000728714887167712 |
| nmrk2.S | 6154.51165200164 | 0.26398558487641 | 0.0647477669448291 | 4.07713805946929 | 4.55934234170957e-05 | 0.000728714887167712 |
| LOC108704363 | 110.23938355695 | -0.740998560592737 | 0.181766354173256 | -4.07665414186847 | 4.56883722188029e-05 | 0.000729752343644705 |
| snrnp40.S | 1115.78818459712 | 0.350985775324392 | 0.0861223761410422 | 4.07543069584593 | 4.59292598982938e-05 | 0.000732674766852232 |
| fen1.L | 1576.5930102747 | 0.370630794699821 | 0.0909430012868595 | 4.0754185528884 | 4.59316567842245e-05 | 0.000732674766852232 |
| h1-0.L | 7054.51130424664 | 0.414283805145834 | 0.101658763852093 | 4.07523945253348 | 4.59670229996454e-05 | 0.000732757780021906 |
| ptpra.S | 1055.18137347209 | -0.384504440158625 | 0.0943618459808336 | -4.07478717867308 | 4.6056446648911e-05 | 0.000733701845828619 |
| sec31b.L | 1245.37688159921 | -0.311884866913871 | 0.076574185764944 | -4.07297660168726 | 4.64160890038973e-05 | 0.000738946570288781 |
| thdl20.S | 351.056584475628 | -0.745719592535569 | 0.183137589262588 | -4.07190897039894 | 4.66294034325292e-05 | 0.000741856402743854 |
| LOC108719641 | 323.690772018373 | -0.582564187637019 | 0.143100086008129 | -4.07102611806903 | 4.68065005733391e-05 | 0.000744186600084228 |
| dctn2.S | 1844.82008399872 | -0.289118945047112 | 0.0710444677504702 | -4.0695490331856 | 4.71042256855064e-05 | 0.000747941214904374 |
| strbp.L | 2075.89023880277 | -0.287600993140813 | 0.0706700238366281 | -4.06963203812803 | 4.70874474839052e-05 | 0.000747941214904374 |
| bmpr2.L | 701.675281423155 | -0.406565047618089 | 0.099937909613608 | -4.06817642264081 | 4.7382501541928e-05 | 0.000751868381750228 |
| ankrd10.L | 1489.01815975365 | 0.330727110225135 | 0.0813126103803468 | 4.06735325158214 | 4.75501338615109e-05 | 0.000754035869472289 |
| LOC108707872 | 4361.05622479722 | -0.32523738559494 | 0.0799684940342968 | -4.06706903165455 | 4.76081434940371e-05 | 0.000754406932421222 |
| LOC108712404 | 41.478990720242 | 0.791682382740397 | 0.194663178597951 | 4.06693442715996 | 4.76356398425189e-05 | 0.000754406932421222 |
| LOC108699457 | 587.885424088088 | -0.43659839899836 | 0.107371782756522 | -4.06623032411036 | 4.77797161195724e-05 | 0.000756195715575826 |
| hdac1.S | 6055.99689422221 | 0.302469261917353 | 0.0744063717940877 | 4.06509892397934 | 4.80120932317985e-05 | 0.000758963882545183 |
| galnt18.S | 718.138806033059 | -0.432365959934495 | 0.106361138063786 | -4.06507459214284 | 4.80171024727071e-05 | 0.000758963882545183 |
| LOC108714648 | 101.697614907658 | 0.705932710915757 | 0.173666419687284 | 4.06487743679468 | 4.80577094919615e-05 | 0.000759111829907485 |
| zbtb47.L | 1874.07564883579 | -0.323911417449825 | 0.0797016184445149 | -4.06405069020423 | 4.8228344792644e-05 | 0.00076131215620045 |
| pex5l.S | 261.512121663815 | -0.591190867341011 | 0.1455092704144 | -4.06290860820991 | 4.84650083459246e-05 | 0.000764551242049928 |
| syt5.L | 561.054024653881 | 0.576562872493218 | 0.141950660345725 | 4.06171321140023 | 4.87138989236401e-05 | 0.000767256494265418 |
| pnpla6.L | 2285.25660455739 | -0.300067960240048 | 0.0738787014793392 | -4.06163013468725 | 4.87312410739911e-05 | 0.000767256494265418 |
| sumo2.S | 2310.67870496975 | 0.282416965003312 | 0.0695306347688902 | 4.0617630766931 | 4.87034924218834e-05 | 0.000767256494265418 |
| nelfb.L | 1149.89680428545 | -0.329396760616084 | 0.0811322463309166 | -4.05999803422874 | 4.90731290816373e-05 | 0.000772138988283224 |
| jazf1.L | 730.757019665284 | -0.429001660063797 | 0.10572102192911 | -4.057865240382 | 4.95233290713204e-05 | 0.000778718288970005 |
| LOC108704435 | 460.406674165201 | -0.484897572364805 | 0.119502476083489 | -4.05763619513656 | 4.95719091274402e-05 | 0.000778977982109983 |
| pafah1b1.S | 2625.62485465159 | -0.315578956350094 | 0.0777914593992051 | -4.05673011905621 | 4.97645292835345e-05 | 0.000781499337048602 |
| cd93.L | 306.619242228139 | -0.572070346883444 | 0.141026486822716 | -4.05647449477056 | 4.98189999083679e-05 | 0.000781849343523185 |
| LOC108710741 | 1559.26143303167 | -0.330810768435959 | 0.0815622927591687 | -4.05592777305504 | 4.99356898359953e-05 | 0.000783174724903596 |
| npy.S | 607.674828861692 | 0.578305203478646 | 0.142627056397866 | 4.05466689199161 | 5.02057953659095e-05 | 0.000786902962980262 |
| znf609.L | 2183.31151382766 | -0.335476764285252 | 0.0827435852008067 | -4.05441416964346 | 5.02600998040515e-05 | 0.00078724620544141 |
| cbl.S | 457.274506889647 | -0.449996331162549 | 0.111109974529935 | -4.05000840893282 | 5.12157925323831e-05 | 0.000801698752436672 |
| wdr70.L | 1264.84094137004 | 0.364943362029783 | 0.0901359056464717 | 4.0488122842095 | 5.14782121927842e-05 | 0.000805287628468448 |
| ntn3.L | 232.135827077881 | -0.69604341060519 | 0.171999408083574 | -4.04677794162516 | 5.19274587815595e-05 | 0.000811792589214418 |
| chrna4.L | 955.465673623175 | -0.511296839728871 | 0.1263698173778 | -4.04603607363204 | 5.20922090435613e-05 | 0.000813844454343588 |
| ogt.L | 10106.070323837 | 0.219341572466717 | 0.054220295952742 | 4.04537763235179 | 5.22388471336571e-05 | 0.000815610894771893 |
| cfl1.S | 16989.7568521533 | -0.217789311949113 | 0.0538390108067743 | -4.0451952717101 | 5.22795288589764e-05 | 0.000815721820231197 |
| kcnc1.S | 1053.22967295163 | -0.570845416905613 | 0.141272647818221 | -4.04073559688734 | 5.32838057240851e-05 | 0.000830858007869655 |
| camkv.L | 1126.25620447134 | 0.395943400049147 | 0.0979988098462865 | 4.040287842987 | 5.33856393463004e-05 | 0.000831911945015409 |
| tyrp1.L | 458.683913569753 | 0.675098464073375 | 0.167127583301214 | 4.0394197698452 | 5.35835926166739e-05 | 0.00083446140963428 |
| kcnj10.L | 644.542356286877 | -0.513320513047791 | 0.127089813035161 | -4.03903743965517 | 5.36709986897239e-05 | 0.000835287150652244 |
| LOC108712767 | 1559.28994749218 | -0.358746750074998 | 0.0888312975896311 | -4.03851750238165 | 5.37900804769994e-05 | 0.000836604491106417 |
| sik2.L | 533.676118853562 | -0.497011645834144 | 0.123090201060907 | -4.03778401164698 | 5.39584983898691e-05 | 0.000838686986489751 |
| nsdhl.L | 562.717770510067 | 0.525775735600414 | 0.130255902359677 | 4.03648300058282 | 5.42584552050824e-05 | 0.000842810045238025 |
| mical3.L | 1113.17119739233 | -0.427647395829568 | 0.105956884665425 | -4.03605105208527 | 5.4358392671422e-05 | 0.000843822870006087 |
| tcf4.L | 728.248059875719 | -0.400550388811741 | 0.0993089192892956 | -4.03337778397228 | 5.49807816554949e-05 | 0.000852939405835628 |
| ncstn.S | 1051.22443869425 | -0.367216370990488 | 0.0910719219907519 | -4.03215791391531 | 5.5267028986934e-05 | 0.000856832930573437 |
| acsbg2.L | 1669.38287275125 | 0.329395640547281 | 0.0817058118011415 | 4.03148360301439 | 5.54258640275181e-05 | 0.000858747411150845 |
| LOC108699158 | 1324.50563779052 | 0.329410687317569 | 0.0817133909374536 | 4.03129831290408 | 5.54706473261196e-05 | 0.000858893502957776 |
| nsf.S | 3637.44649188835 | -0.44518068572614 | 0.110521607557233 | -4.02799683759218 | 5.62541001588548e-05 | 0.000870469496343451 |
| ca7.L | 317.642215180536 | -0.573790440946573 | 0.142477927222884 | -4.02722338912868 | 5.64394049025909e-05 | 0.000872780969257507 |
| hspa4.L | 5301.93462340316 | -0.233986408062096 | 0.0581078642804998 | -4.02675973311617 | 5.65507658240226e-05 | 0.000873391166515452 |
| LOC108713014 | 2050.99500218593 | -0.369099559049397 | 0.0916605088710576 | -4.0268111490481 | 5.65384064906296e-05 | 0.000873391166515452 |
| khsrp.S | 3592.26287302925 | -0.241544455824538 | 0.0600047198757071 | -4.02542427203843 | 5.68726811466452e-05 | 0.000877804902018169 |
| ppme1.S | 1464.67803108197 | -0.354435626961653 | 0.0880630290306069 | -4.02479486412472 | 5.70250021569402e-05 | 0.000879597080922132 |
| ltn1.L | 882.386358172738 | -0.365179679679303 | 0.0907773420567298 | -4.02280647797653 | 5.75087477435218e-05 | 0.000885712767746539 |
| lrwd1.L | 457.154361999778 | 0.431666238144877 | 0.107304339418162 | 4.02282182142407 | 5.7505000068709e-05 | 0.000885712767746539 |
| utp23.L | 660.127000854406 | 0.563183315398319 | 0.140000765862518 | 4.02271596107817 | 5.7530861426866e-05 | 0.000885712767746539 |
| lbx1.L | 206.08702452705 | -0.651397634422407 | 0.161989168181128 | -4.02124192460849 | 5.78921105479172e-05 | 0.000890709900982332 |
| gca.S | 1087.21863950487 | 0.324867577483826 | 0.0808008119065199 | 4.02059793482858 | 5.80506095827805e-05 | 0.000892583233673462 |
| hnrnph1.L | 3463.83802302897 | 0.301099437510524 | 0.0749147425496433 | 4.01922808866354 | 5.8389124623513e-05 | 0.000897220362810642 |
| LOC108697267 | 1111.67694138554 | -0.394699176537314 | 0.0982090890327726 | -4.01896790230486 | 5.84536325275067e-05 | 0.000897643836045037 |
| LOC108699920 | 983.76861490262 | -0.379208769885477 | 0.0943835001638471 | -4.01774430093376 | 5.87579056491087e-05 | 0.000901746405457641 |
| lingo1.S | 1234.58113493634 | -0.398768863215493 | 0.0992905281796123 | -4.01618231392764 | 5.91485050052411e-05 | 0.000907167790781141 |
| LOC100037108 | 1054.00994965433 | 0.478921695292001 | 0.119271637266286 | 4.01538627513565 | 5.93485116308298e-05 | 0.000908015673944384 |
| eif2s3.L | 4061.53852270412 | 0.267908602448053 | 0.0667179804780031 | 4.0155382481396 | 5.93102786600379e-05 | 0.000908015673944384 |
| hp1bp3.S | 2645.3365235738 | -0.276245664397118 | 0.0687971100252578 | -4.01536727771993 | 5.93532925917379e-05 | 0.000908015673944384 |
| mef2d.L | 1118.80786398759 | -0.406802052524331 | 0.101306219741403 | -4.0155683783557 | 5.93027013501044e-05 | 0.000908015673944384 |
| stim2.S | 617.332756812962 | -0.430525176799391 | 0.107241768390062 | -4.01452888424477 | 5.95646494240976e-05 | 0.000910675640723113 |
| tlcd3b.S | 852.826885889964 | -0.425345637843586 | 0.105961547669244 | -4.01415086132273 | 5.96601809725584e-05 | 0.000911562538709015 |
| gsk3b.L | 1016.31609299261 | -0.394515440101965 | 0.0983168494367597 | -4.01269408409724 | 6.00296870489091e-05 | 0.000916505453767907 |
| pa2g4.L | 4097.68798366933 | 0.325931015657493 | 0.0812273276375337 | 4.01257834200723 | 6.00591373342598e-05 | 0.000916505453767907 |
| tmem50a.S | 1038.45027485904 | 0.352394302240183 | 0.0878392969575271 | 4.01180692979108 | 6.02557707956127e-05 | 0.000918928876385409 |
| get1.L | 543.986135317862 | 0.462387355560951 | 0.115262212584714 | 4.01161269762291 | 6.03053769026035e-05 | 0.00091910842313165 |
| LOC108704640 | 752.007703705254 | -0.390230620163113 | 0.0973105384263723 | -4.01015785621587 | 6.06781675815391e-05 | 0.000923631205028766 |
| cdk5r1l.S | 856.424146728012 | -0.438934795401712 | 0.109452951586464 | -4.01026001619488 | 6.06519188915354e-05 | 0.000923631205028766 |
| rab14.L | 1117.44432615586 | -0.36586910768243 | 0.0912393748126877 | -4.00999139279013 | 6.07209612567739e-05 | 0.000923703840182884 |
| ndufs4.S | 430.870826170728 | -0.593657660458918 | 0.148070949428678 | -4.00927840842182 | 6.09045756232597e-05 | 0.000925917246678017 |
| sik3.L | 1179.01546610352 | -0.426553434798127 | 0.106402629810905 | -4.00886176926436 | 6.10121156439806e-05 | 0.000926972068452073 |
| LOC108715794 | 632.268249496101 | -0.450002145218409 | 0.112258240747091 | -4.00863350631184 | 6.10711095064339e-05 | 0.000927288458968316 |
| ddr1.L | 631.579352973456 | -0.445488434772734 | 0.111147675728153 | -4.00807692877282 | 6.12151817463971e-05 | 0.000928895456181744 |
| arf5.S | 1206.2252662248 | -0.340343756157384 | 0.0849632173754809 | -4.00577763731911 | 6.18137798382963e-05 | 0.000937393238072141 |
| mad2l1.S | 1073.62565569852 | 0.369480338468557 | 0.0922669844509862 | 4.00446964498803 | 6.21567714451259e-05 | 0.000942006616024884 |
| ablim3.L | 2293.96886004592 | -0.311039553246567 | 0.0776783540766087 | -4.0041985562646 | 6.22280834252641e-05 | 0.000942499413175416 |
| pwwp2a.S | 1331.61256707835 | -0.394086611316434 | 0.0984416416367573 | -4.00325111166458 | 6.24779249163327e-05 | 0.000945693899013947 |
| hivep3.L | 440.113091119611 | -0.507680459332913 | 0.126836505685612 | -4.00263675342252 | 6.26404386175149e-05 | 0.000947027538527784 |
| sall3.S | 1343.3502940079 | -0.470398651543584 | 0.117522588054571 | -4.00262331974136 | 6.26439966417283e-05 | 0.000947027538527784 |
| tbxas1.L | 133.412830403345 | -0.662587629765119 | 0.16556335214672 | -4.00201869057316 | 6.2804336033288e-05 | 0.000948861032085012 |
| pou3f2.L | 2191.90277729948 | -0.297101645745989 | 0.0742442264937382 | -4.0016801275591 | 6.2894287915331e-05 | 0.000949629478318864 |
| fam168a.S | 752.91357837527 | -0.426647014682762 | 0.106683473104937 | -3.99918564952511 | 6.35608084506779e-05 | 0.000959097068634017 |
| reep5.S | 3143.98937672893 | 0.276155027816685 | 0.0690573753082771 | 3.99892157186555 | 6.36317596497822e-05 | 0.000959571675314593 |
| plekha6.S | 1538.80349972332 | -0.318410937285512 | 0.0796313975503859 | -3.99856020464843 | 6.37289716658043e-05 | 0.000960441462561445 |
| six3.S | 89.1567302643787 | 0.763346532073755 | 0.190915742547105 | 3.99834252476803 | 6.37875979330707e-05 | 0.000960729016854321 |
| kctd12.L | 1485.63367630118 | 0.437838411127432 | 0.10954310470611 | 3.9969509016756 | 6.41636025982229e-05 | 0.000965793408625296 |
| kmt5a.S | 658.290669302811 | 0.387122714716421 | 0.0968828696525241 | 3.99578084448632 | 6.44813645715278e-05 | 0.000969975399938512 |
| rab11fip3.L | 1455.40388028331 | -0.328464504259764 | 0.0822363546066971 | -3.99415204906243 | 6.49261905479339e-05 | 0.000976062421517021 |
| ppp1r16b.L | 329.458455688267 | -0.590368056684247 | 0.147813967509002 | -3.99399371137436 | 6.4969587311855e-05 | 0.000976110794158444 |

| | | | | | | |
|---|---|---|---|---|---|---|
| ddi2.S | 1854.56826820402 | -0.278041939845214 | 0.0696247895064157 | -3.99343311220486 | 6.51234555856819e-05 | 0.000977817818293298 |
| LOC108704618 | 1297.41681720637 | -0.358282796934678 | 0.0897403828025761 | -3.99243669065776 | 6.5397796182662e-05 | 0.000981330488240636 |
| nyap1.L | 1406.48365882324 | -0.413681400935812 | 0.103703052163847 | -3.98909571419566 | 6.63256570780003e-05 | 0.000994639205588234 |
| LOC108708403 | 2164.46769533657 | -0.302460632188863 | 0.075837554123562 | -3.98826987083556 | 6.65569243953052e-05 | 0.000997491623232292 |
| LOC108712074 | 2489.35647293416 | 0.27208183605626 | 0.0682317226129049 | 3.98761493389 | 6.67408738201949e-05 | 0.000999631805541192 |

**KDBarhl1_B_vs_KDCT pAdj<0.001**

| Gene | meanCounts | Log2FC | SD | Wald | pVal | pAdj |
|---|---|---|---|---|---|---|
| LOC108719610 | 156.782034411883 | 3.1810690765492 | 0.19432898583171 | 16.3695038232949 | 3.15777838514499e-60 | 8.14706823367408e-56 |
| sspo.L | 4142.38943792403 | 3.21909264454836 | 0.199534455280629 | 16.1330164257645 | 1.49535073398284e-58 | 1.92900244683786e-54 |
| dmrta2.S | 117.012751895943 | 2.61200347554515 | 0.192165465550229 | 13.592470780669 | 4.43840182143429e-42 | 3.81702556643349e-38 |
| helt.L | 355.626959876354 | 2.28133128682221 | 0.176222836646858 | 12.9457187855504 | 2.48427090071676e-38 | 1.60235473096231e-34 |
| dmrta2.L | 138.305196425389 | 2.36496661877021 | 0.197326827978555 | 11.9850232378297 | 4.2572494937828e-33 | 2.19674073879192e-29 |
| pou4f4.L | 340.333953750134 | 1.62062217031122 | 0.149934045412213 | 10.8089004458971 | 3.12391932479472e-27 | 1.34328530966173e-23 |
| tal2.S | 859.993125967164 | 1.59821619803655 | 0.15111557680077 | 10.5761181730698 | 3.84562495168169e-26 | 1.41738748219125e-22 |
| nefh.L | 2355.09812276355 | -0.99397302639532 | 0.0946628033277826 | -10.5001435775524 | 8.62488410962893e-26 | 2.78152512535533e-22 |
| apcdd1.S | 577.622946124479 | 1.34075647468684 | 0.130052721441784 | 10.3093303994182 | 6.39452885701532e-25 | 1.71398122929839e-21 |
| otx2.S | 470.358645443961 | 1.72288490785298 | 0.167178495995367 | 10.3056610097792 | 6.64333809805577e-25 | 1.71398122929839e-21 |
| rspo2.L | 181.896804776632 | 1.70221222410836 | 0.167178148740188 | 10.1820258026291 | 2.38542843391584e-24 | 5.59491396318441e-21 |
| pou4f4.S | 211.160785658853 | 1.65238378102242 | 0.166036180548116 | 9.9519500844189 | 2.47289180233056e-23 | 5.31671737501071e-20 |
| rbp1.S | 531.306317481309 | 1.34507033905781 | 0.141574743956867 | 9.50077889222675 | 2.08326170569063e-21 | 4.13447323129371e-18 |
| rgr.S | 626.760727070801 | 1.29899404456056 | 0.136976654992836 | 9.48332432726218 | 2.46308483989718e-21 | 4.53911349066766e-18 |
| gucy2d.L | 230.045764219767 | 1.88129046903406 | 0.20091720927526 | 9.36351085016646 | 7.7130224838002e-21 | 1.32663986721363e-17 |
| adgrg2.S | 123.799541142698 | 1.77969057330709 | 0.190422359147471 | 9.34601682950913 | 9.10107143256411e-21 | 1.46754776850096e-17 |
| nr2e1.S | 118.883704211145 | 1.76973218574963 | 0.190202114607164 | 9.30448217889038 | 1.34646823022203e-20 | 2.04346354939579e-17 |
| LOC121393901 | 1763.48099218195 | 1.30921444061879 | 0.140902502589486 | 9.29163369392481 | 1.51938034091554e-20 | 2.17777848864561e-17 |
| rngtt.S | 701.758932352641 | 1.0134079875457 | 0.109329096446329 | 9.26933470124485 | 1.87310598373402e-20 | 2.54348075685988e-17 |
| lef1.L | 611.483845858957 | 1.52527959725695 | 0.165051861220575 | 9.24121416127846 | 2.43715781016604e-20 | 3.14393357511419e-17 |
| glra1.S | 628.88944714456 | -1.07762167600797 | 0.11735743386858 | -9.18238956395998 | 4.21629396108604e-20 | 5.18001829504857e-17 |
| zic3.L | 801.736647368952 | 1.19332328996792 | 0.131477928046598 | 9.07622524706619 | 1.12405596661534e-19 | 1.31821108812163e-16 |
| rgr.L | 282.325895593198 | 1.51911968008455 | 0.169044897494711 | 8.98648644589868 | 2.55260166076371e-19 | 2.86335316729147e-16 |
| stra6.S | 44.7508868074521 | 1.81164332884186 | 0.202401397203722 | 8.95074517207211 | 3.53090625954498e-19 | 3.79572422901085e-16 |
| sp5.S | 226.980832880496 | 1.63736136865134 | 0.183414641444208 | 8.9271028515431 | 4.37310790749678e-19 | 4.51304736053668e-16 |
| otx2.L | 1416.49722874322 | 1.18524214857712 | 0.134890776442049 | 8.78668045243493 | 1.54044152153754e-18 | 1.52859197137187e-15 |
| igfbpl1.S | 1240.73146386321 | 1.01661446309663 | 0.117196964828012 | 8.67440948311697 | 4.15701943163305e-18 | 3.97226301244935e-15 |
| igfbpl1.L | 3414.24136106187 | 0.93163569952425 | 0.10907983538976 | 8.54086088593155 | 1.33225526051821e-17 | 1.2275780614775e-14 |
| otp.S | 3625.94363090596 | -0.921235885121152 | 0.108529559375473 | -8.48834078404402 | 2.09608152884992e-17 | 1.86478977394234e-14 |
| otp.L | 4911.02696403383 | -0.919606050792354 | 0.108476589522507 | -8.47746094194409 | 2.30161592257 19e-17 | 1.97938969341183e-14 |
| nefm.S | 16025.3909828869 | -0.762022448149178 | 0.0905094139995149 | -8.41926175937082 | 3.78864055679 64e-17 | 3.15312665694668e-14 |
| slc4a4.L | 1477.33293108437 | -0.752992036083416 | 0.0895777114507225 | -8.40602002315769 | 4.24158760252116e-17 | 3.41978000453268e-14 |
| apcdd1.L | 421.353262848471 | 1.22229246954597 | 0.145954658579075 | 8.37446698478458 | 5.54743709067878e-17 | 4.33708717998523e-14 |
| nefl.S | 1996.8248665339 | -0.925448119270653 | 0.111763813459887 | -8.28039139522389 | 1.22771092017222e-16 | 9.31615933542451e-14 |
| nmu.S | 78.070086575159 | 1.65886055462668 | 0.201666284453726 | 8.22577040639295 | 1.93940200487957e-16 | 1.42961633502551e-13 |
| LOC108696915 | 4068.48123477839 | -0.615369071387817 | 0.0748460609107821 | -8.22179636308915 | 2.00477383554423e-16 | 1.43675458214003e-13 |
| pitx3.S | 50.9953654701972 | 1.6660253982505 | 0.203296696256699 | 8.19504413464173 | 2.50501232713534e-16 | 1.74673832540788e-13 |
| spa17.L | 1868.2341117073 | -0.699991090810083 | 0.0855295687180068 | -8.1841999357903 | 2.74118650504457e-16 | 1.86112136395131e-13 |
| nkx2-2.L | 533.234259314709 | 1.40708318366827 | 0.172441016957405 | 8.15979404723552 | 3.35596324943059e-16 | 2.22009876500793e-13 |
| nr2e1.L | 113.236747522043 | 1.62234185415289 | 0.203369632730076 | 7.97730630858813 | 1.49561341037173e-15 | 9.64670649689765e-13 |
| wnt2b.L | 60.3092633045348 | 1.59458424360865 | 0.202411660640308 | 7.87792678823121 | 3.32857642509154e-15 | 2.09456760408199e-12 |
| cyp27c1.S | 1284.13950789564 | 0.840248444613843 | 0.106845822438191 | 7.86412070626271 | 3.71699007584096e-15 | 2.28329390373087e-12 |
| XB5848002.S | 196.775260754966 | 1.40046777069367 | 0.180608153583025 | 7.75417799756132 | 8.89175369504494e-15 | 5.33505221702696e-12 |
| lef1.S | 439.93394857345 | 1.25051773312791 | 0.162555082141877 | 7.69288610759318 | 1.43852558692875e-14 | 8.43499094153673e-12 |
| LOC108696667 | 168.661400057737 | -1.5318422709977 | 0.200032000021817 | -7.6579860763809 | 1.88871357436078e-14 | 1.08286244930018e-11 |
| gipr.L | 138.148595458386 | 1.35511250480021 | 0.177205553747777 | 7.64712209149487 | 2.05527167165656e-14 | 1.15273932888564e-11 |
| LOC108700443 | 286.575969523474 | -1.07143853860248 | 0.140500451714031 | -7.62587255436904 | 2.42389210722669e-14 | 1.33056205035009e-11 |
| pitx3.L | 54.0467037259541 | 1.52479051199433 | 0.202302898080476 | 7.53716593515019 | 4.80295020842886e-14 | 2.58158573703051e-11 |
| nefh.S | 1858.9794074709 | -1.01938135966492 | 0.137389460414124 | -7.41964745033759 | 1.17432498379072e-13 | 6.18318052689806e-11 |
| MGC115323 | 2092.82352124555 | -0.705999933159814 | 0.0954931598204701 | -7.39319899443179 | 1.43337711267243e-13 | 7.39622590138972e-11 |
| barhl2.L | 517.10510954355 | 1.21936468235613 | 0.165590251024735 | 7.36374680761844 | 1.78818715731822e-13 | 9.04612326643336e-11 |
| slc39a12.S | 461.743774484411 | 1.13628926171598 | 0.154499125533346 | 7.35466468402066 | 1.91406996704971e-13 | 9.49673175959282e-11 |
| emx1.L | 47.4485194355833 | 1.47453586819206 | 0.200622339690691 | 7.34980895181177 | 1.98490353717722e-13 | 9.66236061493819e-11 |
| rora.L | 642.834601990194 | -0.854757059142449 | 0.116506319539592 | -7.3365724925503 | 2.1913314501215e-13 | 1.04696947061361e-10 |
| scn1b.L | 2370.83008920329 | -0.650758404543486 | 0.0893289094936565 | -7.2849697621093 | 3.21742458397 06e-13 | 1.50926462302621e-10 |
| gucy2d.S | 146.322887937198 | 1.47483204525216 | 0.203358500240005 | 7.25237471515355 | 4.09527002996709e-13 | 1.88674940666341e-10 |
| LOC108704633 | 2319.69689872655 | -0.554329447003717 | 0.0768945970138589 | -7.20895184487161 | 5.63841953986211e-13 | 2.55212673909548e-10 |
| lhx2.L | 714.039594332585 | 0.841140848441365 | 0.117084340801491 | 7.18405930872916 | 6.76714856259183e-13 | 3.01021436060119e-10 |
| chac1.L | 206.952723627476 | 1.31910438539115 | 0.185955559394674 | 7.09365393 36878 | 1.30616517429737e-12 | 5.71170533845289e-10 |
| cox4i2.L | 1455.88645940465 | 0.781384904209809 | 0.111053323049522 | 7.03612355056104 | 1.976616867974e-12 | 8.49945253228819e-10 |
| otx1.L | 650.414519435943 | 1.15092877868824 | 0.16565294911006 | 6.94783150478992 | 3.70943657208778e-12 | 1.56890923868631e-09 |
| tf.L | 120.704156582953 | 1.37032918648617 | 0.197620823077956 | 6.93413358543502 | 4.08718300941427e-12 | 1.70079551036917e-09 |
| LOC108715150 | 1268.5445064191 | -0.660421479967665 | 0.0960856591759328 | -6.87325752491777 | 6.27521315706488e-12 | 2.56984919765514e-09 |
| atp2b4.S | 3694.21179170575 | -0.673410401570945 | 0.098558815651255 | -6.83257400285502 | 8.34043552691231e-12 | 3.36223807178652e-09 |
| LOC108695757 | 5322.68403263409 | -0.492423506880923 | 0.0723132213399628 | -6.80959163146551 | 9.78761761102731e-12 | 3.88493129791546e-09 |
| slit1.S | 1973.25019375614 | -0.557748351841773 | 0.0819808253460956 | -6.80340005711271 | 1.02178384772331e-11 | 3.99424595019111e-09 |
| pygm.L | 8074.38027535992 | -0.653469422858846 | 0.0961667131225771 | -6.79517269167673 | 1.08182861947416e-11 | 4.16584751976618e-09 |
| crx.S | 59.9648889229464 | 1.31349381729512 | 0.193785017649984 | 6.77809787992777 | 1.21768347756303e-11 | 4.62003437075383e-09 |
| hspa12a.S | 513.7226414775 | -0.880372562290956 | 0.130284151836044 | -6.75732658104773 | 1.40561124629209e-11 | 5.25576379048347e-09 |
| rbp4l.S | 29.2511909660974 | 1.27965177355472 | 0.190802263756527 | 6.70669072976839 | 1.99087580987808e-11 | 7.33779941355065e-09 |
| ltbp4.L | 3079.80345045708 | -0.542186722474544 | 0.0813744431312283 | -6.66286246162309 | 2.68544958986289e-11 | 9.62286103034203e-09 |
| paqr6.S | 607.124536496932 | -0.825970375008692 | 0.123959260537868 | -6.66324057940285 | 2.6785465104687e-11 | 9.62286103034203e-09 |
| XB22063245.S | 2213.39506527553 | -0.660684060279927 | 0.0994871161371663 | -6.64090071089225 | 3.1177201309821e-11 | 1.10187916957998e-08 |
| LOC121399515 | 1055.83386692912 | -0.766663453217245 | 0.115492804852627 | -6.63819234622915 | 3.17553313844851e-11 | 1.10714533745907e-08 |
| crx.L | 55.7573777029921 | 1.26495346835416 | 0.191490711616418 | 6.60582151443103 | 3.95318865250264e-11 | 1.35989689646091e-08 |
| kcnh4.L | 1037.21588846878 | -0.62326190425084 | 0.0947801013805999 | -6.57587294349964 | 4.83685453728766e-11 | 1.64198482976344e-08 |
| tfap2e.S | 343.521963661262 | 1.03451258739366 | 0.157490747586983 | 6.56871977080621 | 5.07496886256534e-11 | 1.70044411239202e-08 |
| slc7a10.S | 1710.74340578429 | -0.654111875561382 | 0.0998813242531885 | -6.54889070056058 | 5.7966034772497e-11 | 1.84632555201287e-08 |
| ccnb3.S | 1413.29306329285 | 0.615990397893148 | 0.094022119891205 | 6.55154764225614 | 5.69438145644165e-11 | 1.84632555201287e-08 |
| emilin3.L | 387.739731117739 | -0.879232133123832 | 0.134244261021185 | -6.54949512504735 | 5.77319255050734e-11 | 1.84632555201287e-08 |
| aldoa.L | 11145.2980043162 | -0.473483055933391 | 0.0722655894196909 | -6.55198497286977 | 5.6777256965 29e-11 | 1.84632555201287e-08 |
| atp1b1.L | 6264.91677691865 | -0.575493053392598 | 0.0880565721171792 | -6.53549234947249 | 6.34006232027242e-11 | 1.98443627982492e-08 |
| LOC108697760 | 11777.8334107533 | -0.558458449998259 | 0.0854636248600281 | -6.53445779900981 | 6.38403919478559e-11 | 1.98443627982492e-08 |
| eef1a2.L | 2026.52857041853 | -0.652514664200081 | 0.0999662896997553 | -6.52734703028273 | 6.69448197585004e-11 | 2.05616232115394e-08 |
| barhl2.S | 454.318010795642 | 1.11618098750289 | 0.172711589890249 | 6.4626872360574 | 1.02859751620979e-10 | 3.12209599037795e-08 |
| slc6a3.S | 85.8663339348661 | 1.25555007651986 | 0.194461960730999 | 6.45653305047495 | 1.07128794212294e-10 | 3.21386382636881e-08 |
| nefl.L | 4392.48455502415 | -0.782570457915166 | 0.121355216428791 | -6.44859348402516 | 1.12892920131848e-10 | 3.34785901080653e-08 |
| pcdh7.S | 746.722157201426 | -0.69196514896106 | 0.107442414518041 | -6.44033505822667 | 1.19210042462536e-10 | 3.49502169946979e-08 |
| calb1.L | 2707.65684942868 | -0.522627843847289 | 0.0816145178107201 | -6.40361369357551 | 1.51741762328403e-10 | 4.39880614390201e-08 |
| sema3a.L | 2023.13620582696 | -0.534796509481576 | 0.0835853972791755 | -6.39820503209864 | 1.57214092491816e-10 | 4.5068039847654e-08 |
| rlbp1.S | 204.435798332881 | 1.04427813213364 | 0.163572811543975 | 6.38417914491185 | 1.72319271037e-10 | 4.8855353766534e-08 |
| LOC108707869 | 1501.83963964252 | -0.643125898605353 | 0.100804550129072 | -6.37992925698179 | 1.77169806410885e-10 | 4.96845761456611e-08 |
| nfia.S | 992.068476484675 | -0.630186787143741 | 0.0988431125308008 | -6.37562669778703 | 1.82216277124636e-10 | 5.0550322041028e-08 |
| frmpd1.L | 501.733205542558 | 0.800406358400199 | 0.1261175056915 | 6.34651275420953 | 2.20250295191493e-10 | 6.04516767653246e-08 |
| LOC108701890 | 167.572938110077 | -1.28895744909587 | 0.203357735789011 | -6.33837431408656 | 2.32202147558475e-10 | 6.30612148106174e-08 |
| LOC108699226 | 9415.63642478473 | -0.649671777665872 | 0.102805330211 | -6.31943670948257 | 2.62518435530091e-10 | 7.05518295487121e-08 |
| gpt.S | 544.138412171406 | -0.822168233632312 | 0.130505221423667 | -6.29988765708657 | 2.97861451312381e-10 | 7.92250045758705e-08 |
| wls.L | 747.004438583496 | 0.613259911338564 | 0.0974995175274321 | 6.28987636955248 | 3.1771890049836e-10 | 8.36443636005885e-08 |
| hmgb2.L | 26103.8498814937 | 0.453065663535985 | 0.0721402120066805 | 6.2803483789877 | 3.37815189736086e-10 | 8.80366858100102e-08 |
| LOC108712579 | 368.250986722816 | -0.849832928529222 | 0.135806271018026 | -6.25768546738479 | 3.90732849684672e-10 | 1.00809075218645e-07 |
| LOC108714020 | 25.7683355869744 | 1.22593597187203 | 0.196251637030343 | 6.24675539232566 | 4.19066555050356e-10 | 1.07048684359398e-07 |
| dner.S | 6280.84142001428 | -0.451682408960077 | 0.0725859270626361 | -6.22272701112311 | 4.8858708654902e-10 | 1.23583792480046e-07 |
| lhx2.S | 948.254554364657 | 0.681188911742272 | 0.10982737375629 | 6.20236001685562 | 5.56226536619085e-10 | 1.39326647036625e-07 |
| mapk8ip2.S | 2874.59735810722 | -0.537072369833608 | 0.0867327445102291 | -6.19226767083644 | 5.93047270556427e-10 | 1.47121342118806e-07 |
| ntn1.L | 5856.4311138411 | -0.493666861804794 | 0.0798382948993441 | -6.18333423111281 | 6.276163519012e-10 | 1.54214303610009e-07 |

| | | | | | | |
|---|---|---|---|---|---|---|
| zic3.S | 447.314484161044 | 0.956050052574116 | 0.154727798839853 | 6.17891587512113 | 6.45432744505008e-10 | 1.57095894417257e-07 |
| LOC108710372 | 3861.64470993209 | -0.543133783629191 | 0.0879565658352015 | -6.17502262021946 | 6.61539985946132e-10 | 1.59511510630002e-07 |
| coch.L | 112.634881338041 | -1.142492198744 | 0.185067454255932 | -6.17338258278538 | 6.6844196941479e-10 | 1.5968335935602e-07 |
| gpc1.L | 1072.73786536492 | -0.65775626020303 | 0.106639279208921 | -6.16804863163407 | 6.91378827802231e-10 | 1.6364746566328e-07 |
| lmo3.L | 608.981642834842 | 0.811694789358855 | 0.131707192794997 | 6.16287366037984 | 7.14365026922548e-10 | 1.67551069950925e-07 |
| prdm12.L | 459.923463216879 | 0.917210969310662 | 0.148887016393241 | 6.16044965860637 | 7.25386727743468e-10 | 1.68603401583617e-07 |
| emx2.L | 148.999501744891 | 1.2484094566646 | 0.202917249107997 | 6.15230820520423 | 7.63632932498875e-10 | 1.75908300522062e-07 |
| fst.L | 596.827207803899 | 0.993278103964822 | 0.161526352582997 | 6.14932540778105 | 7.78131879668434e-10 | 1.76103530661803e-07 |
| LOC108719750 | 1457.73650578294 | -0.643070651474493 | 0.104552150309441 | -6.15071664782801 | 7.71336155590592e-10 | 1.76103530661803e-07 |
| hes5.1.S | 942.443557367955 | 0.752770171510055 | 0.122461301599092 | 6.14700449595448 | 7.89598918635466e-10 | 1.77144800876478e-07 |
| LOC108712326 | 76.7304038852918 | 1.24510896067674 | 0.202879100897787 | 6.1371967598774 | 8.39903345493632e-10 | 1.83639884014709e-07 |
| LOC108695500 | 65.495327846809 | -1.2335497576847 | 0.200984363696115 | -6.13754092606827 | 8.38086328971128e-10 | 1.83639884014709e-07 |
| cntnap1.L | 1108.51270430255 | -0.975294864645412 | 0.15888582883831 | -6.13833764644875 | 8.33894762130875e-10 | 1.83639884014709e-07 |
| slc6a4.S | 202.824555532245 | -1.00497521436933 | 0.163808407093032 | -6.13506493472323 | 8.51244170299922e-10 | 1.84555458770908e-07 |
| mapt.S | 10899.5902042932 | -0.425541602973233 | 0.0696101901926053 | -6.11320845117356 | 9.76476852334926e-10 | 2.09942523252009e-07 |
| tfap2e.L | 209.834268711815 | 1.00778388888918 | 0.165330017779847 | 6.09558930932403 | 1.09035016822973e-09 | 2.30582248691205e-07 |
| ptgds.S | 18065.1874376707 | -0.598096410962996 | 0.0981124153621032 | -6.0960318707434 | 1.0873374420353e-09 | 2.30582248691205e-07 |
| pmp2.S | 5163.70241883322 | -0.71121002956653 | 0.116993087918064 | -6.07907733886486 | 1.20876051959616e-09 | 2.53544889476267e-07 |
| map4.L | 5532.97223937178 | -0.440706534306122 | 0.0725390509457793 | -6.07543838194873 | 1.23649454014981e-09 | 2.57270638192462e-07 |
| LOC108708112 | 140.450117175743 | -1.22939435731365 | 0.203301507741372 | -6.04714825272035 | 1.47432083857877e-09 | 3.04299821082658e-07 |
| dner.L | 6711.15592270585 | -0.630541318264562 | 0.104346733490285 | -6.04275090531001 | 1.51508473960744e-09 | 3.10231637157714e-07 |
| stxbp1.S | 15010.8113999687 | -0.578955440067159 | 0.0966850517130959 | -5.98805533853524 | 2.12364814548455e-09 | 4.31418284673239e-07 |
| LOC108699444 | 126.822778627423 | -1.18860259877292 | 0.198876573142117 | -5.97658427030289 | 2.27864706530282e-09 | 4.592897991001e-07 |
| neurod2.S | 1140.82700604756 | -0.642452354784383 | 0.107765574523576 | -5.96157314267212 | 2.49821047267002e-09 | 4.99642094534004e-07 |
| syn1.L | 10708.6472445257 | -0.514558771559184 | 0.0867543492705728 | -5.93121585125788 | 3.00699592011848e-09 | 5.96773036454283e-07 |
| tcf7l2.L | 2667.92183232244 | 1.20065706805081 | 0.20249021473201 | 5.92945723150051 | 3.03937661515627e-09 | 5.98594783748333e-07 |
| paqr6.L | 3087.1669309869 | -0.628002424185084 | 0.106455835267448 | -5.89918272312043 | 3.6530650634l5e-09 | 7.14008171485666e-07 |
| LOC108706170 | 1075.06509238141 | -0.594625620228668 | 0.10086295152177 | -5.8953819143427 | 3.73815759938775e-09 | 7.25146361384992e-07 |
| gja7.S | 2014.09112347778 | 0.567251500489393 | 0.0962941946370646 | 5.89081722555944 | 3.84290406669226e-09 | 7.39902424781047e-07 |
| eef1a2.S | 3134.69205294262 | -0.595024664427067 | 0.101446782049019 | -5.86538727408379 | 4.48085603593884e-09 | 8.563413757572e-07 |
| LOC108706956 | 207.274997560749 | 1.17982073012577 | 0.201647834772115 | 5.85089709224551 | 4.88928676806925e-09 | 9.14084047943382e-07 |
| sim2.S | 35.4437792060384 | 1.16730337678644 | 0.199473791136331 | 5.8519135277709 | 4.8594928050965e-09 | 9.14084047943382e-07 |
| odc1.S | 3565.18543852555 | 0.467652104114792 | 0.0799105165293528 | 5.85219723793193 | 4.85120822164554e-09 | 9.14084047943382e-07 |
| ank1.S | 2302.3732372395 | -0.533921890665488 | 0.0915751780350067 | -5.83042154131968 | 5.52875296423391e-09 | 1.01164415941301e-06 |
| syn2.S | 4609.38527736913 | -0.535155283613625 | 0.0917678643430487 | -5.83161967911877 | 5.48919194395202e-09 | 1.01164415941301e-06 |
| mapt.L | 19386.3683509065 | -0.405100512255889 | 0.0694734933510011 | -5.83100824093008 | 5.5093463381269e-09 | 1.01164415941301e-06 |
| ednrb2.L | 232.438236689278 | 1.16099140192111 | 0.199258479094511 | 5.82655958831461 | 5.65816700169812e-09 | 1.02803315946346e-06 |
| rph3a.S | 1857.78079957403 | -0.590510023875176 | 0.101911335373617 | -5.79435076294807 | 6.85859801877269e-09 | 1.23742537681354e-06 |
| LOC108709778 | 1762.63993597561 | -0.556234398195932 | 0.0960251288123163 | -5.79259205455592 | 6.93083144245173e-09 | 1.2417739667726e-06 |
| sp5.L | 276.05297834953 | 1.00989879166109 | 0.174390270016106 | 5.79102716893447 | 6.99572583715057e-09 | 1.24475673516196e-06 |
| sulf1.S | 1900.56344339141 | -0.644257665209649 | 0.111866060928573 | -5.75918790616046 | 8.45195735604269e-09 | 1.49356506702672e-06 |
| wdr7.L | 2177.84930747007 | -0.587647315031514 | 0.102282745281103 | -5.74532208161295 | 9.17461812644579e-09 | 1.61023909974355e-06 |
| epha4.L | 4345.78567269124 | -0.406693353567827 | 0.0708622882371112 | -5.73920718178048 | 9.51207952757425e-09 | 1.65818683656362e-06 |
| hmga2.L | 4876.78879249367 | 0.45111275497089 | 0.0788209646888793 | 5.72325848524582 | 1.04500084699849e-08 | 1.80946455386316e-06 |
| LOC108707553 | 3491.88688777452 | -0.645862971143558 | 0.11304038931157 | -5.71355933111115 | 1.10637274864332e-08 | 1.89035873609256e-06 |
| ldha.S | 5963.10950754560 | -0.445985976559351 | 0.0780469029064 74 | -5.71433279157521 | 1.10135280021238e-08 | 1.89035873609256e-06 |
| cxcl14.L | 942.987456074912 | -0.708853023127514 | 0.124175044047044 | -5.70849826200961 | 1.13977341738227e-08 | 1.91188119406242e-06 |
| tc2n.L | 208.858255455612 | 0.944927362203563 | 0.165535185369493 | 5.70831729879288 | 1.14098565539665e-08 | 1.91188119406242e-06 |
| plp1.S | 287.494983956113 | -0.949088361340828 | 0.166265054738488 | -5.70828526074595 | 1.14120040265741e-08 | 1.91188119406242e-06 |
| cacna1a.S | 3735.76621042826 | -0.567607551035249 | 0.0995817884823734 | -5.69991320386577 | 1.19868442656695e-08 | 1.99522956164047e-06 |
| not.S | 21.8489503525621 | 1.04027136054 | 0.182590908583237 | 5.69727906286075 | 1.21734581649668e-08 | 1.99983192611957e-06 |
| ptn.S | 9639.34891303288 | -0.391322421859219 | 0.068711220148437 | -5.69517498035756 | 1.2324545591202e-08 | 1.99983192611957e-06 |
| rdh10.L | 793.798449779001 | 0.661683746161507 | 0.116144384950465 | 5.69707908345042 | 1.21877402931012e-08 | 1.99983192611957e-06 |
| slc12a5.L | 4139.43643494924 | -0.684699314022595 | 0.120202634749885 | -5.69620886802775 | 1.22500791350518e-08 | 1.99983192611957e-06 |
| snrpg.S | 2495.79491748458 | 0.508085010019372 | 0.0892533142538339 | 5.69261785141549 | 1.25106187586308e-08 | 2.01733727482922e-06 |
| ache.L | 6534.51257333361 | -0.566941287303238 | 0.0999241823865755 | -5.67371454799519 | 1.39733901032968e-08 | 2.2392140662426e-06 |
| pfkp.S | 5873.61067699205 | -0.535388203220184 | 0.0945122669251192 | -5.66474829816922 | 1.47240508677113e-08 | 2.34494143448736e-06 |
| penk.S | 970.331735351881 | -0.584199838380088 | 0.103226554834485 | -5.65939490392566 | 1.51907640798403e-08 | 2.40442768871091e-06 |
| slc2a1.S | 1291.21679183709 | -0.527588354541347 | 0.0933863224112378 | -5.64952490813429 | 1.60891872676699e-08 | 2.53110385064563e-06 |
| lmx1a.L | 344.623471990967 | 0.90900072975241 | 0.161172974653974 | 5.63990788037189 | 1.70141163637638e-08 | 2.66038910415215e-06 |
| znfx1.L | 108.283314652087 | -1.10790270764709 | 0.19709507084563 | -5.62115887979174 | 1.89680727536721e-08 | 2.9480498617153e-06 |
| proca1.L | 1204.78427376819 | -0.622610062859884 | 0.110924580244028 | -5.61291340016951 | 1.98948179468445e-08 | 3.0552756132654e-06 |
| lhx9.L | 1415.12400968041 | 0.706275862158797 | 0.125817259068874 | 5.61350539175372 | 1.9826842116338e-08 | 3.0552756132654e-06 |
| LOC108708109 | 100.718680754221 | -1.14009684409742 | 0.203341980424454 | -5.60679522112255 | 2.06107274918874e-08 | 3.14648975911654e-06 |
| LOC108708376 | 196.568604555035 | 1.00288068553229 | 0.17897603003721 | 5.60343575239536 | 2.10144031877755e-08 | 3.18924471908594e-06 |
| sv2c.S | 553.148421041548 | -0.691147047789092 | 0.123430188738786 | -5.59949761765137 | 2.14973887676433e-08 | 3.24346567371461e-06 |
| nmu.L | 64.0200633230236 | 1.13838945796392 | 0.203376786447791 | 5.59744048397654 | 2.17539495415688e-08 | 3.26309243123532e-06 |
| LOC108709832 | 11105.6813734492 | 0.656017003349947 | 0.117480176416113 | 5.58406552801169 | 2.34960109514851e-08 | 3.50402937889199e-06 |
| nefm.L | 13465.4012372192 | -0.618936153336342 | 0.110873613757492 | -5.58235753630355 | 2.37279955273245e-08 | 3.5182889919826e-06 |
| LOC108708151 | 1458.82334958647 | -0.645431018738433 | 0.115741029458738 | -5.57651009116462 | 2.45391685745648e-08 | 3.61777456699297e-06 |
| scn4b.L | 899.481118002034 | -0.549478272555529 | 0.0985585805173283 | -5.57514393644216 | 2.47325285098811e-08 | 3.62556383883803e-06 |
| LOC108697003 | 53.5599974089183 | -1.12371534092329 | 0.2021632586176 | -5.55845482807954 | 2.72173376231825e-08 | 3.94498489145005e-06 |
| kcnc1.S | 1072.97114983114 | -0.685768462566035 | 0.123372063335164 | -5.55853929996301 | 2.72041709937833e-08 | 3.94498489145005e-06 |
| snrpb.L | 4871.6558223398 | 0.548544277605623 | 0.0988174439132784 | 5.55108749915675 | 2.83897935118913e-08 | 4.09193671847372e-06 |
| MGC68655 | 2739.95485558831 | 0.508320184793413 | 0.0915954875368222 | 5.54962038483679 | 2.86290557403332e-08 | 4.10349798944775e-06 |
| LOC108707429 | 599.687771757056 | -0.622909141540663 | 0.112414437271608 | -5.54118453696151 | 3.00432407835121e-08 | 4.28240669731 83e-06 |
| LOC108695671 | 1692.21524876977 | -0.494217616502281 | 0.0893267482561779 | -5.5326945864516 | 3.1534829180723e-08 | 4.47032193880579e-06 |
| ldha.L | 1532.52042109379 | -0.569520459404216 | 0.102987533728921 | -5.52999415349903 | 3.2024146072589e-08 | 4.51487961023386e-06 |
| f7.S | 199.252336134486 | 1.05086577557405 | 0.190284390736856 | 5.52260630262252 | 3.34007591018732e-08 | 4.68336730884 96e-06 |
| fry.L | 3611.6918942934 | -0.473763093867149 | 0.0859333951595582 | -5.51314297529479 | 3.52481657863173e-08 | 4.91569014749723e-06 |
| rspo2.S | 35.7038767772253 | 1.11397044125054 | 0.202093540257016 | 5.51215263898999 | 3.54471294216051e-08 | 4.91685988751296e-06 |
| dmbx1.L | 1219.76159995558 | 0.731350165115194 | 0.133079094292823 | 5.49560521884792 | 3.89372697874051e-08 | 5.3720939064976e-06 |
| app.S | 11469.4257929314 | -0.498592147117808 | 0.0909125424850348 | -5.48430539383365 | 4.15097148604021e-08 | 5.69654597552327e-06 |
| LOC108696020 | 248.475104067881 | -0.885847227890462 | 0.161617812272432 | -5.48112374146748 | 4.22632743004124e-08 | 5.76927236481821e-06 |
| rpl28.S | 6048.30405606517 | 0.608878088679056 | 0.111186275908058 | 5.47619824215129 | 4.34560833338321e-08 | 5.90087868427825e-06 |
| sfxn5.L | 388.236969608553 | -0.718320848836676 | 0.131220249487364 | -5.47416158438145 | 4.39587825781342e-08 | 5.91808228147624e-06 |
| kctd15.L | 537.530418832347 | 0.680485469590173 | 0.124316184804603 | 5.47382845330832 | 4.40415425598232e-08 | 5.91808228147624e-06 |
| arpp19.S | 2455.72852230839 | 0.39720690892352 | 0.0726547726733455 | 5.46704496219891 | 4.5760001045982e-08 | 6.11714003619863e-06 |
| LOC108708999 | 3525.29655742698 | -0.57208873133747 | 0.104713762819655 | -5.46335759438577 | 4.67212094111238e-08 | 6.21343918972677e-06 |
| slc40a1.S | 2387.32465343191 | -0.574556572362209 | 0.105230400933393 | -5.45998653683628 | 4.76170690674412e-08 | 6.30010452276915e-06 |
| dhrs3.L | 291.768904339392 | 0.791648573928416 | 0.145160068363661 | 5.45362497312376 | 4.93532319166532e-08 | 6.49649685433496e-06 |
| LOC108717850 | 1732.32440152919 | -0.484177224045773 | 0.0888172702065797 | -5.45138600769453 | 4.99787538039323e-08 | 6.54544085350992e-06 |
| atp2b4.L | 3951.58752560592 | -0.517456442343726 | 0.0949733562726397 | -5.44843798989546 | 5.08141010419105e-08 | 6.62123134788531e-06 |
| actl6a.S | 2368.33467461862 | 0.482184313364647 | 0.0885355920347868 | 5.44622001482964 | 5.14514889166312e-08 | 6.67059504547279e-06 |
| npm1.L | 11561.758167965 | 0.45578010443317 | 0.0837338616071056 | 5.44319939013171 | 5.23320128998718e-08 | 6.75082966408346e-06 |
| nfib.L | 1675.56579648413 | -0.528414382622204 | 0.0971416489308702 | -5.43962747634894 | 5.33920944762766e-08 | 6.81938632419771e-06 |
| atp1a1.L | 10295.1584409084 | -0.488343020458755 | 0.0897748479341817 | -5.43964185622217 | 5.33877853460046e-08 | 6.81938632419771e-06 |
| snu13.L | 3635.79025464394 | 0.507237055109265 | 0.0933369638041816 | 5.4344713437801 | 5.49591282187416e-08 | 6.98495324159376e-06 |
| LOC108713567 | 8204.9133234271 | -0.45229687898768 | 0.0832723785916515 | -5.43153548195902 | 5.5871203680515e-08 | 7.06606399488866e-06 |
| kcnh4.S | 638.555371540979 | -0.644810884481961 | 0.118865943276904 | -5.42468992131617 | 5.80552328053595e-08 | 7.30646344574769e-06 |
| ndrg4.S | 13609.5370334961 | -0.429631877861811 | 0.0792183510915197 | -5.42338829251146 | 5.8479767095759e-08 | 7.324165005197e-06 |
| stra6.L | 102.221021620446 | 1.0897779260775 | 0.201540398310249 | 5.40724308979437 | 6.40022511786763e-08 | 7.97709217589299e-06 |
| pitx2.S | 378.235811900878 | 0.837378930983464 | 0.15493198836861 | 5.40481626680731 | 6.48748587683886e-08 | 8.04697767415589e-06 |
| LOC108702869 | 15782.9944744619 | 0.607310056771238 | 0.112856629916197 | 5.381252809181 | 7.39692255167462e-08 | 9.13112927431604e-06 |
| apex1.S | 1721.3366686525 | 0.426568963681743 | 0.0793514638040656 | 5.37569117483485 | 7.62894564642027e-08 | 9.37270465131633e-06 |
| mast4.S | 1881.08372190309 | -0.481563169104696 | 0.0896111754964527 | -5.37391867071043 | 7.70436186628872e-08 | 9.42049934361369e-06 |
| dach1.S | 1009.71433685339 | -0.5465590277094 | 0.101775965354535 | -5.37021708224993 | 7.8641918586535e-08 | 9.52564084287607e-06 |
| tnnt1.L | 164.390227530781 | -0.968077538621401 | 0.180241430354363 | -5.3710045283047 | 7.82992428654918e-08 | 9.52564084287607e-06 |

| | | | | | | |
|---|---|---|---|---|---|---|
| LOC108699273 | 3715.61431533895 | -0.537215032596835 | 0.100249111435717 | -5.35880094000948 | 8.37760974368156e-08 | 1.0100108943317e-05 |
| rpe65.L | 406.656904198733 | 0.970733323602112 | 0.181300682949926 | 5.35427284557015 | 8.59011580352873e-08 | 1.03081389642345e-05 |
| bsx.L | 26.9156024935515 | 0.966737587691334 | 0.180605542213711 | 5.35275704079664 | 8.66241242635385e-08 | 1.03467703981449e-05 |
| kcne3.S | 61.3818350816973 | 1.08810786597044 | 0.203369734745247 | 5.35039231542229 | 8.77637635214748e-08 | 1.04345857090048e-05 |
| arfgap1l.S | 379.165991024374 | -0.754340822891346 | 0.141218507248055 | -5.34165696544521 | 9.21008169016427e-08 | 1.0900004936066e-05 |
| hsp90b1.L | 10258.7338233 | 0.384588752707228 | 0.0720127776976295 | 5.34056267516935 | 9.26585356525649e-08 | 1.09159370768775e-05 |
| gnat2.S | 50.9849869752778 | 1.06020669686685 | 0.198784925110172 | 5.33343610577742 | 9.63714833351432e-08 | 1.13017466820304e-05 |
| asmt.S | 44.7221419311905 | 0.942217169466975 | 0.177218193594406 | 5.31670676896412 | 1.05662139280196e-07 | 1.23196193129602e-05 |
| atp13a5l.2.L | 758.743682889927 | -0.678899658149275 | 0.127705970857841 | -5.3161152418238 | 1.06006026646402e-07 | 1.23196193129602e-05 |
| lman1l.L | 28.8228228979149 | 1.043669524116 | 0.196440693714113 | 5.31289879089354 | 1.07894969039103e-07 | 1.24829157004882e-05 |
| LOC108699962 | 10088.2088591737 | -0.481373485278691 | 0.0906232301615024 | -5.31181115946563 | 1.08541047184638e-07 | 1.25016027560878e-05 |
| snrpd2.L | 3108.40985018158 | 0.430279338602072 | 0.0810668440077237 | 5.30771049334395 | 1.11010774314126e-07 | 1.27292354546865e-05 |
| ctnna2.L | 1064.00167739161 | -0.655787840252301 | 0.123638507822972 | -5.30407436808661 | 1.13246141592944e-07 | 1.29280993499909e-05 |
| hmgb2.S | 13044.0060197566 | 0.3679544044149 | 0.0694065910927049 | 5.30143317258488 | 1.14897106599925e-07 | 1.30587900893307e-05 |
| LOC108703011 | 511.15956401668 | -0.723289243569423 | 0.136694888007665 | -5.29126768463254 | 1.21471420317643e-07 | 1.37454501938386e-05 |
| LOC108700350 | 3865.51135258902 | -0.607158074269793 | 0.114804769268232 | -5.28861368861096 | 1.23246876381845e-07 | 1.388545594171e-05 |
| LOC108700185 | 71.6077753959641 | 1.0679945350236 | 0.202124640583884 | 5.28384135619707 | 1.26502810539214e-07 | 1.41903152691814e-05 |
| stt3a.L | 2942.09240442414 | 0.441020217472557 | 0.0835416707807145 | 5.27904473720876 | 1.29859099713337e-07 | 1.45037436043468e-05 |
| dusp8.L | 1829.8717918086 | -0.465242199696808 | 0.0882109035217927 | -5.27420286066868 | 1.3333435669732e-07 | 1.48277000120295e-05 |
| znfx1.2.L | 71.79634909068 | -1.06648556714917 | 0.202547408048792 | -5.26536269821193 | 1.39912854068709e-07 | 1.54877540293362e-05 |
| LOC108702289 | 664.128539898114 | -0.663432760666509 | 0.126016926084728 | -5.26463215124331 | 1.40470327242816e-07 | 1.54877540293362e-05 |
| LOC121402164 | 63.8973425049735 | 1.06651442149288 | 0.202796256364279 | 5.25904393213808 | 1.44806294841507e-07 | 1.58978825825995e-05 |
| ache.S | 1321.14406127693 | -0.631798624135958 | 0.12018628323633 | -5.25682804329351 | 1.46561231630455e-07 | 1.6022371932482e-05 |
| cirbp.S | 80843.542236504 | 0.406881500608423 | 0.0774195005836971 | 5.25554282242559 | 1.47588505917078e-07 | 1.60665968466693e-05 |
| atp1b1.S | 4936.99714633098 | -0.407210842174358 | 0.0775894531059559 | -5.24827571111079 | 1.53529408633876e-07 | 1.65173508601871e-05 |
| ptchd4.L | 221.84832963903 | -0.811468039796152 | 0.154607092691842 | -5.24858223298685 | 1.53274219467638e-07 | 1.65173508601871e-05 |
| neurod2.L | 898.677281517633 | -0.61885885670703 | 0.117919850132987 | -5.2481313028222 | 1.53649775443601e-07 | 1.65173508601871e-05 |
| atp6v1a.S | 6555.14309214876 | -0.397667829497294 | 0.0758969899874791 | -5.23957313146277 | 1.60948455902422e-07 | 1.7230166648475e-05 |
| sall3.L | 1393.00971208503 | -0.523204105225699 | 0.0999523366510773 | -5.23453600741869 | 1.65399826973868e-07 | 1.75724441944856e-05 |
| LOC100126660 | 955.611232914183 | -0.846247595127225 | 0.161669936255762 | -5.23441534478285 | 1.65507904622481e-07 | 1.75724441944856e-05 |
| not.L | 23.7429820340257 | 0.97437297540404 | 0.186258690661068 | 5.231288655288 | 1.68332414063658e-07 | 1.7582899930536e-05 |
| tnfrsf21.L | 1108.24898480901 | -0.565844798179847 | 0.108155002520164 | -5.2317949701342 | 1.67871890824545e-07 | 1.7582899930536e-05 |
| vsnl1.S | 5345.30215367615 | -0.41707401454875 | 0.0797080562054013 | -5.23252020440673 | 1.67214368963974e-07 | 1.7582899930536e-05 |
| penk.L | 1260.52928361991 | -0.519074540109186 | 0.0991879180395055 | -5.23324362854803 | 1.66560969358293e-07 | 1.7582899930536e-05 |
| rpsa.S | 32984.9424308455 | 0.454030851427338 | 0.0868161187074538 | 5.22979900722464 | 1.69694431691244e-07 | 1.76536949098149e-05 |
| stx1b.L | 8484.5505098095 | -0.3679383069948 | 0.0703913752013265 | -5.22703677748104 | 1.72248248380672e-07 | 1.78474088683588e-05 |
| slc7a5.L | 4718.64543091536 | 0.395195980326157 | 0.075626466107585 | 5.2256306643227 | 1.73562501540033e-07 | 1.79116501589314e-05 |
| pcdh10.L | 2167.33051937905 | -0.509685770854813 | 0.0976223891597889 | -5.22099259444016 | 1.77966658596476e-07 | 1.82929872182832e-05 |
| col5a3.S | 1023.18869959423 | -0.611273292063216 | 0.11717502837281 | -5.21675394964154 | 1.82085876540905e-07 | 1.86421254553784e-05 |
| ctnnb1.L | 9909.33969763834 | 0.43493788276518 | 0.0834548634109877 | 5.21165412042261 | 1.87164258899825e-07 | 1.90111727543916e-05 |
| hes5.1.L | 789.188380392649 | 0.650570279033541 | 0.124820735120977 | 5.21203691360259 | 1.86778370826578e-07 | 1.90111727543916e-05 |
| vstm2l.L | 2180.68931774183 | -0.535020637795788 | 0.10274928442575 | -5.20704976960115 | 1.91866645358234e-07 | 1.94123900009507e-05 |
| LOC108704587 | 1379.91361116497 | -0.531666652365145 | 0.102214585047769 | -5.20147542659082 | 1.97712617180879e-07 | 1.99257247002605e-05 |
| hsp90b1.S | 5962.76010411335 | 0.42219850136319 | 0.081251739452236 | 5.19617800442759 | 2.03427480311586e-07 | 2.04219026927584e-05 |
| sema3b.S | 392.155787699807 | 0.806692640828947 | 0.155343362257145 | 5.19296498484179 | 2.06971122452078e-07 | 2.06971122452078e-05 |
| psmb7.S | 2502.16639816065 | 0.467928153383764 | 0.0901227994873759 | 5.19211737812594 | 2.07915848294471e-07 | 2.07113084401442e-05 |
| prrt1.S | 5356.75144891327 | -0.449720639480811 | 0.0866281589134864 | -5.19139094171373 | 2.08728835279179e-07 | 2.07123228853954e-05 |
| cabp1.L | 811.57183203142 | -0.552586940146266 | 0.106506508270357 | -5.18829270736747 | 2.12230830536853e-07 | 2.09791395703096e-05 |
| rdh5.L | 108.169915824299 | 0.966922991993796 | 0.186732367569907 | 5.17812206087842 | 2.24130576690104e-07 | 2.20708735824606e-05 |
| shf.L | 446.166465981769 | -0.674107983679215 | 0.130264170278936 | -5.17493016103922 | 2.279962472707e-07 | 2.23661717854907e-05 |
| mrps26.L | 825.369260173712 | 0.561659896931436 | 0.108663137687453 | 5.16881721699342 | 2.35580158540317e-07 | 2.29357286427931e-05 |
| skor1.L | 760.870481072701 | -0.706489290094055 | 0.13667787513888 | -5.16900990285503 | 2.35337429259182e-07 | 2.29357286427931e-05 |
| LOC108713150 | 295.12956230003 | -0.794604984957692 | 0.153910305942453 | -5.1627795818611 | 2.43309580393012e-07 | 2.35991999027809e-05 |
| LOC108700992 | 342.910014481601 | -0.924207309551365 | 0.179337304403542 | -5.15345824241746 | 2.55726038820527e-07 | 2.4710605998388e-05 |
| rps11.L | 10499.2705609255 | 0.528681605498468 | 0.1026249166439 | 5.15159108321566 | 2.58285728029684e-07 | 2.48648200864397e-05 |
| sp8.S | 678.210046965744 | -0.683676750769366 | 0.132757960088885 | -5.14979855303308 | 2.60766381943029e-07 | 2.50103072642443e-05 |
| hmgn2.S | 13292.6497785057 | 0.36358330029801 | 0.0707057289244165 | 5.14220425740432 | 2.71533669378944e-07 | 2.59320092429045e-05 |
| LOC108713861 | 1046.98880457224 | -0.527855245060552 | 0.102677130088286 | -5.14092324753022 | 2.73391725351552e-07 | 2.59320092429045e-05 |
| snrpe.S | 2859.79068969478 | 0.429480837096535 | 0.0835307748786158 | 5.14158808799083 | 2.72425872151257e-07 | 2.59320092429045e-05 |
| rab11fip5.L | 599.686882978615 | -0.539859657364567 | 0.105043553822509 | -5.13938873656884 | 2.75633643075866e-07 | 2.60488937412357e-05 |
| LOC108703510 | 879.545886969384 | -0.687438793958867 | 0.133884611788981 | -5.13456165554227 | 2.82802374524933e-07 | 2.66288367253404e-05 |
| thy1.L | 1090.92235378678 | -0.551149496865206 | 0.107406588055647 | -5.13143101221741 | 2.87547616342092e-07 | 2.67824133632706e-05 |
| bsx.S | 22.6454863728854 | 0.901107892648139 | 0.175562091600454 | 5.13270196563212 | 2.85611982138846e-07 | 2.67824133632706e-05 |
| psma6.S | 2889.45993667797 | 0.427993249598542 | 0.0834023248287225 | 5.13167049572637 | 2.87181922219291e-07 | 2.67824133632706e-05 |
| LOC108716110 | 219.954448568976 | -0.984461056363708 | 0.192498835329757 | -5.11411435127539 | 3.15216368605944e-07 | 2.92538932015588e-05 |
| vsx2.L | 454.368916999019 | -0.643007389291701 | 0.125763516330132 | -5.11282928511471 | 3.17369092831824e-07 | 2.93481096597171e-05 |
| ogdh.S | 1735.96761100985 | -0.39832501259956 | 0.0779802252693665 | -5.10802592867139 | 3.25541987254585e-07 | 2.98896201820936e-05 |
| rab6b.L | 1113.29059679551 | -0.500455463571506 | 0.0979620216035761 | -5.1086681897675 | 3.24437529569 56e-07 | 2.98896201820936e-05 |
| tgm1.S | 298.78335825921 | 0.703631527259928 | 0.13784999304259 | 5.10432762258127 | 3.31972715334955e-07 | 3.02646503732927e-05 |
| nif.S | 7252.86648180764 | -0.535990713892213 | 0.104995940686749 | -5.10487082059029 | 3.31020561305045e-07 | 3.02646503732927e-05 |
| pfkp.L | 4950.03508816896 | -0.38057422929549 | 0.0747220200952679 | -5.09320048909641 | 3.5206904701136e-07 | 3.18715137294495e-05 |
| rps5.L | 13688.2596644909 | 0.54608742907231 | 0.107210614393336 | 5.09359481020057 | 3.51337226617162e-07 | 3.18715137294495e-05 |
| hes5.4.S | 142.774246662364 | 0.900620429311284 | 0.176855025613109 | 5.09242203431356 | 3.53518102836842e-07 | 3.18907938922746e-05 |
| nif.L | 7744.95942394591 | -0.472535652998239 | 0.0928896265090212 | -5.08706591636847 | 3.63645471796053e-07 | 3.25068968150627e-05 |
| hspa5.S | 9908.2346842109 | 0.34897239102552 | 0.0686033253519551 | 5.08681451278331 | 3.64127642618338e-07 | 3.25068968150627e-05 |
| r3hdm1.S | 5391.89385266412 | -0.427824335277248 | 0.0841011735391626 | -5.08701980333279 | 3.63733866526557e-07 | 3.25068968150627e-05 |
| clstn3.L | 6074.17550506355 | -0.462572172974183 | 0.0909542729467353 | -5.08576626460501 | 3.66144752538087e-07 | 3.25742572947678e-05 |
| mtss2.L | 5896.48948953254 | -0.40953754851877 | 0.0806388918792551 | -5.07866042023484 | 3.80105404645207e-07 | 3.37000668035957e-05 |
| bhlhe22.L | 565.684884744449 | -0.669859651783412 | 0.131973575788747 | -5.07571040475308 | 3.86050893473667e-07 | 3.41099762041802e-05 |
| nop58.L | 2215.95012770835 | 0.405713034400593 | 0.0799844514232743 | 5.07239878727901 | 3.92832080906582e-07 | 3.45906747009891e-05 |
| stac2.S | 442.202069559975 | -0.626075193740654 | 0.123460165585394 | -5.0710704199373 | 3.95584350020604e-07 | 3.47145450018081e-05 |
| atp1a2.S | 16310.9553297295 | -0.378750155873101 | 0.074699521743245 | -5.0703156731703 | 3.97156403654583e-07 | 3.47343566586042e-05 |
| map1b.L | 9839.97461603137 | -0.375376967120647 | 0.0740573503847604 | -5.06873342308898 | 4.00471644074556e-07 | 3.49059743821742e-05 |
| prrt2.L | 1414.70873719715 | -0.578525165783424 | 0.114275290350854 | -5.06255695353916 | 4.13670427426621e-07 | 3.5935006826959e-05 |
| kiaa1549l.L | 1170.86434515935 | -0.55293282549817 | 0.109235727774751 | -5.06183129606042 | 4.15248395356888e-07 | 3.59510355711668e-05 |
| hectd1.L | 2332.2365909731 | -0.41127038941016 | 0.0812947756784806 | -5.05900146691747 | 4.21457624615732e-07 | 3.63665776424277e-05 |
| LOC108698757 | 8718.33399388807 | 0.509282807755824 | 0.10083495187048 | 5.05065751813898 | 4.40291863006743e-07 | 3.78651002185799e-05 |
| snrpb.S | 5201.57469852904 | 0.444911643205909 | 0.0881728416405182 | 5.04590342023704 | 4.51383388836904e-07 | 3.86900047574489e-05 |
| mmut.S | 1892.38329753908 | -0.423851054007031 | 0.0840698123793452 | -5.04165576217184 | 4.61520966533074e-07 | 3.94279501210374e-05 |
| meis2.L | 3534.93707279169 | -0.529005862412658 | 0.1051165921992 | -5.03256290320152 | 4.83965729891626e-07 | 4.1208963139287e-05 |
| cadm2.L | 1467.06387386927 | -0.511184204152501 | 0.101790695260894 | -5.02191485029466 | 5.11588641559138e-07 | 4.34177202375847e-05 |
| LOC108713037 | 835.510455466161 | -0.562908415986321 | 0.112293369394723 | -5.01283752567474 | 5.36331819121113e-07 | 4.5368396502704e-05 |
| pc.1.L | 740.263097781788 | -0.516506824586789 | 0.103071618950015 | -5.01114496743564 | 5.41071335458911e-07 | 4.54912590380047e-05 |
| phyhipl.S | 1778.90796904749 | -0.56044816788991 | 0.111842241676665 | -5.01105985974567 | 5.41310718010366e-07 | 4.54912590380047e-05 |
| efnb3.L | 1189.59478555458 | -0.460897541141179 | 0.0921327687920052 | -5.00253652619168 | 5.65808571746043e-07 | 4.73956530878179e-05 |
| stxbp1.L | 17402.4122412017 | -0.436977435083371 | 0.0876170234785928 | -4.98735768158269 | 6.12106789901216e-07 | 5.11079455645676e-05 |
| slc45a4.S | 502.222751594037 | -0.563939577084673 | 0.113393819810011 | -4.97328318271261 | 6.58283909642294e-07 | 5.46100478095536e-05 |
| LOC108697935 | 2466.74934864179 | -0.420907959347705 | 0.0846247970579357 | -4.97381351543501 | 6.56484646524838e-07 | 5.46100478095536e-05 |
| myg1.S | 1133.28214285374 | 0.43156803887279 | 0.0868457921373793 | 4.96936038294294 | 6.71741185313367e-07 | 5.55478287855285e-05 |
| sptbn1.S | 8649.14467267107 | -0.317772935645128 | 0.0639598905456517 | -4.96831581377263 | 6.75369066496175e-07 | 5.5669399091378e-05 |
| LOC108698099 | 2357.49746181364 | -0.470369355922586 | 0.0947328331161526 | -4.96521996070637 | 6.86232479411107e-07 | 5.63847069070273e-05 |
| camk2g.L | 5516.02952203027 | -0.369176925521928 | 0.0744354450820342 | -4.95969259154779 | 7.06048190195086e-07 | 5.78287089112165e-05 |
| tlcd3b.L | 1146.9964068005 | -0.501561865279954 | 0.101145784752565 | -4.95880146174096 | 7.09294079202708e-07 | 5.79107191247781e-05 |
| LOC108718312 | 2962.01386430818 | -0.446395901334931 | 0.0900446822730384 | -4.95749321410613 | 7.14085356512194e-07 | 5.81179879656612e-05 |
| LOC108696074 | 2017.33588469379 | -0.475607422768575 | 0.0959519675066951 | -4.95672402689803 | 7.16916939108301e-07 | 5.81649592106735e-05 |
| rgma.L | 5596.39551415001 | -0.405596088898373 | 0.0818630957339389 | -4.9545657327276 | 7.24920078304301e-07 | 5.862989975000 31e-05 |
| camk2g.S | 6352.45543423111 | -0.360966021327559 | 0.0729297244741255 | -4.94950479972847 | 7.44025430649121e-07 | 5.99870503460853e-05 |
| hmgb3.L | 15900.1322214272 | 0.388489003157383 | 0.0785095233309097 | 4.948304188779 | 7.48628499955129e-07 | 6.01701411178889e-05 |

| | | | | | | |
|---|---|---|---|---|---|---|
| fzd10.L | 970.494599682058 | 0.485053106437572 | 0.0980508973471413 | 4.94695224175543 | 7.53844622447157e-07 | 6.02142144245717e-05 |
| vamp2.S | 4084.04615309588 | -0.432588451580894 | 0.0874389913966442 | -4.94731749155899 | 7.52431963261974e-07 | 6.02142144245717e-05 |
| herc6.S | 49.2627027158805 | -1.00590633328806 | 0.203366368763347 | -4.94627670939345 | 7.56464082086696e-07 | 6.02369546846814e-05 |
| grid2ip.L | 358.647810763374 | -0.648013645425795 | 0.131238201359869 | -4.93769069303894 | 7.90530925420581e-07 | 6.27559934641569e-05 |
| LOC108705545 | 208.025333117862 | -0.98254432289268 | 0.199059833622678 | -4.93592456605341 | 7.9771938182815e-07 | 6.31323927949886e-05 |
| LOC108697306 | 2476.69069067058 | 0.970619687736 15 | 0.196679529838847 | 4.93503156394285 | 8.01377986296056e-07 | 6.32279879095971e-05 |
| kcna1.L | 799.764409029893 | -0.510465291200837 | 0.103587157724613 | -4.9278820117636 | 8.31257872988778e-07 | 6.53855278143612e-05 |
| LOC108710311 | 1535.01491328221 | -0.440384528306626 | 0.0893844445032748 | -4.92685870291996 | 8.35621363195074e-07 | 6.55289701228964e-05 |
| atp2b1.L | 4547.10571731994 | -0.428048669141914 | 0.0869390532659908 | -4.92354877424643 | 8.49886813184061e-07 | 6.64456963034811e-05 |
| clstn3.S | 4193.93570310423 | -0.469299504228783 | 0.0953423927989168 | -4.92225431365631 | 8.55529357578053e-07 | 6.66847656359933e-05 |
| crmp1.L | 6236.4650409342 | -0.502793433562853 | 0.102254908765951 | -4.91705913809659 | 8.78540178184876e-07 | 6.80670768683778e-05 |
| mast1.L | 2378.51351601651 | -0.41113812819057 | 0.0836080695958919 | -4.91744553100855 | 8.76808426262339e-07 | 6.80670768683778e-05 |
| rpl18a.L | 13437.2716990946 | 0.539346563657398 | 0.109837428141053 | 4.91040779801191 | 9.08871834258125e-07 | 7.02062674367055e-05 |
| LOC108695983 | 4680.06973766618 | -0.418711539337961 | 0.0852992366438702 | -4.90873723860049 | 9.16646957296918e-07 | 7.05954970097328e-05 |
| plcb4.S | 870.939120497812 | -0.487946595941962 | 0.0995015290631081 | -4.90391052817375 | 9.39472945988937e-07 | 7.21381012098648e-05 |
| psma3.L | 3622.059572257 | 0.445469216259332 | 0.0908771612327017 | 4.9018830497869 | 9.49223435953619e-07 | 7.26705182421465e-05 |
| LOC108714603 | 2355.19276561847 | -0.44482726779003 | 0.0907982341641817 | -4.89907399504809 | 9.62893795232445e-07 | 7.34989938372695e-05 |
| dnm1l.L | 4210.74417013699 | -0.388442624319886 | 0.0793330333322717 | -4.89635411636116 | 9.76310654684393e-07 | 7.4226669384166e-05 |
| odc1.L | 12968.4161600069 | 0.335524462108513 | 0.0685306333751597 | 4.89597783624352 | 9.78180914364979e-07 | 7.4226669384166e-05 |
| pa2g4.L | 4526.60520495483 | 0.460234268455351 | 0.0940310894402001 | 4.89449404135601 | 9.85607668302609e-07 | 7.43528591877407e-05 |
| agl.S | 1407.55757654666 | -0.420863183234018 | 0.0859866172563322 | -4.89451959692047 | 9.85461570642919e-07 | 7.43528591877407e-05 |
| tcf4.S | 1054.12694009833 | -0.452256272458237 | 0.0924426567359935 | -4.89228986299944 | 9.96695080843179e-07 | 7.49700673054053e-05 |
| snrpa.S | 2366.05930406446 | 0.423068445710034 | 0.0865661545170826 | 4.88722697768153 | 1.02266169158809e-06 | 7.66996268691069e-05 |
| hmgn1.L | 21153.1584822154 | 0.399608907207044 | 0.0818266176796044 | 4.88360534186724 | 1.04163459796958e-06 | 7.78961525438119e-05 |
| LOC108700925 | 2066.85830572659 | 0.714503272685661 | 0.146323689762092 | 4.88303209034282 | 1.04466861647001e-06 | 7.78972552165496e-05 |
| LOC108700932 | 1084.32961157414 | -0.542921331032599 | 0.111261168629474 | -4.87970185573599 | 1.06246331798563e-06 | 7.89958317118995e-05 |
| LOC108711439 | 6834.88417645005 | 0.538671161403219 | 0.110408333723461 | 4.87889947467576 | 1.06679415435343e-06 | 7.90899114434441e-05 |
| znf326.S | 7191.34538640883 | 0.369563791717026 | 0.0757956118782724 | 4.87579402763506 | 1.0837164221592e-06 | 8.01142799189322e-05 |
| hes5.3.L | 462.670074128812 | 0.63073410069929 | 0.129409134585895 | 4.87395347104067 | 1.09386763718601e-06 | 8.06336715411403e-05 |
| LOC108706905 | 6412.41248217859 | 0.440965756908231 | 0.0905480351723806 | 4.86996494257157 | 1.1161805012839e-06 | 8.20440368465088e-05 |
| hes1.S | 270.579663279436 | 0.665474055012159 | 0.136858429080441 | 4.86249958795752 | 1.15912620334825e-06 | 8.49586819499567e-05 |
| slc39a7.L | 1872.85723680941 | 0.408428169417801 | 0.0840310767313911 | 4.86044193772929 | 1.17123992553536e-06 | 8.56033713280799e-05 |
| eda2r.L | 132.837682766595 | 0.893184139961748 | 0.183856313844845 | 4.85805530026835 | 1.18544304266336e-06 | 8.63966963297027e-05 |
| rps5.S | 13642.4135338574 | 0.558273802222991 | 0.114939393435584 | 4.85711456739038 | 1.19108684659296e-06 | 8.65634947664741e-05 |
| nfix.L | 91.1776953790741 | -0.908425640868287 | 0.187069853662211 | -4.856077144897 | 1.19734070005936e-06 | 8.67735675885718e-05 |
| LOC108717266 | 1316.1290131724 | -0.548746071584348 | 0.113037366182704 | -4.85455464963153 | 1.20657593233837e-06 | 8.69052790820542e-05 |
| LOC108701642 | 3679.22534753193 | -0.474559878118035 | 0.0977644749301794 | -4.85411371008694 | 1.2092633794751e-06 | 8.69052790820542e-05 |
| XB5962511.S | 177.178029149657 | -0.779654212803729 | 0.160583864154749 | -4.85512175776517 | 1.20312795688334e-06 | 8.69052790820542e-05 |
| pcna.S | 4444.13246089961 | 0.366827182142339 | 0.0756414274913427 | 4.84955393239138 | 1.23739423088826e-06 | 8.86799198803255e-05 |
| slc4a4.S | 936.729129561915 | -0.601561999142867 | 0.124062091211709 | -4.8488784387514 | 1.24161475488119e-06 | 8.87253843045878e-05 |
| LOC108698154 | 91.4828352189828 | 0.901075373517696 | 0.18585184524978 | 4.84835311861807 | 1.24490655497135e-06 | 8.87253843045878e-05 |
| pnrc2.S | 1781.53500460803 | 0.400764727828323 | 0.0826823331871087 | 4.84704183324628 | 1.25316010376157e-06 | 8.90675776227233e-05 |
| LOC108696239 | 27.3693385635712 | 0.943419740245968 | 0.194839450599629 | 4.84203654517883 | 1.28515120089921e-06 | 9.10903873164824e-05 |
| LOC108717002 | 1889.95309493486 | 0.511086060705927 | 0.105601138159859 | 4.83977795705428 | 1.29984274888122e-06 | 9.18792956743438e-05 |
| kif1a.L | 5691.53537451384 | -0.523121470323805 | 0.108114407538611 | -4.83859165705533 | 1.30762390509101e-06 | 9.21767670801859e-05 |
| h2az1.L | 3393.09833863156 | 0.460404686447847 | 0.0952578985936351 | 4.83324420594148 | 1.34325842318359e-06 | 9.4430701139337e-05 |
| mapk8ip2.L | 4333.47877125022 | -0.473849333329129 | 0.0980667941739876 | -4.83190398259005 | 1.35233476266275e-06 | 9.48104262953777e-05 |
| kcnd1.S | 1018.77017661663 | -0.466528077547488 | 0.0965887432719137 | -4.83004604619539 | 1.36501480172953e-06 | 9.54400593079182e-05 |
| LOC108712865 | 5915.32428589461 | 0.547496501463952 | 0.113406921005789 | 4.82771683252032 | 1.3810727991122e-06 | 9.60422593452687e-05 |
| ndrg4.L | 8177.71907937194 | -0.32361451335677 | 0.0670319105436624 | -4.82776800977467 | 1.38071803044071e-06 | 9.60422593452687e-05 |
| dkc1.L | 7290.56306900721 | 0.45390327328347 | 0.0940593952424438 | 4.82570903324975 | 1.39506056824335e-06 | 9.67542007007485e-05 |
| MGC68699 | 4259.52637415306 | -0.520367240714973 | 0.107980201973181 | -4.8190986051704 | 1.44208272179902e-06 | 9.97472767356966e-05 |
| LOC108697444 | 2393.79296115904 | -0.42985539447635 | 0.0892959565610735 | -4.81381197220843 | 1.48078141505936e-06 | 0.000102006026827907 |
| LOC108701562 | 786.063403055812 | -0.486021811145772 | 0.100969291994679 | -4.81356065338545 | 1.48264573877772e-06 | 0.000102006026827907 |
| LOC108703406 | 2839.42739582784 | -0.418464829844819 | 0.0869691251747333 | -4.81164814529368 | 1.49690715248725e-06 | 0.000102440860833345 |
| LOC108703019 | 330.404800109946 | -0.64208805882422 | 0.133443528502617 | -4.81168375888402 | 1.4966403838101e-06 | 0.000102440860833345 |
| cpne4.L | 1210.82814071339 | -0.50222515184165 | 0.104407554968354 | -4.81023764988919 | 1.50750950861374e-06 | 0.000102893506143477 |
| nos1.L | 1082.1332076383 | -0.591117396574984 | 0.122925989749343 | -4.80875294786769 | 1.51895257514706e-06 | 0.000103401005631647 |
| ppib.S | 6067.23294633452 | 0.398066728819116 | 0.0828175530938777 | 4.80655022936851 | 1.53556916323464e-06 | 0.000104257064240667 |
| hes5.4.L | 198.73822874174 | 0.734017515803976 | 0.153082169413116 | 4.79492496492595 | 1.62735732308766e-06 | 0.000110198999831133 |
| chchd3.S | 832.269136785467 | 0.512522511544595 | 0.106917939374319 | 4.79360633532468 | 1.63809568174925e-06 | 0.000110635781646939 |
| ccdc136.S | 573.109679000149 | -0.566582093906792 | 0.118243256129787 | -4.79166518625717 | 1.65402759128768e-06 | 0.000111420135392225 |
| XB5796436.L | 1768.47608551026 | 0.730669030243214 | 0.15261150068008 | 4.78777174057753 | 1.68643302542175e-06 | 0.000112720134859796 |
| map7d2.S | 2021.97931873629 | -0.577010975918649 | 0.120493946424927 | -4.78871340045412 | 1.67854003226591e-06 | 0.000112720134859796 |
| nkx2-4.S | 47.4274883294456 | 0.85414186965367 | 0.178398894128223 | 4.78782042807822 | 1.68602405375765e-06 | 0.000112720134859796 |
| LOC108704224 | 175.110915478654 | -0.836516799895029 | 0.174774164526122 | -4.78627262881295 | 1.69907222844606e-06 | 0.000113271481896404 |
| wasf1.L | 1455.90829475382 | -0.489204643468917 | 0.102324483557119 | -4.78091485500474 | 1.7449926250687e-06 | 0.000116033014759723 |
| rpl28.L | 6607.11574958801 | 0.528913843818272 | 0.110692016807826 | 4.77824742082823 | 1.76829708562136e-06 | 0.00011728037225972 |
| LOC108714516 | 156.290379983051 | -0.812161376050594 | 0.170007247895233 | -4.77721618404815 | 1.77738657219203e-06 | 0.000117580957852703 |
| LOC108697145 | 48.8333877667923 | -0.970637221883208 | 0.203202889210011 | -4.77669006408688 | 1.78204115764771e-06 | 0.000117587370504632 |
| lin28a.L | 212.421451900937 | 0.948491657784831 | 0.198699336143893 | 4.77350189583903 | 1.81049844189458e-06 | 0.00011885714962056 |
| mvd.L | 830.660070053004 | 0.560530616966795 | 0.117420492592249 | 4.77370350432168 | 1.80868604502595e-06 | 0.00011885714962056 |
| pomp.S | 1071.17038999686 | 0.452680475310324 | 0.0948427091813569 | 4.77296019080091 | 1.81537684795442e-06 | 0.000118874930652853 |
| rps15.L | 7835.09253433281 | 0.44333002482871 | 0.0929002381191617 | 4.77210859524448 | 1.82307156529065e-06 | 0.000119076573125313 |
| wnt8b.S | 91.8052771308071 | 0.95863388030071 | 0.201161040893199 | 4.76550467249605 | 1.88381531462835e-06 | 0.000122733422013665 |
| cabp7.L | 615.032367628977 | -0.593817189179276 | 0.124776409585752 | -4.75905013736735 | 1.94506120806284e-06 | 0.00012640448153154 |
| csnk2b.L | 3105.54128533933 | 0.43577278619379 | 0.0915912725477239 | 4.75779814028382 | 1.95716066317572e-06 | 0.00012687121886918 |
| psat1.S | 1376.45566435824 | 0.631259227232515 | 0.132788882434806 | 4.75385601307728 | 1.99573194823665e-06 | 0.000127135516702483 |
| LOC108707480 | 69.3442496652664 | -0.967131681678333 | 0.203355885828592 | -4.75585782893704 | 1.97605504856514e-06 | 0.000127135516702483 |
| spdef.S | 34.8523937648045 | 0.942575275135183 | 0.198273997967896 | 4.75390260344578 | 1.99527185516268e-06 | 0.000127135516702483 |
| npy.L | 773.529272037343 | 0.683869931926515 | 0.143825272384817 | 4.75486623864521 | 1.98577850139992e-06 | 0.000127135516702483 |
| prpf19.L | 2337.90931651028 | 0.409723244281548 | 0.0861825473482987 | 4.75413244198608 | 1.9930036252925e-06 | 0.000127135516702483 |
| rpl36a.L | 16590.9500960751 | 0.490748312784641 | 0.103193715463917 | 4.7556027087351 | 1.97855235719581e-06 | 0.000127135516702483 |
| LOC108701244 | 62.0224360189754 | -0.966918844436319 | 0.203285035080582 | -4.75646839450349 | 1.97009066221445e-06 | 0.000127135516702483 |
| mmp16.L | 447.58490853488 | -0.625885115815484 | 0.131737357074469 | -4.75100707737503 | 2.02406047717746e-06 | 0.000128622561999469 |
| rpl37.S | 9513.57622605007 | 0.547675693351108 | 0.115336049370372 | 4.74852135425922 | 2.04909255453599e-06 | 0.000129893336380905 |
| rps26.L | 7325.18429421703 | 0.514537596541081 | 0.10840259735057 | 4.74654306369694 | 2.06922685395335e-06 | 0.000130848168705874 |
| rpl23a.L | 14604.0060619818 | 0.472688515428106 | 0.0996269109128275 | 4.74458668945084 | 2.08932489849947e-06 | 0.00013179604494202 |
| LOC121396584 | 162.023335325219 | -0.836334297174718 | 0.176460567153963 | -4.73949682166108 | 2.142495957748e-06 | 0.000134647844576526 |
| sez6l2.S | 6241.24974025052 | -0.406014609848366 | 0.0856704145021585 | -4.73926281561455 | 2.14497147755629e-06 | 0.000134647844576526 |
| rpl8.L | 14873.249470527 | 0.497058317253581 | 0.104955315341856 | 4.73590418583932 | 2.18080606860374e-06 | 0.000136565040218389 |
| psmb1.S | 2364.78457747071 | 0.41718867893006 | 0.08813358184126 | 4.73359496135618 | 2.20577691370423e-06 | 0.000137794296304041 |
| rpl4.L | 35808.9750636693 | 0.37650775432546 | 0.0795689848946391 | 4.73184061382724 | 2.22493088610844e-06 | 0.000138655113192265 |
| pygm.S | 3679.84956353884 | -0.504768746781085 | 0.106695618451978 | -4.73092291984112 | 2.23501379651323e-06 | 0.00013894784566275 |
| khdrbs1.S | 11407.1712012944 | 0.356738443907631 | 0.0754558164005858 | 4.72777926109433 | 2.26988754879776e-06 | 0.000140439085752955 |
| flot1.L | 1820.70032636618 | 0.403586239330592 | 0.0853597083642418 | 4.72806488054567 | 2.26669761303315e-06 | 0.000140439085752955 |
| ahcy.L | 2008.92019687743 | 0.596292223058663 | 0.126179424303869 | 4.72574848354559 | 2.29269285572866e-06 | 0.000141510707363156 |
| tmem255a.L | 908.132861390172 | -0.49973067365379 | 0.105760162096413 | -4.72513150271296 | 2.29966491520463e-06 | 0.000141602278788257 |
| cct5b | 15981.9635362272 | 0.443166680069704 | 0.0938377737261409 | 4.72268962137844 | 2.32745912679758e-06 | 0.000142972489217565 |
| LOC108714718 | 48.0470004467701 | -0.919121561095173 | 0.194658104473142 | -4.72172254827433 | 2.33855555967203e-06 | 0.000143312906032158 |
| rpl12.L | 12290.99702004 | 0.371796756472834 | 0.0788445105319309 | 4.71556933769097 | 2.41037578697717e-06 | 0.000147363111409907 |
| app.L | 9607.52887056634 | -0.465414553521786 | 0.09877814118607 | -4.71171605310001 | 2.45639485216617e-06 | 0.00014982266474205 |
| dbi.S | 2630.70811317163 | 0.515327525431064 | 0.10945517557032 | 4.70811473962682 | 2.50018363849657e-06 | 0.000152133815738707 |
| lrit1.S | 74.5374423969947 | 0.956779666826218 | 0.203320538713191 | 4.70576987884079 | 2.52909680658617e-06 | 0.000153531053199819 |
| dhcr24.L | 719.160313927973 | 0.582300694766384 | 0.123795600122077 | 4.70372690299305 | 2.55454899336084e-06 | 0.000154712122133122 |
| eif2b3.L | 1536.69137139661 | 0.535948053799251 | 0.113972618878288 | 4.70242817155576 | 2.57085672777672e-06 | 0.000155335137181825 |
| ptmap12.S | 14520.4336030361 | 0.423735481999333 | 0.0901455437191592 | 4.70057048321152 | 2.59435688628875e-06 | 0.000156388802958528 |
| lrp1.L | 5894.67936567686 | -0.403335488836672 | 0.08586572236187 | -4.69728172945272 | 2.63646683925022e-06 | 0.000158556746975887 |

| | | | | | | |
|---|---|---|---|---|---|---|
| LOC108708306 | 3079.61627623842 | 0.354175520319777 | 0.0754547549337322 | 4.69387940668325 | 2.68072099524152e-06 | 0.000160752077357294 |
| LOC108701205 | 1472.8288067638 | 0.517064116177784 | 0.110165520173978 | 4.69352039877099 | 2.68543198996099e-06 | 0.000160752077357294 |
| rplp1.L | 17912.2366467621 | 0.485948633901625 | 0.103629149262724 | 4.68930448005159 | 2.74135226173975e-06 | 0.000163719648965013 |
| rpl15.S | 13357.5895117051 | 0.473202498321081 | 0.100960342935312 | 4.68701357942372 | 2.77220579506926e-06 | 0.000165179929590732 |
| sf3b1.S | 5242.56208116445 | 0.365200401526728 | 0.0779722673694623 | 4.68372171090356 | 2.81712426303369e-06 | 0.000167469599046703 |
| rpl36.S | 6950.59201750179 | 0.51641480655371 | 0.11027323683113 | 4.68304750448683 | 2.82640977146197e-06 | 0.000167635338169469 |
| LOC108709337 | 1819.58537691146 | -0.54270746213142 | 0.115900620305763 | -4.68252422376756 | 2.83363689068219e-06 | 0.000167678513255965 |
| LOC443585 | 1831.07198009539 | -0.493705036817362 | 0.105474764746545 | -4.68078822459315 | 2.85774028013873e-06 | 0.000168717847202698 |
| epha10.L | 952.286291431694 | -0.576215702937764 | 0.123172240542797 | -4.67812958827807 | 2.89503571238255e-06 | 0.000170343506561141 |
| rpl10a.S | 26730.3065414961 | 0.500827280277512 | 0.107062744761553 | 4.67788567716006 | 2.89848059613725e-06 | 0.000170343506561141 |
| hsf2.2.L | 1076.23560531907 | 0.486772381142408 | 0.104080206333147 | 4.67689677309357 | 2.91248774408492e-06 | 0.000170777690448616 |
| LOC108717631 | 1261.5387762354 | -0.435663052036822 | 0.0932981347559181 | -4.66957944203903 | 3.01816969734071e-06 | 0.000176573193177756 |
| LOC108713772 | 1301.5667147525 | -0.396504717094107 | 0.0850287638677655 | -4.66318336358201 | 3.11355094378503e-06 | 0.000181690516497719 |
| bzw1.S | 5671.8492061189 | 0.411974079414804 | 0.0883538243152356 | 4.66277586293187 | 3.11972476001898e-06 | 0.000181690516497719 |
| znf706l.L | 1554.76542400556 | 0.437658507869133 | 0.0938884234048193 | 4.66147467384853 | 3.13951706421005e-06 | 0.000182431396974368 |
| nrp2.L | 1502.12016292059 | -0.441619638460253 | 0.0947554589587047 | -4.6606247630832 | 3.15250997224889e-06 | 0.000182774735469711 |
| atp1b2.S | 82.2360182582485 | 0.933477176649093 | 0.200357844315026 | 4.65901986438516 | 3.17718545300535e-06 | 0.00018379234234874 |
| dpysl2.L | 8329.14968940124 | -0.435712514890223 | 0.0935335219000027 | -4.65835676920245 | 3.18743459949352e-06 | 0.00018397273527278 |
| rplp2.L | 9668.66880689557 | 0.529572484280613 | 0.113717518741618 | 4.65691205841271 | 3.20987473220377e-06 | 0.000184854393059949 |
| rpl15.L | 12100.8651553585 | 0.413205660740702 | 0.0888405251781306 | 4.65109430535446 | 3.30178287652215e-06 | 0.000189723826757843 |
| rps12.S | 12448.4979884787 | 0.568043865371562 | 0.122245722174076 | 4.64673818657372 | 3.37224715158016e-06 | 0.000193342170023929 |
| dlx5.S | 48.1161376993256 | 0.871834022544163 | 0.187645693992767 | 4.64617121764473 | 3.38152377899867e-06 | 0.000193444154097929 |
| h2az1.S | 2903.09943569832 | 0.43197916305946 | 0.0930207974351158 | 4.64389872985958 | 3.41895195729516e-06 | 0.000195152567473927 |
| rcan2.L | 602.822346767733 | -0.490216277647918 | 0.105694787569871 | -4.63803645306402 | 3.51734760595805e-06 | 0.000200325757690326 |
| snrpf.L | 2532.54209783575 | 0.462375810805725 | 0.0997376452041933 | 4.63592066826022 | 3.55352244245031e-06 | 0.000201437247922294 |
| rpl18a.S | 10952.0547295272 | 0.531345322944646 | 0.114628876432137 | 4.63535314558557 | 3.56328622751037e-06 | 0.000201437247922294 |
| tcp1.L | 9858.35441483379 | 0.318008318009221 | 0.0686091098194866 | 4.63507424664033 | 3.56809388761583e-06 | 0.000201437247922294 |
| slc7a3.L | 2549.23468508434 | 0.608297312165126 | 0.131205504778764 | 4.63621797776817 | 3.54841873800849e-06 | 0.000201437247922294 |
| rpl26.S | 10260.3536870658 | 0.525497107541845 | 0.113409704705749 | 4.63361675180525 | 3.59331952944962e-06 | 0.00020241843637511 |
| aadac.L | 149.532871619896 | 0.93763840337646 | 0.202390123371905 | 4.63282687788814 | 3.60706164987281e-06 | 0.000202749870515727 |
| psmb7.L | 2779.87911041 | 0.350161040779639 | 0.0756053548407096 | 4.63143174868211 | 3.63145710643856e-06 | 0.000203677376839 38 |
| dctn1.L | 7488.52731980747 | -0.309337811598512 | 0.0668354028699324 | -4.62835261426509 | 3.68586041637399e-06 | 0.000205863833788062 |
| atp6v1a.L | 2840.98653009228 | -0.469853949103546 | 0.10151711850251 | -4.62832235621353 | 3.68639888411181e-06 | 0.000205863833788062 |
| rab6b.S | 1872.41443944486 | -0.46164549631208 | 0.099761945731191 | -4.62747085502909 | 3.70158303134019e-06 | 0.000206265317945091 |
| aplp1.S | 4031.79184388713 | -0.456388666284944 | 0.0986840434140587 | -4.62474631658563 | 3.75057146520189e-06 | 0.000208544706470278 |
| ccdc12.L | 465.808861635319 | 0.572201294362223 | 0.123786362405767 | 4.6224905816892 | 3.79160031859083e-06 | 0.000210372662837943 |
| emilin1.S | 148.066198485764 | -0.794801107358044 | 0.171980923421021 | -4.6214492372059 | 3.81068583709719e-06 | 0.000210977885401518 |
| LOC108704484 | 3143.76391790549 | -0.392136454691499 | 0.0848764673615173 | -4.62008454029147 | 3.83583714320185e-06 | 0.000211915628039845 |
| c21orf62.S | 34.4987946662087 | 0.931707763495068 | 0.201728224797555 | 4.61862867444596 | 3.862844088736e-06 | 0.000212308245669113 |
| racgap1.S | 724.673369491401 | 0.471453276130258 | 0.102082143126853 | 4.61837165334982 | 3.86763083915672e-06 | 0.000212308245669113 |
| rpl39.L | 6137.11565024634 | 0.521085610080197 | 0.112810827456597 | 4.61910990131403 | 3.85389701795776e-06 | 0.000212308245669113 |
| hnrnpdl.L | 24419.5544721593 | 0.322022206888323 | 0.0697342888006135 | 4.61784600412372 | 3.87743819763383e-06 | 0.000212394703819433 |
| aatf.S | 843.675074684061 | 0.476369049370018 | 0.103182648136963 | 4.61675541354291 | 3.89786211555486e-06 | 0.000213061107163804 |
| cdh24.S | 318.196755796208 | -0.601789079715433 | 0.130420722654012 | -4.61421365768612 | 3.94586355724769e-06 | 0.000215228921304419 |
| tcf4.L | 746.988127211487 | -0.45554338785693 | 0.0987596462518337 | -4.61264701875612 | 3.97573144910431e-06 | 0.000215944992393455 |
| grin1.L | 5272.90669086598 | -0.559739897959489 | 0.121347511075859 | -4.61270192521356 | 3.97468100608318e-06 | 0.000215944992393455 |
| lmo3.S | 1393.62384542819 | 0.624032937407156 | 0.135362916174422 | 4.6100730912376 | 4.02527425049738e-06 | 0.000216033018440465 |
| rpsa.L | 19913.7866046729 | 0.513326508031588 | 0.111319652980306 | 4.61128376067074 | 4.00189810390862e-06 | 0.000216033018440465 |
| scrt2.S | 387.920904546 16 | -0.618278770136546 | 0.134084456615068 | -4.61111441060997 | 4.00516014592588e-06 | 0.000216033018440465 |
| fnbp1.L | 2417.25338450566 | -0.359052298091798 | 0.0778664687816241 | -4.61112856034036 | 4.00488749451427e-06 | 0.000216033018440465 |
| syn1.S | 2206.19658454291 | -0.455187974241149 | 0.0987256830349288 | -4.61063383152391 | 4.0144310031402e-06 | 0.000216033018440465 |
| LOC108704221 | 1150.21696426283 | -0.613780102584956 | 0.133142365832609 | -4.60995340398728 | 4.02759232053735e-06 | 0.000216033018440465 |
| rpl11.L | 12711.0431436472 | 0.539099084698213 | 0.117043814975659 | 4.60595961273414 | 4.10568076044286e-06 | 0.000219764654812087 |
| ppp3r1.L | 2222.81805882512 | -0.438151813823848 | 0.0951618914560211 | -4.60427811091101 | 4.13899047412423e-06 | 0.000221088932158189 |
| fgd1.S | 941.292247205091 | -0.487291891907373 | 0.105888851569733 | -4.60191875427479 | 4.18616505788304e-06 | 0.000222906444877373 |
| LOC121398029 | 634.41648583115 | -0.540700509117293 | 0.117499823107332 | -4.60171338831192 | 4.19029557230721e-06 | 0.000222906444877373 |
| cadps.L | 5650.44734821393 | -0.346826154775086 | 0.0754008593232903 | -4.59976395345877 | 4.22969943113991e-06 | 0.000224349033479353 |
| LOC108701467 | 1425.9047235628 | -0.386140802590066 | 0.0839525476998297 | -4.5995126195658 | 4.23480539939708e-06 | 0.000224349033479353 |
| adgrl1.L | 5033.93265557661 | -0.432941341488207 | 0.094153684131858 | -4.59824111483404 | 4.26072772862967e-06 | 0.000225259761861978 |
| mx1.L | 71.6838831047569 | -0.895449978982407 | 0.194783859079689 | -4.59714672054047 | 4.28316013269424e-06 | 0.000225982681847672 |
| spock2.S | 2191.99965189009 | -0.421538684481548 | 0.0917283788003581 | -4.59551002639003 | 4.3169203342134e-06 | 0.000227299070658583 |
| rrm1.S | 4496.06535678511 | 0.344930529989971 | 0.0750841484507666 | 4.59391945047023 | 4.34997344633556e-06 | 0.000228572942801339 |
| nat8.5.L | 155.507578350262 | 0.786967938585131 | 0.171339568651457 | 4.59303093137816 | 4.36854281835853e-06 | 0.000228617453780223 |
| rhbg.L | 2757.07539820659 | 0.642956187138635 | 0.139984967753891 | 4.59303736290472 | 4.36840813186753e-06 | 0.000228617453780223 |
| XB5944457.S | 9676.21561830357 | -0.513755022626853 | 0.11187867350941 | -4.59207288137577 | 4.38865042452067e-06 | 0.000228741779702289 |
| kcnt1.L | 616.751400413898 | -0.492147947399576 | 0.107171561468216 | -4.59215057294405 | 4.38701653136001e-06 | 0.000228741779702289 |
| LOC108719942 | 930.153019227908 | -0.508624575025739 | 0.110874048970098 | -4.58740868354966 | 4.48781620821463e-06 | 0.000233438826959551 |
| hmgcs1.S | 2004.60029845486 | 0.471616392776243 | 0.102835361185173 | 4.58613056190871 | 4.51536295144073e-06 | 0.000233928442062592 |
| magoh.L | 1572.54104824545 | 0.434913569820198 | 0.0948298745288504 | 4.58625060911457 | 4.51276875466093e-06 | 0.000233928442062592 |
| sf3b5.S | 2144.80006746195 | 0.439996105773619 | 0.0959561530477757 | 4.58538709398395 | 4.53146099904285e-06 | 0.000234291971493598 |
| rpl17.L | 11495.8256433249 | 0.445732680314095 | 0.0972293516830143 | 4.58434282033748 | 4.55416516309111e-06 | 0.000234994922415501 |
| inpp5a.1.L | 955.230322012755 | -0.503470874238086 | 0.10985703303385 | -4.58296442507191 | 4.5843005611021e-06 | 0.000236077753445976 |
| rhcg.L | 79.4239895589543 | -0.929033851455351 | 0.202909452649963 | -4.57856368603004 | 4.68179624947184e-06 | 0.000240618213618274 |
| rpl34.S | 6864.34059178489 | 0.485515691655694 | 0.106079165016857 | 4.57691848892799 | 4.71875213505596e-06 | 0.000242035397782195 |
| rpl35a.S | 1919.39496214675 | 0.393105950416373 | 0.085995949878653 | 4.57121470221651 | 4.84905140610967e-06 | 0.000248225250550852 |
| apex1.L | 3072.15665193683 | 0.362365741034075 | 0.0792994913136128 | 4.56958468498865 | 4.88691671315476e-06 | 0.000249174804741883 |
| LOC108707566 | 45.2865455816889 | -0.897816492953579 | 0.196468688100268 | -4.56976886054931 | 4.88262416519181e-06 | 0.000249174804741883 |
| snrpc.S | 2799.9639139442 | 0.379256691138634 | 0.0830255363773742 | 4.56795231547565 | 4.92512037059921e-06 | 0.000250627427142918 |
| rack1.L | 30779.5171939276 | 0.475220848057669 | 0.10404513462572 | 4.56744901880491 | 4.93695700393747e-06 | 0.000250735217916509 |
| MGC80700 | 10165.2513238407 | 0.478521250509822 | 0.104811606484705 | 4.56553683851465 | 4.98217691501615e-06 | 0.000252002750075258 |
| plxna1.L | 7089.86044263482 | -0.466204516526652 | 0.102117201042323 | -4.5653867494217 | 4.98574301415313e-06 | 0.000252002750075258 |
| rpl18.S | 12600.5130198185 | 0.451134288529698 | 0.0988212087628291 | 4.56515655067953 | 4.99121725924251e-06 | 0.000252002750075258 |
| cachd1.S | 962.200132654128 | 0.670820086038772 | 0.147034719636739 | 4.56232437954849 | 5.0590405676191e-06 | 0.000254928216102681 |
| LOC108710030 | 3256.18537272166 | -0.400823358740467 | 0.0879331268548663 | -4.55827482857542 | 5.15755198919559e-06 | 0.000259385655596971 |
| syn2.L | 1059.76955416592 | -0.51040808797354 | 0.111991872815191 | -4.55754578562872 | 5.1754810664505e-06 | 0.000259780956253741 |
| LOC108696619 | 2214.03630006139 | -0.426094961405882 | 0.093563626913182 | -4.55406631255602 | 5.26187570834635e-06 | 0.000263604647136574 |
| nup107.S | 2220.30251105328 | 0.362935960152824 | 0.0797096839799039 | 4.5532229213746 | 5.2830239248843e-06 | 0.000264151196424215 |
| rpl5.L | 26525.1880007605 | 0.477869039589422 | 0.104980797925321 | 4.55196615984345 | 5.31468855505706e-06 | 0.000265060158062174 |
| LOC108702465 | 419.001588466823 | -0.558367116781656 | 0.122672568410236 | -4.55168685238894 | 5.32175046031808e-06 | 0.000265060158062174 |
| LOC108715931 | 4018.07375821771 | -0.324513768980823 | 0.0713099551226931 | -4.55074986967076 | 5.34550646199813e-06 | 0.000265730379035745 |
| rps23.S | 9418.38221928389 | 0.491942872229718 | 0.108225192956368 | 4.54554858061608 | 5.47923570728059e-06 | 0.00027133259356591 |
| synj1.S | 793.492457040999 | -0.530916529517895 | 0.116793932728412 | -4.54575436510447 | 5.47388451306989e-06 | 0.00027133259356591 |
| slc26a1.L | 173.304263126024 | -0.910766698191333 | 0.200450001471087 | -4.54361033428428 | 5.52988395684919e-06 | 0.000271885660050938 |
| LOC108707564 | 6055.99670093347 | -0.426021995215669 | 0.0937649823106384 | -4.54350851157078 | 5.53255703592025e-06 | 0.000271885660050938 |
| LOC108710949 | 4737.39242075477 | -0.408021705246227 | 0.0897875154401576 | -4.54430332820791 | 5.51172409604273e-06 | 0.000271885660050938 |
| psma4.L | 2832.06349009303 | 0.376154490562866 | 0.0827889054771315 | 4.54353742684483 | 5.53179781812269e-06 | 0.000271885660050938 |
| nr2f1.L | 3494.30609344819 | -0.360984411049388 | 0.0795105249335168 | -4.54008335816204 | 5.6231992373829e-06 | 0.000275814715445777 |
| LOC108708507 | 2539.1667029595 | -0.498826657465168 | 0.109969199528097 | -4.53605791081274 | 5.73154481812876e-06 | 0.000280588869780487 |
| ckmt1b.L | 1247.77393099068 | 0.439269366318861 | 0.0968478868147962 | 4.53566289121912 | 5.74228384667043e-06 | 0.000280588869780487 |
| armc6.L | 1460.80543409377 | 0.393455238639227 | 0.0867733852498603 | 4.53428476377047 | 5.77990071599704e-06 | 0.000280869526691181 |
| LOC121395258 | 41.4239946117261 | 0.907064165869938 | 0.200046965663007 | 4.53425605763973 | 5.78068677027198e-06 | 0.000280869526691181 |
| LOC108697176 | 5674.54291808285 | -0.383540234400509 | 0.084580069163247 | -4.53464082253519 | 5.77015933001898e-06 | 0.000280869526691181 |
| dnm1l.S | 2743.5348258462 | -0.346196511124533 | 0.0763692175245646 | -4.53319442658914 | 5.80982917412478e-06 | 0.000281754873481991 |
| rpl27a.L | 8631.05040814136 | 0.510484820468643 | 0.112624356025966 | 4.53263253599379 | 5.82531026026501e-06 | 0.000281975618601946 |
| dynll1.S | 3701.63575595602 | 0.411054454243533 | 0.0907103983322856 | 4.53150313305626 | 5.85654684309855e-06 | 0.000282956757587908 |
| txndc17.S | 549.70157619361 | 0.706164491057607 | 0.155903050314584 | 4.52951042094877 | 5.91205184752971e-06 | 0.000285104556385544 |
| alg10.S | 138.753205892224 | 0.801580192970757 | 0.17701522075091 | 4.52831225230465 | 5.94566763043715e-06 | 0.000285126812017246 |
| atp2b1.S | 3985.52897950506 | -0.40885067729857 | 0.0902800364104397 | -4.52869420034141 | 5.93493189562161e-06 | 0.000285126812017246 |

| | | | | | | |
|---|---|---|---|---|---|---|
| abca3.S | 3324.82783209815 | -0.341440833633845 | 0.0753951538495614 | -4.52868408910118 | 5.93521586153845e-06 | 0.000285126812017246 |
| slc45a4.L | 963.180639222146 | -0.538145881308059 | 0.118880891443314 | -4.52676519139885 | 5.98934257784991e-06 | 0.000286688383132704 |
| iqsec3.S | 1071.83316787346 | -0.492943386298744 | 0.108932203826166 | -4.52523100593269 | 6.03295715012121e-06 | 0.000288241286061347 |
| uba52.L | 9459.73541587935 | 0.516928087971004 | 0.114259238484837 | 4.52416885344117 | 6.06333041860359e-06 | 0.000289156977449118 |
| pcnx2.L | 133.663305484622 | -0.794122125735289 | 0.17561620784878 | -4.52191819572309 | 6.12817442229014e-06 | 0.000290637683998319 |
| galnt14.S | 444.302030444239 | -0.58258230981175 | 0.128834214807678 | -4.5219533544053 | 6.12715637747224e-06 | 0.000290637683998319 |
| glipr2.S | 1322.63997915606 | -0.492012791775304 | 0.10879098757625 | -4.52255101949929 | 6.10987529597244e-06 | 0.000290637683998319 |
| lhx9.S | 791.705819460151 | 0.685261415494874 | 0.151602818396621 | 4.52011000021188 | 6.18075080439305e-06 | 0.000292593340831818 |
| LOC108719602 | 34.191429777878 | -0.899200146497438 | 0.19912828583207 | -4.51568265522938 | 6.31131193131524e-06 | 0.000296159745740688 |
| cabp7.S | 249.987918444485 | -0.655913077357368 | 0.145258141021392 | -4.5154995977869 | 6.31676665704548e-06 | 0.000296159745740688 |
| rps3.S | 10845.9851635206 | 0.558592853739162 | 0.123693513656593 | 4.51594297248252 | 6.30356278729878e-06 | 0.000296159745740688 |
| cct2.L | 8303.2088992462 | 0.35282487002065 | 0.0781411517030437 | 4.51522485055601 | 6.32496201174881e-06 | 0.000296159745740688 |
| nap1l1.L | 9526.24328829362 | 0.362836545501389 | 0.080324786239284 | 4.51711809628121 | 6.26869477729226e-06 | 0.000296159745740688 |
| gjc1.L | 2533.37897690887 | 0.406344378103057 | 0.0899828416324479 | 4.51579846481009 | 6.30786337794145e-06 | 0.000296159745740688 |
| LOC108718675 | 4145.86288035709 | -0.464086322250675 | 0.102858308651013 | -4.51189921686609 | 6.42497171920594e-06 | 0.000300297591223756 |
| map1b.S | 6701.10867207015 | -0.406120885264891 | 0.0900526703073105 | -4.50981502135336 | 6.48841775206017e-06 | 0.000301922798367065 |
| rps14.S | 10493.3673716828 | 0.503133424832631 | 0.111556792957737 | 4.51011015549042 | 6.47939713558674e-06 | 0.000301922798367065 |
| cdk5r1l.L | 1735.32986613623 | -0.437840018898916 | 0.0970905522653256 | -4.50960478319665 | 6.49485089510546e-06 | 0.000301922798367065 |
| lrp1.S | 4735.09592022707 | -0.379178097383866 | 0.0841341646846939 | -4.50682667148233 | 6.58043438203661e-06 | 0.000305351091828318 |
| rps27.L | 16449.9710405333 | 0.455466820510886 | 0.101150657579732 | 4.50285575407023 | 6.70463918740632e-06 | 0.000310555998267654 |
| mab21l1.S | 2251.25778143462 | -0.483244041131511 | 0.107338151954763 | -4.50207155918962 | 6.72943147181455e-06 | 0.000311022587124966 |
| LOC121398856 | 904.66629902569 | -0.485844329236444 | 0.10792283142311 | -4.50177522985565 | 6.73882272104094e-06 | 0.000311022587124966 |
| LOC108717189 | 880.673446397778 | -0.523517767624846 | 0.116379144713128 | -4.49838129430584 | 6.84728109393792e-06 | 0.000315464021827854 |
| serpinf1.L | 916.218204294238 | 0.670480972731363 | 0.149071794654687 | 4.4977051110472 | 6.86908811746692e-06 | 0.000315898815003497 |
| ank1.L | 2030.6014600817 | -0.482948700386562 | 0.107385643444424 | -4.49733022866629 | 6.88120674542502e-06 | 0.000315898815003497 |
| hnrnpab.L | 17976.8343581101 | 0.315314454507353 | 0.0701254657392223 | 4.49643294605589 | 6.91029584270306e-06 | 0.000316109277910885 |
| nop58.S | 2604.937596797 | 0.345360609941617 | 0.0768027249143534 | 4.49672339525383 | 6.90086688977265e-06 | 0.000316109277910885 |
| dnajc5.S | 8562.80331375199 | -0.38319377116936 | 0.0852526435564091 | -4.49480221590797 | 6.96346404278097e-06 | 0.000317977650095131 |
| psmb2.S | 1827.84218259988 | 0.392197965828577 | 0.0872850223257697 | 4.4933020050656 | 7.01272217520868e-06 | 0.000319661187491844 |
| vwa5a2.L | 1180.00026865539 | -0.590697533019399 | 0.131481605324558 | -4.49262489274666 | 7.03506356901412e-06 | 0.000319792862255228 |
| sulf2.L | 3874.17846831598 | -0.303904537713895 | 0.0676476374835102 | -4.49246343285787 | 7.04040099848719e-06 | 0.000319792862255228 |
| sptbn1.L | 13389.3798656258 | -0.396471561021577 | 0.0882812093980299 | -4.49100735847447 | 7.08871017356994e-06 | 0.000320857407856323 |
| r3hdm1.L | 2364.65422250293 | -0.419054232804514 | 0.0933038891149176 | -4.49128366223177 | 7.07951875007708e-06 | 0.000320857407856323 |
| LOC108709287 | 1393.34855351292 | -0.434503597243341 | 0.0967902167462584 | -4.48912722638507 | 7.15155772132577e-06 | 0.000321304768882419 |
| znf706l.S | 3686.3975197157 | 0.327671617255992 | 0.0729967811349404 | 4.48885022272784 | 7.16086209718569e-06 | 0.000321304768882419 |
| LOC108711545 | 1352.12697221143 | -0.429814134085079 | 0.0957392394968288 | -4.48942498753936 | 7.14156900903715e-06 | 0.000321304768882419 |
| cbx3.S | 4028.97618380855 | 0.312798173766358 | 0.0696796139864185 | 4.48909165638212 | 7.15275184764917e-06 | 0.000321304768882419 |
| heg1.L | 299.327460691062 | -0.675928567085131 | 0.150546169837187 | -4.48984233751104 | 7.12759101855755e-06 | 0.000321304768882419 |
| rps17.S | 6588.89235902266 | 0.466948771299389 | 0.104162107309226 | 4.48290442044492 | 7.36339151995621e-06 | 0.000329818578498039 |
| mbp.L | 430.995878287628 | -0.651962756820663 | 0.145539943735189 | -4.47961391277514 | 7.47781746144543e-06 | 0.000334363415087161 |
| rnf103.L | 387.433723953215 | -0.566891506171808 | 0.126615299215802 | -4.47727493978119 | 7.5601859753558e-06 | 0.000337461588519342 |
| rpl27a.S | 5158.40709779115 | 0.533312903440691 | 0.119147483454338 | 4.47607358526443 | 7.60282900411028e-06 | 0.000338778908991442 |
| rps15.S | 10874.8382756898 | 0.468553581496706 | 0.104823686878281 | 4.46992083040146 | 7.82485558378181e-06 | 0.000348071162175122 |
| sez6l2.L | 6865.51631291795 | -0.430999592562554 | 0.0964666388594025 | -4.46786161162642 | 7.9005386818245e-06 | 0.000350832870896854 |
| rps12.L | 26302.2372780177 | 0.507574706953691 | 0.113629418333758 | 4.466930434008 | 7.93499185284959e-06 | 0.000351718823631419 |
| ctnnb1.S | 15580.6742115638 | 0.419785247783396 | 0.0939834594005774 | 4.46658646596719 | 7.94775481306656e-06 | 0.000351718823631419 |
| thoc5.L | 1388.11073760542 | 0.382574943283428 | 0.0856669055669584 | 4.46584291508466 | 7.97541142231576e-06 | 0.000351968071273579 |
| rpl23a.S | 10907.0448046018 | 0.505139410350609 | 0.11312475190791 | 4.46533054730427 | 7.99452261933886e-06 | 0.000351968071273579 |
| prmt1.L | 7111.1997461976 | 0.339135533407577 | 0.0759427900009193 | 4.46567124283256 | 7.98180988626361e-06 | 0.000351968071273579 |
| prmt1.S | 13772.9876289245 | 0.298290871810544 | 0.0668068985389907 | 4.46497110828236 | 8.00795573013918e-06 | 0.000351968071273579 |
| chrm2.L | 495.440894539956 | -0.541252447172969 | 0.12123602199439 | -4.46445238196628 | 8.02737983998139e-06 | 0.000352221768488979 |
| cebpg.L | 975.558569580741 | 0.455530857613765 | 0.102048815664366 | 4.46385246754835 | 8.04990026740258e-06 | 0.000352610232426123 |
| rps10.S | 13687.1852766974 | 0.475103223996727 | 0.106485544817215 | 4.46166871580788 | 8.13238814085702e-06 | 0.000355619684803578 |
| nasp.L | 3213.80192005597 | 0.366413164474567 | 0.0821350682847203 | 4.46110500820916 | 8.15381217720291e-06 | 0.000355953221948959 |
| nkx6-1.L | 719.150435572199 | -0.560503307316392 | 0.125661814686245 | -4.46041073587762 | 8.18027254877814e-06 | 0.000356181725877659 |
| camk4.S | 319.138669210113 | -0.608583484094237 | 0.136472128623098 | -4.45939760912641 | 8.21903254700642e-06 | 0.000356181725877659 |
| LOC108709568 | 132.08304033541 | 0.820579532790305 | 0.184021021075595 | 4.45916193701161 | 8.22807397763895e-06 | 0.000356181725877659 |
| nlgn2.S | 4924.5447307569 | -0.452037374796787 | 0.10136429622567 | -4.45953251419419 | 8.21386126332486e-06 | 0.000356181725877659 |
| h3-3b.S | 3566.9411772813 | 0.439276384639027 | 0.0985079658210648 | 4.45929809815536 | 8.22284907151562e-06 | 0.000356181725877659 |
| LOC108708564 | 17758.3277686368 | 0.469424251383413 | 0.105303291263964 | 4.45783076434624 | 8.27932245982572e-06 | 0.000357799865097996 |
| ankrd13b.S | 590.094353774886 | -0.517377222500795 | 0.116074084667217 | -4.45730176536915 | 8.29977285368164e-06 | 0.000358083845526733 |
| rps17.L | 12363.8335299592 | 0.479530864223989 | 0.107663288510736 | 4.45398678469838 | 8.42902907904892e-06 | 0.000362495195817009 |
| efr3b.L | 984.687103520277 | -0.525053215202838 | 0.117884609197865 | -4.4539589924038 | 8.4301208329537e-06 | 0.000362495195817009 |
| LOC108712711 | 11997.6732776103 | -0.437706375932648 | 0.0983058574728029 | -4.45249537703024 | 8.48780682459355e-06 | 0.000364368412769574 |
| crmp1.S | 6163.40439298714 | -0.499437490975605 | 0.112187395954567 | -4.45181463323976 | 8.51476558227195e-06 | 0.000364918524954512 |
| LOC108717532 | 330.130172269172 | -0.654805578297498 | 0.147199386553396 | -4.44842599979158 | 8.65018401866688e-06 | 0.00037010737592306 |
| rpl37.L | 8900.31116231003 | 0.449682447749315 | 0.101097655694384 | 4.44800074403994 | 8.66732296990561e-06 | 0.000370226709641664 |
| rps4x.S | 18856.4392631898 | 0.467001563421607 | 0.105054603690747 | 4.44532221354464 | 8.77602320807834e-06 | 0.000374250245898217 |
| rps9 | 13122.8506641171 | 0.39655057691406 | 0.0892402259185145 | 4.44363035651827 | 8.84535222101527e-06 | 0.000376584302478868 |
| LOC121393045 | 21151.7449593496 | 0.471740025312717 | 0.106180245960849 | 4.4428228720308 | 8.87862557536428e-06 | 0.000377378154603622 |
| rpl31.L | 9223.76209411266 | 0.518141288142929 | 0.116648855281406 | 4.44188918007771 | 8.91724856342785e-06 | 0.000378396402855984 |
| LOC108706821 | 2626.46458070871 | 0.407039405542849 | 0.0916471204879921 | 4.44137691806892 | 8.93850688218871e-06 | 0.000378675661018832 |
| XB5957215.S | 185.793868878246 | 0.815300011432638 | 0.183701815034652 | 4.43817069133936 | 9.07266596401472e-06 | 0.000383101115992766 |
| sap18.L | 1457.22235325709 | 0.424753585790587 | 0.0956988197775616 | 4.43844121356843 | 9.06127251671145e-06 | 0.000383101115992766 |
| gpc5.L | 1198.14990580758 | -0.520946557852081 | 0.117411075847453 | -4.43694561260068 | 9.12443355602213e-06 | 0.000384657493047992 |
| agpat3.S | 3184.80565279303 | -0.369015622277431 | 0.0831833303741153 | -4.43617273578481 | 9.15723771861511e-06 | 0.000384782952997182 |
| rpl26.L | 10982.2927228798 | 0.482931374840492 | 0.108857258489896 | 4.4363727466581 | 9.1487376199445e-06 | 0.000384782952997182 |
| rps25.S | 11558.9525120003 | 0.509513402448107 | 0.11490744565259 | 4.43411999591884 | 9.24491267169532e-06 | 0.000387835360861365 |
| rps13.L | 11318.7950278014 | 0.392753305509288 | 0.0885873551272961 | 4.43351429721453 | 9.27093564253482e-06 | 0.00038829568113214 |
| vsnl1.L | 4037.12990383392 | -0.366011384072461 | 0.0825826036998903 | -4.43206399016634 | 9.33353070088968e-06 | 0.00038998924571093 |
| LOC108719848 | 385.213824742433 | -0.585111901899614 | 0.132023478474528 | -4.43187763767718 | 9.34160286237808e-06 | 0.00038998924571093 |
| calb2.S | 173.741938780936 | 0.771385340623453 | 0.17421202044409 | 4.4278537075518 | 9.51754160823557e-06 | 0.000396052537891093 |
| sphkap.L | 667.395774043599 | -0.643904993276045 | 0.145419242600514 | -4.42792151685825 | 9.51455072414701e-06 | 0.000396052537891093 |
| hsbp1.L | 2682.71086701611 | 0.411858432343348 | 0.0930321461995872 | 4.42705504675517 | 9.55283601880495e-06 | 0.000396881109960012 |
| LOC108695481 | 144.864407090436 | -0.841874508163731 | 0.190341165347263 | -4.42297653598891 | 9.7350317617936e-06 | 0.000403800352820378 |
| tcp1.S | 6251.27633372727 | 0.340010355955996 | 0.076902697498623 | 4.42130597515235 | 9.81061345656327e-06 | 0.000406282226612091 |
| LOC108700575 | 907.09560702911 | -0.520893303770796 | 0.11783602582636 | -4.4204928002101 | 9.84760668956598e-06 | 0.000407160661207343 |
| rps14.L | 11706.9680847604 | 0.470045448919524 | 0.10637047416766 | 4.41894663531014 | 9.91831318684146e-06 | 0.000408773930064712 |
| tmem145.S | 626.43490239886 | -0.581024610371981 | 0.131482098056613 | -4.41903969407154 | 9.91404390863434e-06 | 0.000408773930064712 |
| eif3l.L | 5433.2712842265 | 0.436584921804517 | 0.0988401718687893 | 4.41707975158203 | 1.00043325299736e-05 | 0.000411661529941467 |
| LOC108703204 | 473.392469453164 | -0.617170869950968 | 0.139760824058736 | -4.41590749129809 | 1.00587097905608e-05 | 0.000413239988210936 |
| pitx2.L | 257.654390005856 | 0.742157657822301 | 0.168092444103054 | 4.41517560044101 | 1.00928027934039e-05 | 0.000413981418235011 |
| LOC108707584 | 34.4710575563103 | 0.859510337449483 | 0.194692290890629 | 4.41471171517687 | 1.01144686498869e-05 | 0.000414211573281081 |
| slc2a1.L | 2821.90343997926 | -0.525270758205049 | 0.119064252522935 | -4.41165796680969 | 1.02582071924914e-05 | 0.000419432243369694 |
| rpl9.L | 13485.3671214254 | 0.464796364834356 | 0.105372081233724 | 4.41100108674327 | 1.02893801700558e-05 | 0.000420041152511772 |
| LOC108715117 | 7591.14024890446 | -0.416063329392501 | 0.0943347683392789 | -4.41049823640963 | 1.03133046071857e-05 | 0.00042035269962937 |
| mars1.S | 1549.70834479483 | 0.464591585854903 | 0.10539516472891 | 4.4080920320196 | 1.04285233494223e-05 | 0.000424378394976492 |
| rps19.S | 17910.465655957 | 0.466570808800726 | 0.105955031177844 | 4.40347951026121 | 1.06528349416726e-05 | 0.000432823844874256 |
| arpc5.L | 1037.89105687214 | 0.436658310553801 | 0.0991771419824655 | 4.40281199705273 | 1.06856759235121e-05 | 0.000433475532746247 |
| lrrn3.S | 1821.41303269778 | 0.435737920440552 | 0.0990132300704634 | 4.40080502505025 | 1.07850002305696e-05 | 0.000436133238164098 |
| LOC108705278 | 231.349089595412 | 0.708606996416586 | 0.161016222392742 | 4.40084226472717 | 1.07831493745005e-05 | 0.000436133238164098 |
| LOC108703551 | 1526.92880576389 | -0.414666197705274 | 0.0942623046345428 | -4.39906704289637 | 1.08717239756859e-05 | 0.000438952235638024 |
| sntb1.L | 327.948103953859 | -0.636138300902045 | 0.144675090904582 | -4.39701331393391 | 1.09750615314561e-05 | 0.000442432167986823 |
| msrb2.S | 681.836863071029 | -0.512853837101395 | 0.11667781298638 | -4.39547008959845 | 1.10533284036841e-05 | 0.000444892157277767 |
| ntn5.L | 433.577895483301 | 0.63017650850318 | 0.143415421456616 | 4.39406377712184 | 1.11251154128606e-05 | 0.000447084077339258 |
| eif3l.S | 3205.83194869385 | 0.427886579697682 | 0.0974469757512123 | 4.39096828197215 | 1.12847002781499e-05 | 0.000452792017381443 |
| psmc3.L | 5134.85449156834 | 0.365076847638926 | 0.083156193876962 | 4.39025441904056 | 1.13218115694189e-05 | 0.000453061192956826 |
| LOC108698084 | 1107.16374184145 | -0.416508823055265 | 0.0948731847209991 | -4.39016382005225 | 1.13265298239207e-05 | 0.000453061192956826 |

| | | | | | | |
|---|---|---|---|---|---|---|
| aqp1.L | 624.996116936664 | -0.700332746206793 | 0.159637020227565 | -4.38703218845139 | 1.14907789919867e-05 | 0.000457668881075699 |
| cdk5r1l.S | 872.720886005937 | -0.510741984256119 | 0.116431838469979 | -4.38661787847498 | 1.15126784425631e-05 | 0.000457668881075699 |
| trim9.L | 1311.65450484088 | -0.437978134306443 | 0.0998266127319828 | -4.38738851614988 | 1.14719761600384e-05 | 0.000457668881075699 |
| sf3b1.L | 4297.99707969861 | 0.336585746730789 | 0.0767253054554878 | 4.38689353835234 | 1.14981032792261e-05 | 0.000457668881075699 |
| nme2.S | 4258.80758774516 | 0.51124178666498 | 0.116644847184631 | 4.38289216372979 | 1.17114090469029e-05 | 0.000464852851400147 |
| rpl21.L | 12814.7672556912 | 0.437260693356568 | 0.0998358878724919 | 4.37979470784121 | 1.18791167198345e-05 | 0.000470063207625352 |
| rps4x.L | 22657.9739788249 | 0.479796626461624 | 0.109540565838052 | 4.38008168746335 | 1.18634827532668e-05 | 0.000470063207625352 |
| npm1.S | 7792.10079737306 | 0.343023360761239 | 0.0783641930676515 | 4.37729717276753 | 1.20160092172595e-05 | 0.000474026051690055 |
| LOC108702162 | 1143.2472735947 | -0.469231523465224 | 0.107189314426405 | -4.37759608759688 | 1.19995464191543e-05 | 0.000474026051690055 |
| gpt.L | 2034.7779286584 | -0.538427107877233 | 0.123031471263464 | -4.37633641496676 | 1.20690692703872e-05 | 0.000475392346833572 |
| lrwd1.L | 495.487689745695 | 0.514374445947984 | 0.117568420604201 | 4.37510722100832 | 1.21372804359772e-05 | 0.00047735035861008 |
| dnajc2.S | 1609.43862973111 | 0.410455086883079 | 0.0938285820347171 | 4.37452083344079 | 1.21699500773669e-05 | 0.000477650611839494 |
| rps15a.S | 8030.93531515766 | 0.483952337221739 | 0.110635227798832 | 4.37430596791204 | 1.21819419608677e-05 | 0.000477650611839494 |
| mn1.L | 1926.49527524869 | -0.335568149733585 | 0.0767321456494767 | -4.37324079619132 | 1.22415570842502e-05 | 0.000479259746242269 |
| cct2.S | 5349.44295571317 | 0.345654072812744 | 0.0790526997667766 | 4.37245121080623 | 1.22859278707253e-05 | 0.000480268089491988 |
| znf385b.L | 742.793835821791 | -0.463566195858695 | 0.106041546499005 | -4.37155257692365 | 1.23366133749891e-05 | 0.000481519856391406 |
| kars1.L | 3378.07504618802 | 0.426440612273615 | 0.0976038245289907 | 4.36909736202961 | 1.24761138313467e-05 | 0.000484965616509934 |
| slc30a10.L | 347.538357183813 | -0.573726487262005 | 0.131317369815058 | -4.36900684250695 | 1.24812856342091e-05 | 0.000484965616509934 |
| LOC108717753 | 2967.01524328818 | -0.522432331674089 | 0.119566648480933 | -4.36938174910371 | 1.245987877157e-05 | 0.000484965616509934 |
| ntn3.S | 339.785804031086 | -0.611209252647914 | 0.140019247677945 | -4.36518023617539 | 1.27017983972551e-05 | 0.000492791576916062 |
| prdm8.L | 444.121989792675 | -0.594107719552697 | 0.136118780555494 | -4.36462710823719 | 1.2733978922078e-05 | 0.000493124699295635 |
| slc66a1.S | 198.390305011211 | -0.757571508288776 | 0.17358071372071 | -4.36437604184358 | 1.27486114120228e-05 | 0.000493124699295635 |
| LOC108706667 | 250.953592607342 | 0.826682885910522 | 0.189453153226698 | 4.36352138684819 | 1.27985422175805e-05 | 0.000493576067583823 |
| llphp3.S | 1713.94449149792 | 0.458309921924004 | 0.105030408525288 | 4.36359268100586 | 1.27943699347808e-05 | 0.000493576067583823 |
| mpst.L | 531.013403248379 | -0.581258814271221 | 0.133364534311044 | -4.35842120451274 | 1.31004071591388e-05 | 0.000504463439859374 |
| gpatch11.L | 1609.07274372655 | 0.379880435943531 | 0.0872771555850602 | 4.35257580746086 | 1.34547325571769e-05 | 0.000516565624963043 |
| tmem63b.S | 2441.16668436964 | -0.371278617228591 | 0.0852970266290908 | -4.35277326656502 | 1.34426155788298e-05 | 0.000516565624963043 |
| ncor2.S | 1315.52593556365 | -0.373186905587064 | 0.0857621061251657 | -4.35141955402093 | 1.35258948642356e-05 | 0.000518526132982585 |
| rps6.L | 21376.8964625762 | 0.453933918888403 | 0.104338030085998 | 4.35060848392729 | 1.35760267576282e-05 | 0.000519675801701494 |
| pnck.S | 219.516334306132 | -0.74333611352625 | 0.170960175537407 | -4.34800743032463 | 1.37379951160306e-05 | 0.000525096702212727 |
| LOC108709624 | 4418.58489746348 | 0.288618276330661 | 0.06648998013307 | 4.34077850155819 | 1.41978789220309e-05 | 0.000541071308993202 |
| gpc1.S | 2494.57511476729 | -0.416010072122898 | 0.095835190032546 | -4.34089056411971 | 1.41906390826931e-05 | 0.000541071308993202 |
| mcm4.L | 2551.78827690968 | 0.344651204620444 | 0.0794165884009978 | 4.33978859530201 | 1.42619853258295e-05 | 0.000541466351652341 |
| LOC121397697 | 53.2394942935721 | 0.882668786908165 | 0.203377260869138 | 4.34005642093938 | 1.42446137299207e-05 | 0.000541466351652341 |
| MGC114789 | 8387.05139842213 | 0.50649360693388 | 0.116713100928621 | 4.33964655984625 | 1.42712061675811e-05 | 0.000541466351652341 |
| cacnb2.S | 330.002710508203 | -0.624205594788667 | 0.143873542101947 | -4.3385711206468 | 1.4341207651206e-05 | 0.000543323285464191 |
| fau.L | 6884.28131330447 | 0.415246493468631 | 0.095764785720426 | 4.33610841756483 | 1.45027431831956e-05 | 0.000548637498719129 |
| lamp5.S | 1592.59365950041 | -0.592745389143878 | 0.136715936661507 | -4.33559834806565 | 1.45364162674017e-05 | 0.00054910620746554 |
| ddx39a.S | 6377.99558609946 | 0.350988327155239 | 0.0809717952308625 | 4.33482209644101 | 1.45878049139461e-05 | 0.000550241764297966 |
| LOC108718251 | 1116.58450501714 | -0.406097368415512 | 0.0937307416617294 | -4.33259527467628 | 1.47361858330133e-05 | 0.000555027145243422 |
| zeb2.S | 1294.9581909395 | 0.388897817868798 | 0.0898017101247614 | 4.33062819548206 | 1.48684553660195e-05 | 0.000559192636214726 |
| psmc2.S | 3076.63487211626 | 0.333999749945945 | 0.0771367340250449 | 4.32997007414783 | 1.49129606650332e-05 | 0.000560050051175921 |
| bcar3.L | 343.992256220572 | 0.556831268569724 | 0.128625918880472 | 4.32907541609999 | 1.49736838038037e-05 | 0.000561513142642637 |
| LOC108695679 | 3462.20509333662 | 0.436054747779694 | 0.100835860250753 | 4.32440152437178 | 1.52946504860298e-05 | 0.000572716955790374 |
| rpl14.S | 7411.58073648455 | 0.471022736396875 | 0.108993317314706 | 4.32157445980702 | 1.54919753493432e-05 | 0.000579265165236313 |
| gabarapl1.L | 3141.01382413227 | -0.415219789815625 | 0.0961748353375774 | -4.31734339194018 | 1.57918378631345e-05 | 0.000589622889824704 |
| LOC108708975 | 22.5642616812518 | 0.82607230595918 | 0.191395387015952 | 4.3160512844039 | 1.58845090044684e-05 | 0.000592225913750412 |
| itga10.L | 153.014622758939 | -0.820036714084214 | 0.190020044866675 | -4.31552742059177 | 1.59222284972346e-05 | 0.000592775606390552 |
| LOC108711739 | 1476.56129944746 | -0.63300981306791 | 0.14669395505655 | -4.31517312914419 | 1.59477867404092e-05 | 0.000592871610810601 |
| efna2.S | 584.155176281653 | 0.658387893991056 | 0.152639822688566 | 4.31334289043546 | 1.60804424306571e-05 | 0.000596085365964012 |
| LOC108695817 | 145.644590271386 | 0.752249685205319 | 0.17439944742139 | 4.31337195345412 | 1.60783277488996e-05 | 0.000596085365964012 |
| pfdn6.L | 1244.15424912653 | 0.420897176559245 | 0.0976226010376814 | 4.31147267216106 | 1.62170819468572e-05 | 0.000600287970245216 |
| LOC108702731 | 3624.88808323205 | -0.498901319044907 | 0.115729583909107 | -4.31092294807471 | 1.62574552337491e-05 | 0.000600920265087001 |
| LOC121394345 | 699.308884533797 | 0.459527917668255 | 0.106637365762737 | 4.30925796395498 | 1.63803215557973e-05 | 0.0006045955595702 |
| rpl22.L | 8417.65572194602 | 0.46443458773563 | 0.107857999526523 | 4.30598184441037 | 1.66246692270209e-05 | 0.000612737808653056 |
| LOC108698377 | 598.705407204009 | -0.560450978110085 | 0.130182851275061 | -4.30510603063932 | 1.66905774798368e-05 | 0.000614289442196561 |
| stt3a.S | 2875.59401674646 | 0.374378249807161 | 0.0869845416355045 | 4.30396301191005 | 1.67769685406694e-05 | 0.000616589442092979 |
| pola1.S | 1141.05921059752 | 0.437183200256693 | 0.101594478860186 | 4.30321809966013 | 1.68339930105490e-05 | 0.000617787029825344 |
| rps28p9.S | 5064.73682808595 | 0.455053295685038 | 0.105822994355569 | 4.30013626486551 | 1.70693120313739e-05 | 0.000624329610516741 |
| LOC108708589 | 3620.22250299028 | -0.376530511670224 | 0.0875664393380312 | -4.29994087365719 | 1.70843684118147e-05 | 0.000624329610516741 |
| bhlhe22.S | 437.021431951319 | -0.597715325222484 | 0.139003237846345 | -4.30001008957217 | 1.70790333515543e-05 | 0.000624329610516741 |
| LOC108708009 | 621.077959454207 | -0.471903787067111 | 0.109768289503215 | -4.29909028557187 | 1.71500602913789e-05 | 0.00062584378432472 |
| pax5.L | 457.769323188678 | -0.652943014573878 | 0.152021944649071 | -4.29505763842936 | 1.74647955200281e-05 | 0.000635531346144887 |
| acss1.S | 226.622440876118 | -0.654878770409916 | 0.152470153524905 | -4.2951276382295 | 1.74592856230214e-05 | 0.000635531346144887 |
| rps27.S | 8769.63427344492 | 0.463182115736414 | 0.107873621215719 | 4.29374772549972 | 1.75682088555478e-05 | 0.00063839406827202 |
| LOC108701569 | 9745.93444258668 | 0.361338266216195 | 0.0841692889819885 | 4.29299416196231 | 1.76279643202529e-05 | 0.000639664528076688 |
| plin3.S | 498.616673154358 | 0.581688701932618 | 0.135541518758106 | 4.29159055662292 | 1.7739782924744e-05 | 0.000642817976767409 |
| fam219a.S | 2605.32091700404 | -0.425255012489125 | 0.0991061670618668 | -4.29090363492376 | 1.77947527147084e-05 | 0.000643905497951581 |
| ptn.L | 11928.4813681014 | -0.272283856108383 | 0.0634703860978121 | -4.2899353989871 | 1.78725098706274e-05 | 0.000645469506953553 |
| kcnj10.L | 662.649603324674 | -0.560428659936555 | 0.130643882242213 | -4.28974285146797 | 1.78880115299144e-05 | 0.000645469506953553 |
| ppid.S | 3414.01719891076 | 0.378633172173348 | 0.0883061449259292 | 4.28773300534118 | 1.80505872379441e-05 | 0.000649519038687528 |
| ercc5.L | 1444.40096045274 | 0.356418706850505 | 0.0831231666961724 | 4.28783840915576 | 1.80420263011252e-05 | 0.000649519038687528 |
| LOC108712252 | 169.667131045802 | -0.71597795430817 | 0.16706740738429 | -4.2855633275093 | 1.8227671389839e-05 | 0.000654977607044353 |
| LOC108701018 | 1220.31778088657 | -0.477161872048459 | 0.111375452960617 | -4.28426425540273 | 1.83344895094459e-05 | 0.000657899623565651 |
| gad1.1.S | 2543.22440724616 | -0.468631936969607 | 0.109417488021629 | -4.28297108115818 | 1.84414147710601e-05 | 0.000660817362629655 |
| mcm7.L | 5115.04514966683 | 0.317123796183298 | 0.0740815196834503 | 4.28074096668596 | 1.86272072246178e-05 | 0.000666549162822662 |
| madd.S | 3426.71372722513 | -0.39017446821829 | 0.0911543395701263 | -4.28037184031292 | 1.86581308826381e-05 | 0.000666730992759089 |
| LOC108713446 | 2906.34814750594 | -0.392716558640729 | 0.0917570527107943 | -4.27996047212324 | 1.86926509748731e-05 | 0.000667040657194641 |
| sec61b.L | 3860.31030262622 | 0.395432495111718 | 0.0924870201991426 | 4.27554584697696 | 1.90669556599391e-05 | 0.000679344166842072 |
| LOC121398252 | 26.4027931652317 | 0.730810124418537 | 0.170938715788262 | 4.27527562172502 | 1.90900977116474e-05 | 0.000679344166842072 |
| rps25.L | 9862.83828312739 | 0.433897553554618 | 0.101523259622853 | 4.27387334849665 | 1.92106184403308e-05 | 0.000682691399119193 |
| h2az2.S | 4035.58088856652 | 0.313878322459717 | 0.0734582050448845 | 4.27288309410679 | 1.92961637172217e-05 | 0.000684132678061853 |
| LOC108714021 | 691.085115591502 | 0.580545348587033 | 0.135870315212737 | 4.27279017994513 | 1.930420890035e-05 | 0.000684132678061853 |
| hes5.3.S | 895.259874633642 | 0.592043265175071 | 0.138697368126095 | 4.26859768987702 | 1.96705677561765e-05 | 0.000696160011123943 |
| ankrd10.L | 1566.87250390519 | 0.354206761756094 | 0.0830592565455833 | 4.26450676887173 | 2.00344270974757e-05 | 0.000707097426969731 |
| LOC108711237 | 2475.01837881909 | -0.35118763581873 | 0.0823456734956887 | -4.26479766222468 | 2.00083438971234e-05 | 0.000707097426969731 |
| skor1.S | 605.740293120694 | -0.65925952246577 | 0.154623871560939 | -4.26363352443902 | 2.01129219361124e-05 | 0.000708898068240026 |
| LOC108708797 | 99.862557332489 | -0.827648376892516 | 0.194184647940476 | -4.26217203919343 | 2.02449485972007e-05 | 0.00071257799973776 |
| rab3a.S | 2691.17259035263 | -0.440921719479805 | 0.103513602296232 | -4.25955342775136 | 2.04835734543271e-05 | 0.000719015231458013 |
| megf6.L | 320.454979443528 | -0.588962224008141 | 0.138267808945833 | -4.25957588030392 | 2.04815160961664e-05 | 0.000719015231458013 |
| erh.L | 2721.50971599296 | 0.383853770403176 | 0.0901637497750177 | 4.25729599047281 | 2.06914329945573e-05 | 0.000725324689211384 |
| znf423.L | 2439.87517753623 | 0.324532247784839 | 0.076250910127487 | 4.25610982534163 | 2.08014556067922e-05 | 0.000728192068731668 |
| lingo3.S | 384.450254134802 | -0.538878364811626 | 0.126632849790475 | -4.25543897735261 | 2.08639263873904e-05 | 0.000729389296469744 |
| chrna2.L | 542.209762422688 | -0.491784823201451 | 0.115616462323109 | -4.25358822887246 | 2.10371994570734e-05 | 0.000733459116206072 |
| LOC121396631 | 107.176094950061 | -0.827408070230856 | 0.194513905359781 | -4.25372195730917 | 2.10246335888713e-05 | 0.000733459116206072 |
| pafah1b1.L | 1776.95461584632 | -0.413202347430796 | 0.0971600076109287 | -4.25280274869306 | 2.11111520060175e-05 | 0.000735044158913971 |
| rps3a.S | 26445.9968860301 | 0.454921796842821 | 0.106979510704397 | 4.25241986850968 | 2.1147289634503e-05 | 0.000735121036049262 |
| LOC100126651 | 187.050928692624 | 0.797679373869235 | 0.187593220720411 | 4.2521759091609 | 2.11703461156822e-05 | 0.000735121036049262 |
| frs2.S | 704.591051156357 | 0.429901925509069 | 0.101241686215468 | 4.2462936126343 | 2.17335789034217e-05 | 0.000753664429715429 |
| rpl31.S | 8308.2807790859 | 0.493195643531299 | 0.116162306609433 | 4.24574595604016 | 2.17867370076796e-05 | 0.000754493711138433 |
| LOC108712655 | 622.48933021863 | -0.57857004494845 | 0.136305027479505 | -4.24467134959819 | 2.18914032104819e-05 | 0.000757102148566264 |
| LOC121394610 | 8058.39200605009 | 0.497448547832945 | 0.117202588865117 | 4.24434777976992 | 2.19230124106047e-05 | 0.000757180348318072 |
| rps10.L | 13799.3434689255 | 0.412529708677523 | 0.0972270728603945 | 4.2429510273323 | 2.20599589070083e-05 | 0.000760891630749753 |
| LOC108710294 | 43.4790758157195 | 0.848255889521368 | 0.200002017971231 | 4.24123665413908 | 2.22291601402222e-05 | 0.000763825859512832 |
| LOC100036976 | 24.0419072726665 | 0.730860761730147 | 0.172332985331705 | 4.24098010211681 | 2.22545867261064e-05 | 0.000763825859512832 |
| LOC108717680 | 515.726298042194 | -0.493132955764784 | 0.116265689866857 | -4.24143146898713 | 2.22098707304908e-05 | 0.000763825859512832 |
| ptmap12.L | 9932.47759839329 | 0.324787995296708 | 0.0765902400713442 | 4.24059246966933 | 2.22930570625257e-05 | 0.000763825859512832 |
| nkap.S | 1161.15405651247 | 0.398451167135804 | 0.0939570472649521 | 4.24078000250689 | 2.22744375882236e-05 | 0.000763825859512832 |

| | | | | | | |
|---|---|---|---|---|---|---|
| banf1.L | 1204.78940178536 | 0.375233539733198 | 0.0885075567173256 | 4.23956500043961 | 2.23953340639439e-05 | 0.000766312491843172 |
| sall2.L | 1841.22707351833 | 0.417484116951514 | 0.0985022287362389 | 4.23832153147944 | 2.25197095762861e-05 | 0.000768529771254208 |
| rps7.L | 13319.539657939 | 0.513593001776592 | 0.121175885328684 | 4.23840932033213 | 2.25109071471635e-05 | 0.000768529771254208 |
| pcdh7.L | 1074.4648460635 | -0.447554756207723 | 0.1056208583571 | -4.23737094328997 | 2.26152333483673e-05 | 0.000770770172242901 |
| snrpe.L | 1295.39378213209 | 0.36949019887401 | 0.0872080800191656 | 4.23688033027223 | 2.2664685369586e-05 | 0.000771436520495143 |
| iqsec3.L | 2088.69418443901 | -0.433400033142196 | 0.102308193830493 | -4.23622015906436 | 2.27313906724899e-05 | 0.000772687588076731 |
| LOC108709952 | 7015.69852155443 | 0.32014312439305 | 0.075578988207049 | 4.23587470522913 | 2.27663705897371e-05 | 0.000772858370020022 |
| phr.L | 1394.36626489106 | 0.456084511675959 | 0.107690945942768 | 4.2351240179313 | 2.28425602748163e-05 | 0.000774425827976688 |
| tmem163.S | 1428.76736879495 | -0.500353729167223 | 0.118162662142676 | -4.234448683655 | 2.29113094213311e-05 | 0.000775737248123809 |
| LOC108714333 | 412.605587300566 | 0.564367587925745 | 0.133320312535403 | 4.23317030385656 | 2.30419882256218e-05 | 0.000779139313526923 |
| slc25a11.S | 1679.54282121665 | -0.368943883571073 | 0.0872010519737118 | -4.23095679719894 | 2.32699367175065e-05 | 0.000785817234701137 |
| map7d1.L | 1887.55520385894 | -0.35173035875261 | 0.0831508715861592 | -4.2300261205098 | 2.33664178517067e-05 | 0.000788043896175207 |
| matr3.S | 2306.23649680964 | 0.362815207219159 | 0.0858139277299152 | 4.2279291580856 | 2.35852021258825e-05 | 0.00079438409249056 |
| cdk14.L | 733.210381491139 | -0.450500243777516 | 0.106563946237116 | -4.22751089543087 | 2.36290736445839e-05 | 0.000794824119987307 |
| inpp5f.L | 1030.64708923859 | -0.377614635179602 | 0.0893471377199905 | -4.22637641021056 | 2.37484609392484e-05 | 0.000797799859677876 |
| rpl11.S | 14315.6398854853 | 0.432664673259048 | 0.102430247197362 | 4.22399325489659 | 2.400112357174e-05 | 0.000805239256373073 |
| pum3.S | 1429.48476981277 | 0.372353047614604 | 0.088165174362351 | 4.22335746861075 | 2.40689607168249e-05 | 0.000806466475966342 |
| rpl7a.S | 20318.4879310458 | 0.386685313873984 | 0.0915780825096912 | 4.2224657175265 | 2.41644162351956e-05 | 0.000808614706703043 |
| med8.L | 335.699428741846 | 0.603214914133766 | 0.142895211404202 | 4.22137948645093 | 2.4281176028942e-05 | 0.000809372534298068 |
| sim1.S | 30.0936067875599 | 0.825462072225444 | 0.195535636457209 | 4.22154287157823 | 2.42635794085543e-05 | 0.000809372534298068 |
| rps7.S | 13250.290504515 | 0.446186360740892 | 0.10568958814361 | 4.22166808082002 | 2.42501025529957e-05 | 0.000809372534298068 |
| adam11.S | 2312.16274209443 | -0.45042560423483 | 0.106720578920669 | -4.22060682944434 | 2.43645559860334e-05 | 0.00081110392830924 |
| rspo3.L | 104.985040851067 | 0.769469585809738 | 0.182354921769118 | 4.21962609149632 | 2.44707829980092e-05 | 0.000813590465655459 |
| ebna1bp2.L | 2061.89448235836 | 0.370135958165792 | 0.0877914612689951 | 4.21608152792544 | 2.4858392213748e-05 | 0.000825413795514411 |
| rps21.S | 5034.20396903167 | 0.453656637879717 | 0.107642798473521 | 4.21446343195278 | 2.50372713080991e-05 | 0.000830284832582206 |
| hes5.2.L | 1142.33713699792 | 0.519691308998792 | 0.123332658460978 | 4.21373637351066 | 2.51180450682935e-05 | 0.00083189417556094 |
| LOC108701416 | 86.3731882594896 | 0.842641484995986 | 0.20010460439677 | 4.21100497680297 | 2.54237144759324e-05 | 0.000840938248050072 |
| foxq1.S | 158.129947968669 | -0.709249812602518 | 0.168459073294756 | -4.21022031482704 | 2.5512177791268e-05 | 0.000842783850210903 |
| psma5.S | 3227.57324040323 | 0.322433679258585 | 0.076591099896618 | 4.20980609618876 | 2.55589950727322e-05 | 0.000843250732578633 |
| dach2.L | 1718.8174604465 | -0.33282257760075 | 0.079074199784641 | -4.20899077710801 | 2.56513857739055e-05 | 0.000845218075308763 |
| hnrnpdl.S | 10849.9265717971 | 0.329614676251484 | 0.0783312308972536 | 4.20795987086986 | 2.57686612808951e-05 | 0.000847999312560067 |
| brd7.L | 3753.15652799329 | 0.356476621852291 | 0.0847321315152874 | 4.20710084211649 | 2.58667734085153e-05 | 0.000849440217509718 |
| ca7.L | 304.918166761584 | -0.751227447307417 | 0.178566060716378 | -4.20700016729728 | 2.58782949985519e-05 | 0.000849440217509718 |
| gabrd.L | 54.6751592770742 | -0.828470532821259 | 0.197076906331138 | -4.20379306862683 | 2.62478918002363e-05 | 0.000860477266132269 |
| LOC121397082 | 308.807429966667 | -0.607069391073474 | 0.144495957251769 | -4.20128979813406 | 2.6539859850672e-05 | 0.000868944649933171 |
| rpl39.S | 6057.0254834139 | 0.451709745333701 | 0.107579801563908 | 4.19883415629245 | 2.68292718423957e-05 | 0.00087727072925223 |
| abca3.L | 1956.86304859638 | -0.360728998395678 | 0.0859173826849987 | -4.19855664968554 | 2.6862165740669e-05 | 0.00087727072925223 |
| nsa2.L | 5430.24699853766 | 0.438264303570015 | 0.104397761939925 | 4.1980239367795 | 2.692541769787e-05 | 0.000878224749184636 |
| ado.L | 2028.274013049 | -0.417797308180467 | 0.0995780139092501 | -4.1956782604765 | 2.72056215003866e-05 | 0.000886243730694412 |
| hnrnpk.S | 13727.4043718203 | 0.296027669447921 | 0.0705882286676129 | 4.19372571086691 | 2.74409766099031e-05 | 0.000892195953322045 |
| kalrn.L | 3371.91462936861 | -0.290890845205794 | 0.0693656033715201 | -4.19358920080016 | 2.74575033696784e-05 | 0.000892195953322045 |
| pitpnc1.S | 912.704313781087 | -0.493214502460619 | 0.11762515571543 | -4.19310392798842 | 2.75163301745235e-05 | 0.000892982790569441 |
| eif1ax.S | 2831.1732521102 | 0.370715781551398 | 0.0884178608969339 | 4.19277030444709 | 2.75568429111181e-05 | 0.000893174054154332 |
| rpl24.S | 8885.77523127592 | 0.466120097452786 | 0.111231273720611 | 4.19054895139993 | 2.78280367627574e-05 | 0.000899236738024286 |
| sult4a1.L | 1752.26374202458 | -0.438852405194866 | 0.104728482792451 | -4.19038253485039 | 2.78484555690467e-05 | 0.000899236738024286 |
| LOC108697808 | 930.406262997452 | -0.394843510479803 | 0.0942202545890725 | -4.19064363816309 | 2.78164253381495e-05 | 0.000899236738024286 |
| hap1.S | 516.084181814056 | 0.542313665097508 | 0.129458584949738 | 4.18909001135815 | 2.80075299110945e-05 | 0.000903242840600298 |
| LOC108701181 | 4847.02714753897 | -0.324129767406631 | 0.0773866239618433 | -4.18844692806923 | 2.8086997415631e-05 | 0.000904674823125194 |
| mafa.L | 408.123649641363 | -0.582284839834288 | 0.139046728199894 | -4.18769177364025 | 2.81805874722914e-05 | 0.000906557552101146 |
| atp1a1.S | 5690.05784036708 | -0.446620433155833 | 0.106752394135951 | -4.18370413863558 | 2.86797315491121e-05 | 0.000921465845538098 |
| parp12.L | 86.572871613614 | -0.81099614817697 | 0.193867724461055 | -4.18324478936094 | 2.87377665123604e-05 | 0.000922182059724999 |
| rps16.L | 9564.47881851012 | 0.493084292786957 | 0.117926553480558 | 4.1812829955066 | 2.89868817281274e-05 | 0.000929020557249299 |
| naca.L | 10300.9531865864 | 0.443808186182714 | 0.106166935049956 | 4.1802863195955 | 2.91142280202062e-05 | 0.000931944271614542 |
| irf7.L | 281.173913745068 | -0.688253257535228 | 0.164693350982972 | -4.17899844424434 | 2.92795688484407e-05 | 0.000936075435303309 |
| stard15.S | 331.782321326208 | -0.538600193196104 | 0.12890049315001 | -4.17841840658671 | 2.93543267664468e-05 | 0.000936145402440456 |
| eif5a.L | 11521.6278680766 | 0.378513103198378 | 0.0905819193219961 | 4.17868274410104 | 2.93202352602239e-05 | 0.000936145402440456 |
| c11orf87.S | 954.181988107481 | -0.410005226631907 | 0.0981397586512992 | -4.1777688499184 | 2.94382599554916e-05 | 0.000936719376828578 |
| aox1.S | 70.8074059587326 | 0.829512749305783 | 0.19855646077397 | 4.17771724008554 | 2.94449385506968e-05 | 0.000936719376828578 |
| dhx32.L | 408.90258669258 | 0.502752911283471 | 0.120386242229954 | 4.17616583066978 | 2.96463730731525e-05 | 0.000941966040994255 |
| LOC108706710 | 6137.17210722895 | -0.300751250727395 | 0.0720320760208433 | -4.17524063363618 | 2.97671230976229e-05 | 0.000944639330773273 |
| uba2.L | 3496.08457009509 | 0.357876093067441 | 0.0857543053788003 | 4.17327260114351 | 3.00255329787929e-05 | 0.00095166922709196 |
| LOC108699406 | 1008.94816773013 | -0.526036339134023 | 0.126080683052249 | -4.17221993408799 | 3.01646257671664e-05 | 0.000954904717537294 |
| rpl18.L | 13614.2483516898 | 0.394243429925696 | 0.0945002513173532 | 4.1718770524931 | 3.02100640450362e-05 | 0.000955171142600409 |
| rpl30.L | 8949.86141692673 | 0.498439098299062 | 0.119546403691373 | 4.16941942967901 | 3.05376545941553e-05 | 0.000964346987183851 |
| ppil4.S | 855.784680045197 | 0.426159724951275 | 0.102234390051923 | 4.16845764653984 | 3.06667729264622e-05 | 0.000967240515284504 |
| stk32b.L | 1104.44704546057 | -0.484340122657322 | 0.116217655431782 | -4.16752618918235 | 3.07923144558376e-05 | 0.000969096642778911 |
| MGC68875 | 71.487842764839 | -0.82377843547649 | 0.197669009933571 | -4.16746376052235 | 3.08007460108026e-05 | 0.000969096642778911 |
| rps16.S | 6575.9843001206 | 0.449422543249953 | 0.107849326999823 | 4.16713349774255 | 3.08453875013778e-05 | 0.000969319120018936 |
| rpl34.L | 9368.67536167373 | 0.47378622160671 | 0.113739431181196 | 4.16554062813915 | 3.10615598165901e-05 | 0.000972104200434227 |
| hmgn1.S | 26573.5615325526 | 0.317635078961404 | 0.0762451504924971 | 4.16597090975197 | 3.10030237318582e-05 | 0.000972104200434227 |
| crnkl1.S | 1404.30771645106 | 0.376591764810019 | 0.0903990786093958 | 4.16588056651804 | 3.10153054451384e-05 | 0.000972104200434227 |
| rps20.L | 18918.4743027305 | 0.456712404735182 | 0.109645084390883 | 4.16537054326127 | 3.10847273394666e-05 | 0.000972104200434227 |
| maf.L | 1170.76594570442 | -0.451858260838258 | 0.108515161213109 | -4.16401040911573 | 3.12705848699888e-05 | 0.000976732554050496 |
| fau.S | 6093.25408400833 | 0.480125699966912 | 0.115314195003026 | 4.16363050493753 | 3.13226857939876e-05 | 0.000977176896595984 |
| rpl13a.L | 21802.1737303143 | 0.436094161716194 | 0.104763137835708 | 4.16266800255721 | 3.14550550275352e-05 | 0.000979949801769827 |
| LOC108702543 | 59.7714434208596 | 0.840598268811341 | 0.201948808654362 | 4.16243242241669 | 3.14875343281855e-05 | 0.000979949801769827 |
| LOC108719919 | 899.300871425902 | -0.608933091343234 | 0.146316673419521 | -4.16174778384479 | 3.15821060302308e-05 | 0.000981708838048139 |

### KDBarhl1_A et B_vs_KDCT COMMON GENES pAdj<0.00[illegible]

**MOBarhl1_1_MOBarhl1_2**

LOC100126660
LOC108695757
LOC108695983
LOC108696020
LOC108696074
LOC108696239
LOC108696915
LOC108697444
LOC108697760
LOC108697808
LOC108697935
LOC108698084
LOC108698099
LOC108698154
LOC108698377
LOC108698757
LOC108699226
LOC108699273
LOC108699406
LOC108699444
LOC108699962
LOC108700185
LOC108700350
LOC108700443
LOC108700932
LOC108701018
LOC108701181
LOC108701569
LOC108701642
LOC108702162
LOC108702465
LOC108702731
LOC108702869
LOC108703011
LOC108703204
LOC108703406
LOC108703510
LOC108703551
LOC108704224
LOC108704587
LOC108704633
LOC108705545
LOC108706170
LOC108706667
LOC108706905

LOC108707429
LOC108707553
LOC108707564
LOC108707869
LOC108708151
LOC108708306
LOC108708507
LOC108708564
LOC108708999
LOC108709287
LOC108709337
LOC108709624
LOC108709778
LOC108709952
LOC108710030
LOC108710372
LOC108711439
LOC108711545
LOC108711739
LOC108712711
LOC108712865
LOC108713037
LOC108713150
LOC108713446
LOC108713567
LOC108713772
LOC108713861
LOC108714021
LOC108714603
LOC108715150
LOC108715931
LOC108717002
LOC108717189
LOC108717266
LOC108717850
LOC108718251
LOC108718312
LOC108718675
LOC108719610
LOC108719750
LOC108719848
LOC121393045
LOC121393901
LOC121394610
LOC121396584
LOC121398252
LOC121398856
LOC121399515

LOC121402164
LOC443585
MGC114789
MGC115323
MGC68655
MGC68699
MGC80700
XB5796436.L
XB5848002.S
XB5957215.S
abca3.L
abca3.S
acss1.S
actl6a.S
adgrl1.L
ado.L
agpat3.S
ahcy.L
aldoa.L
ank1.L
ank1.S
ankrd10.L
apcdd1.L
apcdd1.S
apex1.L
apex1.S
aplp1.S
app.L
app.S
armc6.L
arpp19.S
atp13a5l.2.L
atp1a1.L
atp1a1.S
atp1a2.S
atp1b1.L
atp1b2.S
atp2b1.L
atp2b1.S
atp2b4.L
atp2b4.S
atp6v1a.L
atp6v1a.S
barhl2.L
barhl2.S
bhlhe22.L
bsx.L
bsx.S

bzw1.S
c11orf87.S
ca7.L
cachd1.S
cacna1a.S
cadps.L
calb1.L
camk2g.L
camk2g.S
camk4.S
cbx3.S
cct5b
cdk5r1l.L
cdk5r1l.S
chrm2.L
cirbp.S
ckmt1b.L
clstn3.L
clstn3.S
cntnap1.L
col5a3.S
cox4i2.L
cpne4.L
crmp1.L
crmp1.S
crx.L
crx.S
csnk2b.L
ctnna2.L
ctnnb1.L
ctnnb1.S
cyp27c1.S
dach1.S
dach2.L
dbi.S
dctn1.L
ddx39a.S
dhcr24.L
dhrs3.L
dhx32.L
dlx5.S
dmrta2.L
dmrta2.S
dnajc5.S
dner.L
dner.S
dnm1l.L
dnm1l.S

dpysl2.L
dusp8.L
dynll1.S
ednrb2.L
eef1a2.L
efna2.S
efnb3.L
efr3b.L
eif3l.L
eif5a.L
emilin3.L
emx2.L
epha10.L
epha4.L
erh.L
fam219a.S
fau.L
fau.S
fgd1.S
fnbp1.L
foxq1.S
frmpd1.L
fry.L
fst.L
galnt14.S
gipr.L
glipr2.S
glra1.S
gnat2.S
gpc1.L
gpc1.S
gpc5.L
gpt.S
grid2ip.L
grin1.L
gucy2d.L
gucy2d.S
h2az1.L
h2az1.S
h2az2.S
h3-3b.S
hectd1.L
helt.L
hes5.1.L
hes5.1.S
hes5.2.L
hes5.3.L
hes5.3.S

hes5.4.L
hmga2.L
hmgb2.L
hmgb2.S
hmgb3.L
hmgn1.L
hmgn1.S
hmgn2.S
hnrnpab.L
hnrnpdl.L
hnrnpk.S
hsbp1.L
hsf2.2.L
hspa12a.S
igfbpl1.L
igfbpl1.S
iqsec3.S
itga10.L
kalrn.L
kars1.L
kcnc1.S
kcne3.S
kcnh4.L
kcnh4.S
kcnj10.L
kctd15.L
khdrbs1.S
kiaa1549l.L
kif1a.L
ldha.L
ldha.S
lef1.L
lef1.S
lhx2.L
lhx2.S
lhx9.L
lhx9.S
lman1l.L
lmo3.L
lmo3.S
lmx1a.L
lrit1.S
lrp1.L
lrp1.S
lrrn3.S
lrwd1.L
ltbp4.L
mab21l1.S

madd.S
maf.L
mafa.L
magoh.L
map1b.S
map4.L
map7d1.L
map7d2.S
mapk8ip2.L
mapk8ip2.S
mapt.L
mapt.S
mast1.L
matr3.S
mcm4.L
mcm7.L
megf6.L
meis2.L
mmp16.L
mn1.L
msrb2.S
mtss2.L
mvd.L
myg1.S
naca.L
nap1l1.L
nasp.L
ncor2.S
nefh.L
nefh.S
nefl.L
nefl.S
nefm.S
nfia.S
nfib.L
nif.L
nif.S
nkx2-2.L
nkx2-4.S
nkx6-1.L
nlgn2.S
nme2.S
nmu.S
not.L
not.S
npm1.L
npm1.S
npy.L

nr2e1.L
nr2e1.S
nr2f1.L
nsa2.L
ntn1.L
ntn3.S
ntn5.L
odc1.L
odc1.S
ogdh.S
otp.L
otp.S
otx1.L
otx2.L
otx2.S
pa2g4.L
pafah1b1.L
paqr6.L
paqr6.S
pc.1.L
pcdh10.L
pcdh7.L
pcdh7.S
pcna.S
pfkp.L
pfkp.S
phr.L
phyhipl.S
pitpnc1.S
pitx2.S
pitx3.L
pitx3.S
plcb4.S
plxna1.L
pmp2.S
pomp.S
pou4f4.L
pou4f4.S
prdm12.L
proca1.L
prpf19.L
prrt1.S
prrt2.L
psma3.L
psma4.L
psma5.S
psma6.S
psmb1.S

psmb7.S
psmc2.S
psmc3.L
ptchd4.L
ptgds.S
ptmap12.L
ptmap12.S
ptn.S
pygm.L
pygm.S
r3hdm1.L
rab11fip5.L
rab6b.L
rack1.L
rbp1.S
rdh5.L
rgma.L
rgr.L
rgr.S
rlbp1.S
rngtt.S
rora.L
rpe65.L
rpl10a.S
rpl11.L
rpl11.S
rpl12.L
rpl13a.L
rpl14.S
rpl15.L
rpl15.S
rpl17.L
rpl18.L
rpl18.S
rpl18a.L
rpl18a.S
rpl21.L
rpl22.L
rpl23a.L
rpl23a.S
rpl24.S
rpl26.L
rpl26.S
rpl27a.L
rpl27a.S
rpl28.L
rpl28.S
rpl30.L

rpl31.L
rpl31.S
rpl34.L
rpl34.S
rpl35a.S
rpl36.S
rpl36a.L
rpl37.L
rpl37.S
rpl39.L
rpl39.S
rpl4.L
rpl5.L
rpl7a.S
rpl8.L
rpl9.L
rplp1.L
rplp2.L
rps10.L
rps10.S
rps11.L
rps12.L
rps12.S
rps13.L
rps14.L
rps14.S
rps15.L
rps15.S
rps15a.S
rps16.L
rps16.S
rps17.L
rps17.S
rps19.S
rps20.L
rps21.S
rps23.S
rps25.L
rps25.S
rps26.L
rps27.L
rps27.S
rps28p9.S
rps3.S
rps3a.S
rps4x.L
rps4x.S
rps5.L

rps5.S
rps6.L
rps7.L
rps7.S
rps9
rpsa.L
rpsa.S
rrm1.S
rspo2.L
rspo2.S
sap18.L
scn1b.L
scrt2.S
sez6l2.L
sez6l2.S
sf3b1.S
sf3b5.S
sfxn5.L
shf.L
sim1.S
sim2.S
skor1.L
skor1.S
slc12a5.L
slc25a11.S
slc2a1.L
slc2a1.S
slc39a12.S
slc39a7.L
slc40a1.S
slc45a4.L
slc45a4.S
slc4a4.L
slc4a4.S
slc6a3.S
slc6a4.S
slc7a10.S
slit1.S
snrpa.S
snrpb.S
snrpc.S
snrpe.S
snrpf.L
snrpg.S
sntb1.L
sp5.L
sp5.S
sp8.S

sptbn1.L
sptbn1.S
sspo.L
stac2.S
stk32b.L
stra6.L
stra6.S
stt3a.L
stt3a.S
stx1b.L
stxbp1.L
stxbp1.S
sulf1.S
sulf2.L
sult4a1.L
sv2c.S
syn1.L
syn1.S
syn2.S
synj1.S
tal2.S
tcf4.L
tcp1.L
tf.L
tfap2e.L
tfap2e.S
tlcd3b.L
tmem145.S
tmem255a.L
tmem63b.S
tnfrsf21.L
trim9.L
txndc17.S
uba2.L
uba52.L
vamp2.S
vstm2l.L
vsx2.L
vwa5a.2.L
wasf1.L
wdr7.L
wls.L
wnt2b.L
wnt8b.S
zeb2.S
zic3.L
zic3.S
znf326.S

znf706l.L
znfx1.L

1

**KDBarhl1_A et B_vs_KDCT COMMON GENES pAdj<0.00**

**MOBarhl1_1_MOBarhl1_2**
LOC100126660
LOC108695757
LOC108695983
LOC108696020
LOC108696074
LOC108696239
LOC108696915
LOC108697444
LOC108697760
LOC108697808
LOC108697935
LOC108698084
LOC108698099
LOC108698154
LOC108698377
LOC108698757
LOC108699226
LOC108699273
LOC108699406
LOC108699444
LOC108699962
LOC108700185
LOC108700350
LOC108700443
LOC108700932
LOC108701018
LOC108701181
LOC108701569
LOC108701642
LOC108702162
LOC108702465
LOC108702731
LOC108702869
LOC108703011
LOC108703204
LOC108703406
LOC108703510
LOC108703551
LOC108704224
LOC108704587
LOC108704633
LOC108705545
LOC108706170
LOC108706667
LOC108706905

LOC108707429
LOC108707553
LOC108707564
LOC108707869
LOC108708151
LOC108708306
LOC108708507
LOC108708564
LOC108708999
LOC108709287
LOC108709337
LOC108709624
LOC108709778
LOC108709952
LOC108710030
LOC108710372
LOC108711439
LOC108711545
LOC108711739
LOC108712711
LOC108712865
LOC108713037
LOC108713150
LOC108713446
LOC108713567
LOC108713772
LOC108713861
LOC108714021
LOC108714603
LOC108715150
LOC108715931
LOC108717002
LOC108717189
LOC108717266
LOC108717850
LOC108718251
LOC108718312
LOC108718675
LOC108719610
LOC108719750
LOC108719848
LOC121393045
LOC121393901
LOC121394610
LOC121396584
LOC121398252
LOC121398856
LOC121399515

LOC121402164
LOC443585
MGC114789
MGC115323
MGC68655
MGC68699
MGC80700
XB5796436.L
XB5848002.S
XB5957215.S
abca3.L
abca3.S
acss1.S
actl6a.S
adgrl1.L
ado.L
agpat3.S
ahcy.L
aldoa.L
ank1.L
ank1.S
ankrd10.L
apcdd1.L
apcdd1.S
apex1.L
apex1.S
aplp1.S
app.L
app.S
armc6.L
arpp19.S
atp13a5l.2.L
atp1a1.L
atp1a1.S
atp1a2.S
atp1b1.L
atp1b2.S
atp2b1.L
atp2b1.S
atp2b4.L
atp2b4.S
atp6v1a.L
atp6v1a.S
barhl2.L
barhl2.S
bhlhe22.L
bsx.L
bsx.S

bzw1.S
c11orf87.S
ca7.L
cachd1.S
cacna1a.S
cadps.L
calb1.L
camk2g.L
camk2g.S
camk4.S
cbx3.S
cct5b
cdk5r1l.L
cdk5r1l.S
chrm2.L
cirbp.S
ckmt1b.L
clstn3.L
clstn3.S
cntnap1.L
col5a3.S
cox4i2.L
cpne4.L
crmp1.L
crmp1.S
crx.L
crx.S
csnk2b.L
ctnna2.L
ctnnb1.L
ctnnb1.S
cyp27c1.S
dach1.S
dach2.L
dbi.S
dctn1.L
ddx39a.S
dhcr24.L
dhrs3.L
dhx32.L
dlx5.S
dmrta2.L
dmrta2.S
dnajc5.S
dner.L
dner.S
dnm1l.L
dnm1l.S

dpysl2.L
dusp8.L
dynll1.S
ednrb2.L
eef1a2.L
efna2.S
efnb3.L
efr3b.L
eif3l.L
eif5a.L
emilin3.L
emx2.L
epha10.L
epha4.L
erh.L
fam219a.S
fau.L
fau.S
fgd1.S
fnbp1.L
foxq1.S
frmpd1.L
fry.L
fst.L
galnt14.S
gipr.L
glipr2.S
glra1.S
gnat2.S
gpc1.L
gpc1.S
gpc5.L
gpt.S
grid2ip.L
grin1.L
gucy2d.L
gucy2d.S
h2az1.L
h2az1.S
h2az2.S
h3-3b.S
hectd1.L
helt.L
hes5.1.L
hes5.1.S
hes5.2.L
hes5.3.L
hes5.3.S

hes5.4.L
hmga2.L
hmgb2.L
hmgb2.S
hmgb3.L
hmgn1.L
hmgn1.S
hmgn2.S
hnrnpab.L
hnrnpdl.L
hnrnpk.S
hsbp1.L
hsf2.2.L
hspa12a.S
igfbpl1.L
igfbpl1.S
iqsec3.S
itga10.L
kalrn.L
kars1.L
kcnc1.S
kcne3.S
kcnh4.L
kcnh4.S
kcnj10.L
kctd15.L
khdrbs1.S
kiaa1549l.L
kif1a.L
ldha.L
ldha.S
lef1.L
lef1.S
lhx2.L
lhx2.S
lhx9.L
lhx9.S
lman1l.L
lmo3.L
lmo3.S
lmx1a.L
lrit1.S
lrp1.L
lrp1.S
lrrn3.S
lrwd1.L
ltbp4.L
mab21l1.S

madd.S
maf.L
mafa.L
magoh.L
map1b.S
map4.L
map7d1.L
map7d2.S
mapk8ip2.L
mapk8ip2.S
mapt.L
mapt.S
mast1.L
matr3.S
mcm4.L
mcm7.L
megf6.L
meis2.L
mmp16.L
mn1.L
msrb2.S
mtss2.L
mvd.L
myg1.S
naca.L
nap1l1.L
nasp.L
ncor2.S
nefh.L
nefh.S
nefl.L
nefl.S
nefm.S
nfia.S
nfib.L
nif.L
nif.S
nkx2-2.L
nkx2-4.S
nkx6-1.L
nlgn2.S
nme2.S
nmu.S
not.L
not.S
npm1.L
npm1.S
npy.L

nr2e1.L
nr2e1.S
nr2f1.L
nsa2.L
ntn1.L
ntn3.S
ntn5.L
odc1.L
odc1.S
ogdh.S
otp.L
otp.S
otx1.L
otx2.L
otx2.S
pa2g4.L
pafah1b1.L
paqr6.L
paqr6.S
pc.1.L
pcdh10.L
pcdh7.L
pcdh7.S
pcna.S
pfkp.L
pfkp.S
phr.L
phyhipl.S
pitpnc1.S
pitx2.S
pitx3.L
pitx3.S
plcb4.S
plxna1.L
pmp2.S
pomp.S
pou4f4.L
pou4f4.S
prdm12.L
proca1.L
prpf19.L
prrt1.S
prrt2.L
psma3.L
psma4.L
psma5.S
psma6.S
psmb1.S

psmb7.S
psmc2.S
psmc3.L
ptchd4.L
ptgds.S
ptmap12.L
ptmap12.S
ptn.S
pygm.L
pygm.S
r3hdm1.L
rab11fip5.L
rab6b.L
rack1.L
rbp1.S
rdh5.L
rgma.L
rgr.L
rgr.S
rlbp1.S
rngtt.S
rora.L
rpe65.L
rpl10a.S
rpl11.L
rpl11.S
rpl12.L
rpl13a.L
rpl14.S
rpl15.L
rpl15.S
rpl17.L
rpl18.L
rpl18.S
rpl18a.L
rpl18a.S
rpl21.L
rpl22.L
rpl23a.L
rpl23a.S
rpl24.S
rpl26.L
rpl26.S
rpl27a.L
rpl27a.S
rpl28.L
rpl28.S
rpl30.L

rpl31.L
rpl31.S
rpl34.L
rpl34.S
rpl35a.S
rpl36.S
rpl36a.L
rpl37.L
rpl37.S
rpl39.L
rpl39.S
rpl4.L
rpl5.L
rpl7a.S
rpl8.L
rpl9.L
rplp1.L
rplp2.L
rps10.L
rps10.S
rps11.L
rps12.L
rps12.S
rps13.L
rps14.L
rps14.S
rps15.L
rps15.S
rps15a.S
rps16.L
rps16.S
rps17.L
rps17.S
rps19.S
rps20.L
rps21.S
rps23.S
rps25.L
rps25.S
rps26.L
rps27.L
rps27.S
rps28p9.S
rps3.S
rps3a.S
rps4x.L
rps4x.S
rps5.L

rps5.S
rps6.L
rps7.L
rps7.S
rps9
rpsa.L
rpsa.S
rrm1.S
rspo2.L
rspo2.S
sap18.L
scn1b.L
scrt2.S
sez6l2.L
sez6l2.S
sf3b1.S
sf3b5.S
sfxn5.L
shf.L
sim1.S
sim2.S
skor1.L
skor1.S
slc12a5.L
slc25a11.S
slc2a1.L
slc2a1.S
slc39a12.S
slc39a7.L
slc40a1.S
slc45a4.L
slc45a4.S
slc4a4.L
slc4a4.S
slc6a3.S
slc6a4.S
slc7a10.S
slit1.S
snrpa.S
snrpb.S
snrpc.S
snrpe.S
snrpf.L
snrpg.S
sntb1.L
sp5.L
sp5.S
sp8.S

sptbn1.L
sptbn1.S
sspo.L
stac2.S
stk32b.L
stra6.L
stra6.S
stt3a.L
stt3a.S
stx1b.L
stxbp1.L
stxbp1.S
sulf1.S
sulf2.L
sult4a1.L
sv2c.S
syn1.L
syn1.S
syn2.S
synj1.S
tal2.S
tcf4.L
tcp1.L
tf.L
tfap2e.L
tfap2e.S
tlcd3b.L
tmem145.S
tmem255a.L
tmem63b.S
tnfrsf21.L
trim9.L
txndc17.S
uba2.L
uba52.L
vamp2.S
vstm2l.L
vsx2.L
vwa5a.2.L
wasf1.L
wdr7.L
wls.L
wnt2b.L
wnt8b.S
zeb2.S
zic3.L
zic3.S
znf326.S

znf706l.L
znfx1.L

1
